# Compensatory sequence variation between *trans*-species small RNAs and their target sites

**DOI:** 10.1101/675900

**Authors:** Nathan R. Johnson, Claude W. dePamphilis, Michael J. Axtell

## Abstract

*Trans*-species small regulatory RNAs (sRNAs) are delivered to host plants from diverse pathogens and parasites and can target host mRNAs. How *trans*-species sRNAs can be effective on diverse hosts has been unclear. Multiple species of the parasitic plant *Cuscuta* produce *trans*-species sRNAs that collectively target many host mRNAs. Confirmed target sites are nearly always in highly conserved, protein-coding regions of host mRNAs. *Cuscuta trans*-species sRNAs can be grouped into superfamilies that have variation in a three-nucleotide period. These variants compensate for synonymous-site variation in host mRNAs. By targeting host mRNAs at highly conserved protein-coding sites, and simultaneously expressing multiple variants to cover synonymous-site variation, *Cuscuta trans*-species sRNAs may be able to successfully target homologous mRNAs from diverse hosts.

**One Sentence Summary:** The parasitic plant *Cuscuta* produces a diverse set of sRNAs that compensate for sequence variation in mRNA targets in diverse hosts.

## Main Text

Small regulatory RNAs (sRNAs) produced in one organism can sometimes function to silence mRNAs in another organism. These *trans*-species sRNAs seem especially prominent in plant/pathogen and plant/parasite interactions. Fungal plant pathogens produce sRNAs with complementarity to host mRNAs (*1*) and host plants produce *trans*-species sRNAs that silence mRNAs in both pathogenic fungi (*2, 3*) and oomycetes (*4*). The parasitic plant *Cuscuta campestris* produces *trans*-species microRNAs (miRNAs) which silence host plant mRNAs (*5*). Silencing by plant *trans*-species sRNAs relies on extensive complementarity between the sRNA and target mRNA (*6*). *Trans*-species silencing is expected to benefit the source organism while being detrimental to the target organism in parasitic/pathogenic relationships. This implies that target sites are not under purifying selection to maintain complementarity *to trans*-species sRNAs. How could such a system be stable over evolutionary time and/or be useful against multiple species? One suggestion is a “shotgun” strategy, in which a very diverse set of *trans*- species sRNAs is deployed to hit target mRNAs randomly. The plant response to *Phytophthora*, may make use of this strategy (*4*). However, the fact that the *trans*-species sRNAs delivered to hosts from the parasitic plant *C. campestris* are miRNAs (*5*) argues against the shotgun hypothesis in this case. MiRNAs are defined by the precise excision of a single mature, functional small RNA (*7*), which implies selection for the miRNA to target a particular target sequence. We examined *Cuscuta trans*-species sRNAs and their targets in detail to shed light on how this system may be evolutionarily stable and robust against diverse hosts.

We analyzed sRNA expression from four *Cuscuta* species (Fig. 1A). Specimens from two or three distinct populations of *C. pentagona* and *C. gronovii*, respectively, were included, making a total of seven separate sRNA expression studies (identified with acronyms for brevity; Fig. 1A, S1; Table S1). All four *Cuscuta* species are generalists with documented hosts spanning multiple plant families (Fig. S2). RNA samples (three biological replicates each) from host-parasite interfaces and parasite stems growing on the host *Arabidopsis thaliana* were obtained and used for sRNA sequencing (Fig. 1B). Libraries were condensed to highly expressed sRNA variants and filtered to remove any sRNAs that came from the host (Fig. S3). Differential expression analysis revealed several hundred *Cuscuta* sRNAs in each experiment that were significantly up-regulated in the interface tissue relative to parasite stems (FDR < 0.1); we dubbed these haustorially-induced (HI) sRNAs (Fig. 1A; Data S1). HI-sRNAs are mostly 21 or 22 nucleotides long (Fig. 1A), sizes consistent with either miRNAs or short interfering RNAs (siRNAs). Distinguishing miRNAs from siRNAs requires a genome assembly (*7*), a criterion met so far for only one of the four species (*C. campestris*) included in this study *(5, 8)*. The majority of *C. campestris* HI-sRNAs (295/408) come from *MIRNA* hairpins. *C. campestris-*derived HI-sRNAs were recovered from 40 of the 42 novel *MIRNA* loci described by Shahid et al. (*5*), including representatives from all five miRNA families previously demonstrated to target host mRNAs.

**Fig. 1.**
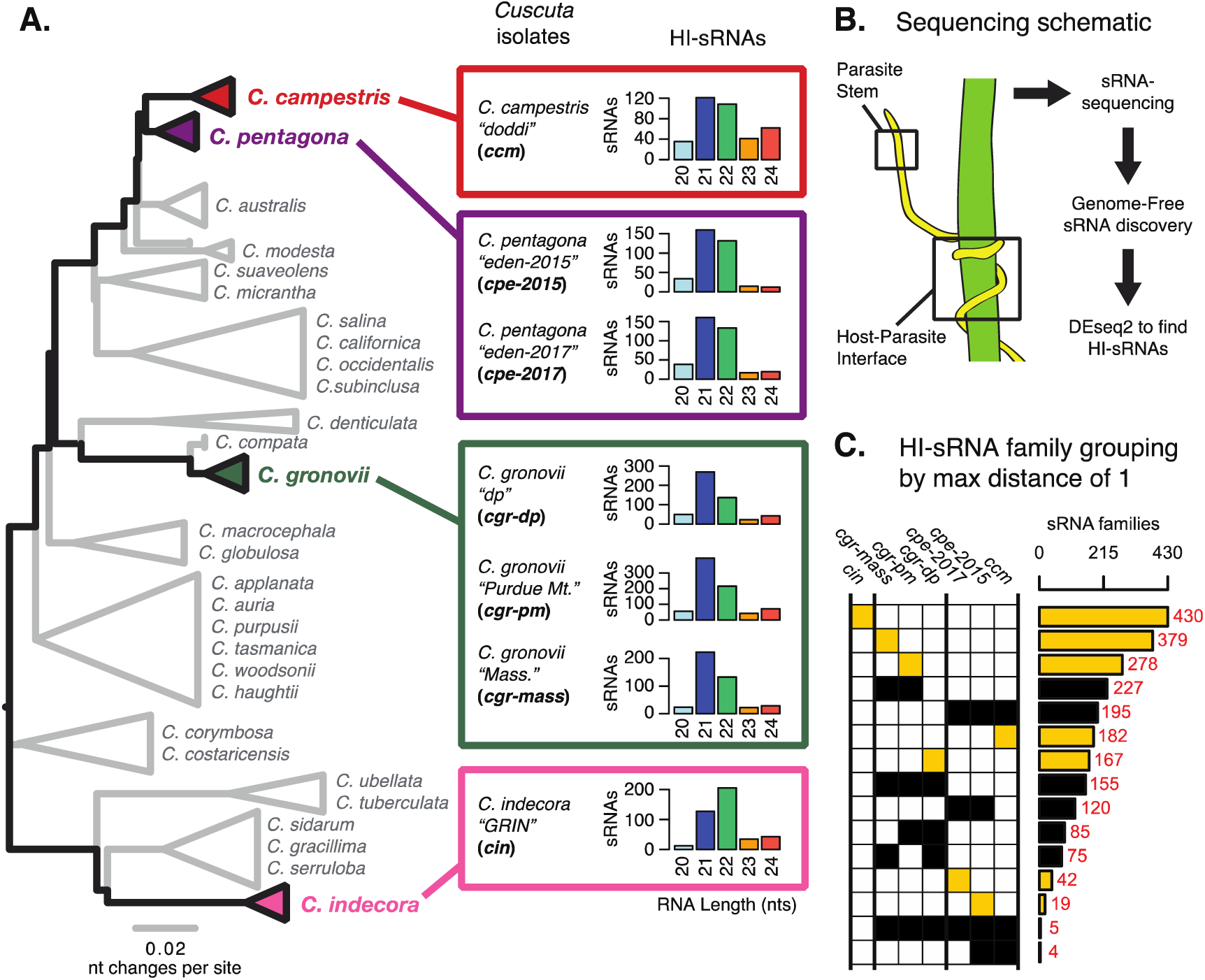
Haustorium-induced small RNAs (HI-sRNAs) are present in multiple *Cuscuta* species. (**A**) Phylogeny of select *Cuscuta* species. Size distribution of HI sRNAs for each sequenced isolate and acronyms are shown. (**B**) Sampling and sequencing schematic to discern HI-sRNAs. (**C**) HI-sRNA family counts and membership for each isolate, showing only the top 15 groups. Families were grouped strictly using a maximum edit distance of one nucleotide. Yellow indicates families present in a single isolate.

We next examined conservation of HI-sRNA accumulation between isolates and species. Some canonical plant miRNAs are highly conserved, with several ubiquitous families found in multiple plant orders, or even broadly in all land plants (*9*). Surprisingly, when using a strict cutoff (maximum edit distance of 1) we found that the majority of HI-sRNAs were not observed in more than one species (Fig. 1C). In many cases HI-sRNAs were unique to single isolates of a single species (Fig. 1C). This result implies that HI-sRNAs could be rapidly differentiating in expression, sequence, or both within these *Cuscuta* species.

Our previous work showed that *C. campestris* HI-sRNAs can target host mRNAs. We thus looked for evidence of interactions between our broader sets of HI-sRNAs with host (*A. thaliana*) mRNAs using two complementary methods: secondary siRNA accumulation (*5*) and degradome analysis (*10*). Secondary siRNAs can accumulate from mRNAs as a result of an initial miRNA- or siRNA-directed targeting event, especially when the initiating sRNA is 22 nucleotides long (*11*). Degradome analysis made use of the NanoPARE method (*12*), which recovers 5’ ends of both capped and uncapped mRNAs. NanoPARE libraries were made from just one isolate from each of the four *Cuscuta* species, and comprised three biological replicates from the host portion of the interface (Table S1). *A. thaliana xrn4* mutants were used as hosts for these experiments because they over-accumulate 5’ remnants of sRNA-mediated mRNA cleavage (*12, 13*). The *MPK3* mRNA is an example with both degradome and secondary siRNA evidence of targeting by a *C. gronovii* HI-sRNA (Fig. 2A-D). Altogether these two analyses yielded a set of 54 *A. thaliana* mRNAs targeted by *Cuscuta* HI-sRNAs confirmed by a single method and seven more confirmed by both (Fig. 2E, S4A, S5). Based on RNA-seq analysis, accumulation of confirmed target mRNAs is generally down-regulated in parasitized host stems (Fig. S6). This greatly expands on the set of six mRNAs previously identified as host targets of *C. campestris* miRNAs (*5*), and demonstrates that *trans*-species sRNAs are used by multiple *Cuscuta* species. Repeating this analysis to examine possible self-targeting of *C. campestris* mRNAs by HI-sRNAs found only four confirmed targeting interactions, an indication that HI-sRNAs may largely function in *trans* in the host (Fig. S7). One explanation for the lack of endogenous targeting may be purifying selection in the parasite, as target predictions show that *C. campestris* homologs of targeted *A. thaliana* mRNAs invariably have lower complementarity to HI-sRNAs (Fig. S7).

**Fig. 2.**
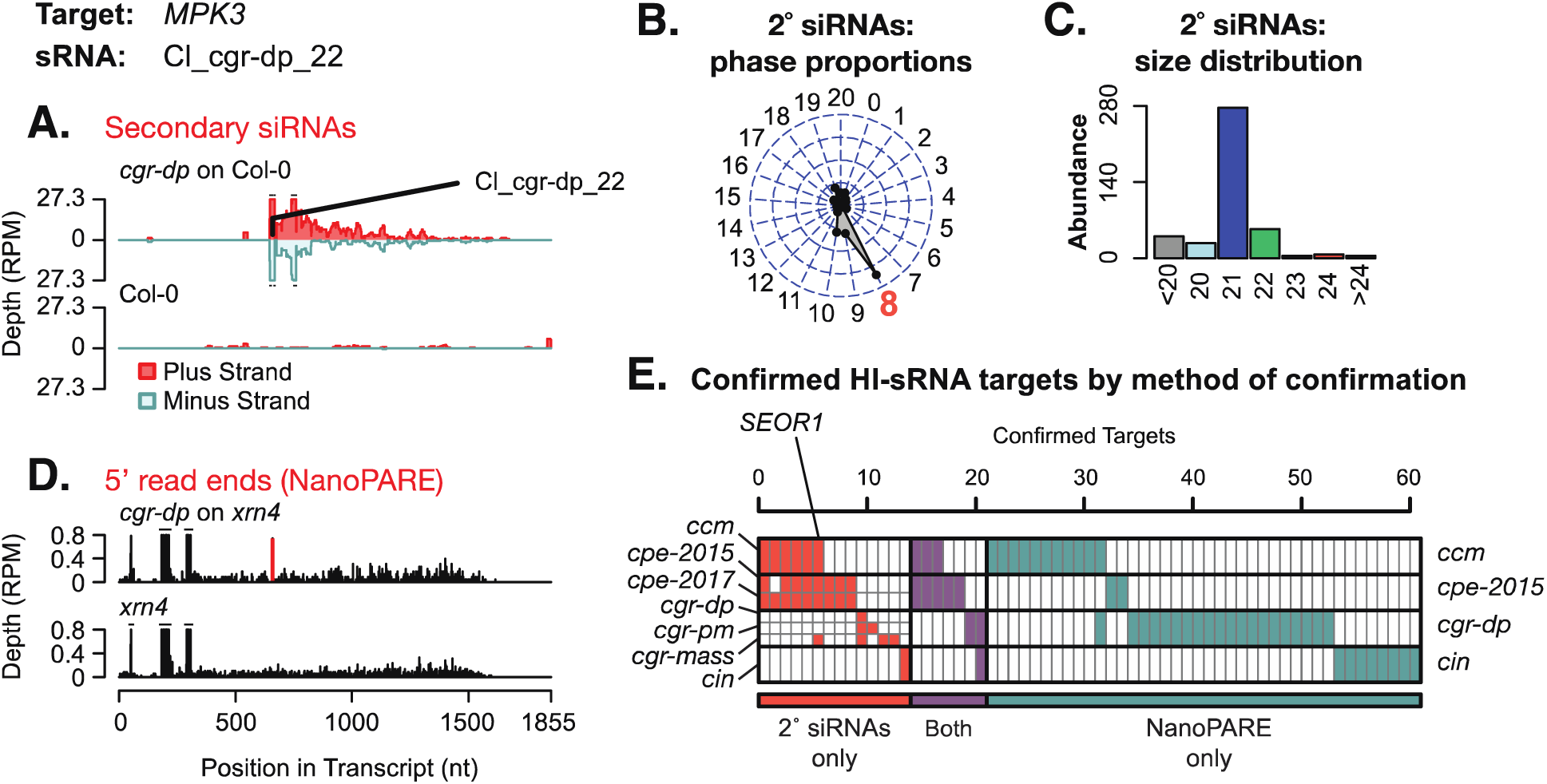
Host targets of *Cuscuta* HI-sRNAs. (**A**) Secondary siRNA accumulation from *A. thaliana MPK3*. (**B**) Phasing analysis of secondary siRNAs from *MPK3*. Expected phase for cut-site shown in red. (**C**) Size distribution of *MPK3* secondary siRNAs. **(D)** Frequency of 5’ ends from the *MPK3* mRNA, with the predicted HI-sRNA cut site shown in red. **(E)** Host mRNAs with confirmed targeting by a *Cuscuta* HI-sRNA. Full details in Figs. S4-S5.

Some mRNAs were confirmed to be targeted by HI-sRNAs from multiple species, with the most frequent interaction being with *SEOR1. SEOR1* encodes a phloem protein that acts to reduce sap loss after wounding (*14*). *C. campestris* growth is enhanced when *A. thaliana seor1* mutants are used as hosts (*5*). However, the majority of mRNAs confirmed as targets are unique to a single *Cuscuta* species or isolate. A possible explanation could be that *Cuscuta trans*-species sRNAs function to regulate similar host processes, while not necessarily the same target mRNAs. Additionally, our current analysis is likely to have missed many targets, both due to lack of sensitivity of our methods (secondary siRNA accumulation and/or degradome analysis both can miss true targets), and because *A. thaliana* is unlikely to be a major host of *Cuscuta* in nature.

Numerous target mRNAs are known to be involved in the same processes, both on a gene ontology level (Fig. S8) and when manually examining known pathways. Genes involved in auxin signaling repeatedly appear, including the previously identified targets *TIR1, AFB2*, and *AFB3* (*5*) and new targets *PXY* (*15*) and *ARK2 (16)* with a proposed role in auxin response. Auxin signaling is involved in many processes in the plant, and is potentially connected to defense against *Cuscuta* through its role in glucosinolate production (*17, 18*). Phloem protein mRNAs are targeted, adding *OPS (19)* to the previously identified *SEOR1*. A receptor-like kinase gene (*CuRe1*) from tomato is a resistance gene that prevents *Cuscuta reflexa* infestation (*20*). Receptor-like kinases and kinases in general are well represented in the set of HI-sRNA targets, including several that are involved in defense responses. This includes the well-known defense regulator *MPK3 (21)* and previously discovered *BIK1 (22)*. Another pathway found to be targeted is brassinosteroid (BR) signaling, with targets *BRI1 (23), MAPKKK5* (*24*), *PICKLE* (*25*). BR has a clear role in defense, with connections to both *BIK1* and *MPK3* (*26*). An overall theme of targeting host immunity and vascular system function emerges from this set of confirmed targets.

Plant miRNAs that initially seem unrelated based on divergent sequences can sometimes be grouped into superfamilies (*27*). To identify potential superfamilies among *Cuscuta* HI-sRNAs, we clustered them with a cutoff of five substitutions and barring indels (Fig. S9). This clustering strategy gives low rates of grouping by random chance (Fig. S10). Many superfamilies of *Cuscuta* HI-sRNAs were found, with a substantial portion of them shared between species and isolates (Fig. 3A). 19 superfamilies were shared between all isolates except *C. indecora*, and another 20 superfamilies were present in at least one isolate each of *C. campestris, C. pentagona*, and *C. gronovii*. Leveraging the prior *C. campestris* miRNA annotations, we can extrapolate that 158 out of 332 superfamilies which contain *C. campestris* HI-sRNAs are likely miRNAs. Furthermore, we extrapolate that of the superfamilies present in *C. campestris* with proven target relationships, 22 out of 23 are likely to be miRNAs (Fig. S4B).

**Fig. 3.**
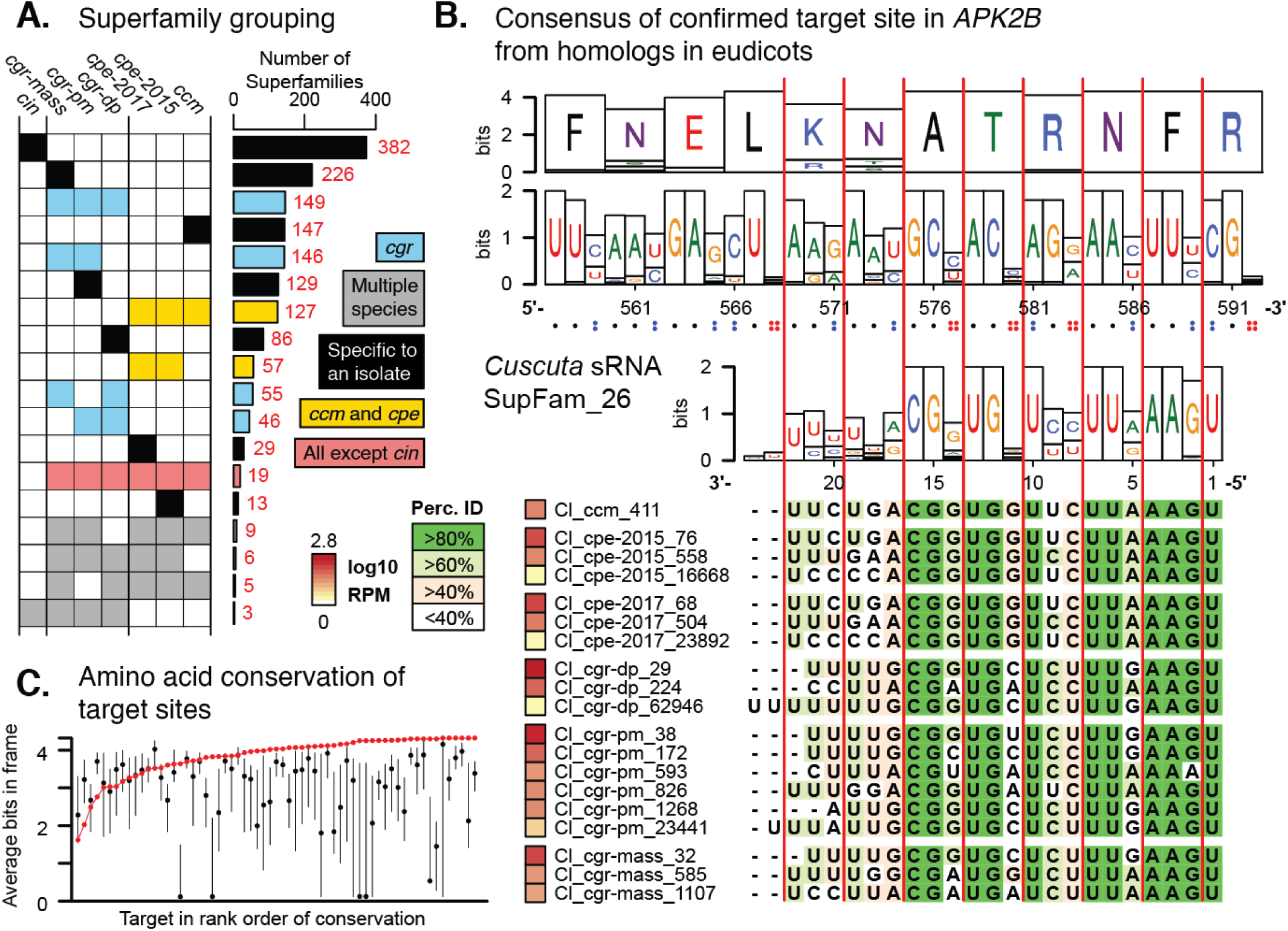
*Cuscuta* HI-sRNAs form superfamilies that co-vary with target sites across eudicots. (**A**) sRNA superfamily count and membership for each *Cuscuta* isolate. Colors indicate general groupings of superfamilies. (**B**) An example HI-sRNA superfamily aligned to target sites from homologs in 36 eudicot genomes. Nucleotide and amino acid Shannon entropy from the alignments are shown as bits. Vertical red lines indicate the frame. Dots indicate the number of possible synonymous nucleotides at each codon. 17 additional examples in Fig. S11. (**C**) Average conservation of target sites from homologs. Confirmed target site shown (red point), with all other possible sites shown by 25-75% quartiles (black line) and median (black point).

HI-sRNAs within a superfamily vary at several positions both between and within species (Fig. 3B). In many cases variation within superfamilies occurred in a three-nucleotide period (Fig. 3B, S11). This pattern led us to investigate nucleotide variation in corresponding target sites among possible hosts. All four *Cuscuta* species in this study are generalists that parasitize eudicot hosts, so we aligned homologous target mRNAs from 36 eudicot species (Table S3). Analysis of translated target site conservation shows that HI-sRNAs target highly-conserved protein-coding positions (Fig 3C). Positional variations in HI-sRNA superfamilies precisely correspond to variable positions in homologous target sequences (Fig. 3B). This variation is frequently apparent at synonymous sites, accounting for the three-nucleotide periodicity of superfamily variation. Modeling correlation of positional variation between HI-sRNA superfamilies and eudicot target sites found 18 significant examples of this type of co-variation (Fig. S11, S4C). Importantly, HI-sRNA superfamily variation occurs within single *Cuscuta* species (Fig. 3B, S11), such that multiple HI-sRNA variants are commonly deployed by a given parasite during infestation. By targeting conserved sites, and making several HI-sRNA variants that collectively cover many/most possible synonymous target variants, *Cuscuta* may ensure successful targeting across a wide range of hosts.

HI-sRNA superfamily diversity could also enable repression of multiple mRNAs with homologous target-sites within a single host. We examined target predictions within *A. thaliana* and found ten examples of gene family-specific motifs potentially targeted by *Cuscuta* HI-sRNA superfamilies (Fig. S12). These include a HI-sRNA superfamily predicted to target the mRNA region encoding the eponymous WRKY motifs within the well-known family of defense-related transcription factors (*28*) (Fig. S12). Most of the possible targets, including the *WRKY*s, predicted in this analysis have not yet been experimentally confirmed by secondary siRNA accumulation or degradome analysis. However, false negatives are common with these methods of confirmation. The striking patterns of sequence covariation between the HI-sRNA superfamilies and their possible target mRNA families make a strong case for the reality of these interactions.

We conclude that multiple *Cuscuta* species use *trans*-species HI-sRNAs to target a substantial number of host mRNAs. Many if not most of these HI-siRNAs are likely to be miRNAs. Host genes involved in pathogen defense, hormone signaling, and vascular system function are common targets of *Cuscuta trans*-species HI-sRNAs. *Cuscuta trans*-species HI-sRNAs nearly always interact with highly conserved target sites within the coding sequences of host mRNAs. HI-sRNAs can often be grouped into superfamilies that have nucleotide diversity that corresponds with target site variation primarily at synonymous sites. It seems likely that host target sites are under purifying selection because they code for critical amino acids that have little variation even among distantly related eudicots. By targeting these constrained sites, and deploying an array of variants that cover most possible permutations of synonymous site variation, *Cuscuta* HI-sRNAs are likely to be robust *against* the development of host resistance. This is also a suitable strategy for a generalist parasite that interacts with diverse hosts. The strategy used by *Cuscuta* provides a novel paradigm for the molecular evolution of *trans*-species sRNA targeting during parasitism.

## Acknowledgments

We thank T. Phifer for help generating preliminary data; L.S. Berghardt for greenhouse support; C. Praul for next-generation sequencing support; J. Westwood (Virginia Tech) for providing multiple *Cuscuta* seeds and thoughtful insight into the work; C. Depew for informing us about *C. gronovii* locations in the wild; and M. Schon and M. Nodine for early access to the NanoPARE method.

## Author contributions

NRJ performed sequencing experiments and analysis. NRJ, CWD, and MJA conceived and planned experiments. NRJ and MJA wrote the manuscript.

## Data and materials availability

Supplement contains additional data and libraries have been deposited to the NCBI sequence read archive under the BioProject: PRJNA543296.

## Supplementary Materials

Materials and Methods

Figures S1-S12

Tables S1-S3

External Data S1-S2

References (*29-46*)

