## Supplemental Materials for "Compensatory sequence variation between *trans*-species small RNAs and their target sites"

**This PDF file includes:**

Materials and Methods
Figs. S1 to S12
Tables S1 to S3
Captions for Data S1 to S2

**Other Supplementary Materials for this manuscript include the following:**

Data S1 to S2

#### Materials and Methods

Seed sources

*Cuscuta campestris* (isolate doddi) was originally acquired from a tomato field in California, followed by several generations of selfing in the Westwood laboratory and provided to us as a gift. *C. gronovii* (isolate DP) was a gift C. dePamphilis, from an unknown source. *C. gronovii* (isolate mass) was collected in Massachusetts and provided as a gift by J. Westwood. *C. gronovii* (isolate PM) was collected from a road-side farm near State College, PA by M. Axtell (Coordinates: 40.866 N, 77.888 W). *C. pentagona* (isolates eden-2015 and -2017) were purchased from ebay seller eden_wilds in 2018, both collected from locations in upstate New York. *C. indecora* (isolate *cin*/GRIN) was acquired from the U.S. national plant germplasm system (www.ars-grin.gov) under the accession: PI 675068.

Genotyping *Cuscuta*

*Cuscuta* seed were scarified by nicking with a razor blade under dissecting microscope and germinated on wet paper towel under growth lighting at ~28°C, harvesting seedlings after 3-5 days of growth for DNA extraction. DNA was extracted using Edwards method (*29*). 2 uL of template was used in a 20 uL PCR reaction using *Taq* polymerase using 0.5 uM final concentration of each primer (Forward: MJA1, Reverse: MJA2, Table S2) (*30*). PCR was performed for 30 cycles and enzymatically cleaned-up using 0.5 uL of Exo1 (NEB) and 1 uL of Antarctic Phosphatase (NEB) with 5 uL PCR product, followed by incubation (15 minutes at 37°C) and inactivation (15 minutes at 80°C). Sanger sequencing was performed by the Penn State genomics core using MJA1 primer. Sequences were trimmed of low quality bases and aligned using muscle (*31*) to published TrnL-F sequences (*32*, *33*). Nucleotide phylogeny was constructed using MEGA7 (*34*) with a maximum likelihood method and 500 bootstraps (Data S2).

Growth conditions

Host *A. thaliana* (Col-0 and *xrn4*) was sown on wet potting medium, followed by with 3 days of stratification at 4°C. Plants were placed into long day (16 day/8 night) growth conditions at ~23°C under cool-white-fluorescent lamps. Hosts were allowed to grow to maturity (4-5 weeks old), ready for attachment when first inflorescences were longer than 5 cm. *Cuscuta* seeds were scarified and germinated as above. Seedlings were ready for attachment once completely emerged from their seed husk and roughly ~2 cm in length, 3-5 days depending on the species. Seedlings were placed in soil next to the primary bolts of host plants. House-built far-red supplementary LED lighting was used to induce attachment under fluorescent lights, allowing 4-5 days for attachment of the parasite. Once attached, parasitized hosts were removed from far-red lighting to prevent secondary attachment. For *C. indecora*, experimental attachments came from tendrils from a previously established *C. indecora* colony. 5 cm tendril tips were cut off of the colony and affixed to primary bolts of host plants with scotch tape. Plants were allowed to grow for 10 days after attachment followed by tissue harvest.

Tissue collection and RNA extraction

All tissues were collected by the following methods and immediately submerged in liquid nitrogen to preserve RNA stability. Guide to tissues gathered is found in Table S1. Interface (IN) tissue was collected by taking both the host and parasite portions of the interface, trimming away any stems above and below the connection. Parasite stem (PS) was harvested ~4 cm above the interface, approximately 4 cm long each. In NanoPARE experiment using *xrn4* as a host, we collected only host interface (HIN); similar method to interface collection, except removing any parasite tissue which can be pulled away. Control stem (CS) tissues were harvested from non-parasitized *A. thaliana*, collecting stems from the same region where *Cuscuta* would have been attached. 1-3 tissues were pooled for each biological replicate. RNA was extracted by grinding tissue in a liquid nitrogen cooled mortar, with Tri-reagent (Sigma) added while still cold. Tri-reagent extraction was performed as per the manufacturer's suggestions with a second sodium-acetate–ethanol precipitation and wash step.

Sequencing library preparation

All small-RNA-seq libraries were prepared using a protocol based on the NEBnext small-RNA library kit (NEB), described as follows. **Step 1)** 3' SR Adaptor (NJ410) was pre-adenylated using 5' adenylation kit (NEB) as per manufacturer's instructions. **Step 2)** 500 ng of total RNA, 1 µL adenylated adapter (5 µM) and water to 5.25 µL were denatured for 2 minutes at 70°C and immediately moved to ice. Entire reaction was combined with premixed 100 u RNA Ligase 2, truncated KQ (NEB), 1 µL 10x T4 RNA reaction buffer, 10 u RNAse inhibitor and 3 µL 50% PEG8000, to a total volume of 10 µL and incubated 1 hour at 25°C. **Step 3)** Primer hybridization was performed adding 0.5 µL SR RT primer (NJ391, 10 µM) and 2.25 µL water to the prior reaction and incubated as follows: 5 minutes at 75°C, 15 minutes at 37°C, 15 minutes at 25°C, and holding at 4°C. **Step 4)** 5' SR RNA adaptor (NJ411) was diluted to 10 µM and denatured for 2 minutes at 70°C, moved to ice and used immediately for ligation. Ligation was performed combining the prior reaction with 0.5 µL denatured adapter (NJ411), 5 u RNA Ligase 1 (NEB), 0.25 µL 10x RNA ligase buffer, 10 u RNAse inhibitor, 0.5 µL ATP (10 mM), and water to 15 uL and incubated for 1 hour at 25°C. **Step 5)** Reverse transcription was performed immediately following ligation, combining the prior reaction with 100 u Protoscript II reverse transcriptase (NEB), 4.5 µL 5x first strand synthesis buffer, 1 µL dNTPs (10 uM), 1.5 µL DTT (0.1 M), and 10 u RNAse inhibitor equaling 23 µL in total volume and incubated for 1 hour at 50°C followed by heat-killing for 15 minutes at 70°C. **Step 6)** Library amplification was performed, combining 5 µL of cDNA with 25 µL LongAmp Taq 2x master mix (NEB), 1.25 µL SR primer (NJ412, 10 µM), 1.25 µL barcode primer ("NEB" primers, 10 µM), and water to 50 µL total volume. Reaction was performed as follows: 30 seconds initial denature at 94°C, 15 cycles of 15 seconds at 94°C, 30 seconds at 62°C, and 15 seconds at 70°C, followed by final extension of 5 minutes at 70°C. Reactions were purified and size selected for sRNAs 15-40 nt in length by PAGE. Extracted bands were quantified by qPCR and quality-controlled by high-sensitivity DNA chip (Agilent). Sequencing was performed on a NextSeq550 (Illumina) with the high-output kit (75 nt, single-end, single barcode) by the Penn State genomics core. Sequencing libraries were de-multiplexed and adaptor trimmed using cutadapt (*35*) (cutadapt -a AGATCGGAAGA -m 15 -j 8 -o *output.fq input.fq*).

NanoPARE and mRNA-seq libraries were prepared using the protocol described in (*12*), with the following details: NanoPARE and mRNA-seq were performed on interfaces (IN) of four isolates (*ccm*, *cpe-2015*, *cgr-dp*, and *cin*) and control stems (CS), grown on Col-0 *A. thaliana.* NanoPare was also performed on host interfaces (HIN) of the same isolates and control stem grown on *xrn4* mutant Col-0 *A. thaliana*. The Nextera DNA flex kit (Illumina) was used for tagmentation of 110 ng pre-amplified PCR product. Libraries were amplified using different barcoded i7 and i5 primer sets, described in Table S2), allowing for either amplification of 5' ends (NanoPARE) or all tagged entities (mRNA-seq). The sequencing of NanoPARE data made use of custom read 1 sequencing and i5 index sequencing primers (NJ395 and NJ416, reverse complements of each other), which sequence out from the template switching oligo adapter. Sequencing was performed on a NextSeq550 (Illumina) with the high-output kit (75 nt, single-end, double barcoded) by the Penn State genomics core. Sequencing libraries were de-multiplexed and NanoPARE libraries were trimmed using an in-house script to remove any residual untemplated 5' nucleotides caused by reverse transcription of the template-switching oligo.

Genome-free sRNA discovery

Genome-free sRNA discovery was performed using a set of in-house scripts, corresponding to the following pipeline (Fig. S3). Reads were filtered by size, retaining lengths of 20 to 24 nt, and condensed to unique sequences with a count of abundances for each tissue. For each *Cuscuta* species, unique reads were further condensed by sequence similarity to their most abundant variants. This process found similar variants for sRNAs in rank order of abundance, clustering sRNAs with a levenshtein edit distance of 2 or less. Reads which do not cluster to a variant with abundance of 0.5 reads per million (RPM) or higher are discarded. Most abundant sRNA variants of each cluster are reported, with the abundance as the combined abundance of all clustered reads. Host sRNAs were then filtered, removing an sRNA if it met one of the following criteria: **1)** it is closely similar to an annotated miRNA; **2)** it aligns perfectly to the *A. thaliana* genome or transcriptome (*36*); **3)** it is present in non-parasitized *A. thaliana* control libraries at an RPM greater than 1/100 its expression in parasite libraries. Differential expression analysis was then performed with DEseq2 (*37*) to identify sRNAs up-regulated in the interface tissue relative to the parasite stem, using the command the "results" command with a false-discovery rate of 0.1 (Benjamini–Hochberg correction). This pipeline resulted in our list of HI-sRNAs.

Superfamilies were constructed using an in-house script that corresponds to the following pipeline. All by all comparisons of HI-sRNA sequences were performed, measuring modified hamming distance (Fig. S9), and sequences with a distance of 5 or less were clustered together, ordered by overall size of the superfamily. To test this distance cutoff, HI-sRNA sequences were shuffled using UShuffle (*38*), set to retain di-nucleotide structure (10 random replicates) (Fig. S10).

Target confirmation

Target prediction of HI-sRNAs was performed using the script GSTAr.pl (https://github.com/MikeAxtell/GSTAr) under default settings, using HI-sRNA from a given isolate as the query and *A. thaliana* ARAPORT11 transcriptome (*36*) as the subject.

To find secondary siRNAs produced from targeting of *A. thaliana* genes, sRNA-annotation was performed on the *A. thaliana* genome (*39*), using ShortStack (*40*) (https://github.com/MikeAxtell/ShortStack) with gene locations from the ARAPORT11 annotation (*36*) as the basis for sRNA loci. Differential expression analysis was performed with DEseq2 (*37*) to identify loci up-regulated in the interface (IN) relative to control stem (CS), using the "results" command with a false-discovery rate of 0.1 (Benjamini–Hochberg correction). Up-regulated loci were then filtered to retain loci which met the following criteria: **1)** have a strong predicted HI-sRNA target site (complementarity score (*41*) 6 or less); **2)** are unstranded; **3)** have a predominant sRNA length of 21/22 nt; **4)** have a minimum depth of 20 reads. Plots of loci with predicted targets, radar plots of sRNA phasing, and length distribution plots to find examples where HI-sRNAs are clearly causative in the locus.

To confirm targeting of *A. thaliana* genes using degradome data, host stem of interface (HIN) NanoPARE libraries were aligned to the ARAPORT11 (*36*) transcriptome using bowtie (*42*) (bowtie -p 8 -f -v 3 -S -a). Using an in-house script, frequency at 5' positions of alignments were intersected with predicted HI-sRNA target sites, retaining interactions which met the following criteria: **1)** have a strong prediction score (complementarity score (*41*) 6 or less); **2)** the target site is greater than 100 nt from the start of the transcript (to avoid miscalls with the transcriptional start site); **3)** the target is not in an organellar genome; **4)** the target peak is greater than the median peak depth in the gene; **5)** the target peak is found in all 3 replicates; **6)** the target peak is at least 10 fold higher than detected in control stems. Candidates were then examined by eye, filtering out hits with low expression compared to the rest of the gene, hits which appear to be in the transcriptional start site, or hits with low prominence compared to surrounding peaks.

Confirmation of targeting in *C. campestris* genes (Fig. S7) was performed using the similar methods as above, with the following changes. Secondary siRNAs were annotated with ShortStack (*40*) to the *C. campestris* genome (*8*), using gene annotations as the basis for loci. Different NanoPARE libraries were used, coming from mixed host-parasite interface (IN), and were aligned to the *C. campestris* transcriptome (*8*). No direct control was present to compare peak expression, so the few confirmed examples could not be subjected to this filter.

*mRNA-seq analysis*

mRNA-seq libraries were aligned to the *A. thaliana* genome (*39*) using HISAT2 (*43*) (hisat2 -p 2 --max-intronlen 5000 -x genome.fa -U library.fq.gz). Gene expression was quantified by minBamCov (*44*) (multiBamCov -bams alignment.bam -bed annotation.gff) using the ARAPORT11 annotation (*36*). DEseq2 (*37*) was used to accurately estimate log fold change of mRNAs for each condition, using the "lfcShrink" command (type=apeglm).

Identification of *C. campestris* miRNAs

To identify sRNAs that were derived from miRNA hairpins, *de novo* annotation of sRNA loci in the *C. campestris* genome (*39*) was performed using ShortStack (*40*). Next, loci containing a HI-sRNA from *C. campestris* were extracted and screened by eye to find miRNAs with the criteria that they have a clear concise hairpin with 2 matching regions of expression which have a clear 2 nt offset (factors consistent with miRNA processing). Superfamilies were annotated to identify which contained confirmed miRNAs.

Discovery of target homologs in eudicots

cDNA and CDS libraries of 36 eudicot species (Table S3) available in phytozome v12.1.6 (*45*) were downloaded for local analysis. Nucleotide queries of *A. thaliana* target transcripts were searched against translated CDS libraries from eudicots using blastx (*46*) (blastx -query target.fa -db eudicot.db -outfmt 6 -num_threads 6 -evalue 0.001 -task blastx-fast), extracting the best hit for each species based on bit score. Conservation of target site and coding sequence of homologs was calculated by aligning their translated coding sequences using muscle (*31*) and measuring the average conservation (shannon entropy) of every 8 amino acid window, flagging the window which corresponds to the target site. RNA superfamilies and transcripts of best-hit homologs of target were each aligned using muscle (*31*) and oriented to each other using in-house scripts. Conservation for each position in homologs and superfamily were calculated and used to construct a linear model in R.

Discovery of conserved motifs in *A. thaliana* targets

Conserved motifs targeted by sRNA superfamilies in *A. thaliana* were found by first extracting all targets of a superfamily with very strong predicted targeting (complementarity score (*41*) 3 or less). Using an in-house script, sequences of target sites were translated for the correct frame and clustered using a greedy algorithm with a maximum edit distance in a cluster of 3 or less. Conservation of target sites and surrounding nucleotide sequences were then calculated and oriented adjacent to multiple sequence alignments of the targeting superfamily, highlighting confirmed interactions.

Code availability

ShortStack (*40*) and GSTAr.pl are both freely available at https://github.com/MikeAxtell. Muscle (*31*) is freely available at https://www.drive5.com/muscle/. MEGA7 (*34*) is freely available at https://www.megasoftware.net/. Blast-suite (*46*) is freely available at https://blast.ncbi.nlm.nih.gov. Cutadapt (*35*) is freely available at https://cutadapt.readthedocs.io/en/stable/. Bamtools (*44*) is freely available at https://bedtools.readthedocs.io/en/latest/index.html#. The R package DEseq2 (*37*) is freely available at https://bioconductor.org/packages/release/bioc/html/DESeq2.html. HISAT2 (*43*) and bowtie (*42*) are both freely available at https://ccb.jhu.edu/software. Ushuffle (*38*) is freely available at https://github.com/guma44/ushuffle. All scripts and code used in this publication will be made available upon request.

Data availability

sRNA-seq data from this work are available at the NCBI SRA under BioProject PRJNA543296

.

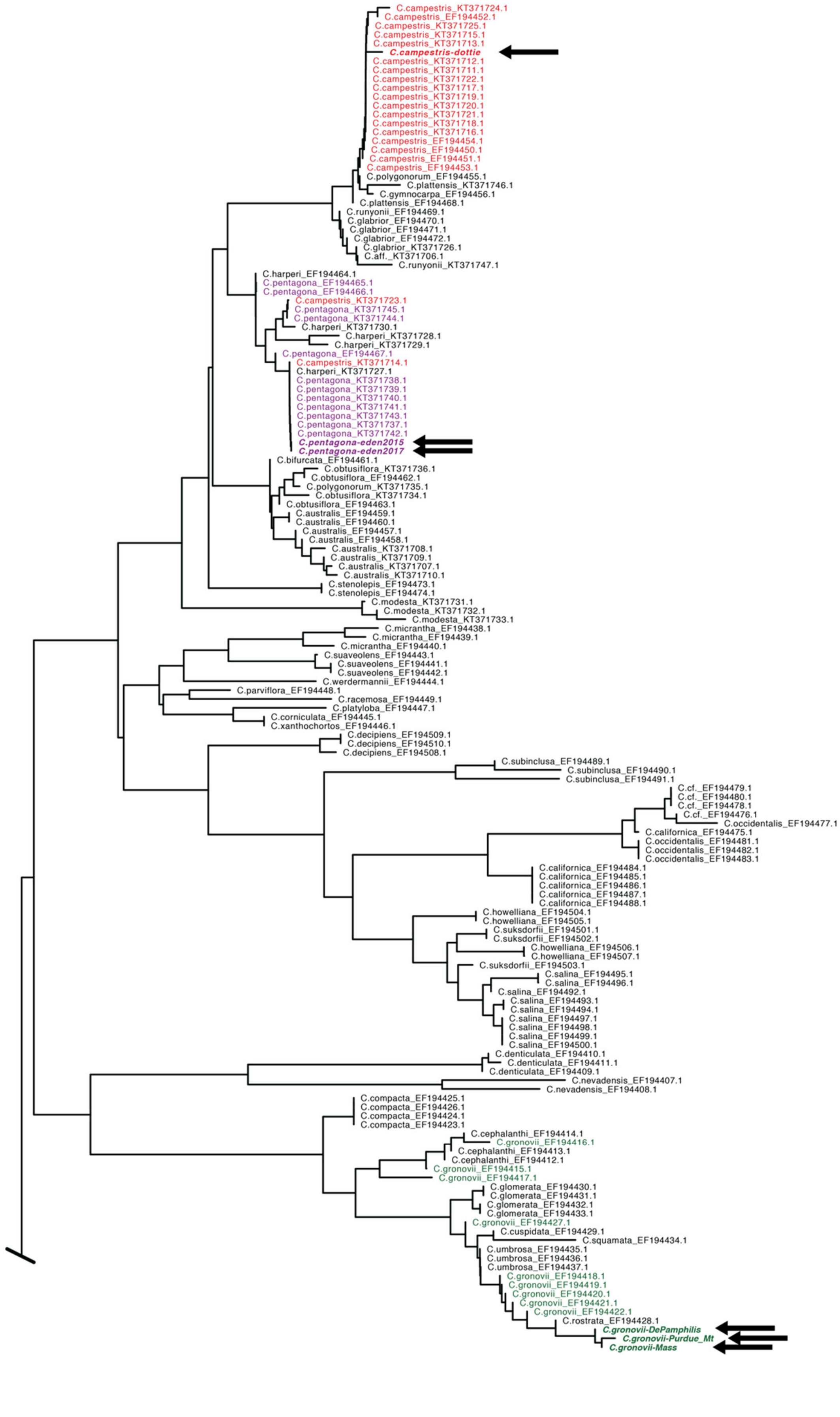
C.campestris_KT371724.1
C.campestris_EF194452.1
C.campestris_KT371725.1
C.campestris_KT371715.1
C.campestris_KT371713.1
C.campestris-dottie
C.campestris_KT371712.1
C.campestris_KT371711.1
C.campestris_KT371722.1
C.campestris_KT371717.1
C.campestris_KT371719.1
C.campestris_KT371720.1
C.campestris_KT371721.1
C.campestris_KT371718.1
C.campestris_KT371716.1
C.campestris_EF194454.1
C.campestris_EF194450.1
C.campestris_EF194451.1
C.campestris_EF194453.1
C.polygonorum_EF194455.1
C.plattensis_KT371746.1
C.gymnocarpa_EF194456.1
C.plattensis_EF194468.1
C.runyonii_EF194469.1
C.glabrior_EF194470.1
C.glabrior_EF194471.1
C.glabrior_EF194472.1
C.glabrior_KT371726.1
C.aff._KT371706.1
C.runyonii_KT371747.1
C.harperi_EF194464.1
C.pentagona_EF194465.1
C.pentagona_EF194466.1
C.campestris_KT371723.1
C.pentagona_KT371745.1
C.pentagona_KT371744.1
C.harperi_KT371730.1
C.harperi_KT371728.1
C.harperi_KT371729.1
C.pentagona_EF194467.1
C.campestris_KT371714.1
C.harperi_KT371727.1
C.pentagona_KT371738.1
C.pentagona_KT371739.1
C.pentagona_KT371740.1
C.pentagona_KT371741.1
C.pentagona_KT371743.1
C.pentagona_KT371737.1
C.pentagona_KT371742.1
C.pentagona-eden2015
C.pentagona-eden2017
C.bifurcata_EF194461.1
C.obtusiflora_KT371736.1
C.obtusiflora_EF194462.1
C.polygonorum_KT371735.1
C.obtusiflora_KT371734.1
C.obtusiflora_EF194463.1
C.australis_EF194459.1
C.australis_EF194460.1
C.australis_EF194457.1
C.australis_EF194458.1
C.australis_KT371708.1
C.australis_KT371709.1
C.australis_KT371707.1
C.australis_KT371710.1
C.stenolepis_EF194473.1
C.stenolepis_EF194474.1
C.modesta_KT371731.1
C.modesta_KT371732.1
C.modesta_KT371733.1
C.micrantha_EF194438.1
C.micrantha_EF194439.1
C.micrantha_EF194440.1
C.suaveolens_EF194443.1
C.suaveolens_EF194441.1
C.suaveolens_EF194442.1
C.werdermannii_EF194444.1
C.parviflora_EF194448.1
C.racemosa_EF194449.1
C.platyloba_EF194447.1
C.corniculata_EF194445.1
C.xanthochortos_EF194446.1
C.decipiens_EF194509.1
C.decipiens_EF194510.1
C.decipiens_EF194508.1
C.subinclusa_EF194489.1
C.subinclusa_EF194490.1
C.subinclusa_EF194491.1
C.cf._EF194479.1
C.cf._EF194480.1
C.cf._EF194478.1
C.cf._EF194476.1
C.occidentalis_EF194477.1
C.californica_EF194475.1
C.occidentalis_EF194481.1
C.occidentalis_EF194482.1
C.occidentalis_EF194483.1
C.californica_EF194484.1
C.californica_EF194485.1
C.californica_EF194486.1
C.californica_EF194487.1
C.californica_EF194488.1
C.howelliana_EF194504.1
C.howelliana_EF194505.1
C.suksdorfii_EF194501.1
C.suksdorfii_EF194502.1
C.howelliana_EF194506.1
C.howelliana_EF194507.1
C.suksdorfii_EF194503.1
C.salina_EF194495.1
C.salina_EF194496.1
C.salina_EF194492.1
C.salina_EF194493.1
C.salina_EF194494.1
C.salina_EF194497.1
C.salina_EF194498.1
C.salina_EF194499.1
C.salina_EF194500.1
C.denticulata_EF194410.1
C.denticulata_EF194411.1
C.denticulata_EF194409.1
C.nevadensis_EF194407.1
C.nevadensis_EF194408.1
C.compacta_EF194425.1
C.compacta_EF194426.1
C.compacta_EF194424.1
C.compacta_EF194423.1
C.cephalanthi_EF194414.1
C.gronovii_EF194416.1
C.cephalanthi_EF194413.1
C.cephalanthi_EF194412.1
C.gronovii_EF194415.1
C.gronovii_EF194417.1
C.glomerata_EF194430.1
C.glomerata_EF194431.1
C.glomerata_EF194432.1
C.glomerata_EF194433.1
C.gronovii_EF194427.1
C.cuspidata_EF194429.1
C.squamata_EF194434.1
C.umbrosa_EF194435.1
C.umbrosa_EF194436.1
C.umbrosa_EF194437.1
C.gronovii_EF194418.1
C.gronovii_EF194419.1
C.gronovii_EF194420.1
C.gronovii_EF194421.1
C.gronovii_EF194422.1
C.rostrata_EF194428.1
C.gronovii-DePamphilis
C.gronovii-Purdue_Mt
C.gronovii-Mass

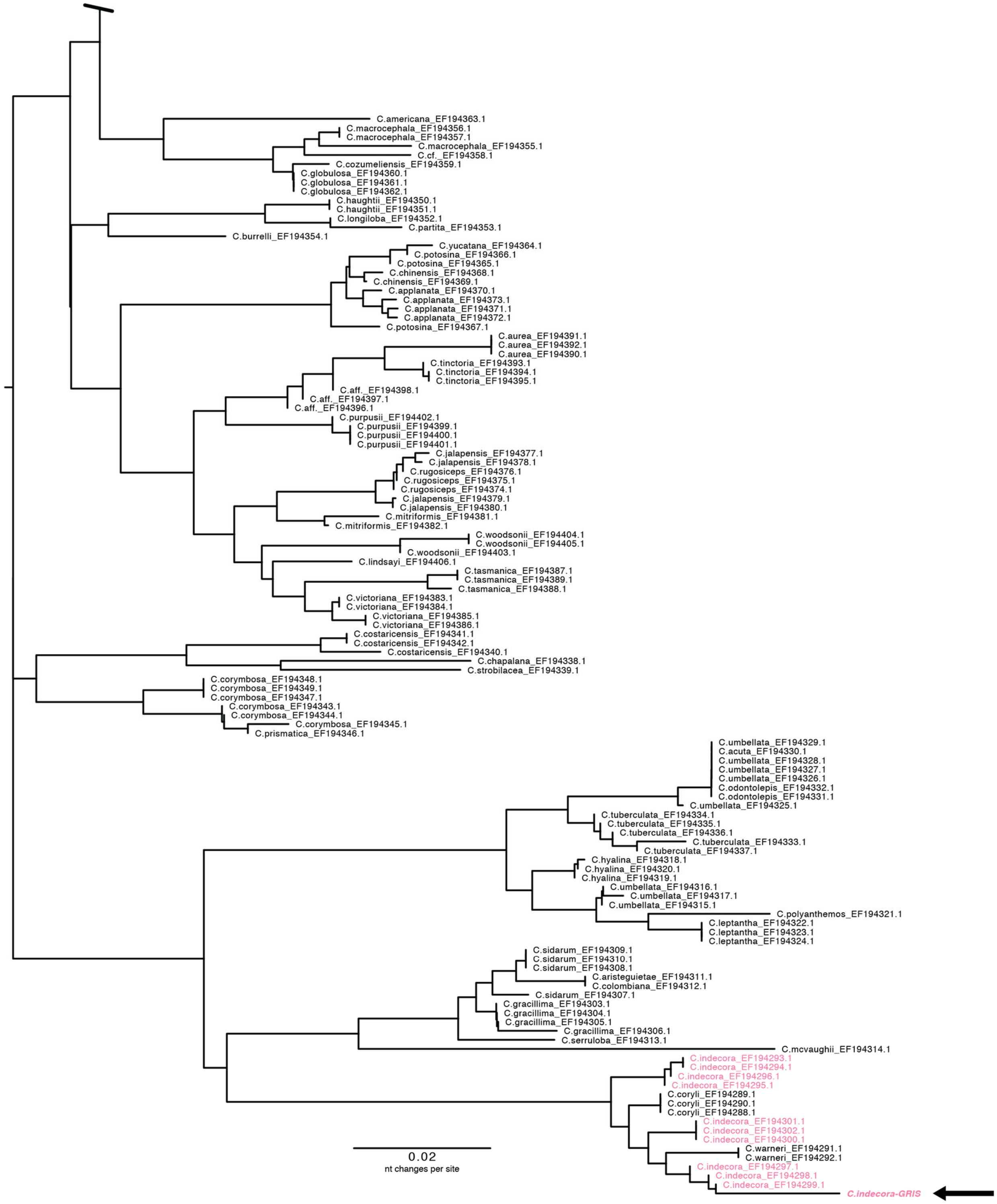


**Fig. S1. Unabridged phylogeny of Cuscuta**

Phylogeny based on TrnL-F sequencing using vouchered samples and primers (*34*, *35*). Isolates used in this study are in bold and indicated with arrows. Samples identified as members of species examined in this study are highlighted with color; red - *C. campestris*, purple - *C. pentagona*, green - *C. gronovii*, pink - *C. indecora*.

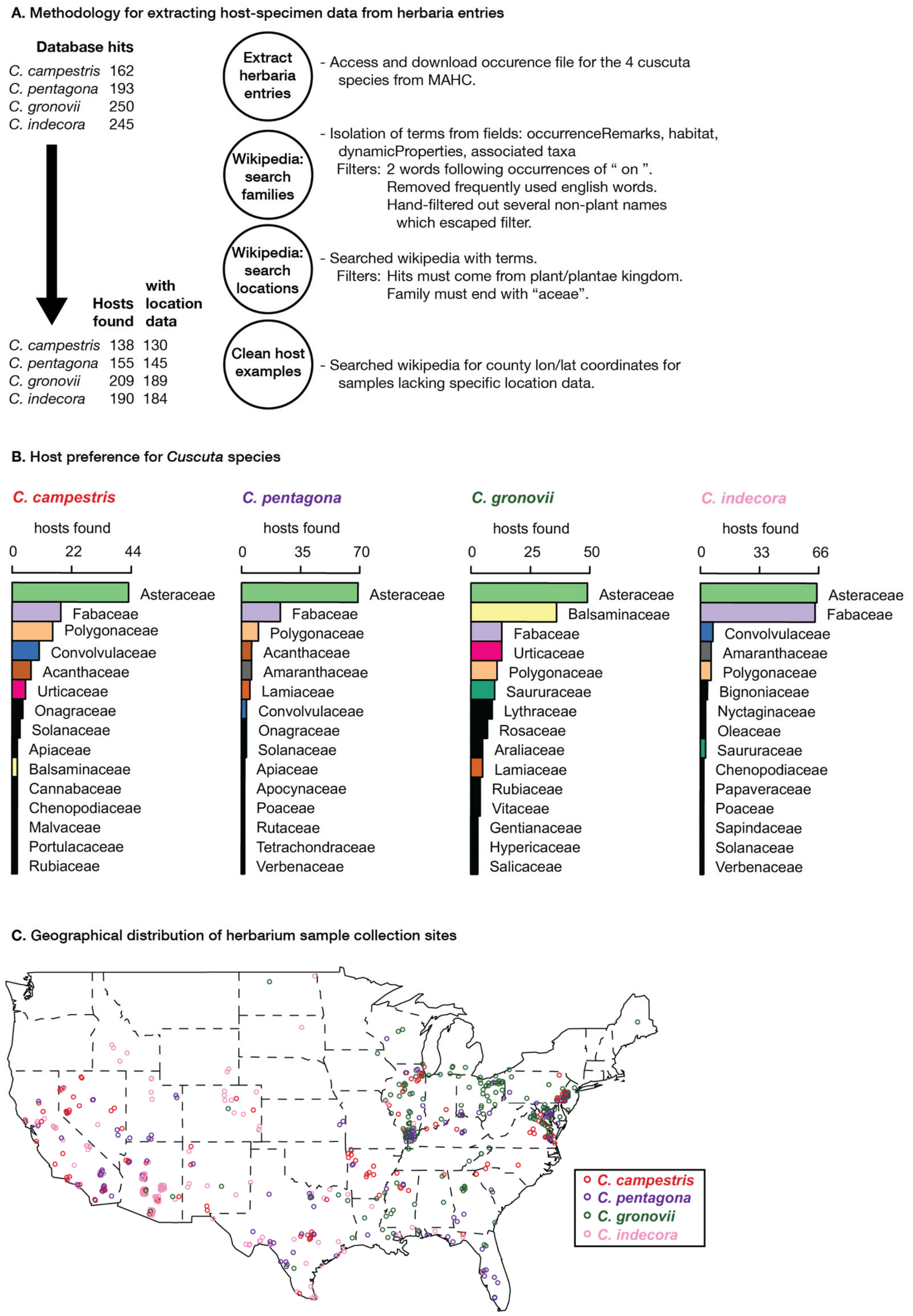


**Fig. S2. Host preference in *Cuscuta* species**

(**A**) Pipeline for processing herbaria data on interactions with each *Cuscuta* species of interest. (**B**) Ranked list of most identified host families for each species. Top 15 are shown for each species, with the top 10 overall identified with consistent colors (all others in black). (**C**) Geographical listings for each sample, where latitude and longitude or a searchable county are found.

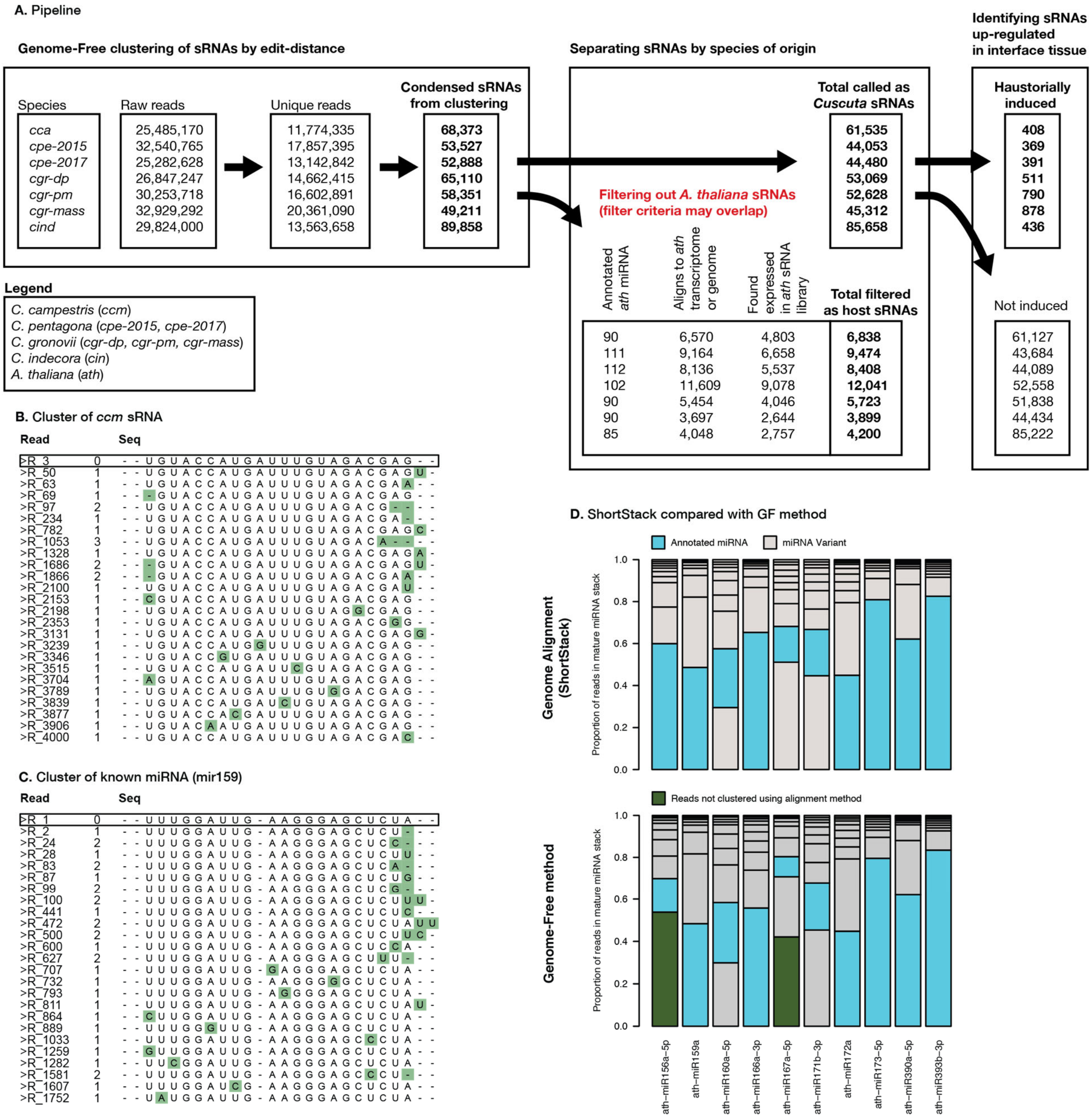


**Fig. S3. Genome-free HI-sRNA discovery pipeline**

(**A**) Discovery of HI-sRNAs in *Cuscuta* isolates. Three major steps include condensing reads to representative sRNAs in a genome-free manner, filtering reads which may be originating from *A. thaliana*, and performing differential expression with DEseq2 to find reads up-regulated in the interface tissue (FDR < 0.1, null hypothesis: sRNA not differentially expressed). (**B**) Example of a *C. campestris* sRNA discovered by this method, with the top 25 constituent sRNA sequences ranked by expression. Highest expressed read is deemed as representative sRNA sequence and is shown with black box. Green boxes show variations from representative sequences with total distance shown to left. (**C**) Same as B but with a known miRNA, showing similar variation to the novel sRNA in B. (**D**) Comparing the proportion of reads present in annotated miRNAs, using both genome-alignment (ShortStack) and genome-free based approaches. Reads are ranked by size, with the canonical miRNA (blue) and the variants (grey) showing the proportion of reads they make up in the sRNA. Reads grouped in the locus by the genome-free method that are absent in the alignment approach are shown in green.

**A.** Attributes of confirmed target-sites for *Cuscuta* sRNAs

| | Avg. Target site conservation (bits) | Correlation coeff. | P-value | Count of homologs found in phytozome | | | Count of sRNAs in superfamily | Confirmed as miRNA in *C. campestris* | |
|---|---|---|---|---|---|---|---|---|---|
| 1 | 4.32 | 0.38 | 0.08 | 33 | AT5G61480 | PXY | 5 | NoAl | SupFam_124 |
| 2 | 3.83 | 0.54 | 0.03 | 35 | AT1G20696 | HMGB3 | 10 | Y | SupFam_20 |
| 3 | 4.32 | -0.13 | 0.62 | 9 | AT3G14280 | LL-diamino... | 10 | Y | SupFam_20 |
| 4 | 3'UTR | | | 27 | AT3G20910 | NF-YA9 | 6 | Y | SupFam_72 |
| 5 | 4.1 | 0.57 | 0 | 36 | AT5G02290 | NAK | 5 | NoAl | SupFam_124 |
| 6 | 3.98 | | | 31 | AT4G01650 | Polyketide... | 5 | Y | SupFam_134 |
| 7 | 3.93 | | | 34 | AT2G17220 | Kin3 | 3 | Y | SupFam_320 |
| 8 | 4.11 | -0.04 | 0.86 | 36 | AT2G46110 | KPHMT1 | 13 | Y | SupFam_50 |
| 9 | 3.03 | 0.4 | 0.09 | 34 | AT3G52610 | GATA_zinc_... | 12 | Y | SupFam_18 |
| 10 | 4.26 | 0.46 | 0.03 | 34 | AT4G28490 | HAE | 5 | NoAl | SupFam_124 |
| 11 | 3.85 | 0.73 | 0 | 36 | AT2G39660 | BIK1 | 25 | Y | SupFam_1 |
| 12 | 4.26 | | | 36 | AT2G39660 | BIK1 | 4 | Y | SupFam_190 |
| 13 | 3.96 | | | 36 | AT5G60570 | Galactose_... | 3 | Y | SupFam_281 |
| 14 | 4.32 | 0.23 | 0.3 | 35 | AT1G46264 | HSFB4 | 5 | Y | SupFam_116 |
| 15 | 4.32 | | | 33 | AT3G27300 | G6PD5 | 3 | Y | SupFam_312 |
| 16 | 3.84 | 0.56 | 0.02 | 36 | AT3G26810 | AFB2 | 11 | Y | SupFam_27 |
| 17 | 4.03 | 0.62 | 0.01 | 35 | AT3G62980 | TIR1 | 11 | Y | SupFam_27 |
| 18 | 4.2 | | | 36 | AT5G07350 | Tudor1 | 3 | Y | SupFam_287 |
| 19 | 3.75 | 0.52 | 0.04 | 36 | AT1G12820 | AFB3 | 11 | Y | SupFam_27 |
| 20 | 2.76 | | | 36 | AT1G12820 | AFB3 | 4 | Y | SupFam_196 |
| 21 | 3.83 | -0.04 | 0.88 | 31 | AT1G67310 | Calmodulin... | 8 | Y | SupFam_39 |
| 22 | 3.6 | 0.76 | 0 | 34 | AT3G01680 | SEOR1 | 15 | Y | SupFam_5 |
| 23 | 2.49 | -0.27 | 0.25 | 33 | AT3G01680 | SEOR1 | 13 | Y | SupFam_19 |
| 24 | 4.28 | -0.24 | 0.31 | 32 | AT3G53840 | Protein_ki... | 4 | | SupFam_257 |
| 25 | 5'UTR | | | 13 | AT3G51990 | Protein_ki... | 3 | Y | SupFam_269 |
| 26 | 4.02 | -0.28 | 0.22 | 36 | AT3G51990 | Protein_ki... | 4 | | SupFam_257 |
| 27 | 4.3 | -0.23 | 0.33 | 36 | AT4G00330 | CRCK2 | 4 | | SupFam_257 |
| 28 | 4.04 | 0.63 | 0.01 | 36 | AT4G03190 | AFB1 | 11 | Y | SupFam_27 |
| 29 | 4.3 | 0.88 | 0 | 36 | AT5G65430 | GRF8 | 13 | Y | SupFam_24 |
| 30 | 4.26 | -0.22 | 0.36 | 35 | AT1G21590 | Protein_ki... | 4 | | SupFam_257 |
| 31 | 4.16 | -0.2 | 0.4 | 35 | AT3G52870 | IQ_calmodu... | 5 | | SupFam_188 |
| 32 | 3.01 | 0.22 | 0.32 | 34 | AT3G09830 | PCRK1 | 6 | | SupFam_81 |
| 33 | 3.72 | | | 35 | AT1G10900 | Phosphatid... | 3 | | SupFam_370 |
| 34 | 4.06 | 0.71 | 0 | 35 | AT1G65800 | ARK2 | 15 | Y | SupFam_49 |
| 35 | 4.3 | 0.65 | 0 | 36 | AT1G63910 | AtMYB103 | 10 | | SupFam_37 |
| 36 | 4.27 | 0.51 | 0.02 | 36 | AT5G24010 | Protein_ki... | 15 | Y | SupFam_49 |
| 37 | 3.63 | 0.08 | 0.75 | 7 | AT3G06380 | TLP9 | 17 | | SupFam_14 |
| 38 | 3.31 | 0.1 | 0.66 | 35 | AT2G36570 | PXC1 | 5 | | SupFam_161 |
| 39 | 3.04 | | | 35 | AT2G25170 | PKL | 4 | | SupFam_206 |
| 40 | 4.09 | 0.79 | 0 | 33 | AT2G02800 | APK2B | 19 | Y | SupFam_26 |
| 41 | 2.03 | | | 8 | AT4G28300 | DUF1421_co... | 1 | | SupFam_897 |
| 42 | 4.32 | 0.57 | 0.01 | 36 | AT4G39400 | BRI1 | 8 | | SupFam_59 |
| 43 | 3.52 | 0.24 | 0.32 | 7 | AT5G39610 | NAC6 | 5 | | SupFam_187 |
| 44 | 3.17 | | | 36 | AT3G51670 | SEC14_cyto... | 10 | | SupFam_37 |
| 45 | 3.81 | -0.36 | 0.12 | 36 | AT1G67230 | LINC1 | 14 | Y | SupFam_3 |
| 46 | 4.3 | | | 35 | AT4G27630 | GTG2 | 1 | | SupFam_895 |
| 47 | 3.64 | -0.26 | 0.26 | 36 | AT4G23630 | BTI1 | 4 | | SupFam_207 |
| 48 | 3.5 | -0.39 | 0.1 | 36 | AT5G24760 | GroES-like... | 16 | | SupFam_4 |
| 49 | 3.39 | 0.01 | 0.97 | 34 | AT5G46580 | PPR_repeat... | 19 | Y | SupFam_26 |
| 50 | 4.17 | 0.12 | 0.65 | 33 | AT3G27925 | DEG1 | 9 | | SupFam_33 |
| 51 | 4.07 | 0.43 | 0.07 | 34 | AT5G56460 | Protein_ki... | 16 | | SupFam_4 |
| 52 | 4.25 | 0.53 | 0.02 | 36 | AT1G53440 | Leucine-ri... | 8 | | SupFam_59 |
| 53 | 3.53 | -0.08 | 0.74 | 13 | AT4G02400 | U3_ribonuc... | 10 | | SupFam_37 |
| 54 | 4.13 | 0.6 | 0 | 36 | AT5G66850 | MAPKKK5 | 5 | | SupFam_157 |
| 55 | 3.27 | | | 30 | AT4G00830 | LIF2 | 2 | | SupFam_490 |
| 56 | 4.3 | | | 36 | AT3G45640 | MPK3 | 1 | | SupFam_1385 |
| 57 | 3.86 | 0.56 | 0.01 | 35 | AT3G45640 | MPK3 | 4 | | SupFam_203 |
| 58 | 4.07 | 0.2 | 0.38 | 35 | AT4G38470 | STY46 | 5 | | SupFam_189 |
| 59 | 4.25 | | | 34 | AT5G11200 | UAP56b | 1 | | SupFam_1585 |
| 60 | 3'UTR | | | 25 | AT1G17590 | NF-YA8 | 1 | | SupFam_1364 |
| 61 | 4.32 | | | 36 | AT4G29040 | RPT2a | 1 | | SupFam_1368 |
| 62 | 4.12 | | | 36 | AT3G46740 | TOC75-III | 1 | | SupFam_1423 |
| 63 | 3.98 | | | 36 | AT1G02840 | SR34 | 2 | | SupFam_742 |
| 64 | 3.69 | | | 36 | AT1G18540 | Ribosomal_... | 1 | | SupFam_1431 |
| 65 | 1.62 | | | 23 | AT3G09070 | OPS | 1 | | SupFam_1431 |
| 66 | 4.25 | | | 34 | AT5G11170 | UAP56a | 1 | | SupFam_1585 |

Confirmation status | Target interaction | Target | sRNA superfamily

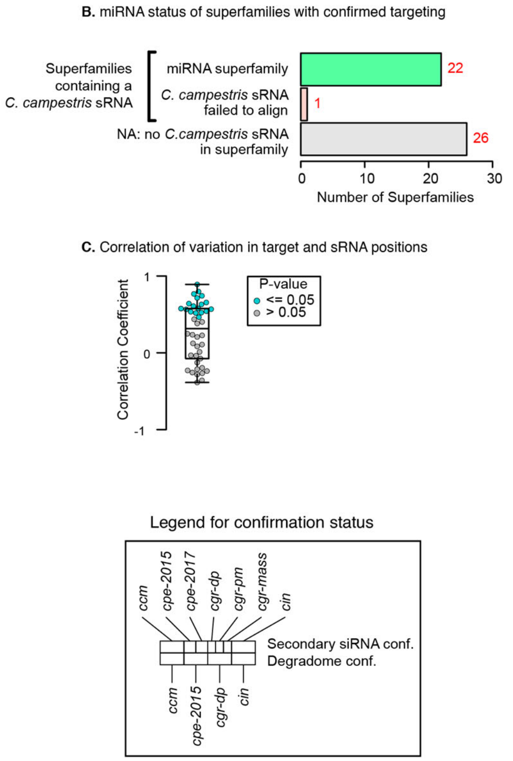


**Fig. S4. Summary of *Cuscuta* HI-sRNA and host gene target relationships**
(**A**) Complete list of target interactions between sRNAs and host genes. Confirmation status diagram indicates in what species the interaction is confirmed. Target gene information includes the number of homologs found in 36 eudicot transcriptomes. sRNA counts in superfamilies and the presence of a confirmed miRNA in the family is shown (NoAl: *ccm* sRNA failed to align to *ccm* genome). Target interaction columns indicate the conservation at the translated target site in an alignment of found homologs (5'/3' UTR: not considered for conservation analysis). Correlation coefficient and P-value for variation in positions in target and sRNA superfamily shown. (**B**) Breakdown of superfamilies with confirmed targeting by the presence of a confirmed miRNA, where possible. (**C**) Correlation of positional variation in target-sites and their sRNAs, indicating the interactions with a significant correlation.

**Fig. S5. Target confirmation data for every confirmed HI-sRNA-target interaction**
Details of confirmed HI-sRNA targets including HI-sRNA-target complementarity, site, score, superfamily and the status of *C. campestris* superfamily members as a confirmed miRNA. Targeting confirmation for target mRNA is shown in upper right, with confirmed interactions in species highlighted in red. sRNA distribution at target locus is shown for experimental interface and control, demonstrating secondary siRNA phasing and size distribution for up-regulated loci. Degradome sequencing is shown where confirmed hits were discovered in NanoPARE data.

**Allenscore:** 2

**Target Interaction:**

```
5'- AUUGGAUUGAAGCUAUUGAUCCU   AT3G52870.1
     |||||  ||||||||||||||||
3'- UAACCGAACUUCGAUAACUAGGU   CI_cpe-2015_676
```

**Target Site:** 559

**Superfamily:** SupFam_188

**miRNA in ccm:** N/A

**Confirmed Targeting**

| *2nd-siRNA* | *NanoPARE* |
|---|---|
| *ccm* | *ccm* |
| *cpe-2015* | *cpe-2015* |
| *cpe-2017* | |
| *cgr-dp* | *cgr-dp* |
| *cgr-pm* | |
| *cgr-mass* | |
| *cin* | *cin* |

### IQ_calmodulin-binding_motif_family_prote

AT3G52870 - CI_cpe-2015_676

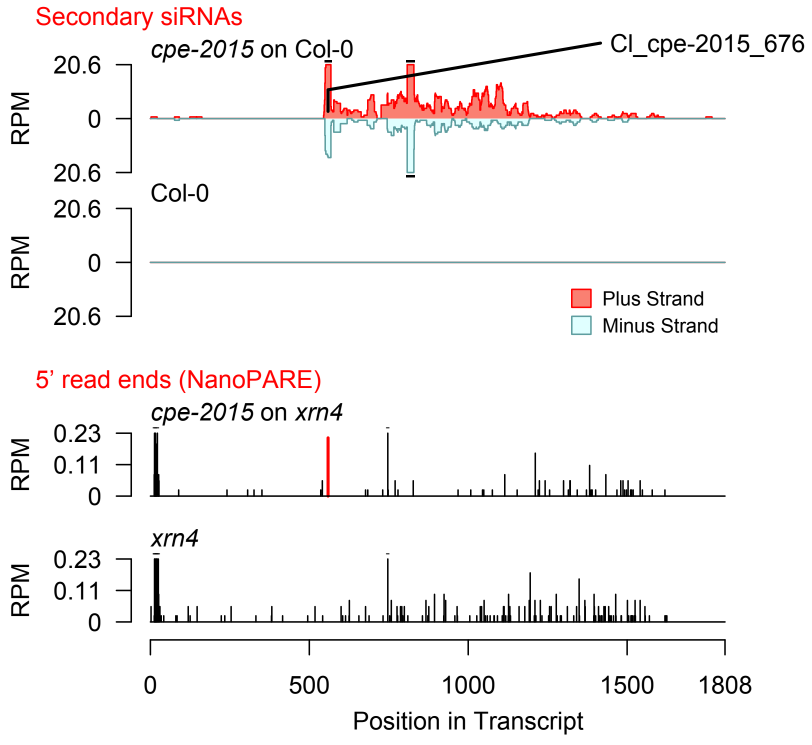


2nd siRNAs: Phase diagram

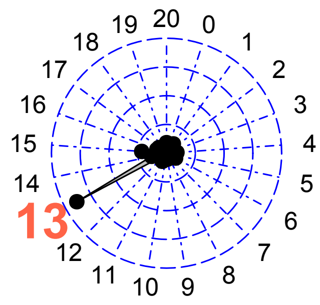


2nd siRNAs: Size distribution

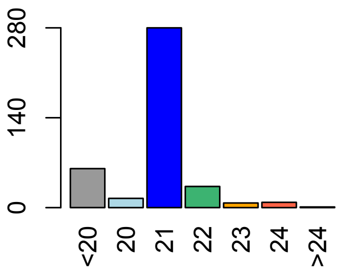

**Allenscore:** 2

**Target Interaction:**

```
5'- GGAUUGAAGCUAUUGAUCCU   AT3G52870.1
     | |  | | | | | | | | | | | | | | | |
3'- CCGAACUUCGAUAACUAGGU   CI_cpe-2015_4190
```

**Target Site:** 559

**Superfamily:** SupFam_188

**miRNA in ccm:** N/A

**Confirmed Targeting**

| *2nd-siRNA* | *NanoPARE* |
|---|---|
| *ccm* | *ccm* |
| *cpe-2015* | *cpe-2015* |
| *cpe-2017* | |
| *cgr-dp* | *cgr-dp* |
| *cgr-pm* | |
| *cgr-mass* | |
| *cin* | *cin* |

### IQ_calmodulin-binding_motif_family_prote

AT3G52870 - CI_cpe-2015_4190

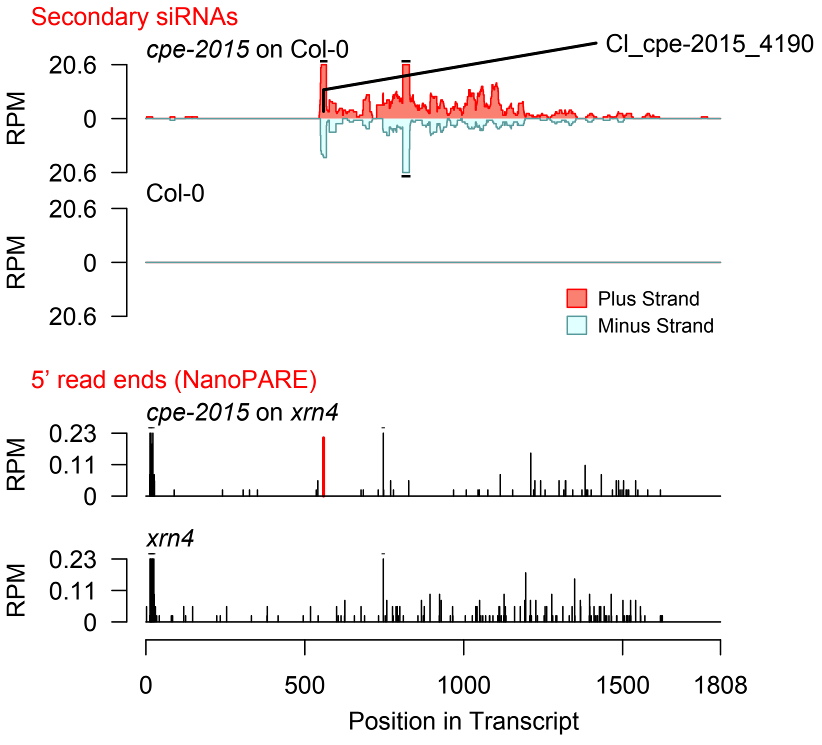


2nd siRNAs: Phase diagram

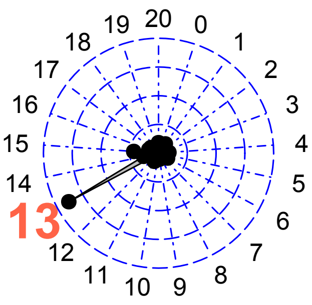


2nd siRNAs: Size distribution

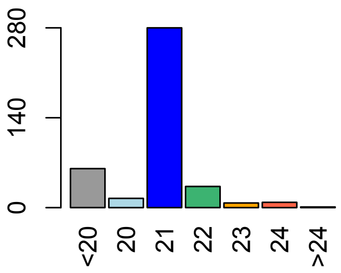

**Allenscore:** 2

**Target Interaction:**

```
5'- AUUGGAUUGAAGCUAUUGAUCCU   AT3G52870.1
     |||||  |||||||||||||||||
3'- UAACCGAACUUCGAUAACUAGGU   CI_cpe-2017_757
```

**Target Site:** 559

**Superfamily:** SupFam_188

**miRNA in ccm:** N/A

**Confirmed Targeting**

| *2nd-siRNA* | *NanoPARE* |
|---|---|
| *ccm* | *ccm* |
| *cpe-2015* | *cpe-2015* |
| *cpe-2017* | |
| *cgr-dp* | *cgr-dp* |
| *cgr-pm* | |
| *cgr-mass* | |
| *cin* | *cin* |

### IQ_calmodulin-binding_motif_family_prote

AT3G52870 - CI_cpe-2017_757

Secondary siRNAs

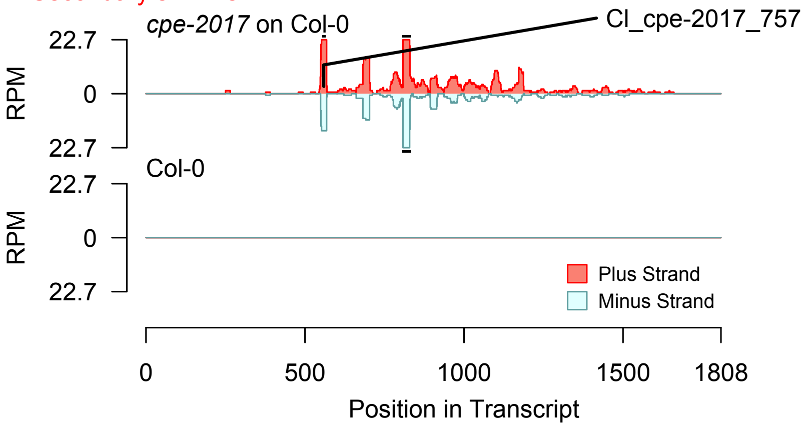


2nd siRNAs: Phase diagram

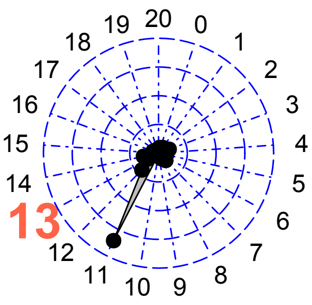


2nd siRNAs: Size distribution

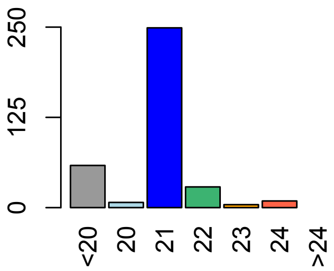


Degradome hits not found for sRNA

**Allenscore:** 2

**Target Interaction:**

```
5'- GGAUUGAAGCUAUUGAUCCU    AT3G52870.1
     | |   | | | | | | | | | | | | | | | |
3'- CCGAACUUCGAUAACUAGGU    CI_cpe-2017_3761
```

**Target Site:** 559

**Superfamily:** SupFam_188

**miRNA in ccm:** N/A

**Confirmed Targeting**

| *2nd-siRNA* | *NanoPARE* |
|---|---|
| *ccm* | *ccm* |
| *cpe-2015* | *cpe-2015* |
| *cpe-2017* | |
| *cgr-dp* | *cgr-dp* |
| *cgr-pm* | |
| *cgr-mass* | |
| *cin* | *cin* |

### IQ_calmodulin-binding_motif_family_prote

AT3G52870 - CI_cpe-2017_3761

Secondary siRNAs

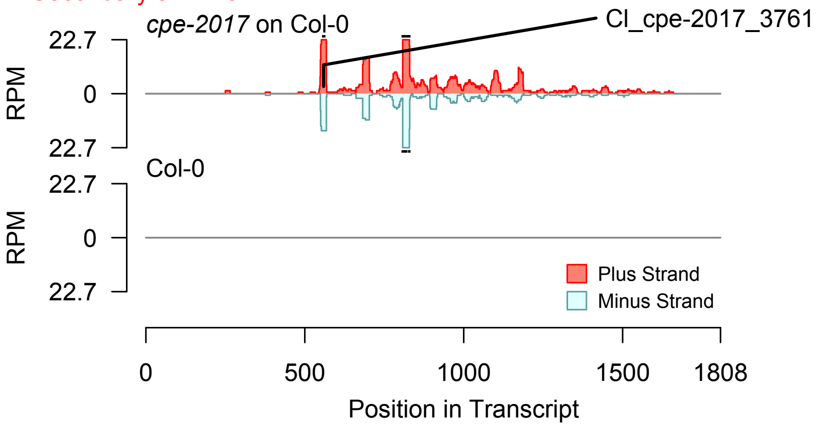


2nd siRNAs: Phase diagram

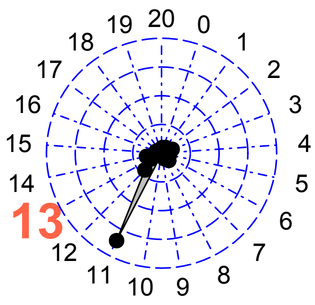


2nd siRNAs: Size distribution

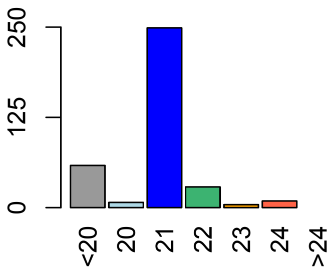


Degradome hits not found for sRNA

**Allenscore:** 2

**Target Interaction:**

```
5'- CUCCUGAGCAUUCUGCUGCUG   AT4G00830.1
      ||||||  |||||||||||||
3'- AAGGACUAGUAAGACGACGAC   Cl_cgr-dp_538
```

**Target Site:** 1081

**Superfamily:** SupFam_490

**miRNA in ccm:** N/A

**Confirmed Targeting**

| *2nd-siRNA* | *NanoPARE* |
|---|---|
| *ccm* | *ccm* |
| *cpe-2015* | *cpe-2015* |
| *cpe-2017* | |
| *cgr-dp* | *cgr-dp* |
| *cgr-pm* | |
| *cgr-mass* | |
| *cin* | *cin* |

### LIF2

AT4G00830 - Cl_cgr-dp_538

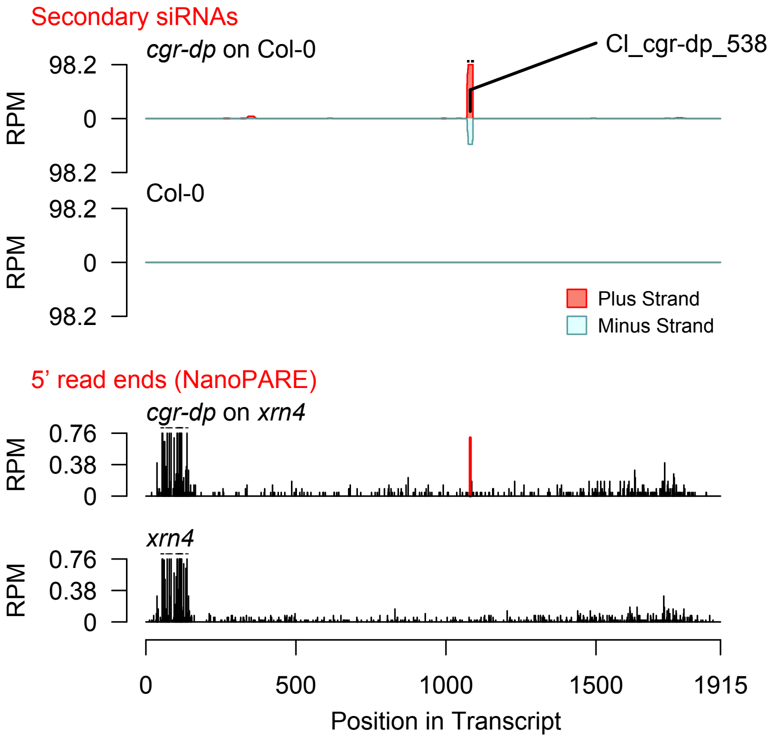


Diff. Exp. secondary siRNA locus not found

**Allenscore:** 3

**Target Interaction:**

```
5'- AUGGAUGUCAUCUGCCAAGCA   AT5G11170.1
     | | | | | | | | | | | | : : | | | | | |
3'- UACCUACAGUAGGUGGUUCGA   Cl_cin_42282
```

**Target Site:** 382

**Superfamily:** SupFam_1585

**miRNA in ccm:** N/A

**Confirmed Targeting**

| *2nd-siRNA* | *NanoPARE* |
|---|---|
| *ccm* | *ccm* |
| *cpe-2015* | *cpe-2015* |
| *cpe-2017* | |
| *cgr-dp* | *cgr-dp* |
| *cgr-pm* | |
| *cgr-mass* | |
| *cin* | *cin* |

### UAP56a

AT5G11170 - Cl_cin_42282

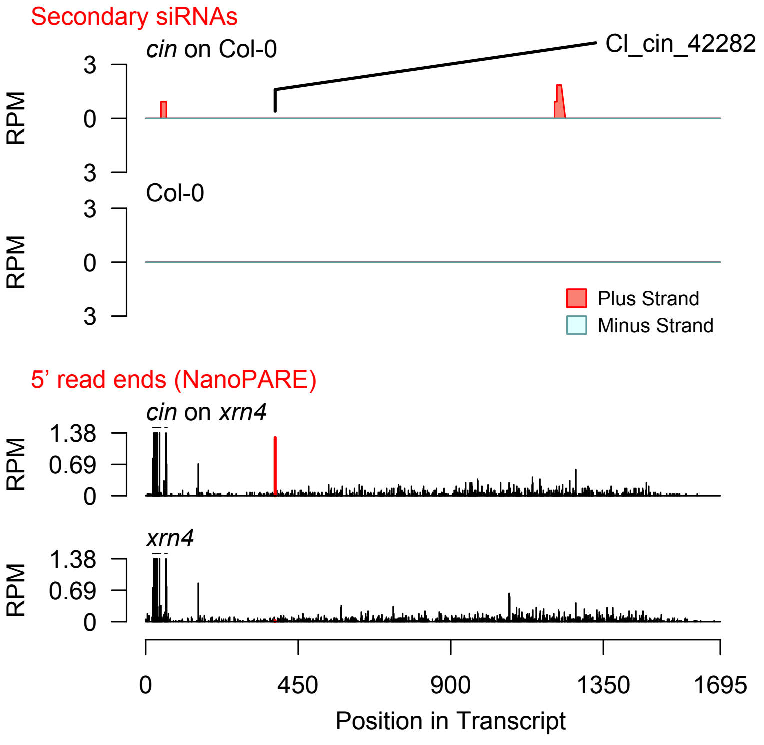


Diff. Exp. secondary siRNA locus not found

**Allenscore:** 4

**Target Interaction:**

```
5'- GUGUGCGAUGAAAGAAGUUGAG    AT5G66850.1
    | | | | | | : | | : | | | | | | |   | | | :
3'- CACACGUUAUUUUCUUC - ACUU   CI_cgr-dp_5622
```

**Target Site:** 1391

**Superfamily:** SupFam_157

**miRNA in ccm:** N/A

**Confirmed Targeting**

| *2nd-siRNA* | *NanoPARE* |
|---|---|
| *ccm* | *ccm* |
| *cpe-2015* | *cpe-2015* |
| *cpe-2017* | |
| *cgr-dp* | *cgr-dp* |
| *cgr-pm* | |
| *cgr-mass* | |
| *cin* | *cin* |

### MAPKKK5

AT5G66850 - CI_cgr-dp_5622

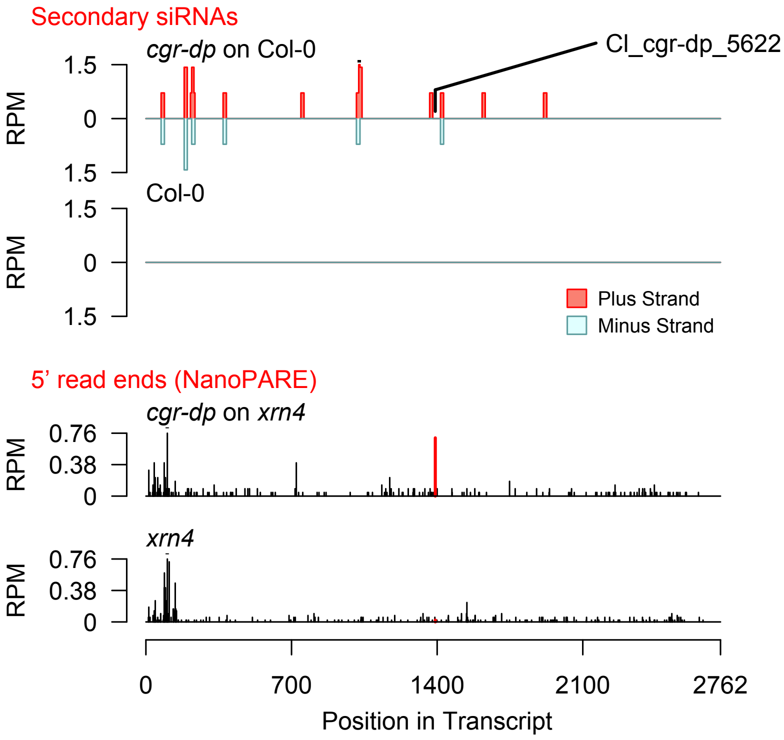


Diff. Exp. secondary siRNA locus not found

**Allenscore:** 1.5

**Target Interaction:**

```
5'- GUGUGCGAUGAAAGAAGUUGA   AT5G66850.1
     : | | : | | : | | | | | | | | | | | | | |
3'- UACGCGUUACUUUCUUCAACU   CI_cgr-dp_866
```

**Target Site:** 1391

**Superfamily:** SupFam_157

**miRNA in ccm:** N/A

**Confirmed Targeting**

| *2nd-siRNA* | *NanoPARE* |
|---|---|
| *ccm* | *ccm* |
| *cpe-2015* | *cpe-2015* |
| *cpe-2017* | |
| *cgr-dp* | *cgr-dp* |
| *cgr-pm* | |
| *cgr-mass* | |
| *cin* | *cin* |

### MAPKKK5

AT5G66850 - CI_cgr-dp_866

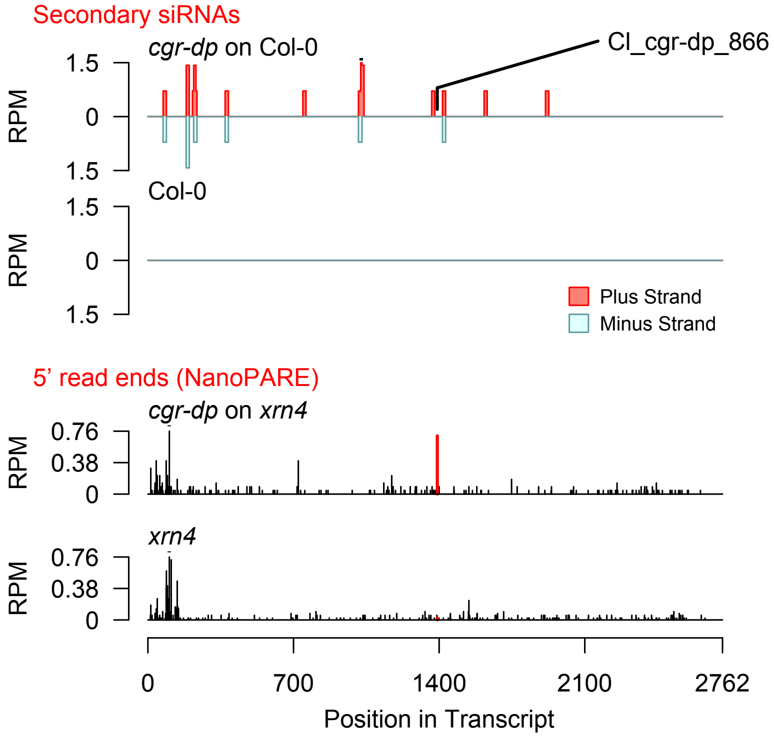


Diff. Exp. secondary siRNA locus not found

**Allenscore:** 5.5

**Target Interaction:**

```
5'- GAGAUGUUGUUCCACCACCA   AT3G45640.1
     : | | | | | | | : | | | |   | |   | | |
3'- UUCUACAAUAAGGAGGCGGU   CI_cgr-dp_22
```

**Target Site:** 659

**Superfamily:** SupFam_203

**miRNA in ccm:** N/A

**Confirmed Targeting**

| *2nd-siRNA* | *NanoPARE* |
|---|---|
| *ccm* | *ccm* |
| *cpe-2015* | *cpe-2015* |
| *cpe-2017* | |
| *cgr-dp* | *cgr-dp* |
| *cgr-pm* | |
| *cgr-mass* | |
| *cin* | *cin* |

### MPK3

AT3G45640 - CI_cgr-dp_22

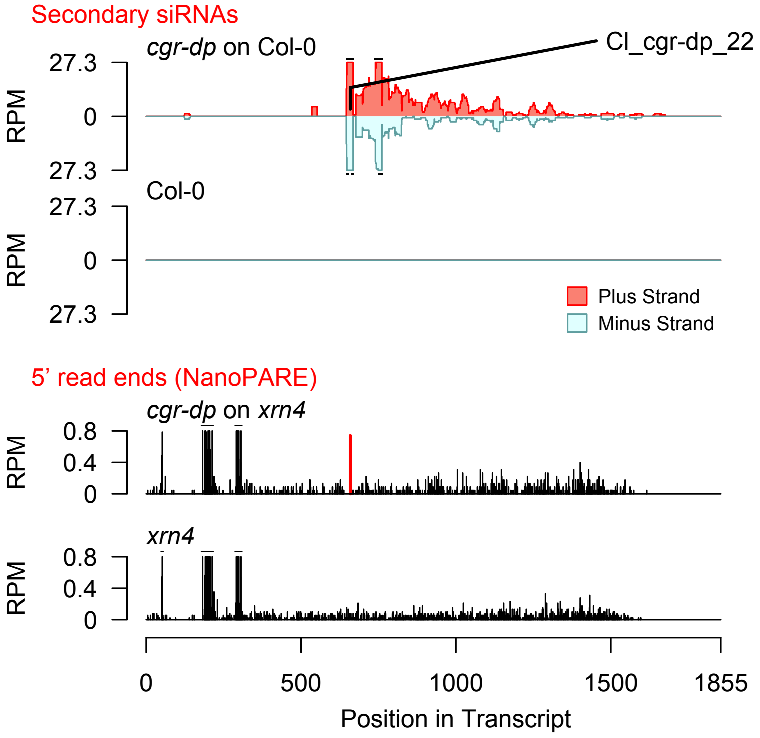


2nd siRNAs: Phase diagram

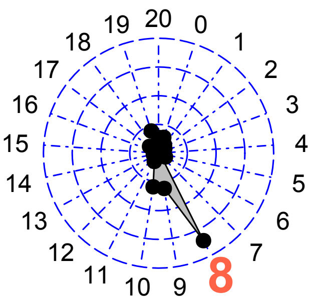


2nd siRNAs: Size distribution

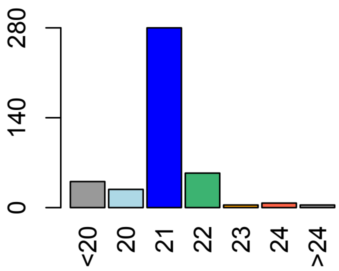

**Allenscore:** 5.5

**Target Interaction:**

```
5'- GAUGCUAAGCGUACGCUUCGUGA    AT3G45640.1
    |||||:|||||  ||:|||||  ||
3'- CUACGGUUCGCCUGUGAAGC-CU    Cl_cin_3787
```

**Target Site:** 591

**Superfamily:** SupFam_1385

**miRNA in ccm:** N/A

**Confirmed Targeting**

| *2nd-siRNA* | *NanoPARE* |
|---|---|
| *ccm* | *ccm* |
| *cpe-2015* | *cpe-2015* |
| *cpe-2017* | |
| *cgr-dp* | *cgr-dp* |
| *cgr-pm* | |
| *cgr-mass* | |
| *cin* | *cin* |

### MPK3

AT3G45640 - Cl_cin_3787

Secondary siRNAs

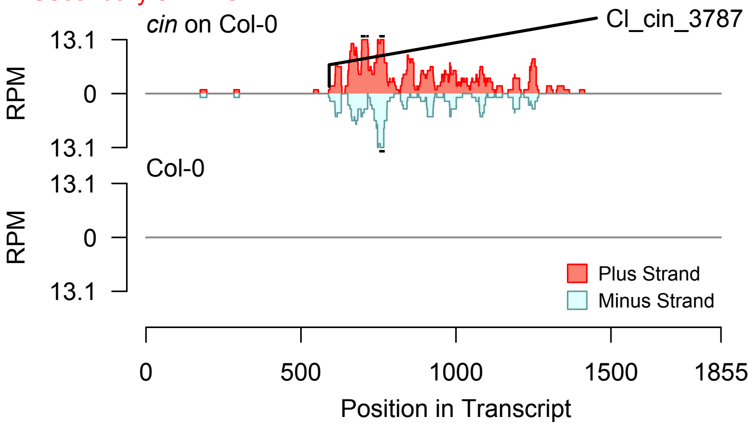


Degradome hits not found for sRNA

2nd siRNAs: Phase diagram

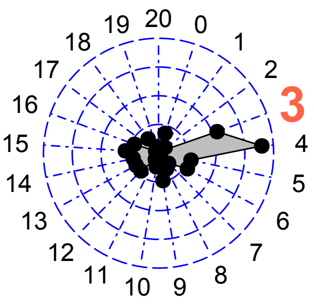


2nd siRNAs: Size distribution

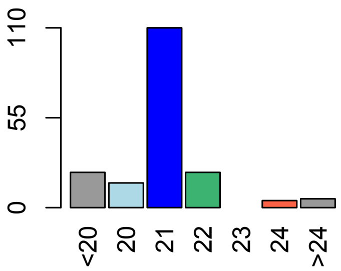

**Allenscore:** 6

**Target Interaction:**

```
5'- GCUCUUGGAAGAAGAUGAAAGGCUCC   AT4G02400.1
     : | : |       | | | | | | | | | | | | | : | | | |
3'- UGGG- - -CUUCUUCUACUUUUCGAGU   CI_cgr-dp_279
```

**Target Site:** 1850

**Superfamily:** SupFam_37

**miRNA in ccm:** N/A

**Confirmed Targeting**

| *2nd-siRNA* | *NanoPARE* |
|---|---|
| *ccm* | *ccm* |
| *cpe-2015* | *cpe-2015* |
| *cpe-2017* | |
| *cgr-dp* | *cgr-dp* |
| *cgr-pm* | |
| *cgr-mass* | |
| *cin* | *cin* |

### U3_ribonucleoprotein_family_protein

AT4G02400 - CI_cgr-dp_279

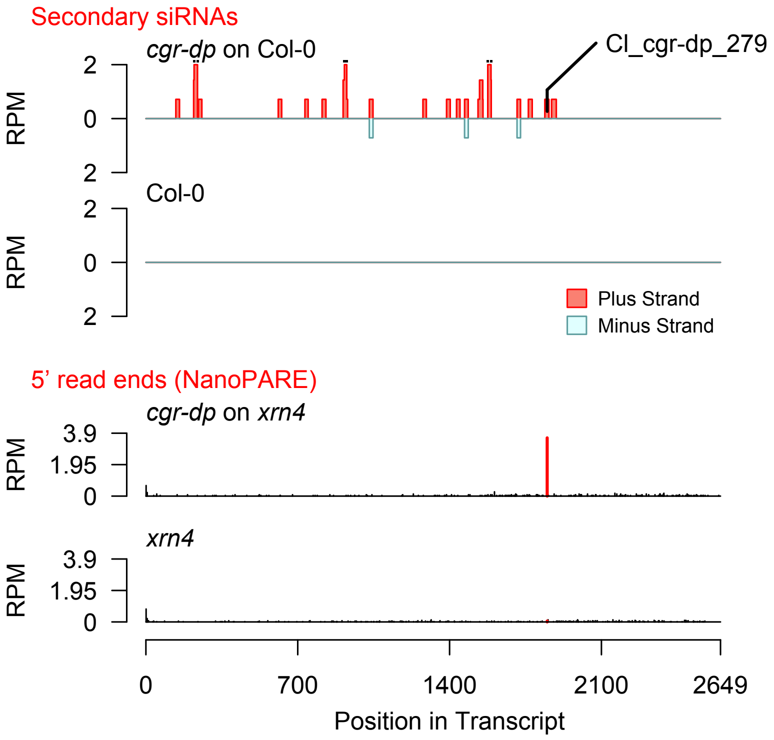


Diff. Exp. secondary siRNA locus not found

**Allenscore:** 3

**Target Interaction:**

```
5'- GGAAGAAGAUGAAAGGCUCC   AT4G02400.1
     | | | | | | | | | | | | | | : | | | |
3'- GCUUCUUCUACUUUUCGAGU   CI_cgr-dp_37384
```

**Target Site:** 1850

**Superfamily:** SupFam_37

**miRNA in ccm:** N/A

**Confirmed Targeting**

| *2nd-siRNA* | *NanoPARE* |
|---|---|
| *ccm* | *ccm* |
| *cpe-2015* | *cpe-2015* |
| *cpe-2017* | |
| *cgr-dp* | *cgr-dp* |
| *cgr-pm* | |
| *cgr-mass* | |
| *cin* | *cin* |

### U3_ribonucleoprotein_family_protein

AT4G02400 - CI_cgr-dp_37384

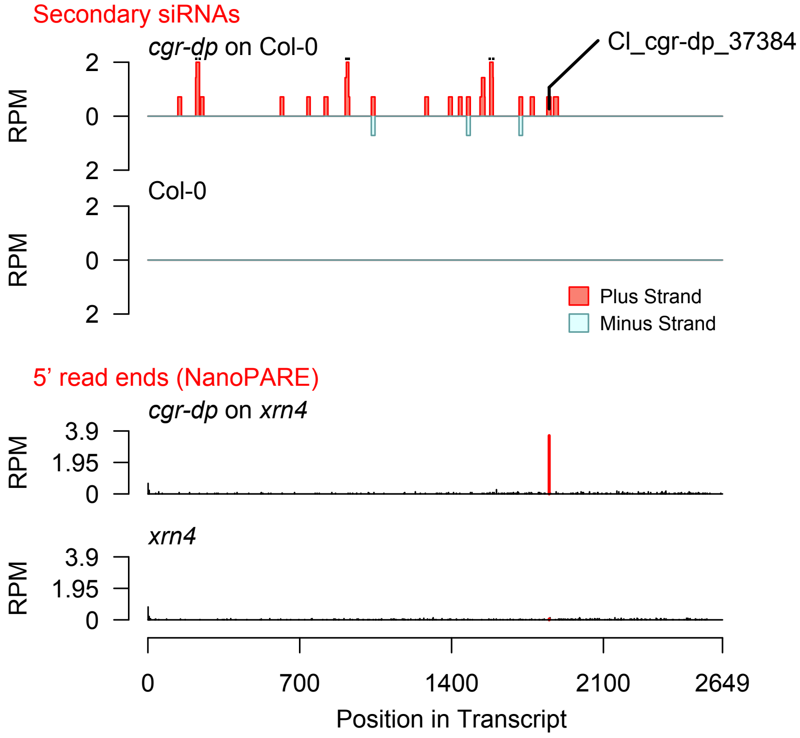


Diff. Exp. secondary siRNA locus not found

**Allenscore:** 3.5

**Target Interaction:**

```
5'- AGAGGAGGAGGAGGAAGACUA   AT3G09070.1
     |   ||||||:||  ||||||||||
3'- UGUCCUCUUCAUCCUUCUGAU   CI_cin_11140
```

**Target Site:** 1028

**Superfamily:** SupFam_1431

**miRNA in ccm:** N/A

**Confirmed Targeting**

| *2nd-siRNA* | *NanoPARE* |
|---|---|
| *ccm* | *ccm* |
| *cpe-2015* | *cpe-2015* |
| *cpe-2017* | |
| *cgr-dp* | *cgr-dp* |
| *cgr-pm* | |
| *cgr-mass* | |
| *cin* | *cin* |

### OPS

AT3G09070 - CI_cin_11140

Secondary siRNAs

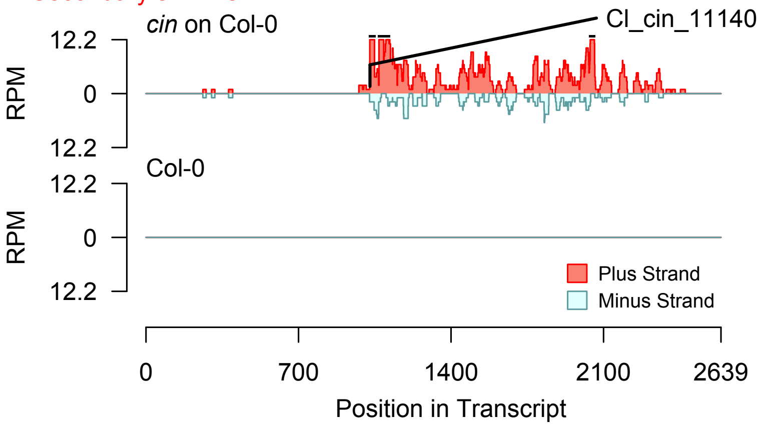


Degradome hits not found for sRNA

2nd siRNAs: Phase diagram

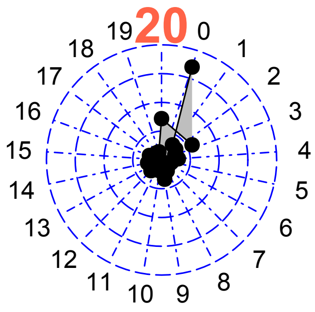


2nd siRNAs: Size distribution

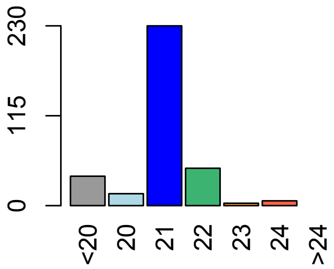

**Allenscore:** 4

**Target Interaction:**

```
5'- UUGGCUACUUAGCUCCAGAGUA    AT1G21590.1
     | | | |   | | | | | | | |   | | | | | | | |
3'- AACCAAUGAAUCUAGGUCUCAC    CI_cpe-2015_439
```

**Target Site:** 2056

**Superfamily:** SupFam_257

**miRNA in ccm:** N/A

**Confirmed Targeting**

| *2nd-siRNA* | *NanoPARE* |
|---|---|
| *ccm* | *ccm* |
| *cpe-2015* | *cpe-2015* |
| *cpe-2017* | |
| *cgr-dp* | *cgr-dp* |
| *cgr-pm* | |
| *cgr-mass* | |
| *cin* | *cin* |

### Protein_kinase_superfamily_protein

AT1G21590 - CI_cpe-2015_439

Secondary siRNAs

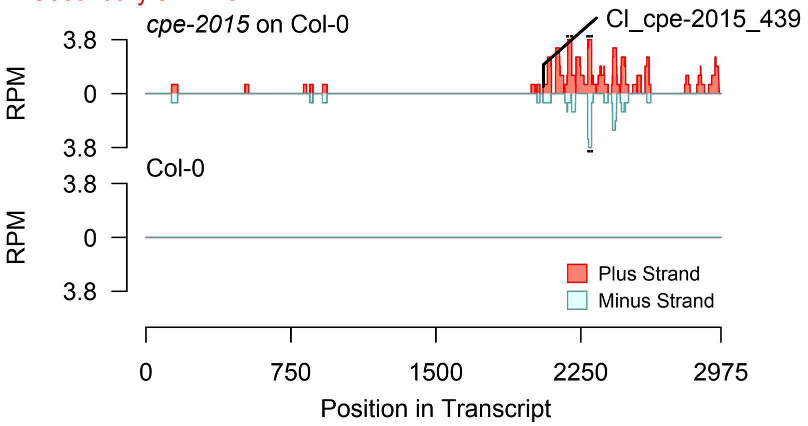


2nd siRNAs: Phase diagram

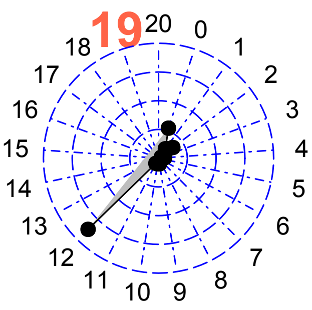


2nd siRNAs: Size distribution

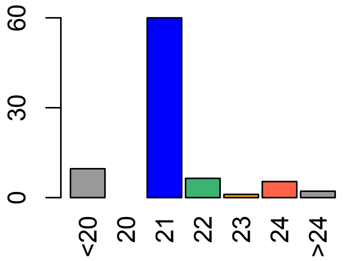


Degradome hits not found for sRNA

**Allenscore:** 4.5

**Target Interaction:**

```
5'- GGCUACUUAGCUCCAGAGUA   AT1G21590.1
     : |   | | | | | | |   | | | | | | | |
3'- UCAAUGAAUCUAGGUCUCAC   CI_cpe-2015_45087
```

**Target Site:** 2056

**Superfamily:** SupFam_257

**miRNA in ccm:** N/A

**Confirmed Targeting**

| *2nd-siRNA* | *NanoPARE* |
|---|---|
| *ccm* | *ccm* |
| *cpe-2015* | *cpe-2015* |
| *cpe-2017* | |
| *cgr-dp* | *cgr-dp* |
| *cgr-pm* | |
| *cgr-mass* | |
| *cin* | *cin* |

### Protein_kinase_superfamily_protein

AT1G21590 - CI_cpe-2015_45087

Secondary siRNAs

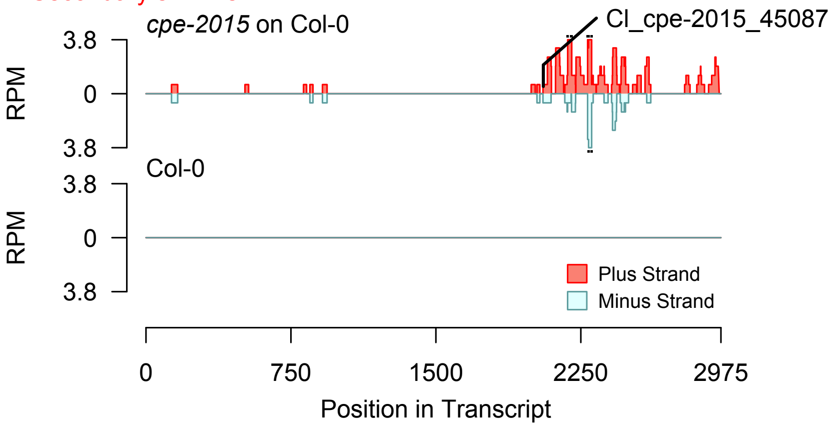


2nd siRNAs: Phase diagram

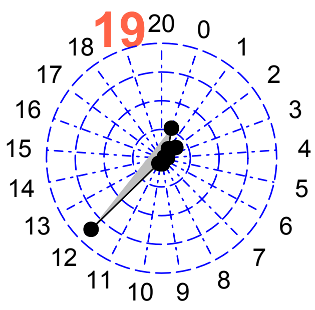


2nd siRNAs: Size distribution

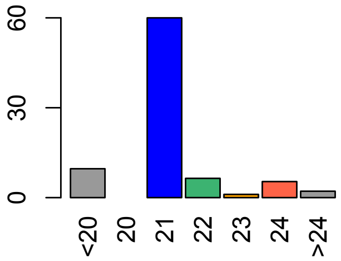


Degradome hits not found for sRNA

**Allenscore:** 4

**Target Interaction:**

```
5'- UUGGCUACUUAGCUCCAGAGUA    AT1G21590.1
      | | | |   | | | | | | | |   | | | | | | | |
3'- AACCAAUGAAUCUAGGUCUCAC    CI_cpe-2017_503
```

**Target Site:** 2056

**Superfamily:** SupFam_257

**miRNA in ccm:** N/A

**Confirmed Targeting**

| *2nd-siRNA* | *NanoPARE* |
|---|---|
| *ccm* | *ccm* |
| *cpe-2015* | *cpe-2015* |
| *cpe-2017* | |
| *cgr-dp* | *cgr-dp* |
| *cgr-pm* | |
| *cgr-mass* | |
| *cin* | *cin* |

### Protein_kinase_superfamily_protein

AT1G21590 - CI_cpe-2017_503

Secondary siRNAs

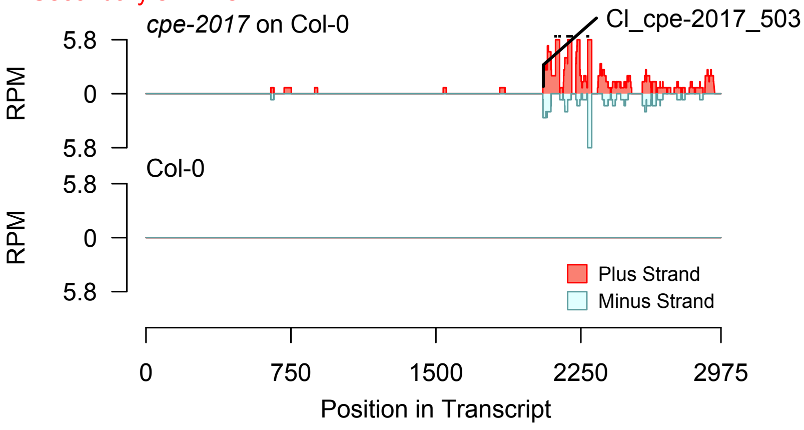


2nd siRNAs: Phase diagram

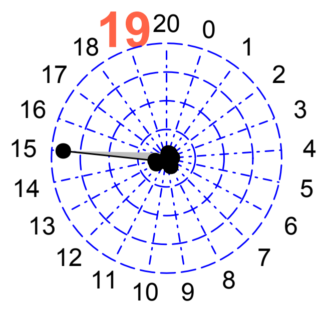


2nd siRNAs: Size distribution

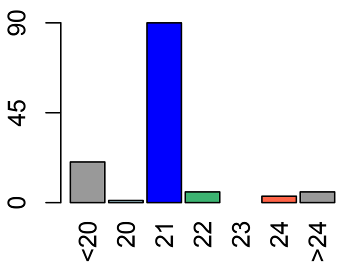


Degradome hits not found for sRNA

**Allenscore:** 2.5

**Target Interaction:**

```
5'- UAGGUUACAUGGCUCCAGAGUA   AT1G53440.1
     ||||:|||||| ||||||||||
3'- AUCCGAUGUACGGAGGUCUCAU   Cl_cgr-mass_20246
```

**Target Site:** 2600

**Superfamily:** SupFam_59

**miRNA in ccm:** N/A

**Confirmed Targeting**

| *2nd-siRNA* | *NanoPARE* |
|---|---|
| *ccm* | *ccm* |
| *cpe-2015* | *cpe-2015* |
| *cpe-2017* | |
| *cgr-dp* | *cgr-dp* |
| *cgr-pm* | |
| *cgr-mass* | |
| *cin* | *cin* |

### Leucine-rich_repeat_transmembrane_protei

AT1G53440 - Cl_cgr-mass_20246

Secondary siRNAs

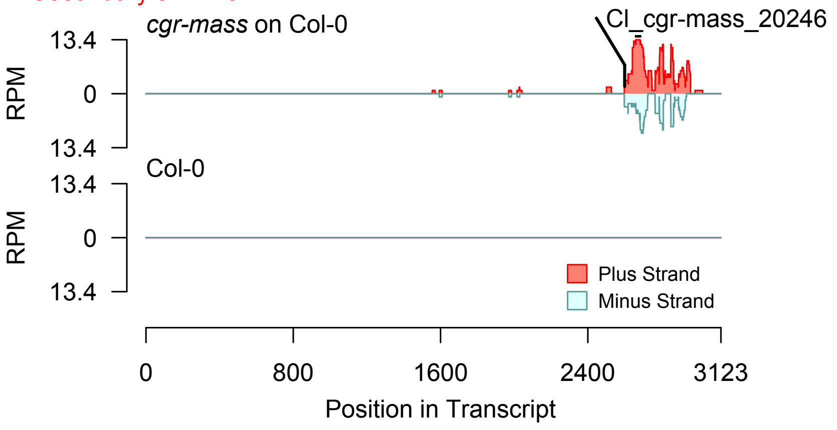


2nd siRNAs: Phase diagram

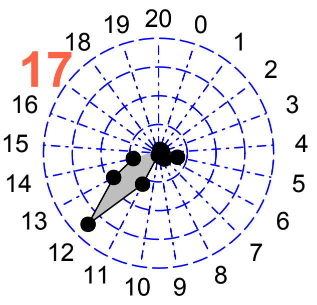


2nd siRNAs: Size distribution

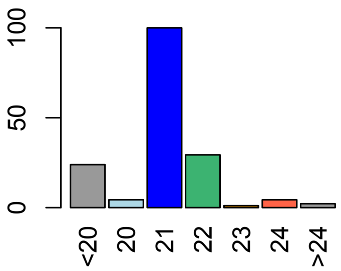


Degradome hits not found for sRNA

**Allenscore:** 4

**Target Interaction:**

```
5'- GAGAAUGGCUGGCAGAGGUGA   AT5G56460.1
     |||||||| ||||||:||||
3'- GUCUUACCGUCCGUCUUCACU   Cl_cgr-dp_8691
```

**Target Site:** 820

**Superfamily:** SupFam_4

**miRNA in ccm:** N/A

**Confirmed Targeting**

| *2nd-siRNA* | *NanoPARE* |
|---|---|
| *ccm* | *ccm* |
| *cpe-2015* | *cpe-2015* |
| *cpe-2017* | |
| *cgr-dp* | *cgr-dp* |
| *cgr-pm* | |
| *cgr-mass* | |
| *cin* | *cin* |

### Protein_kinase_superfamily_protein

AT5G56460 - Cl_cgr-dp_8691

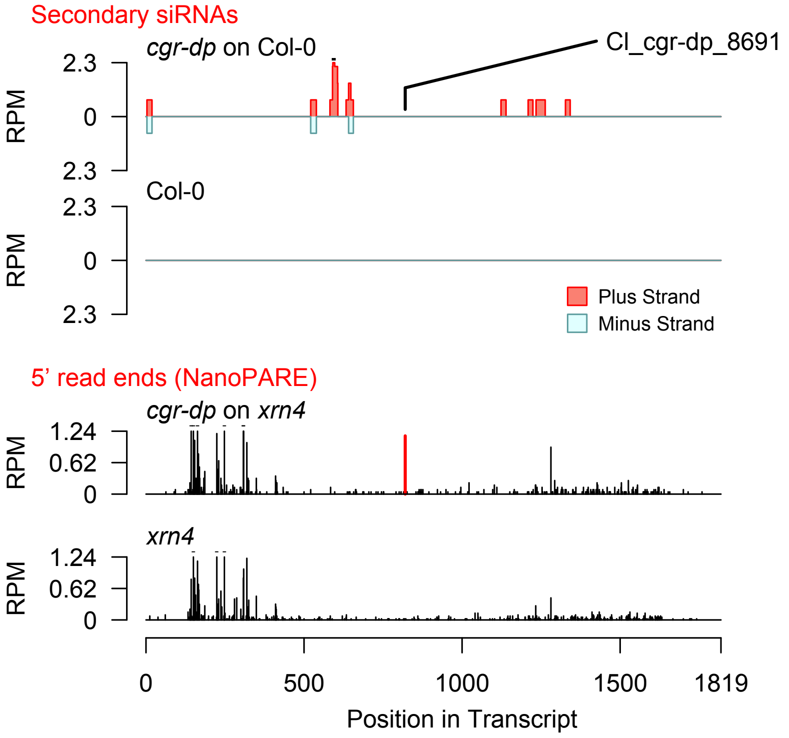


Diff. Exp. secondary siRNA locus not found

**Allenscore:** 5.5

**Target Interaction:**

```
5'- GAGAAUGGCUGGCAGAGGUGA   AT5G56460.1
     ||||||:|  ||  ||||||||
3'- GUCUUACUGUCCAUCUCCACU   CI_cgr-dp_67
```

**Target Site:** 820

**Superfamily:** SupFam_4

**miRNA in ccm:** N/A

**Confirmed Targeting**

| *2nd-siRNA* | *NanoPARE* |
|---|---|
| *ccm* | *ccm* |
| *cpe-2015* | *cpe-2015* |
| *cpe-2017* | |
| *cgr-dp* | *cgr-dp* |
| *cgr-pm* | |
| *cgr-mass* | |
| *cin* | *cin* |

### Protein_kinase_superfamily_protein

AT5G56460 - CI_cgr-dp_67

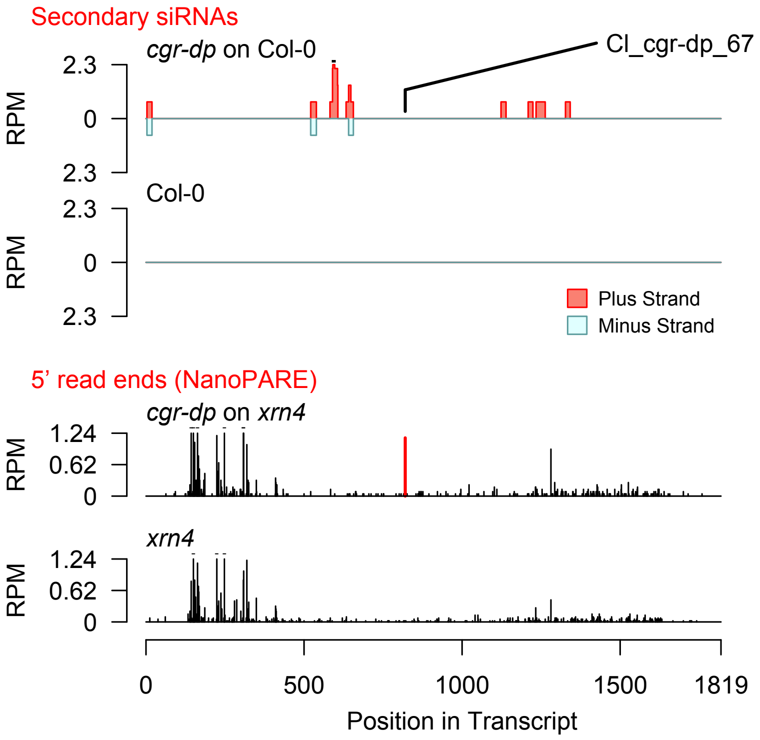


Diff. Exp. secondary siRNA locus not found

**Allenscore:** 5

**Target Interaction:**

```
5'- GAGAAUGGCUGGCAGAGGUGA   AT5G56460.1
      | | | | | | | |   | | |   | | | | | | |
3'- AUCUUACCGUCCGGCUCCACU   Cl_cgr-dp_1335
```

**Target Site:** 820

**Superfamily:** SupFam_4

**miRNA in ccm:** N/A

**Confirmed Targeting**

| *2nd-siRNA* | *NanoPARE* |
|---|---|
| *ccm* | *ccm* |
| *cpe-2015* | *cpe-2015* |
| *cpe-2017* | |
| *cgr-dp* | *cgr-dp* |
| *cgr-pm* | |
| *cgr-mass* | |
| *cin* | *cin* |

### Protein_kinase_superfamily_protein

AT5G56460 - Cl_cgr-dp_1335

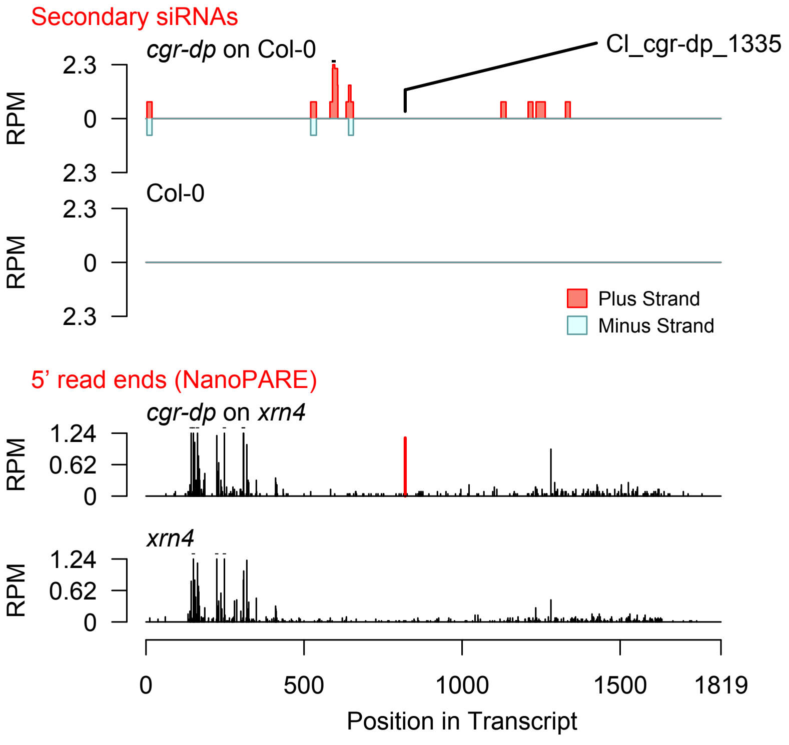


Diff. Exp. secondary siRNA locus not found

**Allenscore:** 3.5

**Target Interaction:**

```
5'- GGAGACUAGUACUAGGAGACAUAAUCA   AT3G27925.1
     | |      | | | | | | | : | | | | | | | | | | | | | |
3'- CC - - - GAUCAUGGUCCUCUGUAUUAGU   Cl_cgr-dp_83
```

**Target Site:** 1305

**Superfamily:** SupFam_33

**miRNA in ccm:** N/A

**Confirmed Targeting**

| *2nd-siRNA* | *NanoPARE* |
|---|---|
| *ccm* | *ccm* |
| *cpe-2015* | *cpe-2015* |
| *cpe-2017* | |
| *cgr-dp* | *cgr-dp* |
| *cgr-pm* | |
| *cgr-mass* | |
| *cin* | *cin* |

### DEG1

AT3G27925 - Cl_cgr-dp_83

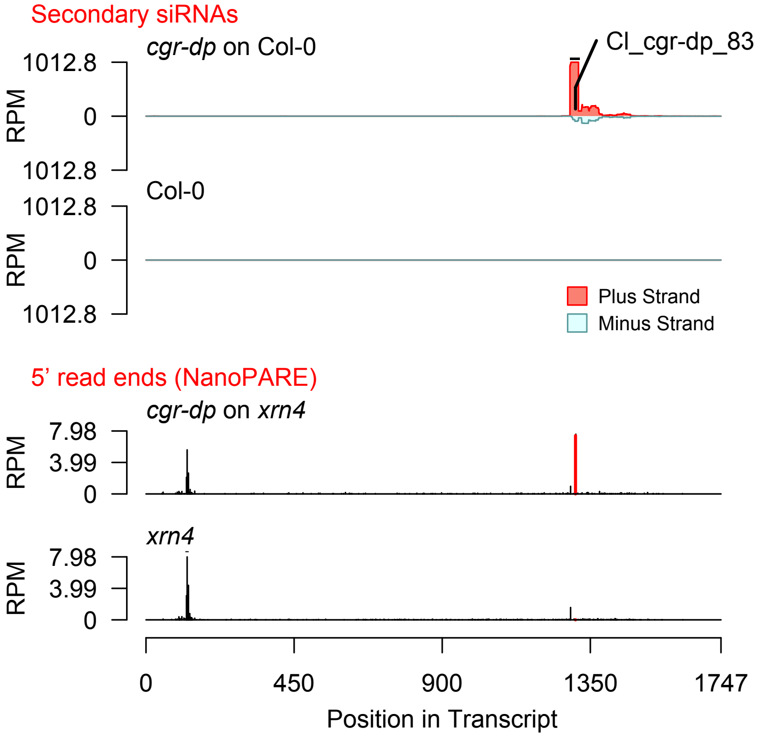


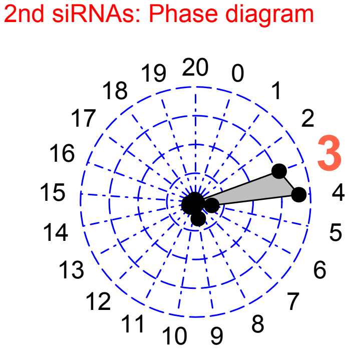


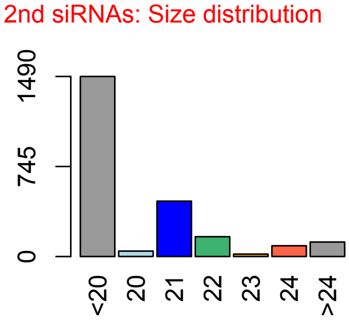

**Allenscore:** 0.5

**Target Interaction:**

```
5'- AGUACUAGGAGACAUAAUCA   AT3G27925.1
     |||||:||||||||||||||
3'- UCAUGGUCCUCUGUAUUAGU   Cl_cgr-dp_775
```

**Target Site:** 1305

**Superfamily:** SupFam_33

**miRNA in ccm:** N/A

**Confirmed Targeting**

| *2nd-siRNA* | *NanoPARE* |
|---|---|
| *ccm* | *ccm* |
| *cpe-2015* | *cpe-2015* |
| *cpe-2017* | |
| *cgr-dp* | *cgr-dp* |
| *cgr-pm* | |
| *cgr-mass* | |
| *cin* | *cin* |

### DEG1

AT3G27925 - Cl_cgr-dp_775

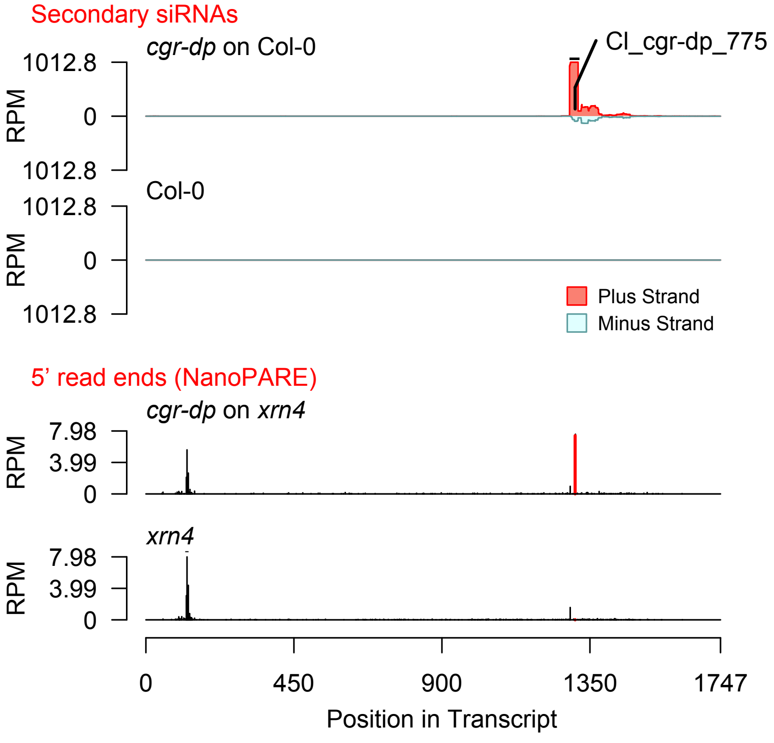


2nd siRNAs: Phase diagram

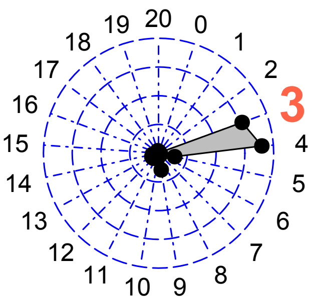


2nd siRNAs: Size distribution

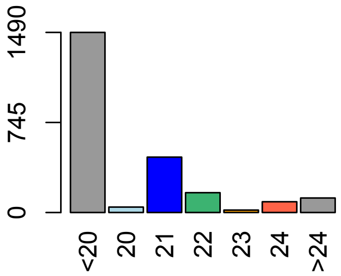

**Allenscore:** 3.5

**Target Interaction:**

```
5'- GGAGACUAGUACUAGGAGACAUAAUCA    AT3G27925.1
     | |       | | | | | | | : | | | | | | | | | | | | | |
3'- CC - - - GAUCAUGGUCCUCUGUAUUAGU    CI_cgr-pm_154
```

**Target Site:** 1305

**Superfamily:** SupFam_33

**miRNA in ccm:** N/A

**Confirmed Targeting**

| *2nd-siRNA* | *NanoPARE* |
|---|---|
| *ccm* | *ccm* |
| *cpe-2015* | *cpe-2015* |
| *cpe-2017* | |
| *cgr-dp* | *cgr-dp* |
| *cgr-pm* | |
| *cgr-mass* | |
| *cin* | *cin* |

### DEG1

AT3G27925 - CI_cgr-pm_154

Secondary siRNAs

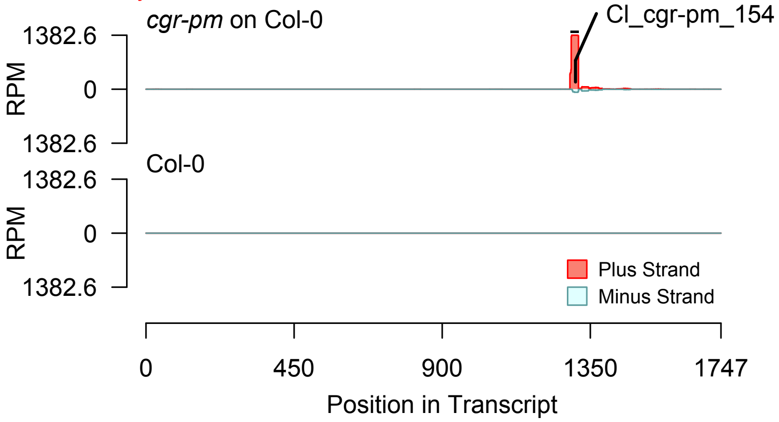


Degradome hits not found for sRNA

2nd siRNAs: Phase diagram

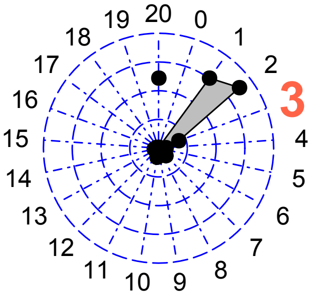


2nd siRNAs: Size distribution

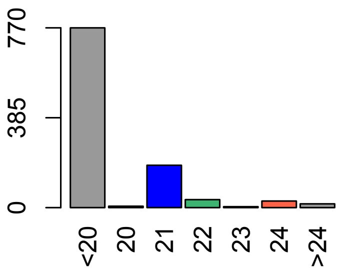

**Allenscore:** 1.5

**Target Interaction:**

```
5'- CUAGUACUAGGAGACAUAAUCA    AT3G27925.1
      | | | | | | : | | | | | | | | | | | | | |
3'- AAUCAUGGUCCUCUGUAUUAGU    Cl_cgr-pm_456
```

**Target Site:** 1305

**Superfamily:** SupFam_33

**miRNA in ccm:** N/A

**Confirmed Targeting**

| *2nd-siRNA* | *NanoPARE* |
|---|---|
| *ccm* | *ccm* |
| *cpe-2015* | *cpe-2015* |
| *cpe-2017* | |
| *cgr-dp* | *cgr-dp* |
| *cgr-pm* | |
| *cgr-mass* | |
| *cin* | *cin* |

### DEG1

AT3G27925 - Cl_cgr-pm_456

Secondary siRNAs

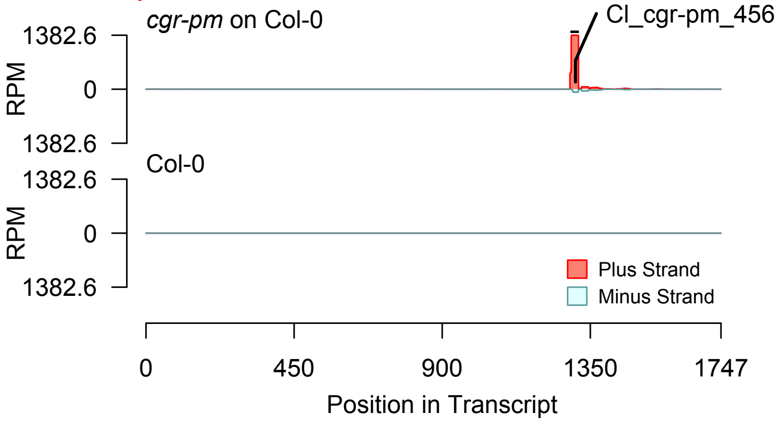


Degradome hits not found for sRNA

2nd siRNAs: Phase diagram

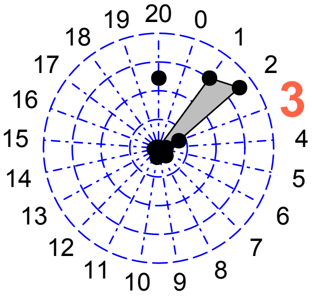


2nd siRNAs: Size distribution

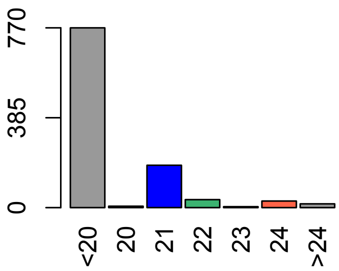

**Allenscore:** 1.5

**Target Interaction:**

5'- AGUACUAGGAGACAUAAUCA    AT3G27925.1
     | | | | : | | | | | | | | | | | | | |
3'- CCAUGGUCCUCUGUAUUAGU    Cl_cgr-pm_11629

**Target Site:** 1305

**Superfamily:** SupFam_33

**miRNA in ccm:** N/A

**Confirmed Targeting**

| *2nd-siRNA* | *NanoPARE* |
|---|---|
| *ccm* | *ccm* |
| *cpe-2015* | *cpe-2015* |
| *cpe-2017* | |
| *cgr-dp* | *cgr-dp* |
| *cgr-pm* | |
| *cgr-mass* | |
| *cin* | *cin* |

### DEG1

AT3G27925 - Cl_cgr-pm_11629

Secondary siRNAs

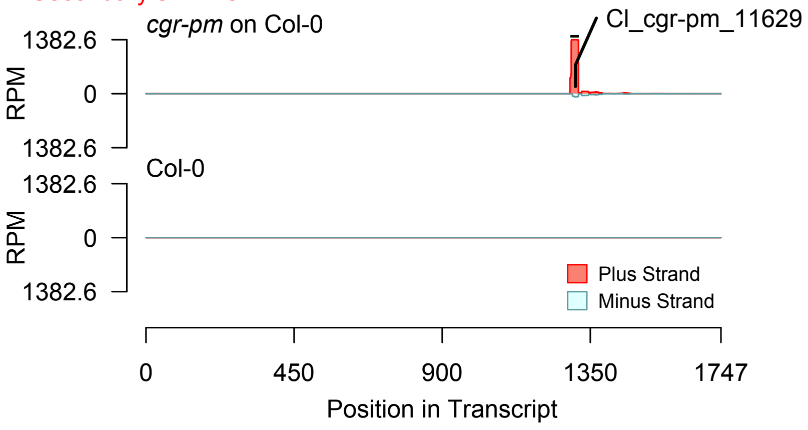


2nd siRNAs: Phase diagram

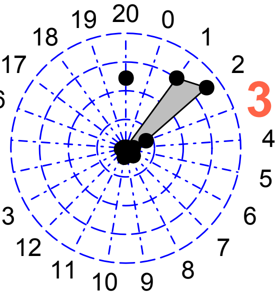


2nd siRNAs: Size distribution

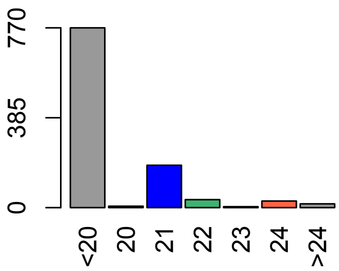


Degradome hits not found for sRNA

**Allenscore:** 3.5

**Target Interaction:**

```
5'- GGAGACUAGUACUAGGAGACAUAAUCA   AT3G27925.1
     | |      | | | | | | | : | | | | | | | | | | | | | |
3'- CC - - - GAUCAUGGUCCUCUGUAUUAGU   CI_cgr-mass_424
```

**Target Site:** 1305

**Superfamily:** SupFam_33

**miRNA in ccm:** N/A

**Confirmed Targeting**

| *2nd-siRNA* | *NanoPARE* |
|---|---|
| *ccm* | *ccm* |
| *cpe-2015* | *cpe-2015* |
| *cpe-2017* | |
| *cgr-dp* | *cgr-dp* |
| *cgr-pm* | |
| *cgr-mass* | |
| *cin* | *cin* |

### DEG1

AT3G27925 - CI_cgr-mass_424

2nd siRNAs: Phase diagram

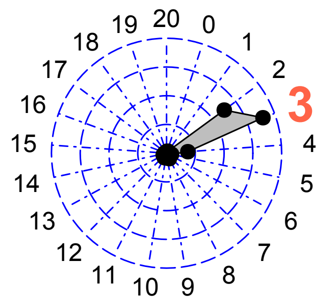


Secondary siRNAs

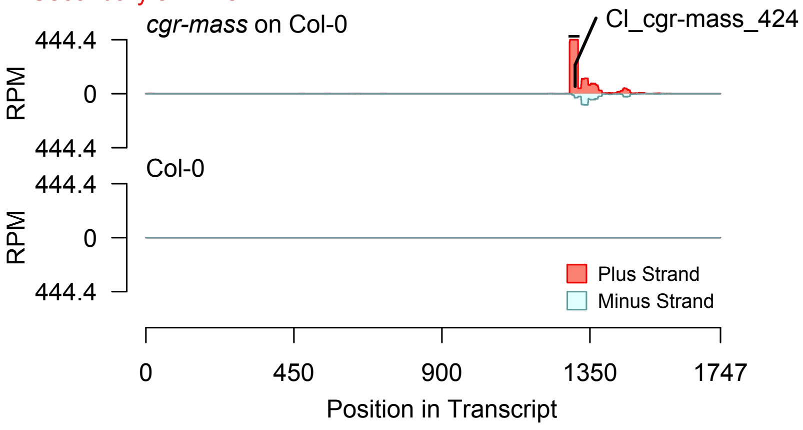


2nd siRNAs: Size distribution

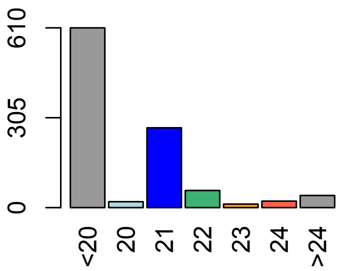


Degradome hits not found for sRNA

**Allenscore:** 0.5

**Target Interaction:**

```
5'- AGUACUAGGAGACAUAAUCA   AT3G27925.1
     | | | | | : | | | | | | | | | | | | | |
3'- UCAUGGUCCUCUGUAUUAGU   Cl_cgr-mass_5502
```

**Target Site:** 1305

**Superfamily:** SupFam_33

**miRNA in ccm:** N/A

**Confirmed Targeting**

| *2nd-siRNA* | *NanoPARE* |
|---|---|
| *ccm* | *ccm* |
| *cpe-2015* | *cpe-2015* |
| *cpe-2017* | |
| *cgr-dp* | *cgr-dp* |
| *cgr-pm* | |
| *cgr-mass* | |
| *cin* | *cin* |

### DEG1

AT3G27925 - Cl_cgr-mass_5502

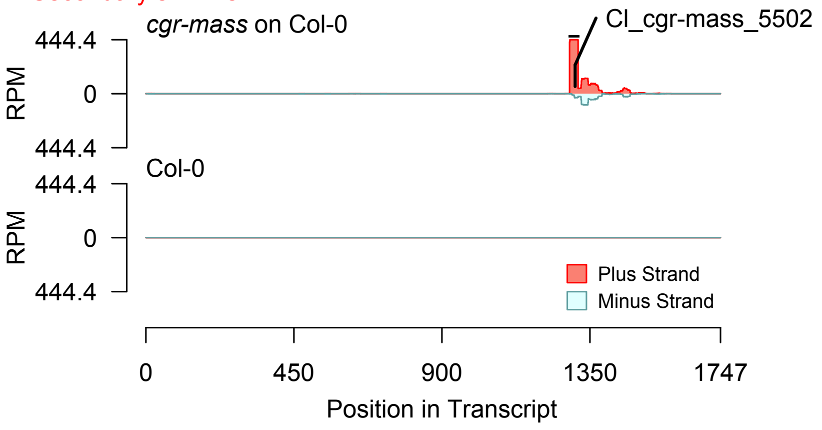


2nd siRNAs: Phase diagram

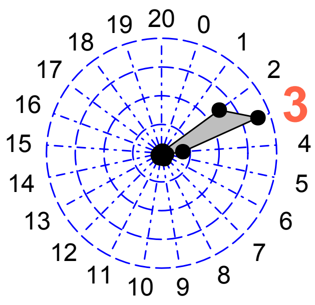


2nd siRNAs: Size distribution

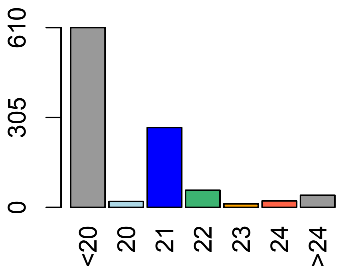


Degradome hits not found for sRNA

**Allenscore:** 4.5

**Target Interaction:**

```
5'- UGUGGUUACAUUGCACCAGAAUA   AT4G28490.1
     : :   | | | | | |   : | | | | | | | | | | | | |
3'- GUCCCAAUGCGACGUGGUCUUAU   CI_ccm_818
```

**Target Site:** 2815

**Superfamily:** SupFam_124

**miRNA in ccm:** N

**Confirmed Targeting**

| *2nd-siRNA* | *NanoPARE* |
|---|---|
| *ccm* | *ccm* |
| *cpe-2015* | *cpe-2015* |
| *cpe-2017* | |
| *cgr-dp* | *cgr-dp* |
| *cgr-pm* | |
| *cgr-mass* | |
| *cin* | *cin* |

### HAE

AT4G28490 - CI_ccm_818

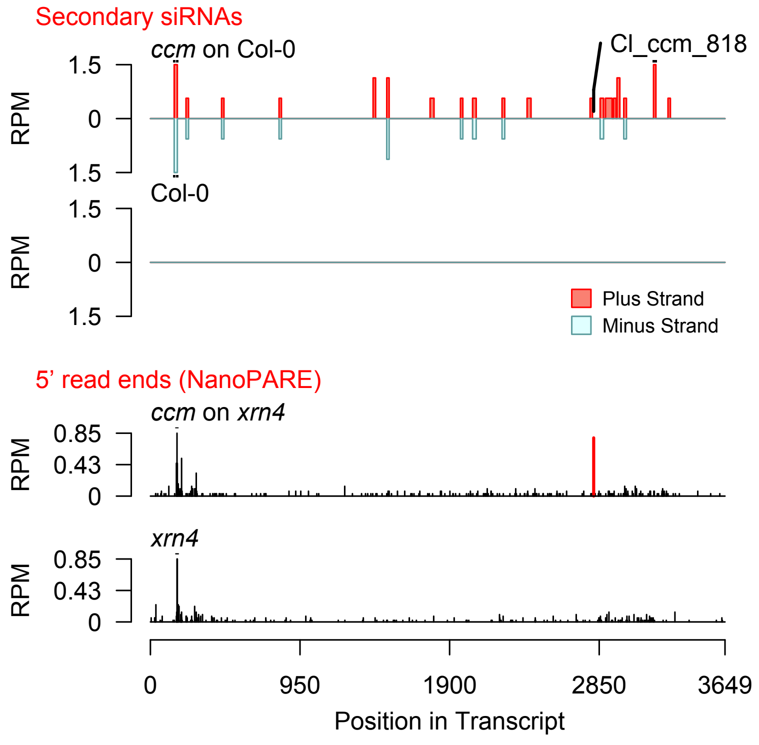


Diff. Exp. secondary siRNA locus not found

**Allenscore:** 5

**Target Interaction:**

5'- UAAACGCCAAGAGGAACUUCC AT5G46580.1
|  |  |  |  |  |  |  |  |  |  | : |  |  |  |  |  |  |
3'- UUUUGCGGUGCUCUUUGAAGU Cl_cgr-dp_29

**Target Site:** 1929

**Superfamily:** SupFam_26

**miRNA in ccm:** Y

**Confirmed Targeting**

| *2nd-siRNA* | *NanoPARE* |
|---|---|
| *ccm* | *ccm* |
| *cpe-2015* | *cpe-2015* |
| *cpe-2017* | |
| *cgr-dp* | *cgr-dp* |
| *cgr-pm* | |
| *cgr-mass* | |
| *cin* | *cin* |

### PPR_repeat-containing_protein

AT5G46580 - Cl_cgr-dp_29

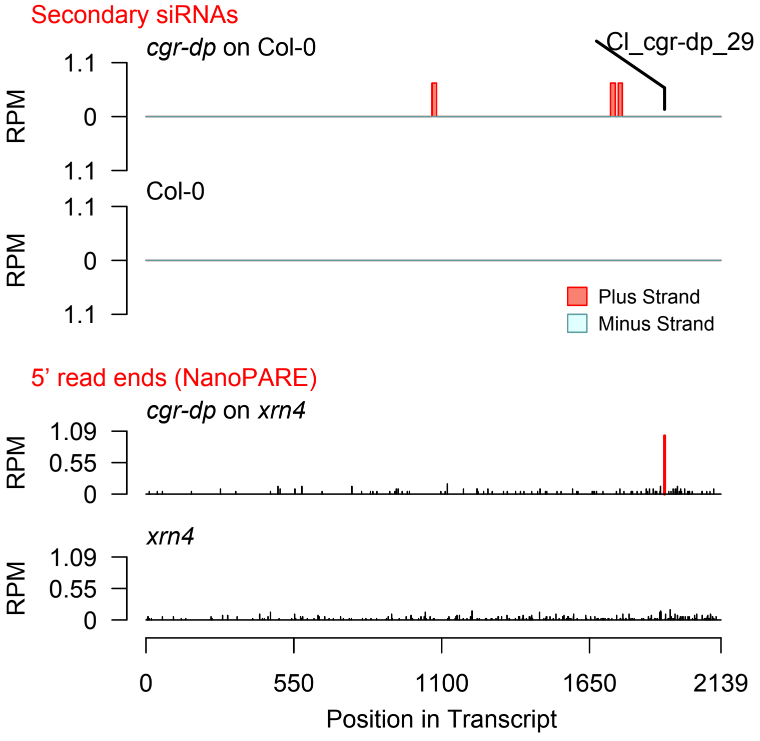


Diff. Exp. secondary siRNA locus not found

**Allenscore:** 4.5

**Target Interaction:**

5'- AAAGAUGCUACGACUCGUUCA AT3G52610.1
| | | | : | | | | | | | | | | | | | |
3'- AUCCUAUGAUGCUGAGCAAGC Cl_ccm_205

**Target Site:** 1115

**Superfamily:** SupFam_18

**miRNA in ccm:** Y

**Confirmed Targeting**

| *2nd-siRNA* | *NanoPARE* |
|---|---|
| *ccm* | *ccm* |
| *cpe-2015* | *cpe-2015* |
| *cpe-2017* | |
| *cgr-dp* | *cgr-dp* |
| *cgr-pm* | |
| *cgr-mass* | |
| *cin* | *cin* |

### GATA_zinc_finger_protein

AT3G52610 - Cl_ccm_205

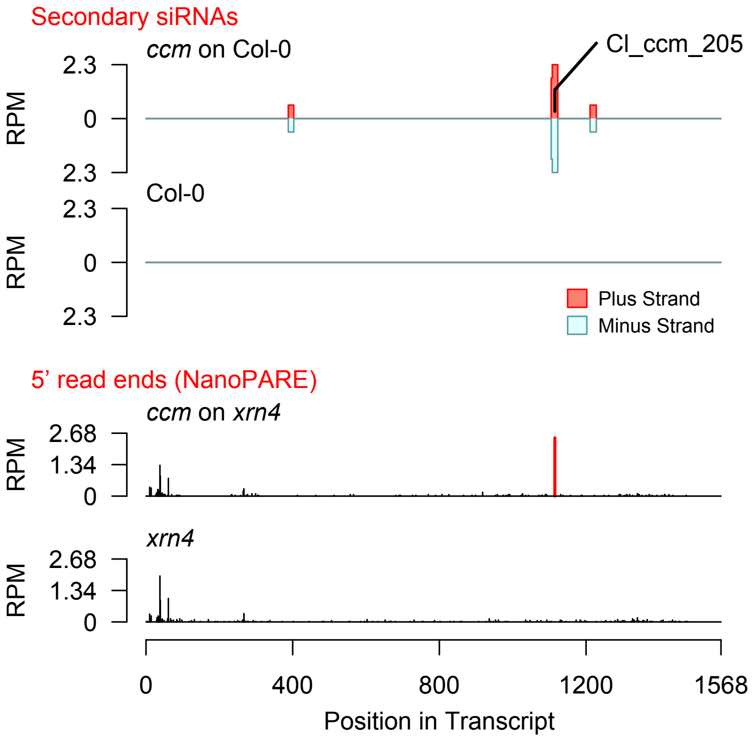


Diff. Exp. secondary siRNA locus not found

**Allenscore:** 2.5

**Target Interaction:**

```
5'- UAUAGUUGAAGCAGGAAUUGCG   AT2G46110.1
    : | | | | : | | | | | | | | | | | | : | | :
3'- GUAUCGACUUCGUCCUUAGCGU   CI_ccm_30
```

**Target Site:** 681

**Superfamily:** SupFam_50

**miRNA in ccm:** Y

**Confirmed Targeting**

| *2nd-siRNA* | *NanoPARE* |
|---|---|
| *ccm* | *ccm* |
| *cpe-2015* | *cpe-2015* |
| *cpe-2017* | |
| *cgr-dp* | *cgr-dp* |
| *cgr-pm* | |
| *cgr-mass* | |
| *cin* | *cin* |

### KPHMT1

AT2G46110 - CI_ccm_30

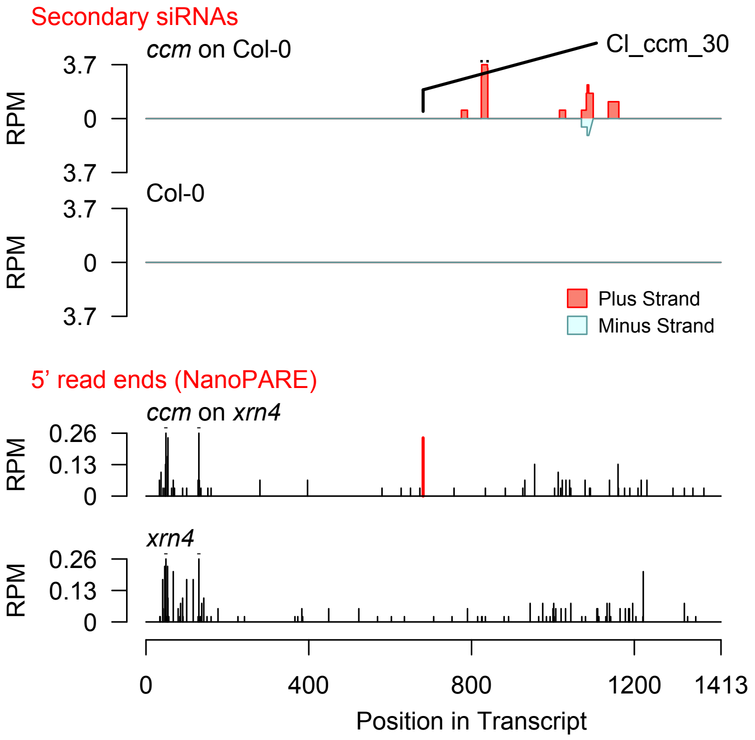


Diff. Exp. secondary siRNA locus not found

**Allenscore:** 5

**Target Interaction:**

```
5'- ACAGGAGAAGAAGGAAGACCA   AT1G18540.1
     ||||||||||  ||||||||  |
3'- UGUCCUCUUCAUCCUUCUGAU   CI_cin_11140
```

**Target Site:** 607

**Superfamily:** SupFam_1431

**miRNA in ccm:** N/A

**Confirmed Targeting**

| *2nd-siRNA* | *NanoPARE* |
|---|---|
| *ccm* | *ccm* |
| *cpe-2015* | *cpe-2015* |
| *cpe-2017* | |
| *cgr-dp* | *cgr-dp* |
| *cgr-pm* | |
| *cgr-mass* | |
| *cin* | *cin* |

### Ribosomal_protein_L6_family_protein

AT1G18540 - CI_cin_11140

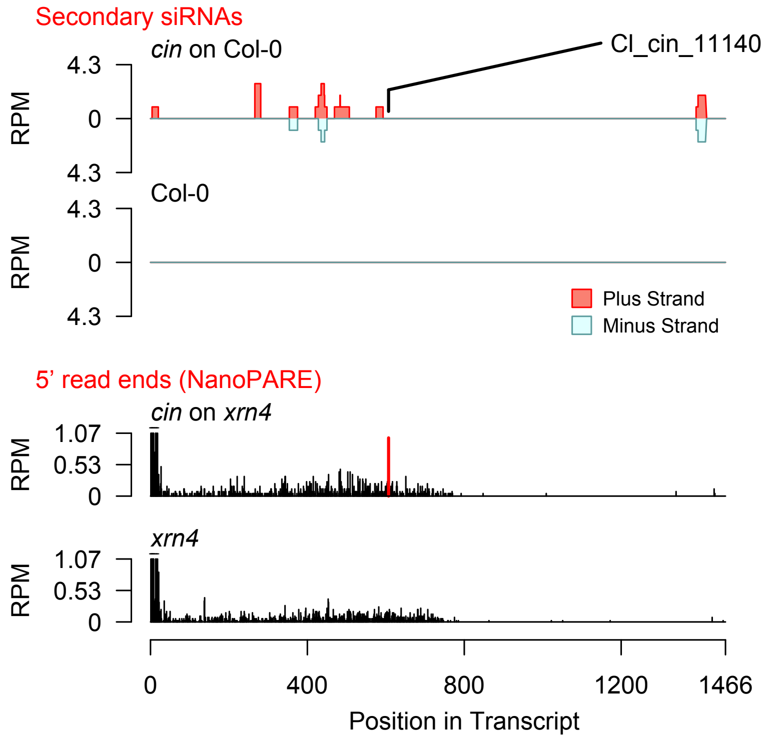


Diff. Exp. secondary siRNA locus not found

**Allenscore:** 0.5

**Target Interaction:**

5'- GAUUGCUGUUAAGAAACUGAA AT2G17220.1
:||||||||||||||||||||
3'- UUAACGACAAUUCUUUGACUU CI_ccm_9385

**Target Site:** 799

**Superfamily:** SupFam_320

**miRNA in ccm:** Y

**Confirmed Targeting**

| *2nd-siRNA* | *NanoPARE* |
|---|---|
| *ccm* | *ccm* |
| *cpe-2015* | *cpe-2015* |
| *cpe-2017* | |
| *cgr-dp* | *cgr-dp* |
| *cgr-pm* | |
| *cgr-mass* | |
| *cin* | *cin* |

### Kin3

AT2G17220 - CI_ccm_9385

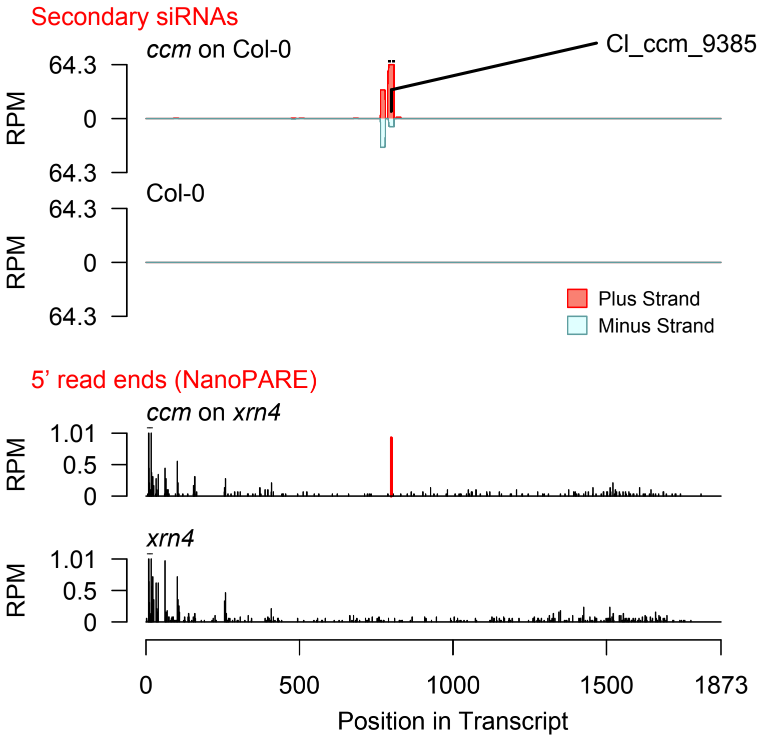


Diff. Exp. secondary siRNA locus not found

**Allenscore:** 2

**Target Interaction:**

```
5'- UCGGGAUGCUGAAGAUGCUA   AT1G02840.1
        | | | | | | | | | | | | | | | | | |
3'- CUCCCUACGACUUCUACGAU   CI_cin_3533
```

**Target Site:** 370

**Superfamily:** SupFam_742

**miRNA in ccm:** N/A

**Confirmed Targeting**

| *2nd-siRNA* | *NanoPARE* |
|---|---|
| *ccm* | *ccm* |
| *cpe-2015* | *cpe-2015* |
| *cpe-2017* | |
| *cgr-dp* | *cgr-dp* |
| *cgr-pm* | |
| *cgr-mass* | |
| *cin* | *cin* |

# SR34

AT1G02840 - CI_cin_3533

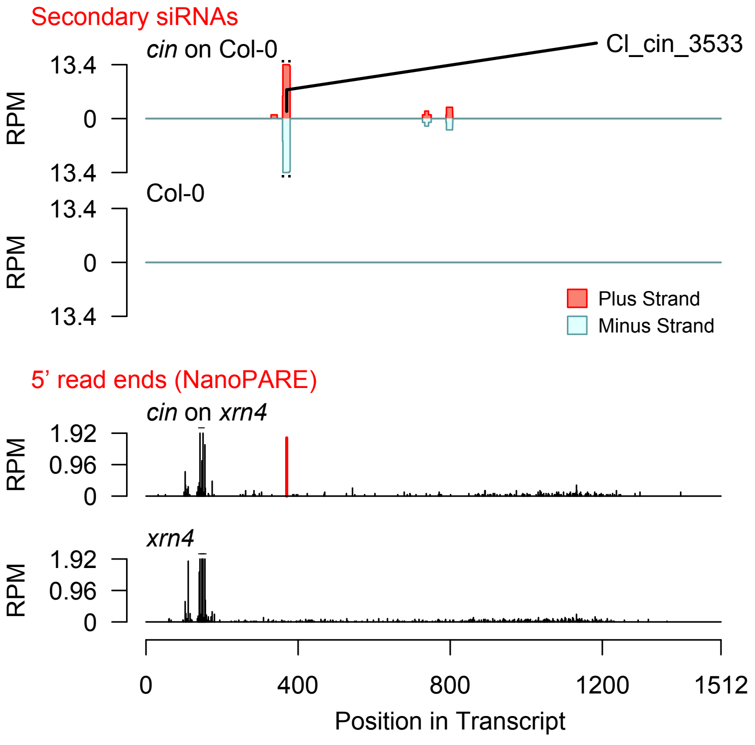


Diff. Exp. secondary siRNA locus not found

**Allenscore:** 3

**Target Interaction:**

5'- AGGAAGAUUGUGCAAAGAGAUA  AT4G01650.1
|  |  |  |  |  |  |  |  |  |  |  | : |  |  |  |  |  |  |  |  |
3'- UCCUUCUAACAUGUGUCUCUAU  CI_ccm_6791

**Target Site:** 1000

**Superfamily:** SupFam_134

**miRNA in ccm:** Y

**Confirmed Targeting**

| *2nd-siRNA* | *NanoPARE* |
|---|---|
| *ccm* | *ccm* |
| *cpe-2015* | *cpe-2015* |
| *cpe-2017* | |
| *cgr-dp* | *cgr-dp* |
| *cgr-pm* | |
| *cgr-mass* | |
| *cin* | *cin* |

### Polyketide_cyclase_dehydrase_and_lipid_t

AT4G01650 - CI_ccm_6791

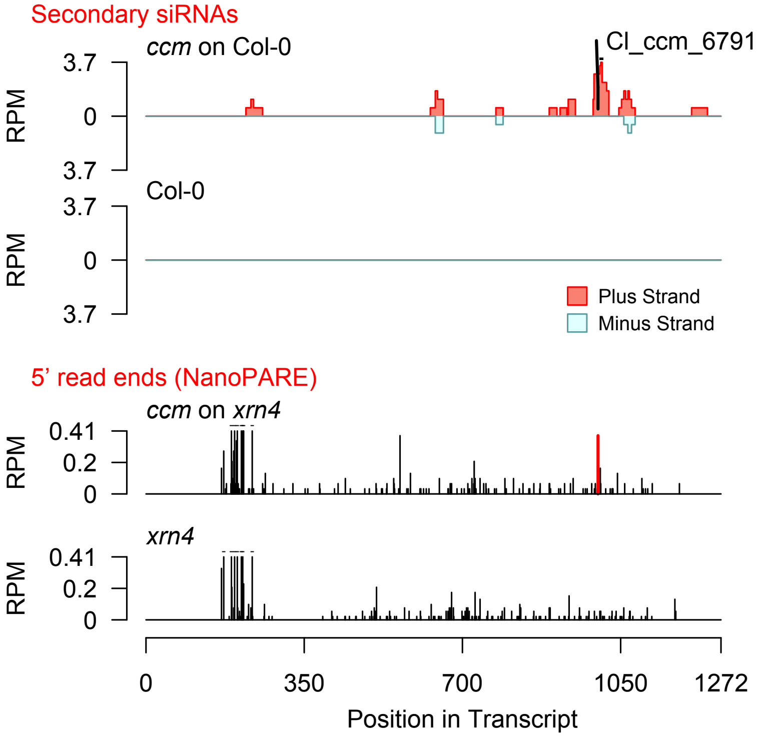


Diff. Exp. secondary siRNA locus not found

**Allenscore:** 3

**Target Interaction:**

```
5'- UGGACUUGUGAUGGAGGAGAUAA    AT3G46740.1
     | : |   | | | | | : | | | | | : | | | | | | |
3'- AUC - GAACAUUACCUUCUCUAUU    CI_cin_10092
```

**Target Site:** 1871

**Superfamily:** SupFam_1423

**miRNA in ccm:** N/A

**Confirmed Targeting**

| *2nd-siRNA* | *NanoPARE* |
|---|---|
| *ccm* | *ccm* |
| *cpe-2015* | *cpe-2015* |
| *cpe-2017* | |
| *cgr-dp* | *cgr-dp* |
| *cgr-pm* | |
| *cgr-mass* | |
| *cin* | *cin* |

### TOC75-III

AT3G46740 - CI_cin_10092

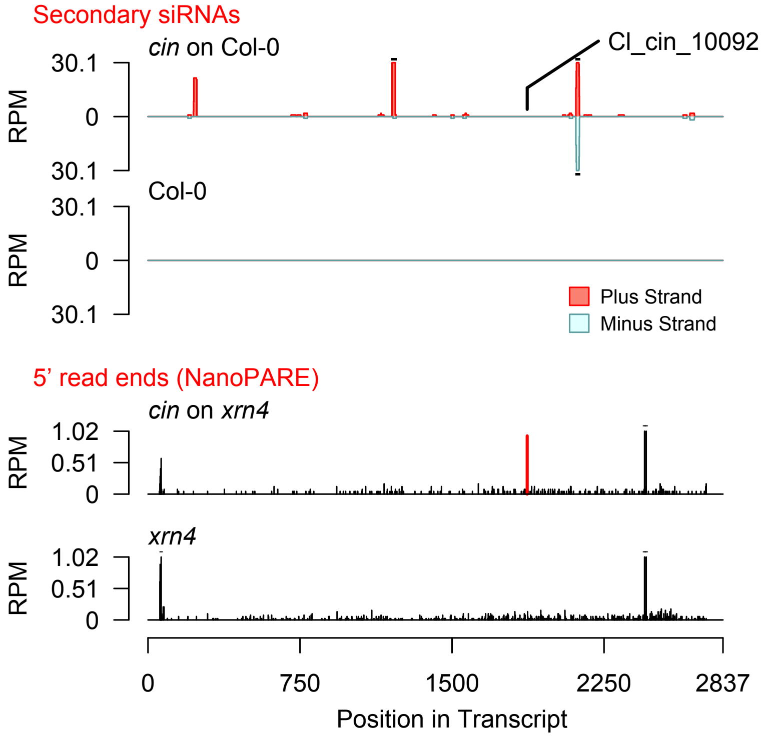


Diff. Exp. secondary siRNA locus not found

**Allenscore:** 5

**Target Interaction:**

```
5'- AAGAAUGACAGGAGGAGGUGC   AT5G24760.1
     |||||||||||  :||||||
3'- GUCUUACUGUCCAUCUCCACU   Cl_cgr-dp_67
```

**Target Site:** 1124

**Superfamily:** SupFam_4

**miRNA in ccm:** N/A

**Confirmed Targeting**

| *2nd-siRNA* | *NanoPARE* |
|---|---|
| *ccm* | *ccm* |
| *cpe-2015* | *cpe-2015* |
| *cpe-2017* | |
| *cgr-dp* | *cgr-dp* |
| *cgr-pm* | |
| *cgr-mass* | |
| *cin* | *cin* |

### GroES-like_zinc-binding_dehydrogenase_fa

AT5G24760 - Cl_cgr-dp_67

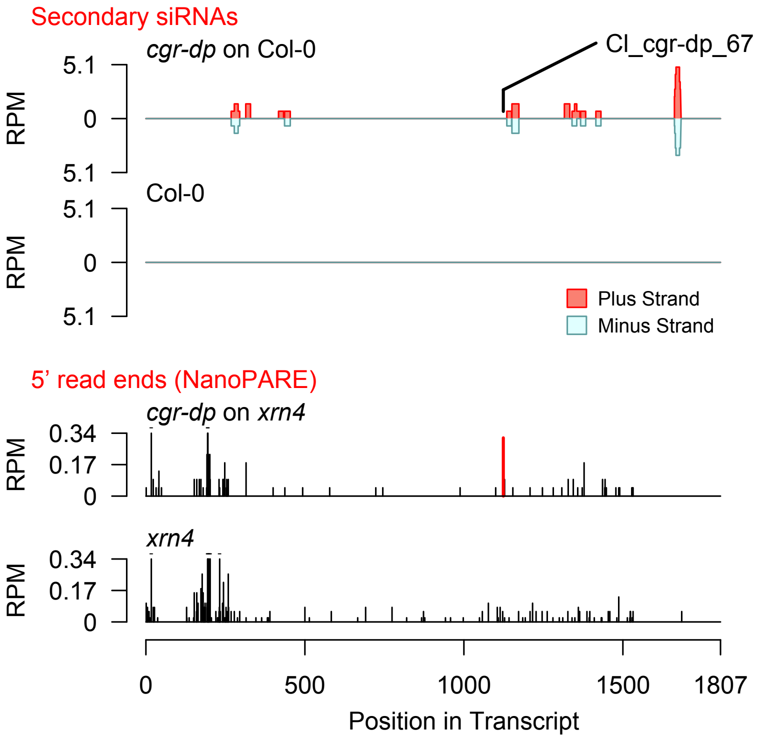


Diff. Exp. secondary siRNA locus not found

**Allenscore:** 3.5

**Target Interaction:**

```
5'- CUGGUCUGCAAAUCAUGGCACA   AT1G12820.1
     ||  |||||:|||||||||||  |||
3'- GAGCAGAUGUUUAGUACCAUGU   CI_ccm_2
```

**Target Site:** 472

**Superfamily:** SupFam_27

**miRNA in ccm:** Y

**Confirmed Targeting**

| *2nd-siRNA* | *NanoPARE* |
|---|---|
| *ccm* | *ccm* |
| *cpe-2015* | *cpe-2015* |
| *cpe-2017* | |
| *cgr-dp* | *cgr-dp* |
| *cgr-pm* | |
| *cgr-mass* | |
| *cin* | *cin* |

### AFB3

AT1G12820 - CI_ccm_2

Secondary siRNAs

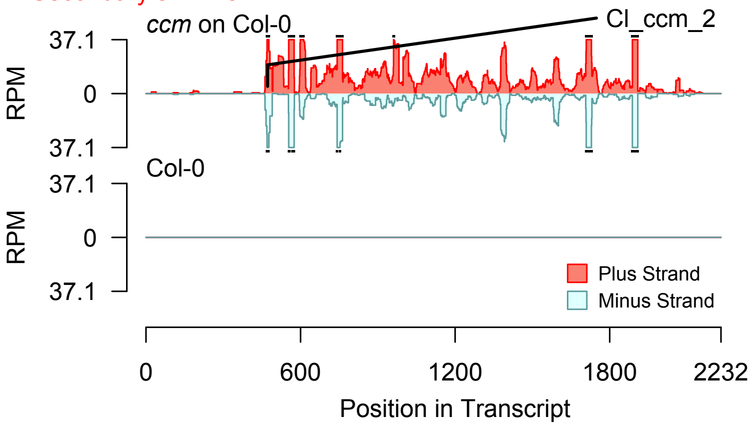


2nd siRNAs: Phase diagram

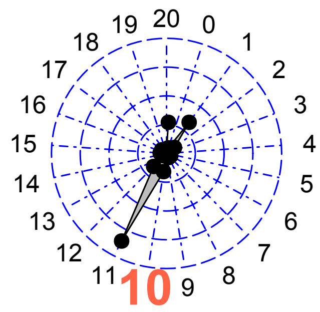


2nd siRNAs: Size distribution

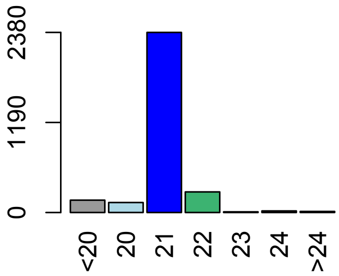


Degradome hits not found for sRNA

**Allenscore:** 4

**Target Interaction:**

```
5'- UCUGGUCUGCAAAUCAUGGC   AT1G12820.1
      ||  ||||:||||||||||
3'- UGAGCAGAUGUUUAGUACCA   CI_ccm_98
```

**Target Site:** 469

**Superfamily:** SupFam_27

**miRNA in ccm:** Y

**Confirmed Targeting**

| *2nd-siRNA* | *NanoPARE* |
|---|---|
| *ccm* | *ccm* |
| *cpe-2015* | *cpe-2015* |
| *cpe-2017* | |
| *cgr-dp* | *cgr-dp* |
| *cgr-pm* | |
| *cgr-mass* | |
| *cin* | *cin* |

### AFB3

AT1G12820 - CI_ccm_98

Secondary siRNAs

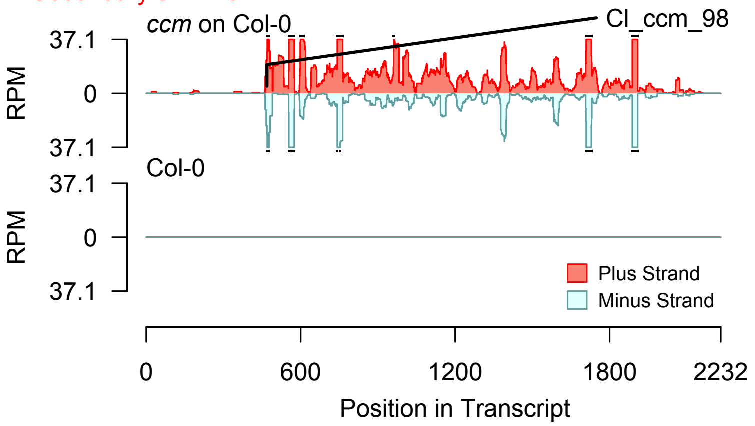


Degradome hits not found for sRNA

2nd siRNAs: Phase diagram

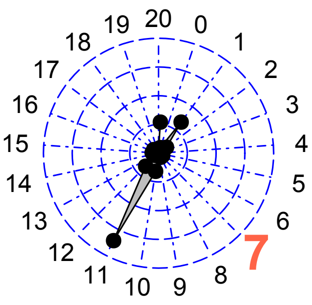


2nd siRNAs: Size distribution

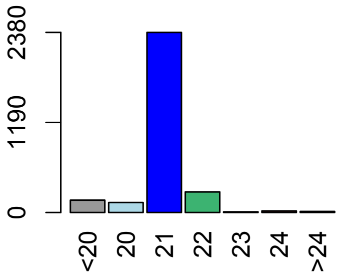

**Allenscore:** 4

**Target Interaction:**

```
5'- CUGGUCUGCAAAUCAUGGCACA   AT1G12820.1
     ||||  |||||||||||||  |||
3'- AACCAUACGUUUAGUACCAUGU   Cl_ccm_614
```

**Target Site:** 472

**Superfamily:** SupFam_27

**miRNA in ccm:** Y

**Confirmed Targeting**

| *2nd-siRNA* | *NanoPARE* |
|---|---|
| *ccm* | *ccm* |
| *cpe-2015* | *cpe-2015* |
| *cpe-2017* | |
| *cgr-dp* | *cgr-dp* |
| *cgr-pm* | |
| *cgr-mass* | |
| *cin* | *cin* |

### AFB3

AT1G12820 - Cl_ccm_614

Secondary siRNAs

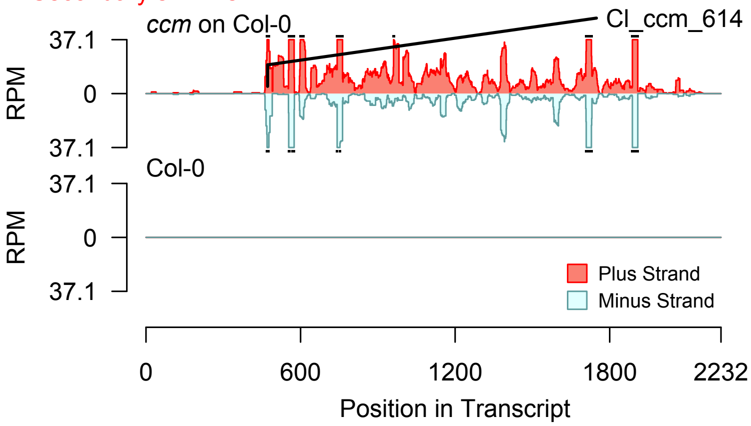


2nd siRNAs: Phase diagram

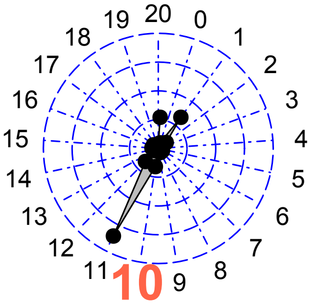


2nd siRNAs: Size distribution

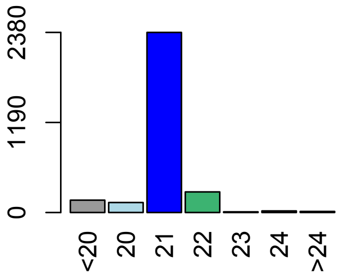


Degradome hits not found for sRNA

**Allenscore:** 5

**Target Interaction:**

```
5'- CAGAUCCUGAAU-CUUUUGCA   AT1G12820.1
     ||||||  ||||  |||||:||
3'- AUCUAGGUCUUACGAAAAUGU   CI_ccm_2562
```

**Target Site:** 1123

**Superfamily:** SupFam_196

**miRNA in ccm:** Y

**Confirmed Targeting**

| *2nd-siRNA* | *NanoPARE* |
|---|---|
| *ccm* | *ccm* |
| *cpe-2015* | *cpe-2015* |
| *cpe-2017* | |
| *cgr-dp* | *cgr-dp* |
| *cgr-pm* | |
| *cgr-mass* | |
| *cin* | *cin* |

### AFB3

AT1G12820 - CI_ccm_2562

Secondary siRNAs

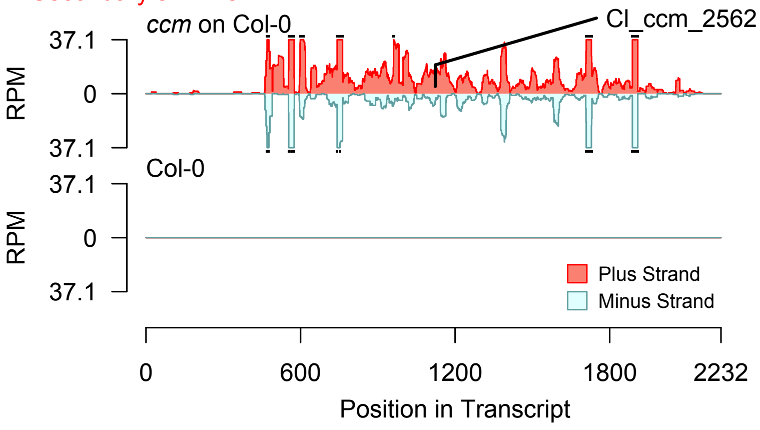


2nd siRNAs: Phase diagram

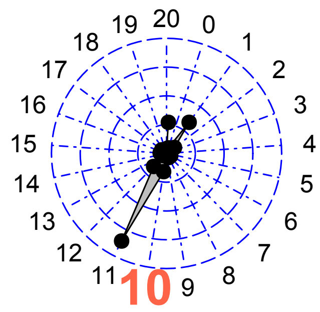


2nd siRNAs: Size distribution

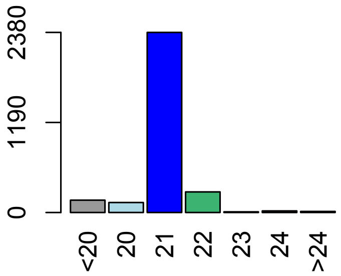


Degradome hits not found for sRNA

**Allenscore:** 3

**Target Interaction:**

```
5'- CUGGUCUGCAAAUCAUGGCACA   AT1G12820.1
    ||  |||||||||||||||  |||
3'- GAGCAGACGUUUAGUACCAUGU   CI_cpe-2015_7
```

**Target Site:** 472

**Superfamily:** SupFam_27

**miRNA in ccm:** Y

**Confirmed Targeting**

| *2nd-siRNA* | *NanoPARE* |
|---|---|
| *ccm* | *ccm* |
| *cpe-2015* | *cpe-2015* |
| *cpe-2017* | |
| *cgr-dp* | *cgr-dp* |
| *cgr-pm* | |
| *cgr-mass* | |
| *cin* | *cin* |

### AFB3

AT1G12820 - CI_cpe-2015_7

Secondary siRNAs

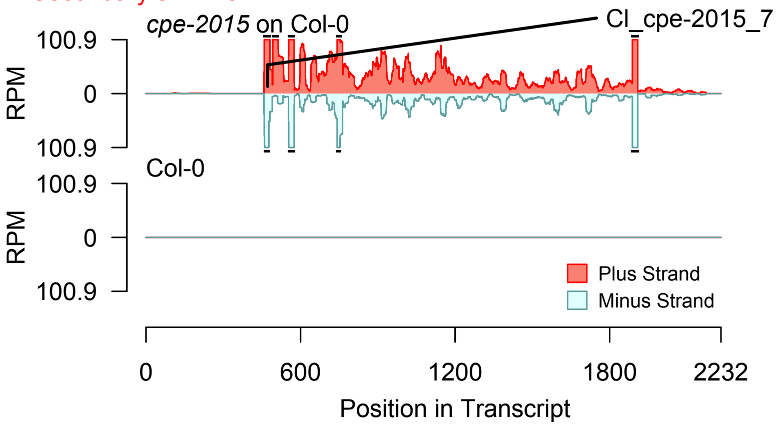


2nd siRNAs: Phase diagram

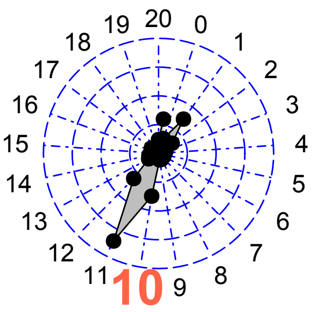


2nd siRNAs: Size distribution

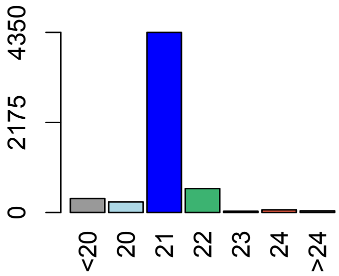


Degradome hits not found for sRNA

**Allenscore:** 3

**Target Interaction:**

```
5'- CUGGUCUGCAAAUCAUGGCACA   AT1G12820.1
    || |||||||||||||||| |||
3'- GAGCAGACGUUUAGUACCAUGU   Cl_cpe-2017_11
```

**Target Site:** 472

**Superfamily:** SupFam_27

**miRNA in ccm:** Y

**Confirmed Targeting**

| *2nd-siRNA* | *NanoPARE* |
|---|---|
| *ccm* | *ccm* |
| *cpe-2015* | *cpe-2015* |
| *cpe-2017* | |
| *cgr-dp* | *cgr-dp* |
| *cgr-pm* | |
| *cgr-mass* | |
| *cin* | *cin* |

### AFB3

AT1G12820 - Cl_cpe-2017_11

Secondary siRNAs

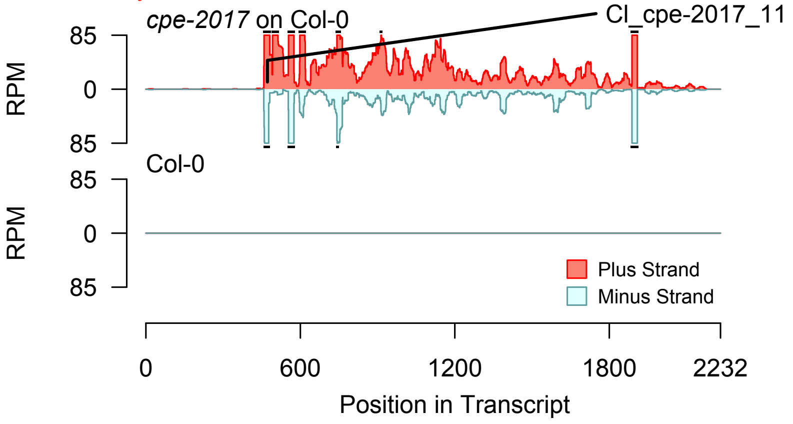


Degradome hits not found for sRNA

2nd siRNAs: Phase diagram

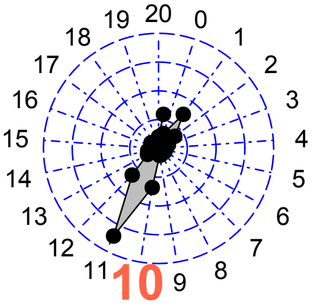


2nd siRNAs: Size distribution

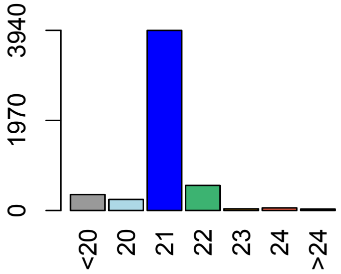

**Allenscore:** 5.5

**Target Interaction:**

```
5'- UUCCUGUACGGAGGAGAUGACU   AT3G01680.1
     |  |||:||||||||||||  |||
3'- AUGGAUAUGCCUCCUCUCCUGU   CI_ccm_535
```

**Target Site:** 1753

**Superfamily:** SupFam_5

**miRNA in ccm:** Y

**Confirmed Targeting**

| *2nd-siRNA* | *NanoPARE* |
|---|---|
| *ccm* | *ccm* |
| *cpe-2015* | *cpe-2015* |
| *cpe-2017* | |
| *cgr-dp* | *cgr-dp* |
| *cgr-pm* | |
| *cgr-mass* | |
| *cin* | *cin* |

### SEOR1

AT3G01680 - CI_ccm_535

Secondary siRNAs

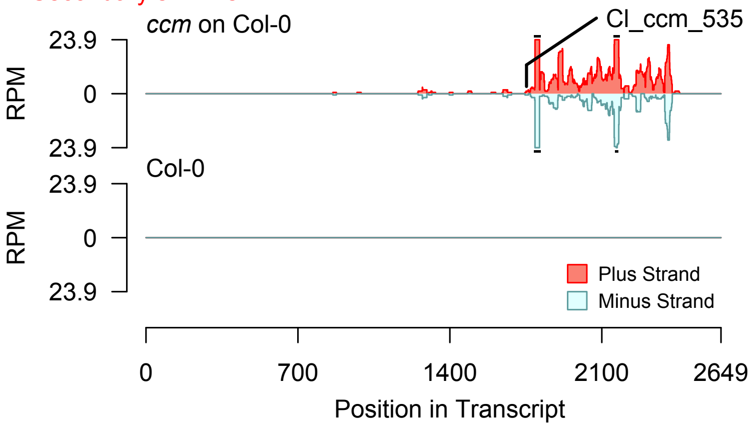


2nd siRNAs: Phase diagram

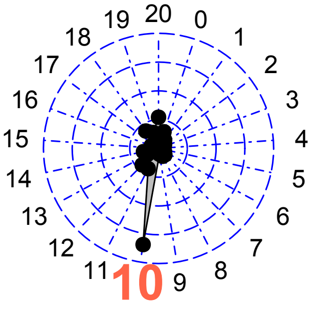


2nd siRNAs: Size distribution

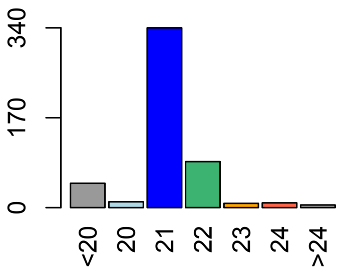


Degradome hits not found for sRNA

**Allenscore:** 5.5

**Target Interaction:**

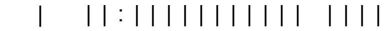


**Target Site:** 1753

**Superfamily:** SupFam_5

**miRNA in ccm:** Y

**Confirmed Targeting**

| *2nd-siRNA* | *NanoPARE* |
|---|---|
| *ccm* | *ccm* |
| *cpe-2015* | *cpe-2015* |
| *cpe-2017* | |
| *cgr-dp* | *cgr-dp* |
| *cgr-pm* | |
| *cgr-mass* | |
| *cin* | *cin* |

### SEOR1

AT3G01680 - CI_ccm_6321

Secondary siRNAs

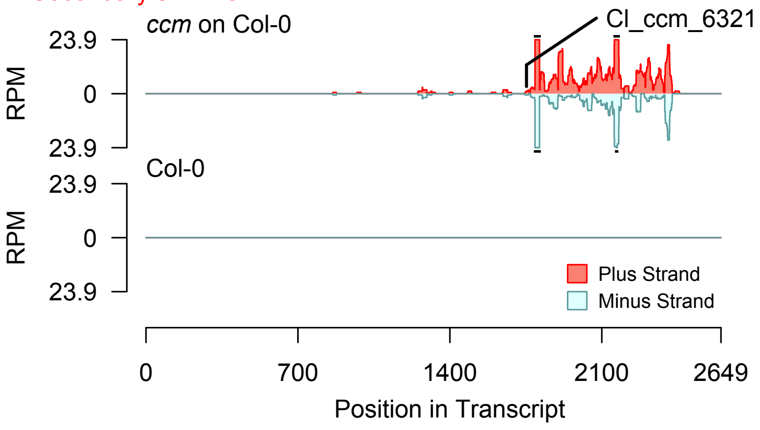


2nd siRNAs: Phase diagram

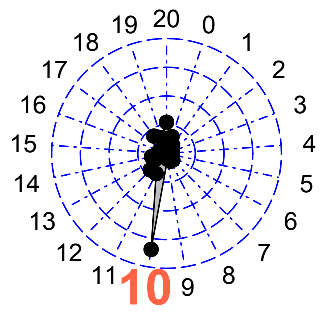


2nd siRNAs: Size distribution

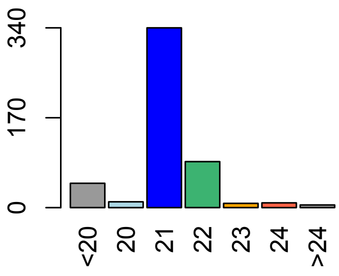


Degradome hits not found for sRNA

**Allenscore:** 6

**Target Interaction:**

```
5'- GGCCACUUUGAUUUCCACAUC   AT3G01680.1
     : | | | | | | | : : | | | : | |   | | | | |
3'- GUGGUGAAGUUAGAGUUGUAG   CI_ccm_315
```

**Target Site:** 2250

**Superfamily:** SupFam_19

**miRNA in ccm:** Y

**Confirmed Targeting**

| *2nd-siRNA* | *NanoPARE* |
|---|---|
| *ccm* | *ccm* |
| *cpe-2015* | *cpe-2015* |
| *cpe-2017* | |
| *cgr-dp* | *cgr-dp* |
| *cgr-pm* | |
| *cgr-mass* | |
| *cin* | *cin* |

### SEOR1

AT3G01680 - CI_ccm_315

Secondary siRNAs

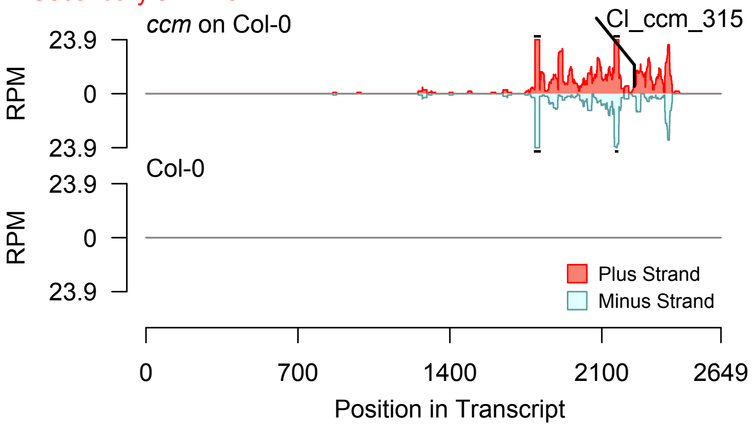


2nd siRNAs: Phase diagram

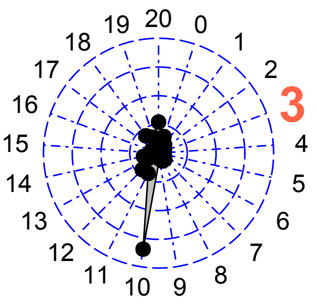


2nd siRNAs: Size distribution

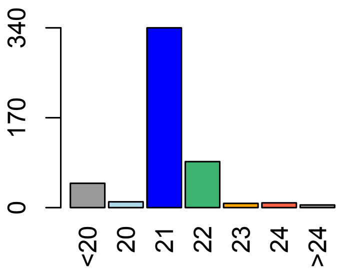


Degradome hits not found for sRNA

**Allenscore:** 5

**Target Interaction:**

```
5'- UCCUGUACGGAGGAGAUGACU   AT3G01680.1
      | | | | | | | | | | |   | | | | | |
3'- UCGACAUGCCUCCGCUACUGU   CI_cgr-mass_4275
```

**Target Site:** 1753

**Superfamily:** SupFam_5

**miRNA in ccm:** Y

**Confirmed Targeting**

| *2nd-siRNA* | *NanoPARE* |
|---|---|
| *ccm* | *ccm* |
| *cpe-2015* | *cpe-2015* |
| *cpe-2017* | |
| *cgr-dp* | *cgr-dp* |
| *cgr-pm* | |
| *cgr-mass* | |
| *cin* | *cin* |

### SEOR1

AT3G01680 - CI_cgr-mass_4275

Secondary siRNAs

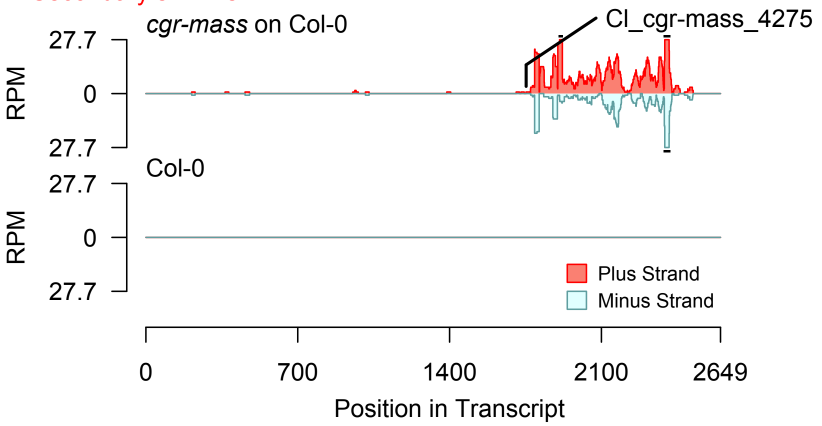


2nd siRNAs: Phase diagram

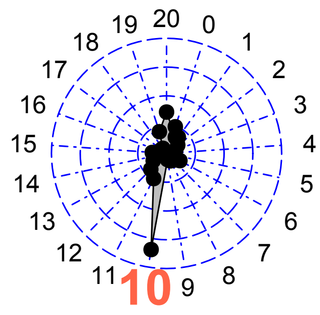


2nd siRNAs: Size distribution

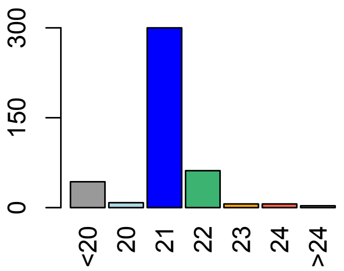


Degradome hits not found for sRNA

**Allenscore:** 5.5

**Target Interaction:**

```
5'- UUCCUGUACGGAGGAGAUGACU    AT3G01680.1
     |   | | | : | |   | | | | | | | |   | | | |
3'- AUGGAUAUACCUCCUCUUCUGA    CI_cgr-mass_5049
```

**Target Site:** 1753

**Superfamily:** SupFam_5

**miRNA in ccm:** Y

**Confirmed Targeting**

| 2nd-siRNA | NanoPARE |
|---|---|
| *ccm* | *ccm* |
| *cpe-2015* | *cpe-2015* |
| *cpe-2017* | |
| *cgr-dp* | *cgr-dp* |
| *cgr-pm* | |
| *cgr-mass* | |
| *cin* | *cin* |

### SEOR1

AT3G01680 - CI_cgr-mass_5049

Secondary siRNAs

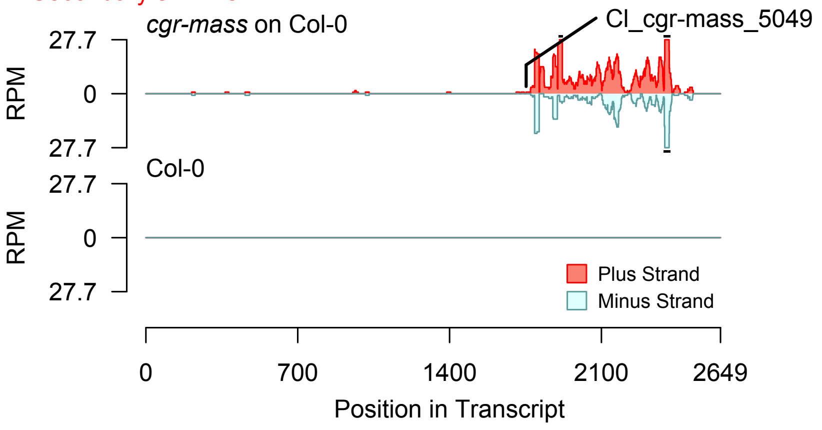


Degradome hits not found for sRNA

2nd siRNAs: Phase diagram

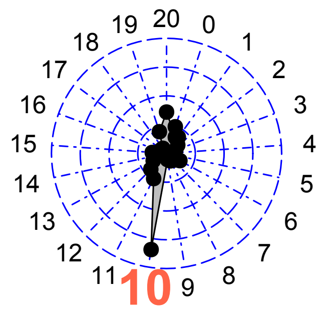


2nd siRNAs: Size distribution

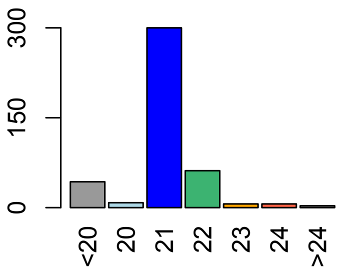

**Allenscore:** 3.5

**Target Interaction:**

```
5'- UUCCUGUACGGAGGAGAUGACU   AT3G01680.1
     | : | | | | | | | | | | | | | | |   | | |
3'- AGGGACAUGCCUCCUCUUCUGU   Cl_cpe-2015_74
```

**Target Site:** 1753

**Superfamily:** SupFam_5

**miRNA in ccm:** Y

**Confirmed Targeting**

| 2nd-siRNA | NanoPARE |
|---|---|
| *ccm* | *ccm* |
| *cpe-2015* | *cpe-2015* |
| *cpe-2017* | |
| *cgr-dp* | *cgr-dp* |
| *cgr-pm* | |
| *cgr-mass* | |
| *cin* | *cin* |

### SEOR1

AT3G01680 - Cl_cpe-2015_74

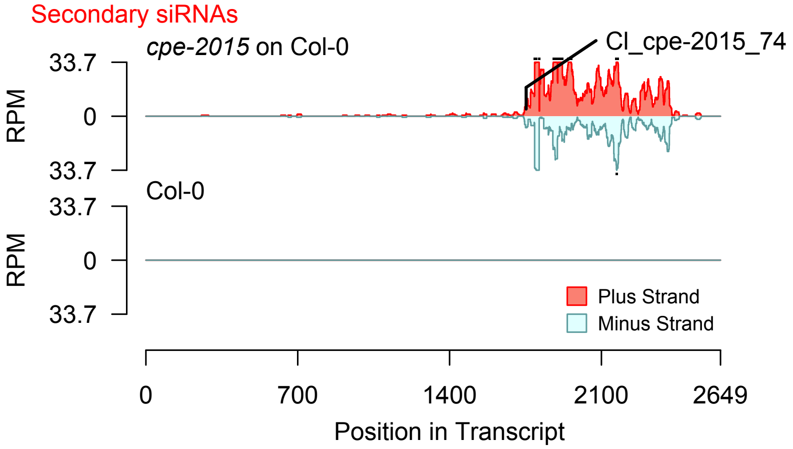


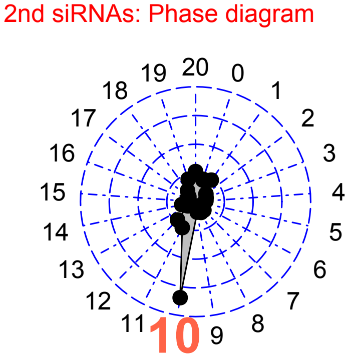


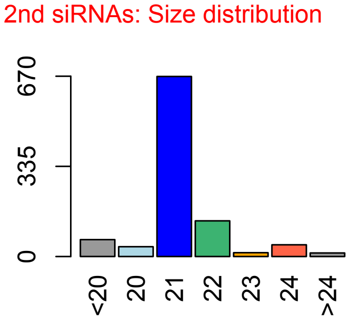


Degradome hits not found for sRNA

**Allenscore:** 5.5

**Target Interaction:**

```
5'- UUCCUGUACGGAGGAGAUGACU    AT3G01680.1
     |   | | | : | | | | | | | | | | | |    | | |
3'- AUGGAUAUGCCUCCUCUCCUGU    CI_cpe-2015_154
```

**Target Site:** 1753

**Superfamily:** SupFam_5

**miRNA in ccm:** Y

**Confirmed Targeting**

| *2nd-siRNA* | *NanoPARE* |
|---|---|
| *ccm* | *ccm* |
| *cpe-2015* | *cpe-2015* |
| *cpe-2017* | |
| *cgr-dp* | *cgr-dp* |
| *cgr-pm* | |
| *cgr-mass* | |
| *cin* | *cin* |

### SEOR1

AT3G01680 - CI_cpe-2015_154

Secondary siRNAs

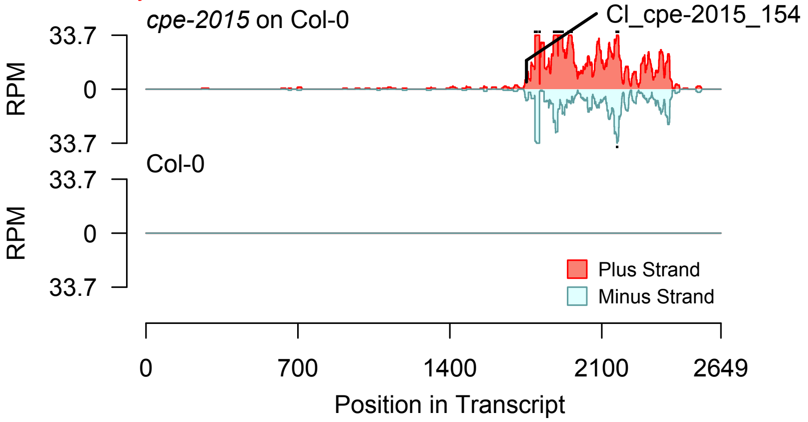


2nd siRNAs: Phase diagram

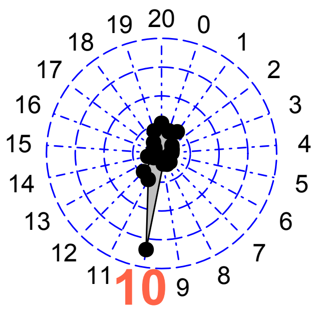


2nd siRNAs: Size distribution

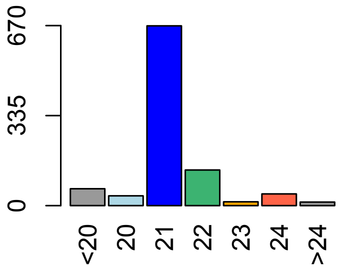


Degradome hits not found for sRNA

**Allenscore:** 5.5

**Target Interaction:**

```
5'- UUCCUGUACGGAGGAGAUGACU    AT3G01680.1
     |     | | : | | | | | | | | | | |   | | | |
3'- ACAGAUAUGCCUCCUCUCCUGA    CI_cpe-2015_1417
```

**Target Site:** 1753

**Superfamily:** SupFam_5

**miRNA in ccm:** Y

**Confirmed Targeting**

| *2nd-siRNA* | *NanoPARE* |
|---|---|
| *ccm* | *ccm* |
| *cpe-2015* | *cpe-2015* |
| *cpe-2017* | |
| *cgr-dp* | *cgr-dp* |
| *cgr-pm* | |
| *cgr-mass* | |
| *cin* | *cin* |

### SEOR1

AT3G01680 - CI_cpe-2015_1417

Secondary siRNAs

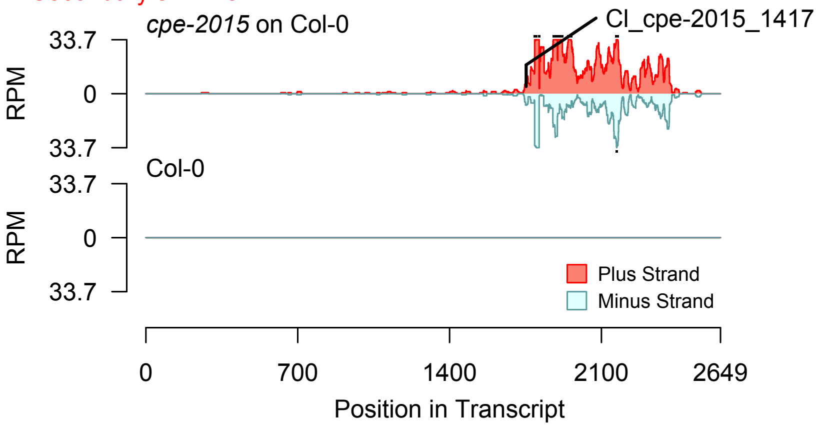


2nd siRNAs: Phase diagram

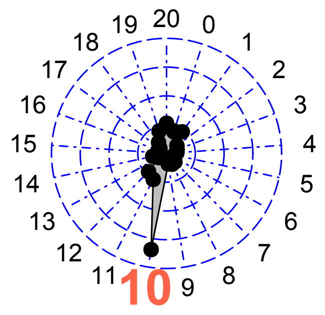


2nd siRNAs: Size distribution

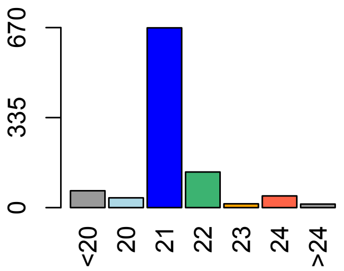


Degradome hits not found for sRNA

**Allenscore:** 6

**Target Interaction:**

```
5'- GGCCACUUUGAUUUCCACAUC   AT3G01680.1
     :|||||||::||:||  |||||
3'- GUGGUGAAGUUAGAGUUGUAG   CI_cpe-2015_191
```

**Target Site:** 2250

**Superfamily:** SupFam_19

**miRNA in ccm:** Y

**Confirmed Targeting**

| *2nd-siRNA* | *NanoPARE* |
|---|---|
| *ccm* | *ccm* |
| *cpe-2015* | *cpe-2015* |
| *cpe-2017* | |
| *cgr-dp* | *cgr-dp* |
| *cgr-pm* | |
| *cgr-mass* | |
| *cin* | *cin* |

### SEOR1

AT3G01680 - CI_cpe-2015_191

Secondary siRNAs

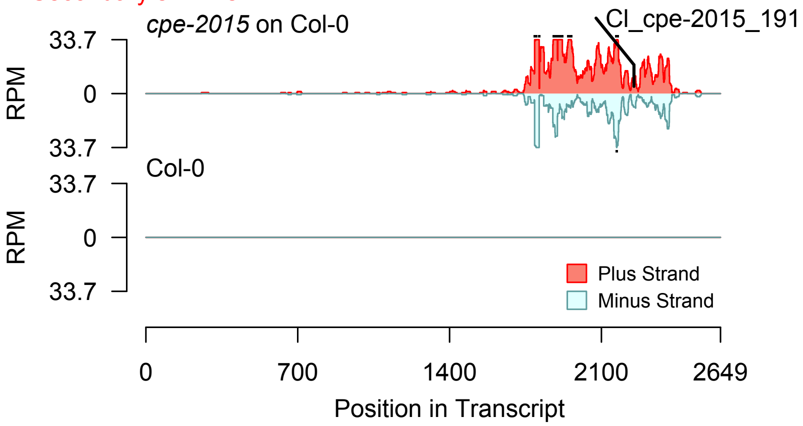


Degradome hits not found for sRNA

2nd siRNAs: Phase diagram

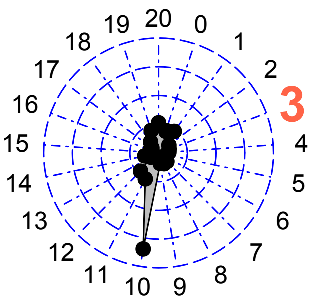


2nd siRNAs: Size distribution

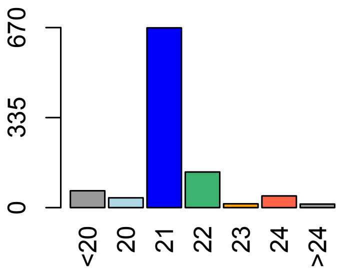

**Allenscore:** [illegible]

**Target Interaction:**

```
5'- AUUGCUUCGAGUGUCAGCGUC   AT3G01680.1
     | : | : | | | | | |   |   | | | : | : | |
3'- UGAUGAAGCU - AGAGUUGUAG   Cl_cpe-2015_3720
```

**Target Site:** [illegible]

**Superfamily:** SupFam_19

**miRNA in ccm:** Y

**Confirmed Targeting**

| *2nd-siRNA* | *NanoPARE* |
|---|---|
| *ccm* | *ccm* |
| *cpe-2015* | *cpe-2015* |
| *cpe-2017* | |
| *cgr-dp* | *cgr-dp* |
| *cgr-pm* | |
| *cgr-mass* | |
| *cin* | *cin* |

### SEOR1

AT3G01680 - Cl_cpe-2015_3720

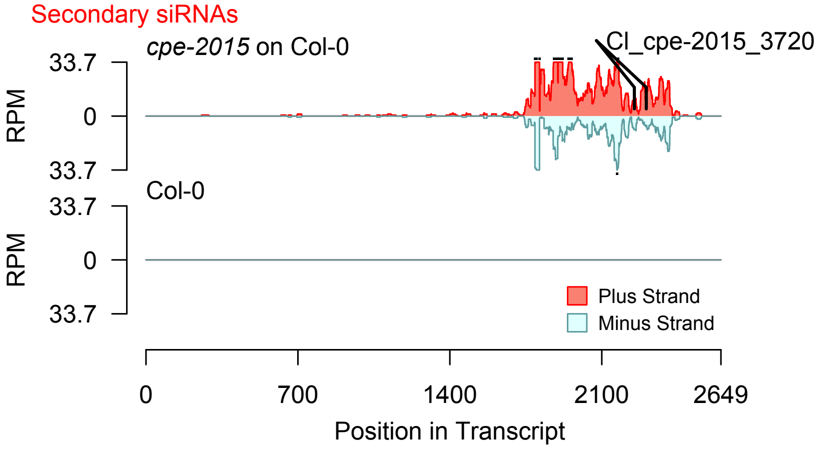


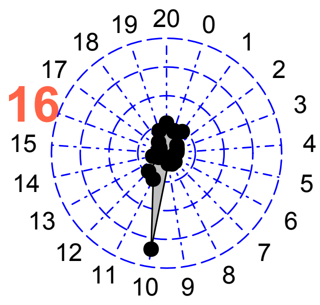


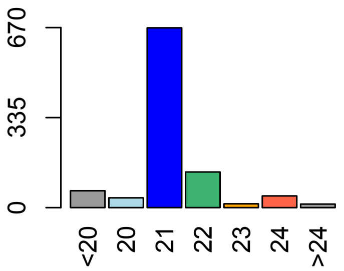


Degradome hits not found for sRNA

**Allenscore:** 3.5

**Target Interaction:**

```
5'- UUCCUGUACGGAGGAGAUGACU   AT3G01680.1
     | : | | | | | | | | | | | | | | |   | | |
3'- AGGGACAUGCCUCCUCUUCUGU   Cl_cpe-2017_89
```

**Target Site:** 1753

**Superfamily:** SupFam_5

**miRNA in ccm:** Y

**Confirmed Targeting**

| *2nd-siRNA* | *NanoPARE* |
|---|---|
| *ccm* | *ccm* |
| *cpe-2015* | *cpe-2015* |
| *cpe-2017* | |
| *cgr-dp* | *cgr-dp* |
| *cgr-pm* | |
| *cgr-mass* | |
| *cin* | *cin* |

### SEOR1

AT3G01680 - Cl_cpe-2017_89

Secondary siRNAs

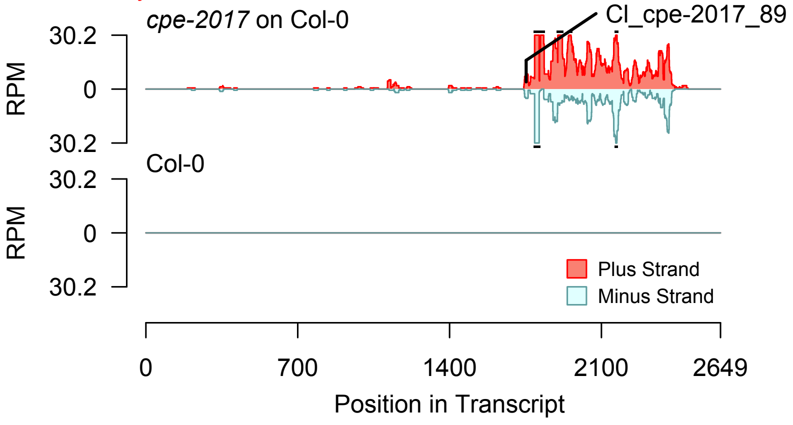


2nd siRNAs: Phase diagram

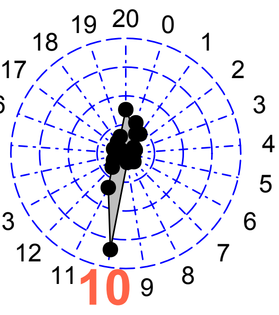


2nd siRNAs: Size distribution

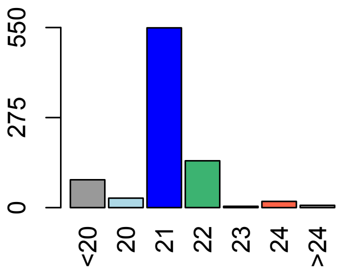


Degradome hits not found for sRNA

**Allenscore:** 5.5

**Target Interaction:**

```
5'- UUCCUGUACGGAGGAGAUGACU   AT3G01680.1
    | |||:||||||||||| |||
3'- AUGGAUAUGCCUCCUCUCCUGU   CI_cpe-2017_181
```

**Target Site:** 1753

**Superfamily:** SupFam_5

**miRNA in ccm:** Y

**Confirmed Targeting**

| *2nd-siRNA* | *NanoPARE* |
|---|---|
| *ccm* | *ccm* |
| *cpe-2015* | *cpe-2015* |
| *cpe-2017* | |
| *cgr-dp* | *cgr-dp* |
| *cgr-pm* | |
| *cgr-mass* | |
| *cin* | *cin* |

### SEOR1

AT3G01680 - CI_cpe-2017_181

Secondary siRNAs

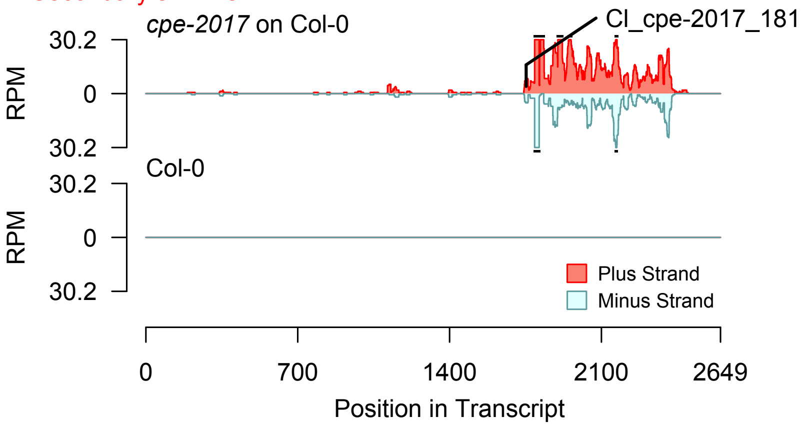


2nd siRNAs: Phase diagram

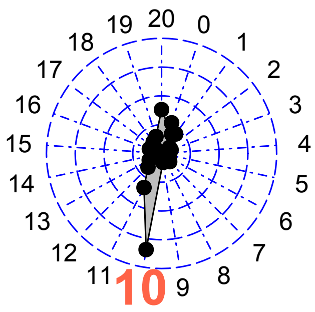


2nd siRNAs: Size distribution

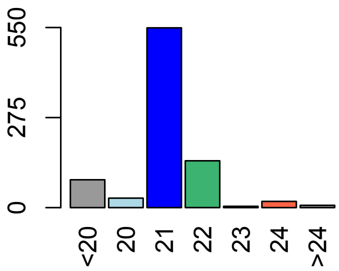


Degradome hits not found for sRNA

**Allenscore:** 5.5

**Target Interaction:**

```
5'- UUCCUGUACGGAGGAGAUGACU    AT3G01680.1
     |    | | : | | | | | | | | | | |   | | | |
3'- ACAGAUAUGCCUCCUCUCCUGA    CI_cpe-2017_1111
```

**Target Site:** 1753

**Superfamily:** SupFam_5

**miRNA in ccm:** Y

**Confirmed Targeting**

| *2nd-siRNA* | *NanoPARE* |
|---|---|
| *ccm* | *ccm* |
| *cpe-2015* | *cpe-2015* |
| *cpe-2017* | |
| *cgr-dp* | *cgr-dp* |
| *cgr-pm* | |
| *cgr-mass* | |
| *cin* | *cin* |

### SEOR1

AT3G01680 - CI_cpe-2017_1111

Secondary siRNAs

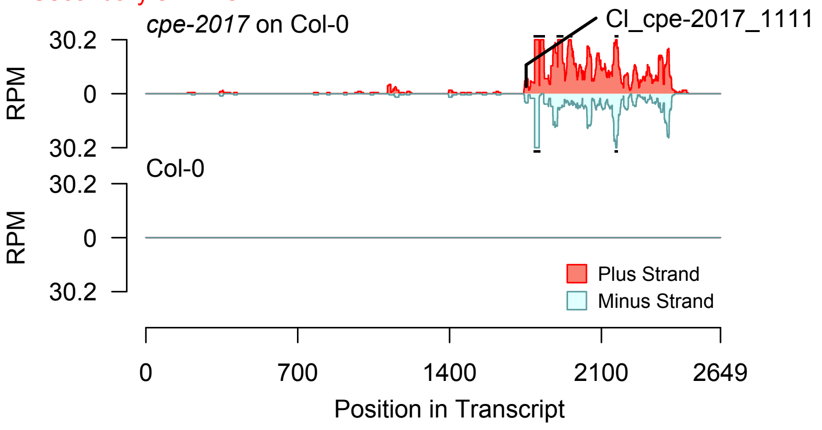


2nd siRNAs: Phase diagram

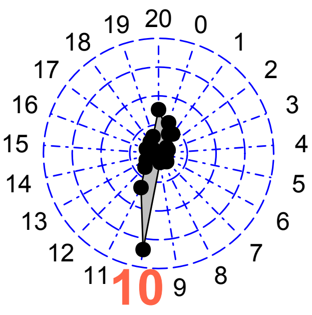


2nd siRNAs: Size distribution

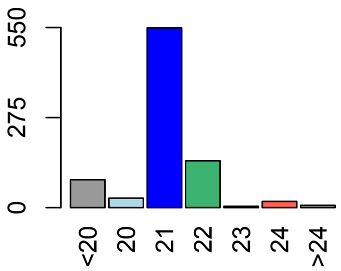


Degradome hits not found for sRNA

**Allenscore:** 6

**Target Interaction:**

```
5'- GGCCACUUUGAUUUCCACAUC   AT3G01680.1
     : | | | | | | | : : | | : | |   | | | | |
3'- GUGGUGAAGUUAGAGUUGUAG   Cl_cpe-2017_172
```

**Target Site:** 2250

**Superfamily:** SupFam_19

**miRNA in ccm:** Y

**Confirmed Targeting**

| *2nd-siRNA* | *NanoPARE* |
|---|---|
| *ccm* | *ccm* |
| *cpe-2015* | *cpe-2015* |
| *cpe-2017* | |
| *cgr-dp* | *cgr-dp* |
| *cgr-pm* | |
| *cgr-mass* | |
| *cin* | *cin* |

### SEOR1

AT3G01680 - Cl_cpe-2017_172

Secondary siRNAs

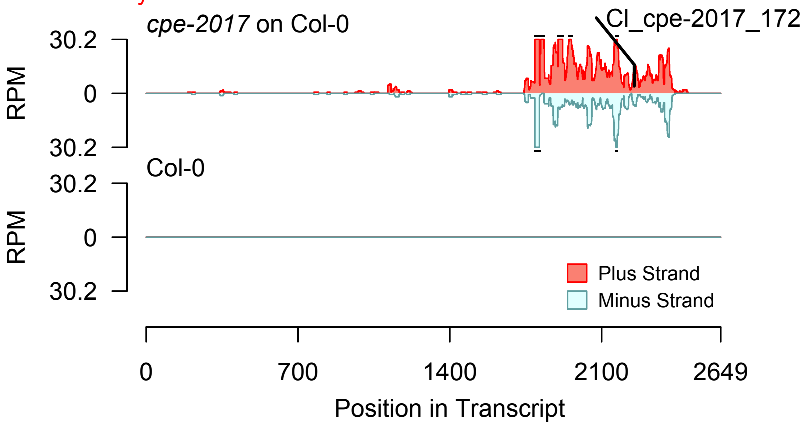


2nd siRNAs: Phase diagram

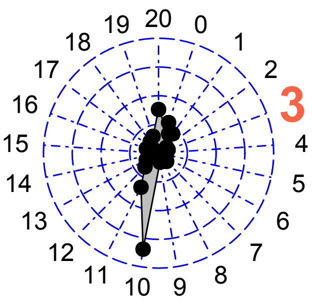


2nd siRNAs: Size distribution

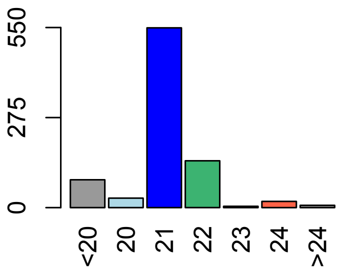


Degradome hits not found for sRNA

**Allenscore:** [illegible]

**Target Interaction:**

```
5'- AUUGCUUCGAGUGUCAGCGUC    AT3G01680.1
    | : | : | | | | | |  |   | | | : | : | |
3'- UGAUGAAGCU - AGAGUUGUAG  CI_cpe-2017_2830
```

**Target Site:** [illegible]

**Superfamily:** SupFam_19

**miRNA in ccm:** Y

**Confirmed Targeting**

| *2nd-siRNA* | *NanoPARE* |
|---|---|
| *ccm* | *ccm* |
| *cpe-2015* | *cpe-2015* |
| *cpe-2017* | |
| *cgr-dp* | *cgr-dp* |
| *cgr-pm* | |
| *cgr-mass* | |
| *cin* | *cin* |

### SEOR1

AT3G01680 - CI_cpe-2017_2830

Secondary siRNAs

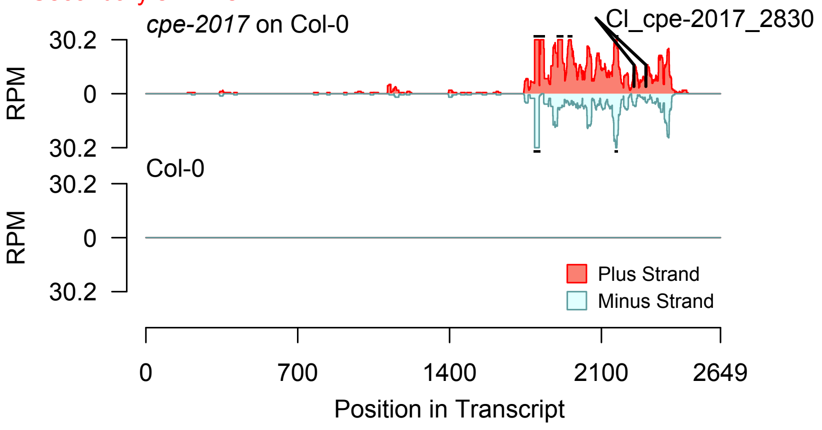


Degradome hits not found for sRNA

2nd siRNAs: Phase diagram

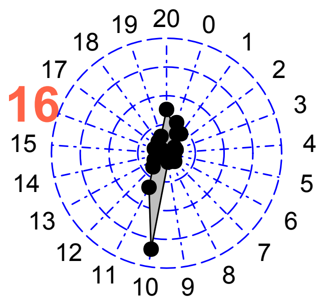


2nd siRNAs: Size distribution

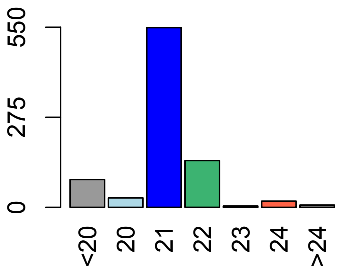

**Allenscore:** 5.5

**Target Interaction:**

```
5'- UUGUAUGGUGAGCCUGGAACU   AT4G29040.1
      ||||||| :  |||  ||||||||
3'- UACAUACUUCUCAGACCUUGU   CI_cin_1915
```

**Target Site:** 776

**Superfamily:** SupFam_1368

**miRNA in ccm:** N/A

**Confirmed Targeting**

| *2nd-siRNA* | *NanoPARE* |
|---|---|
| *ccm* | *ccm* |
| *cpe-2015* | *cpe-2015* |
| *cpe-2017* | |
| *cgr-dp* | *cgr-dp* |
| *cgr-pm* | |
| *cgr-mass* | |
| *cin* | *cin* |

### RPT2a

AT4G29040 - CI_cin_1915

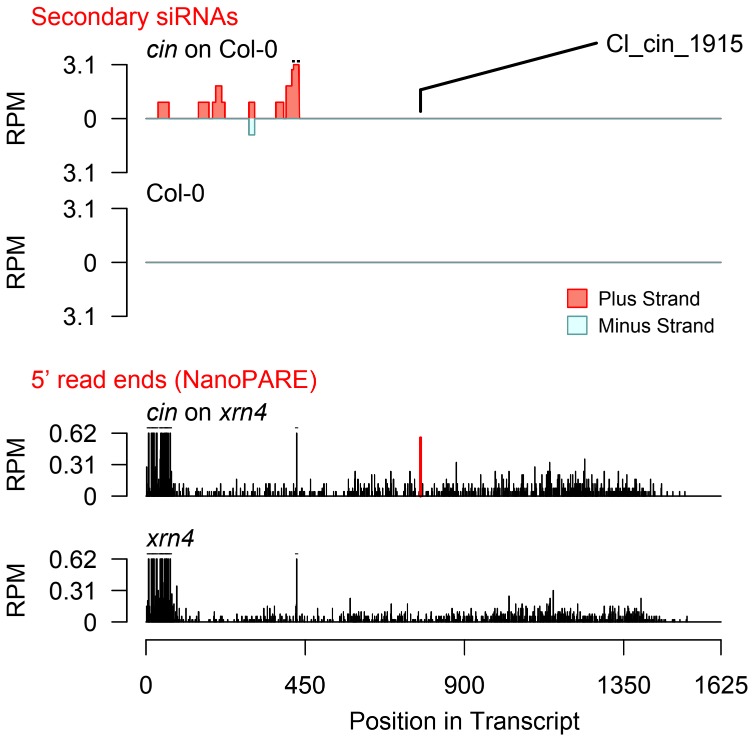


Diff. Exp. secondary siRNA locus not found

**Allenscore:** 5

**Target Interaction:**

```
5'- GUCCCCUCCAAAGAUUCCUGA   AT4G23630.1
     |||  ||  |||||:|||||  ||
3'- CAGAGGUGGUUUUUAAGGUCU   CI_cgr-dp_91
```

**Target Site:** 1080

**Superfamily:** SupFam_207

**miRNA in ccm:** N/A

**Confirmed Targeting**

| *2nd-siRNA* | *NanoPARE* |
|---|---|
| *ccm* | *ccm* |
| *cpe-2015* | *cpe-2015* |
| *cpe-2017* | |
| *cgr-dp* | *cgr-dp* |
| *cgr-pm* | |
| *cgr-mass* | |
| *cin* | *cin* |

### BTI1

AT4G23630 - CI_cgr-dp_91

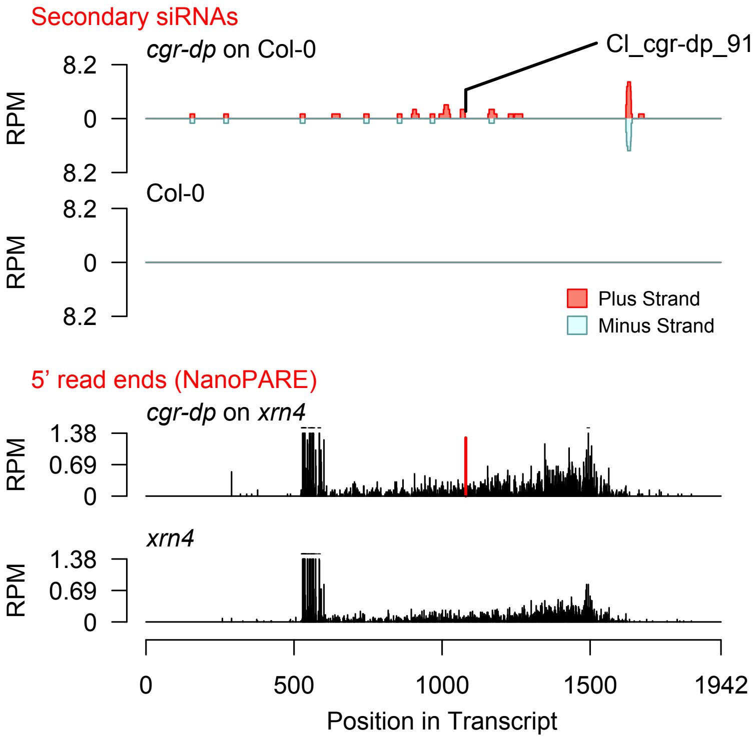


Diff. Exp. secondary siRNA locus not found

**Allenscore:** 4.5

**Target Interaction:**

```
5'- UGCAGUGUCGAGGGUUGGAAGU   AT4G27630.1
     | | | |   | |   | : | | | : | | | | | | |
3'- ACGU - AC - GUUCCUAACCUUCU   CI_cgr-dp_190
```

**Target Site:** 1649

**Superfamily:** SupFam_895

**miRNA in ccm:** N/A

**Confirmed Targeting**

| *2nd-siRNA* | *NanoPARE* |
|---|---|
| *ccm* | *ccm* |
| *cpe-2015* | *cpe-2015* |
| *cpe-2017* | |
| *cgr-dp* | *cgr-dp* |
| *cgr-pm* | |
| *cgr-mass* | |
| *cin* | *cin* |

### GTG2

AT4G27630 - CI_cgr-dp_190

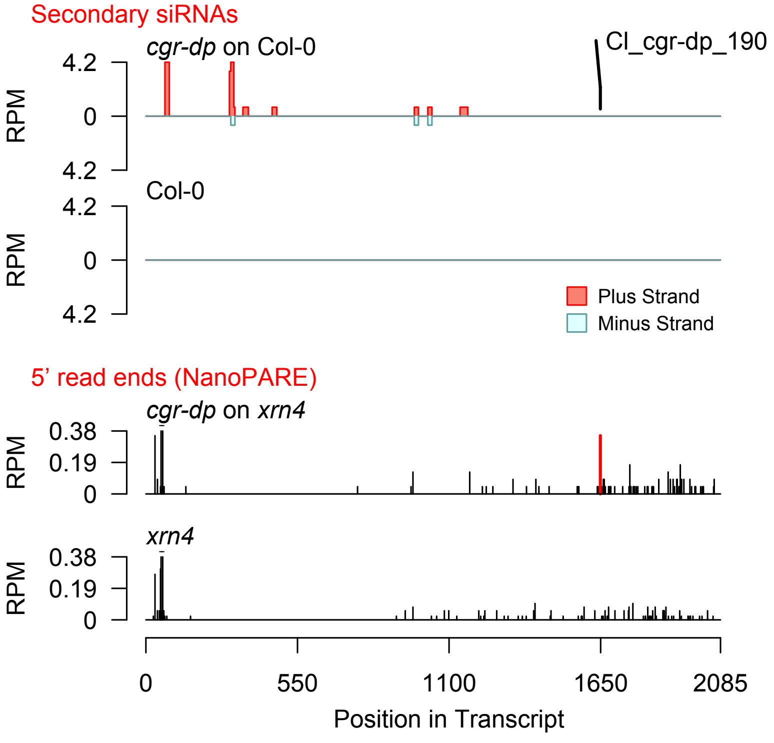


Diff. Exp. secondary siRNA locus not found

**Allenscore:** 5

**Target Interaction:**

```
5'- UUGAGAAAGAAUGGGAAGAAC   AT1G67230.1
     :  ||||||| ||||||||||
3'- GUCUCUUUCAUACCCUUCUUC   CI_cgr-dp_10674
```

**Target Site:** 1930

**Superfamily:** SupFam_3

**miRNA in ccm:** Y

**Confirmed Targeting**

| *2nd-siRNA* | *NanoPARE* |
|---|---|
| *ccm* | *ccm* |
| *cpe-2015* | *cpe-2015* |
| *cpe-2017* | |
| *cgr-dp* | *cgr-dp* |
| *cgr-pm* | |
| *cgr-mass* | |
| *cin* | *cin* |

### LINC1

AT1G67230 - CI_cgr-dp_10674

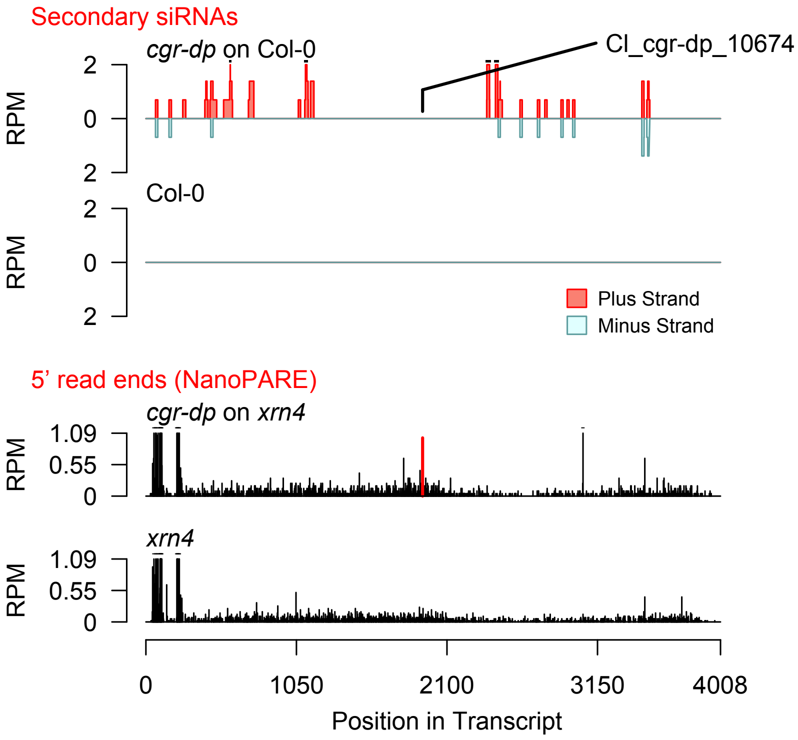


Diff. Exp. secondary siRNA locus not found

**Allenscore:** 6

**Target Interaction:**

```
5'- AACCGAAGAAGAUG - AAAGCUACA    AT3G51670.1
    |  | | | | | | | | | | |   | | | | | |   | |
3'- UGGGCUUCUUCUACUUUUCGA - GU    CI_cgr-dp_279
```

**Target Site:** 1326

**Superfamily:** SupFam_37

**miRNA in ccm:** N/A

**Confirmed Targeting**

| *2nd-siRNA* | *NanoPARE* |
|---|---|
| *ccm* | *ccm* |
| *cpe-2015* | *cpe-2015* |
| *cpe-2017* | |
| *cgr-dp* | *cgr-dp* |
| *cgr-pm* | |
| *cgr-mass* | |
| *cin* | *cin* |

### SEC14_cytosolic_factor_family_protein___

AT3G51670 - CI_cgr-dp_279

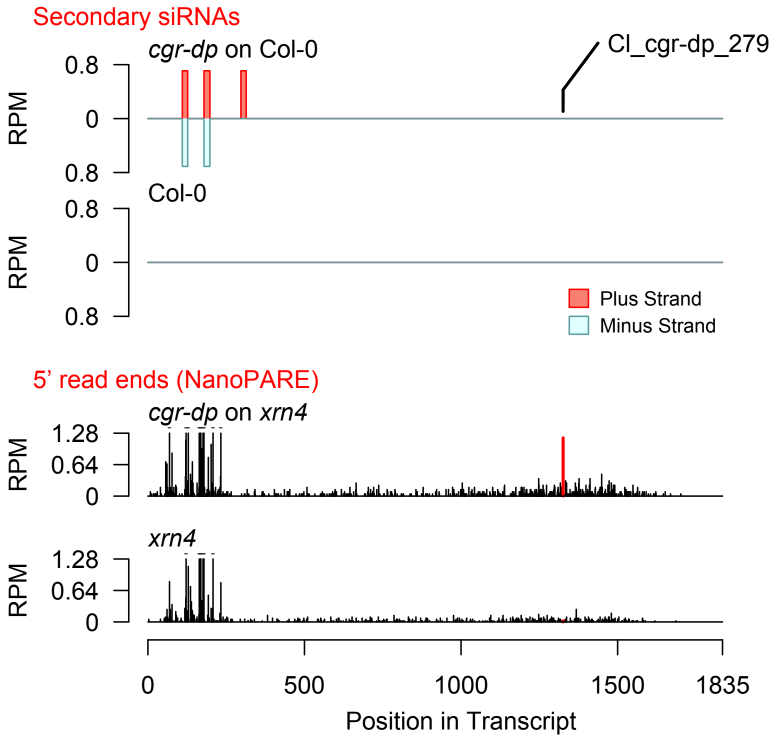


Diff. Exp. secondary siRNA locus not found

**Allenscore:** 5

**Target Interaction:**

```
5'- CUCACGUGACCUGCUUCUCCG   AT5G39610.1
       |||||  |||||||||||:
3'- UUUUGCACGGGACGAAGAGGU   CI_cgr-dp_26729
```

**Target Site:** 904

**Superfamily:** SupFam_187

**miRNA in ccm:** N/A

**Confirmed Targeting**

| *2nd-siRNA* | *NanoPARE* |
|---|---|
| *ccm* | *ccm* |
| *cpe-2015* | *cpe-2015* |
| *cpe-2017* | |
| *cgr-dp* | *cgr-dp* |
| *cgr-pm* | |
| *cgr-mass* | |
| *cin* | *cin* |

### NAC6

AT5G39610 - CI_cgr-dp_26729

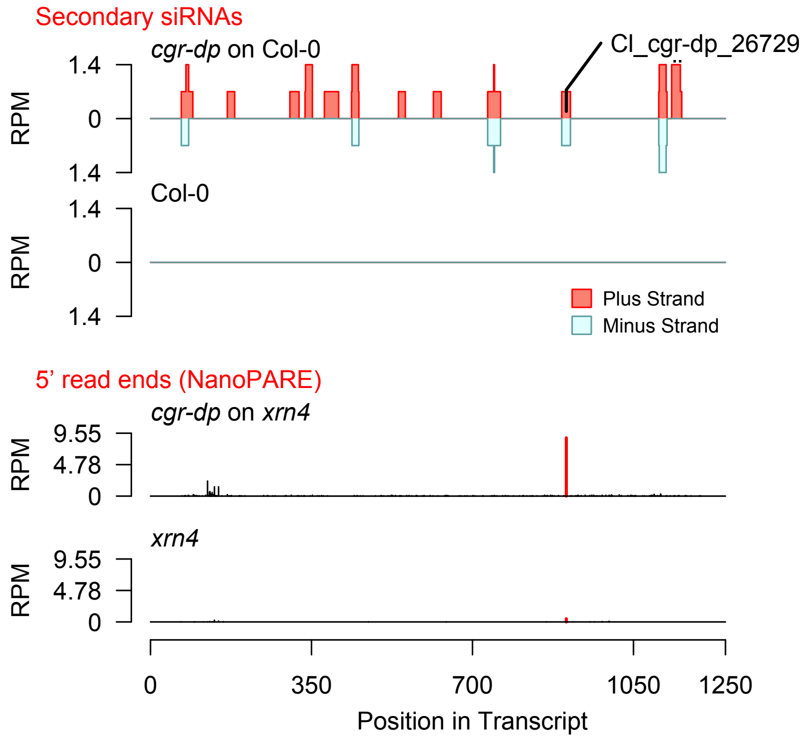


Diff. Exp. secondary siRNA locus not found

**Allenscore:** 4.5

**Target Interaction:**

```
5'- CGGGUUACGUUCCUCCAGAGUA    AT4G39400.1
      : | | : | | | : |   | | | | | | | | | | |
3'- AUCCGAUGUACGGAGGUCUCAU    CI_cgr-mass_20246
```

**Target Site:** 3530

**Superfamily:** SupFam_59

**miRNA in ccm:** N/A

**Confirmed Targeting**

| *2nd-siRNA* | *NanoPARE* |
|---|---|
| *ccm* | *ccm* |
| *cpe-2015* | *cpe-2015* |
| *cpe-2017* | |
| *cgr-dp* | *cgr-dp* |
| *cgr-pm* | |
| *cgr-mass* | |
| *cin* | *cin* |

### BRI1

AT4G39400 - CI_cgr-mass_20246

Secondary siRNAs

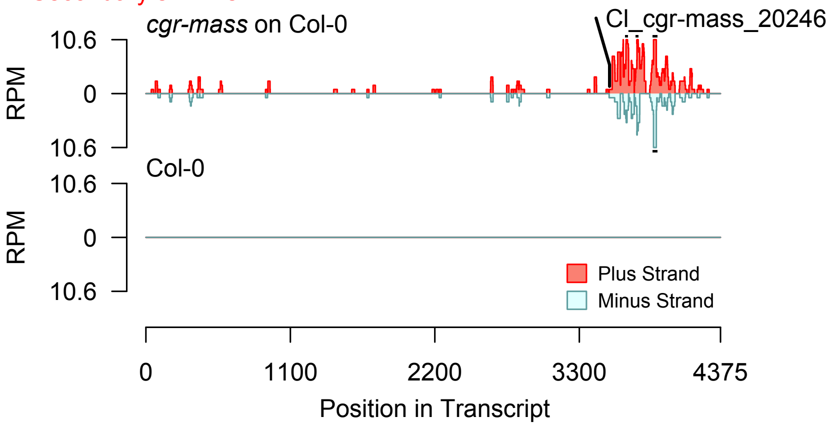


2nd siRNAs: Phase diagram

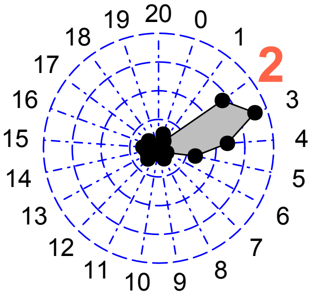


2nd siRNAs: Size distribution

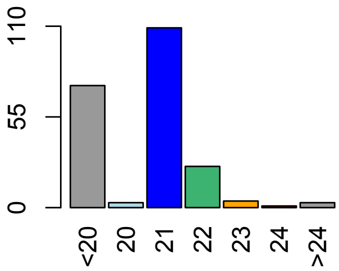


Degradome hits not found for sRNA

**Allenscore:** 3.5

**Target Interaction:**

```
5'- AAAGGGAAGUCAUCCUUGGCU   AT1G17590.1
        | | | : | | | | | | | | | | | | | | |
3'- AGUCCUUUCAGUAGGAACCGU   CI_cin_1673
```

**Target Site:** 1329

**Superfamily:** SupFam_1364

**miRNA in ccm:** N/A

**Confirmed Targeting**

| *2nd-siRNA* | *NanoPARE* |
|---|---|
| *ccm* | *ccm* |
| *cpe-2015* | *cpe-2015* |
| *cpe-2017* | |
| *cgr-dp* | *cgr-dp* |
| *cgr-pm* | |
| *cgr-mass* | |
| *cin* | *cin* |

# NF-YA8

AT1G17590 - CI_cin_1673

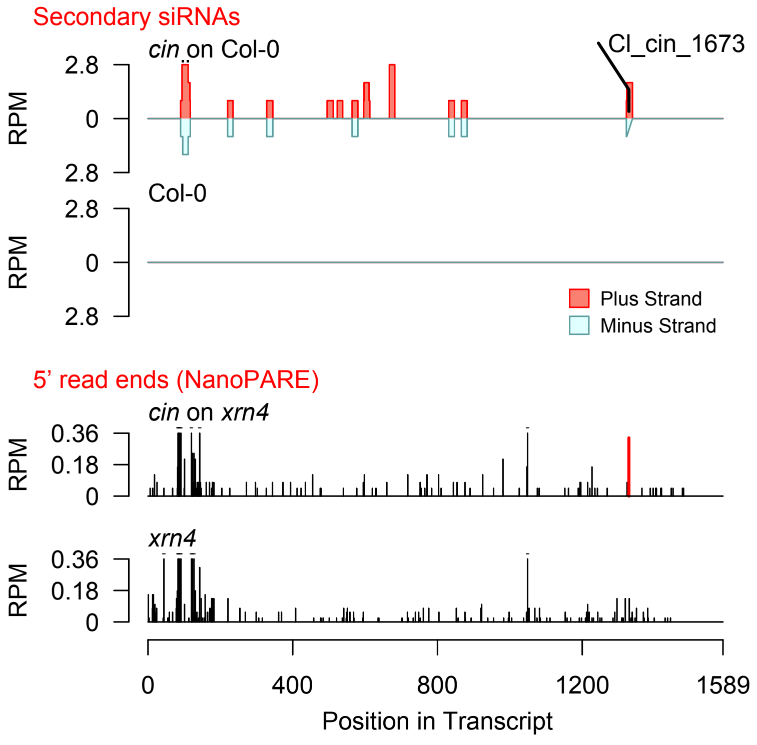


Diff. Exp. secondary siRNA locus not found

**Allenscore:** 3

**Target Interaction:**

```
5'- CAGGGAUACGCUGCUCCAGAAUA   AT5G02290.1
     | | | | |   | | | | | | | |   | | | | | | | |
3'- GUCCCAAUGCGACGUGGUCUUAU   CI_ccm_818
```

**Target Site:** 1120

**Superfamily:** SupFam_124

**miRNA in ccm:** N

**Confirmed Targeting**

| *2nd-siRNA* | *NanoPARE* |
|---|---|
| *ccm* | *ccm* |
| *cpe-2015* | *cpe-2015* |
| *cpe-2017* | |
| *cgr-dp* | *cgr-dp* |
| *cgr-pm* | |
| *cgr-mass* | |
| *cin* | *cin* |

### NAK

AT5G02290 - CI_ccm_818

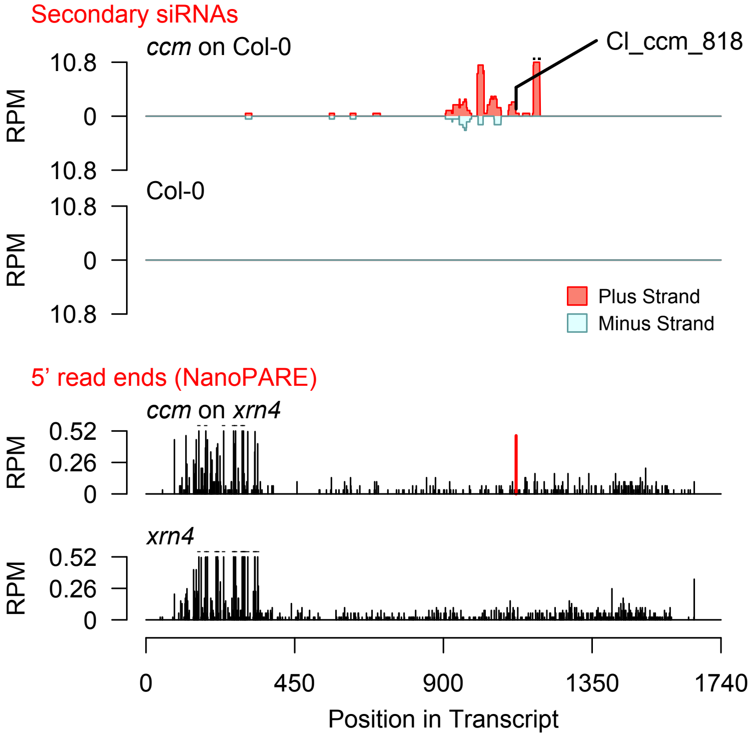


Diff. Exp. secondary siRNA locus not found

**Allenscore:** 4.5

**Target Interaction:**

```
5'- CCGUUUCUACUCCACCAUCACA   AT4G28300.1
        | | | | | : |   | | | | | | | : | | | |
3'- UUCAAAGGUAAGGUGGUGGUGU   CI_cgr-dp_307
```

**Target Site:** 990

**Superfamily:** SupFam_897

**miRNA in ccm:** N/A

**Confirmed Targeting**

| *2nd-siRNA* | *NanoPARE* |
|---|---|
| *ccm* | *ccm* |
| *cpe-2015* | *cpe-2015* |
| *cpe-2017* | |
| *cgr-dp* | *cgr-dp* |
| *cgr-pm* | |
| *cgr-mass* | |
| *cin* | *cin* |

### DUF1421_containing_protein

AT4G28300 - CI_cgr-dp_307

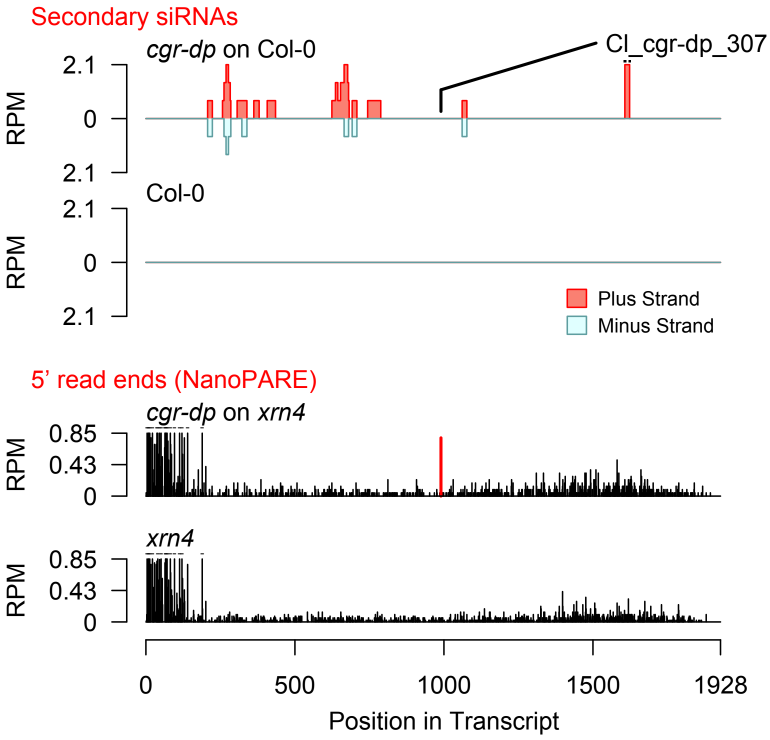


Diff. Exp. secondary siRNA locus not found

**Allenscore:** 3.5

**Target Interaction:**

```
5'- GAGAUGUUGAGAUUGAAGUUG   AT5G07350.1
     : | | | | | | | | | | | | | | | | | |     |
3'- UUCUACAACUCUAACUUCAUC   Cl_ccm_935
```

**Target Site:** 2216

**Superfamily:** SupFam_287

**miRNA in ccm:** Y

**Confirmed Targeting**

| *2nd-siRNA* | *NanoPARE* |
|---|---|
| *ccm* | *ccm* |
| *cpe-2015* | *cpe-2015* |
| *cpe-2017* | |
| *cgr-dp* | *cgr-dp* |
| *cgr-pm* | |
| *cgr-mass* | |
| *cin* | *cin* |

### Tudor1

AT5G07350 - Cl_ccm_935

Secondary siRNAs

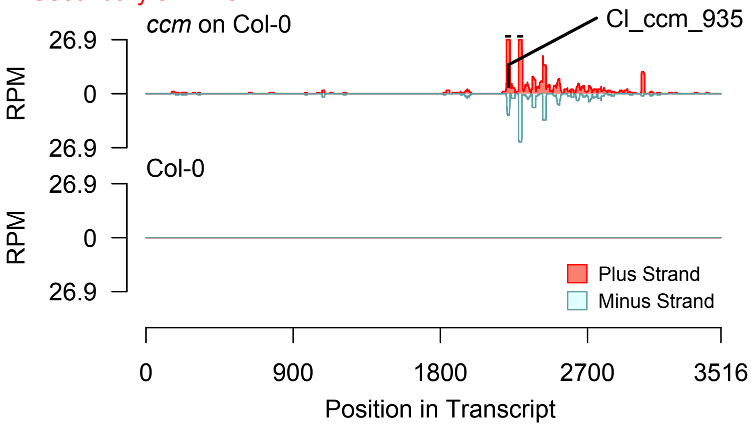


2nd siRNAs: Phase diagram

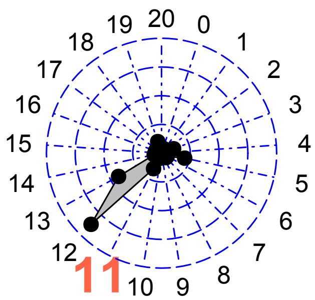


2nd siRNAs: Size distribution

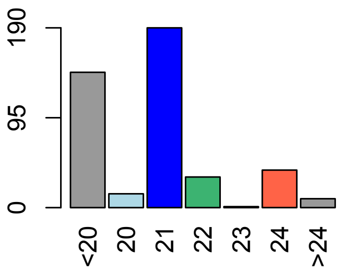


Degradome hits not found for sRNA

**Allenscore:** 3.5

**Target Interaction:**

```
5'- GAGAUGUUGAGAUUGAAGUUG   AT5G07350.1
     :||||||||||||||||||| |
3'- UUCUACAACUCUAACUUCAUC   Cl_cpe-2015_358
```

**Target Site:** 2216

**Superfamily:** SupFam_287

**miRNA in ccm:** Y

**Confirmed Targeting**

| *2nd-siRNA* | *NanoPARE* |
|---|---|
| *ccm* | *ccm* |
| *cpe-2015* | *cpe-2015* |
| *cpe-2017* | |
| *cgr-dp* | *cgr-dp* |
| *cgr-pm* | |
| *cgr-mass* | |
| *cin* | *cin* |

### Tudor1

AT5G07350 - Cl_cpe-2015_358

Secondary siRNAs

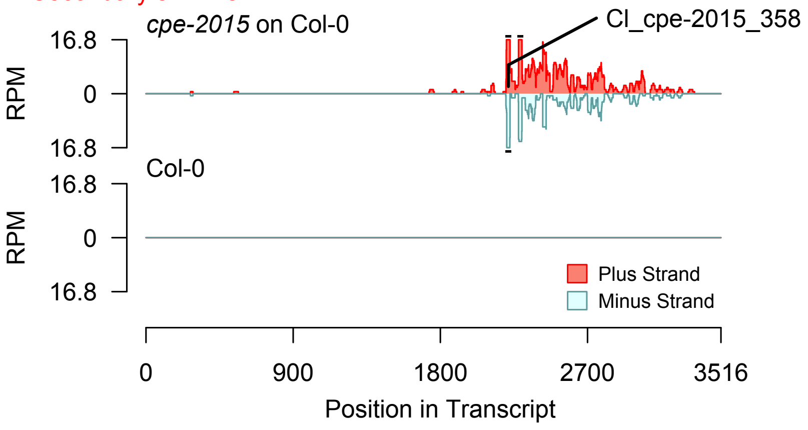


2nd siRNAs: Phase diagram

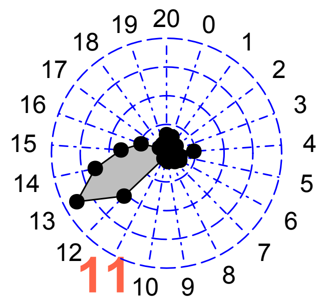


2nd siRNAs: Size distribution

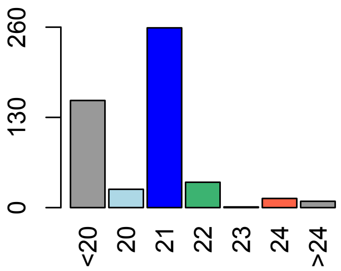


Degradome hits not found for sRNA

**Allenscore:** 3.5

**Target Interaction:**

```
5'- GAGAUGUUGAGAUUGAAGUUG   AT5G07350.1
    : | | | | | | | | | | | | | | | | | |   |
3'- UUCUACAACUCUAACUUCAUC   Cl_cpe-2017_269
```

**Target Site:** 2216

**Superfamily:** SupFam_287

**miRNA in ccm:** Y

**Confirmed Targeting**

| *2nd-siRNA* | *NanoPARE* |
|---|---|
| *ccm* | *ccm* |
| *cpe-2015* | *cpe-2015* |
| *cpe-2017* | |
| *cgr-dp* | *cgr-dp* |
| *cgr-pm* | |
| *cgr-mass* | |
| *cin* | *cin* |

### Tudor1

AT5G07350 - Cl_cpe-2017_269

Secondary siRNAs

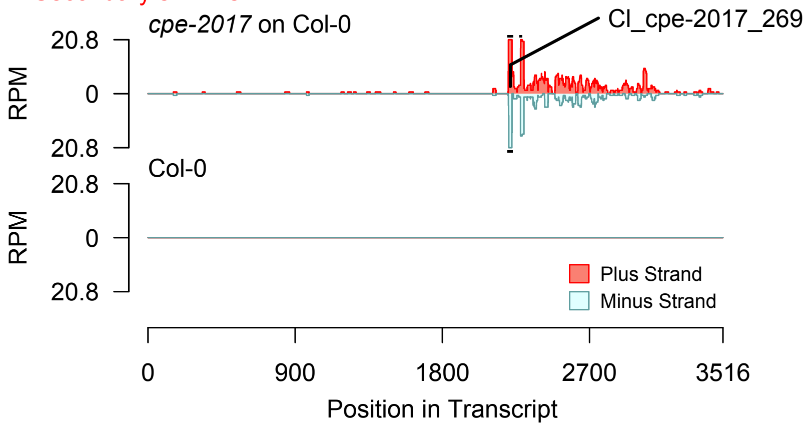


2nd siRNAs: Phase diagram

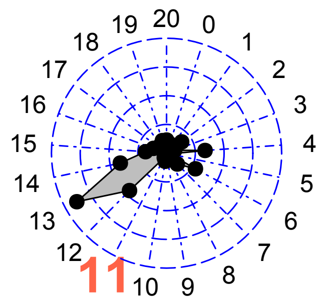


2nd siRNAs: Size distribution

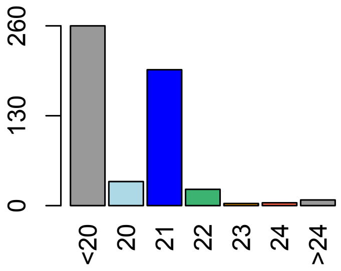


Degradome hits not found for sRNA

**Allenscore:** 1.5

**Target Interaction:**

```
5'- GCGGCAAUUCAUUCUUGGCUU   AT3G20910.1
     : | | | | | | | | | | | | : | | | | | | | |
3'- UGCCGUUAAGUAGGAACCGAA   Cl_ccm_2887
```

**Target Site:** 1073

**Superfamily:** SupFam_72

**miRNA in ccm:** Y

**Confirmed Targeting**

| *2nd-siRNA* | *NanoPARE* |
|---|---|
| *ccm* | *ccm* |
| *cpe-2015* | *cpe-2015* |
| *cpe-2017* | |
| *cgr-dp* | *cgr-dp* |
| *cgr-pm* | |
| *cgr-mass* | |
| *cin* | *cin* |

# NF-YA9

AT3G20910 - Cl_ccm_2887

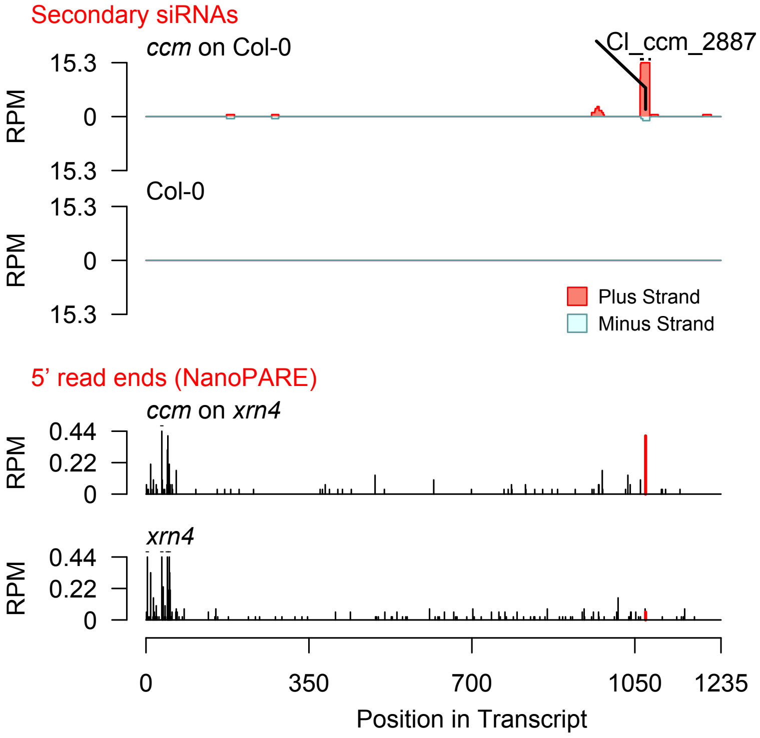


Diff. Exp. secondary siRNA
locus not found

**Allenscore:** 4

**Target Interaction:**

```
5'- GGCGGCAAUUCAUUCUUGGCUU   AT3G20910.1
    : : | | | | | |   | | | | : | | | | | | |
3'- UUGCCGUUUAGUAGGAACCGAU   CI_ccm_2720
```

**Target Site:** 1073

**Superfamily:** SupFam_72

**miRNA in ccm:** Y

**Confirmed Targeting**

| *2nd-siRNA* | *NanoPARE* |
|---|---|
| *ccm* | *ccm* |
| *cpe-2015* | *cpe-2015* |
| *cpe-2017* | |
| *cgr-dp* | *cgr-dp* |
| *cgr-pm* | |
| *cgr-mass* | |
| *cin* | *cin* |

# NF-YA9

AT3G20910 - CI_ccm_2720

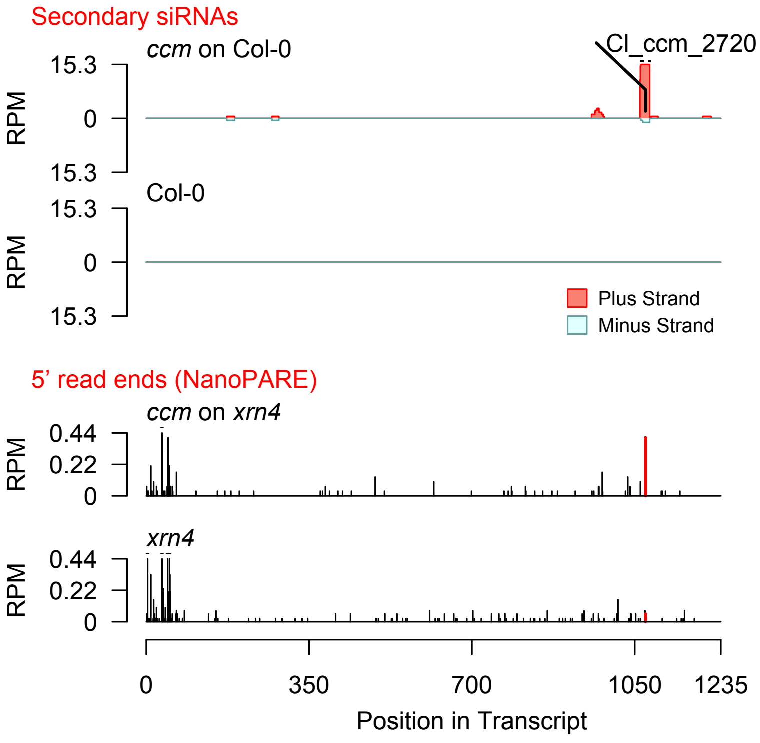


Diff. Exp. secondary siRNA locus not found

**Allenscore:** 6

**Target Interaction:**

```
5'- UCAAGAACGCUACGAGGAACUUUC    AT2G02800.1
       | | : | | | | | | : | | | | | : | | | | | | :
3'- UUUUUUUGCGGUGCUCUUUGAAGU    Cl_cgr-dp_62946
```

**Target Site:** 582

**Superfamily:** SupFam_26

**miRNA in ccm:** Y

**Confirmed Targeting**

| *2nd-siRNA* | *NanoPARE* |
|---|---|
| *ccm* | *ccm* |
| *cpe-2015* | *cpe-2015* |
| *cpe-2017* | |
| *cgr-dp* | *cgr-dp* |
| *cgr-pm* | |
| *cgr-mass* | |
| *cin* | *cin* |

### APK2B

AT2G02800 - Cl_cgr-dp_62946

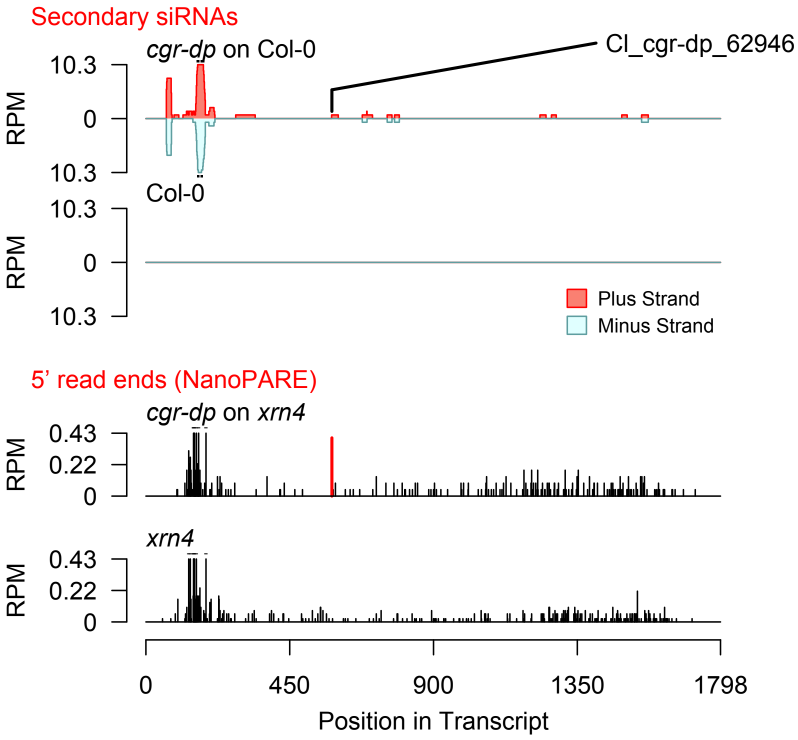


Diff. Exp. secondary siRNA locus not found

**Allenscore:** 4

**Target Interaction:**

```
5'- AGAACGCUACGAGGAACUUUC   AT2G02800.1
     | : | | | | | : | | | | | : | | | | | :
3'- UUUUGCGGUGCUCUUUGAAGU   Cl_cgr-dp_29
```

**Target Site:** 582

**Superfamily:** SupFam_26

**miRNA in ccm:** Y

**Confirmed Targeting**

| *2nd-siRNA* | *NanoPARE* |
|---|---|
| *ccm* | *ccm* |
| *cpe-2015* | *cpe-2015* |
| *cpe-2017* | |
| *cgr-dp* | *cgr-dp* |
| *cgr-pm* | |
| *cgr-mass* | |
| *cin* | *cin* |

### APK2B

AT2G02800 - Cl_cgr-dp_29

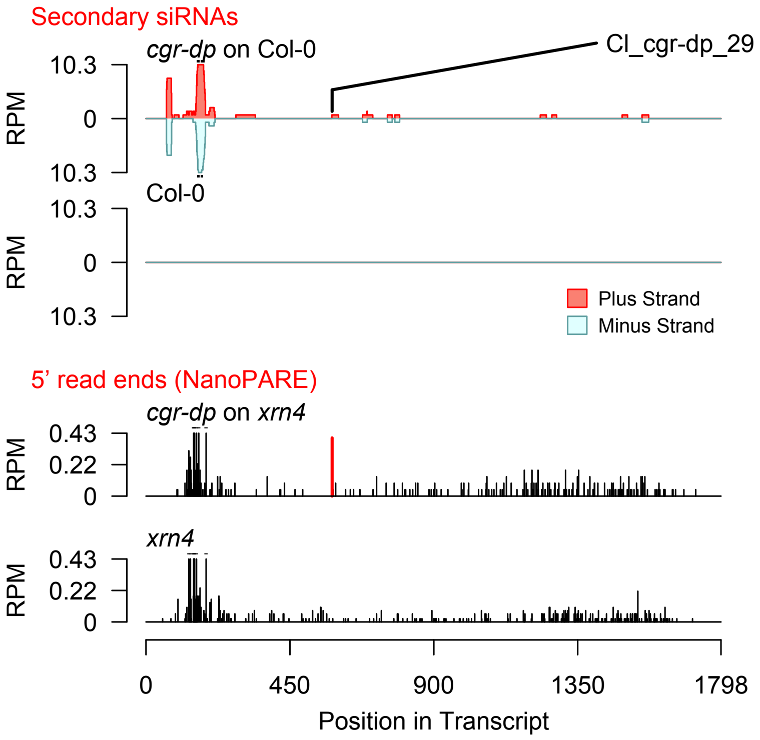


Diff. Exp. secondary siRNA locus not found

**Allenscore:** 3.5

**Target Interaction:**

```
5'- AGAGAGGUGGAGAUUCUUGA   AT3G14280.1
     ||||:|||||:||:|||||
3'- GCUCUUCACCUUUAGGAACU   Cl_ccm_1979
```

**Target Site:** 470

**Superfamily:** SupFam_20

**miRNA in ccm:** Y

**Confirmed Targeting**

| *2nd-siRNA* | *NanoPARE* |
|---|---|
| *ccm* | *ccm* |
| *cpe-2015* | *cpe-2015* |
| *cpe-2017* | |
| *cgr-dp* | *cgr-dp* |
| *cgr-pm* | |
| *cgr-mass* | |
| *cin* | *cin* |

### LL-diaminopimelate_aminotransferase

AT3G14280 - Cl_ccm_1979

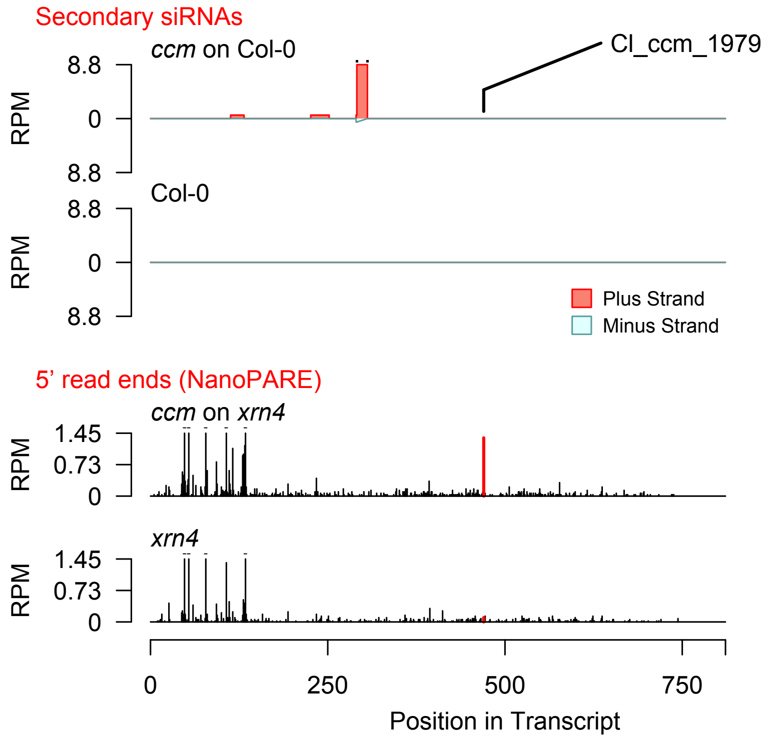


Diff. Exp. secondary siRNA locus not found

**Allenscore:** 3

**Target Interaction:**

```
5'- CUCGCAU - UCUUGAAAGAUUC    AT2G25170.1
      | | | | | |   | | | | | | | | | | | |
3'- AAGCGUAUAGAACUUUCUAAU    Cl_cgr-dp_80
```

**Target Site:** 3195

**Superfamily:** SupFam_206

**miRNA in ccm:** N/A

**Confirmed Targeting**

| 2nd-siRNA | NanoPARE |
|---|---|
| *ccm* | *ccm* |
| *cpe-2015* | *cpe-2015* |
| *cpe-2017* | |
| *cgr-dp* | *cgr-dp* |
| *cgr-pm* | |
| *cgr-mass* | |
| *cin* | *cin* |

### PKL

AT2G25170 - Cl_cgr-dp_80

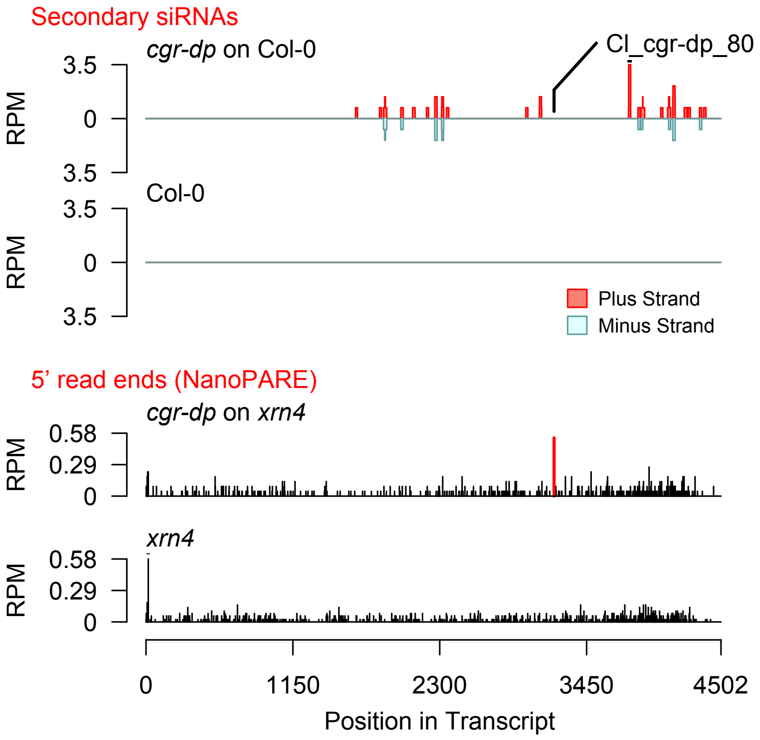


Diff. Exp. secondary siRNA locus not found

**Allenscore:** 4.5

**Target Interaction:**

```
5'- UCCAAACCGACACACACGGCAA    AT2G36570.1
    | | | | | | | | : | | | | | | | |     | | |
3'- AGGUUUGGUUGUGUGUGG-GUU    CI_cgr-pm_525
```

**Target Site:** 408

**Superfamily:** SupFam_161

**miRNA in ccm:** N/A

**Confirmed Targeting**

| *2nd-siRNA* | *NanoPARE* |
|---|---|
| *ccm* | *ccm* |
| *cpe-2015* | *cpe-2015* |
| *cpe-2017* | |
| *cgr-dp* | *cgr-dp* |
| *cgr-pm* | |
| *cgr-mass* | |
| *cin* | *cin* |

### PXC1

AT2G36570 - CI_cgr-pm_525

Secondary siRNAs

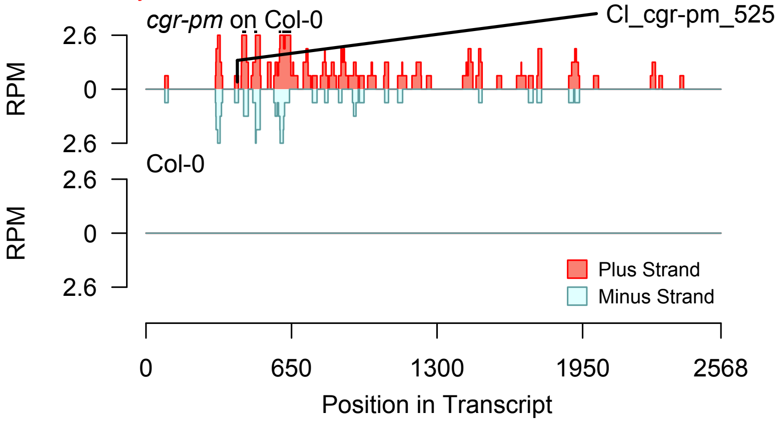


2nd siRNAs: Phase diagram

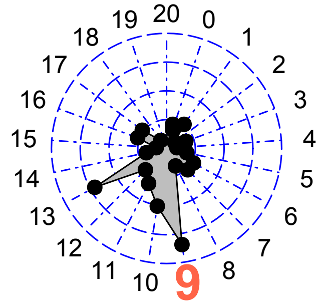


2nd siRNAs: Size distribution

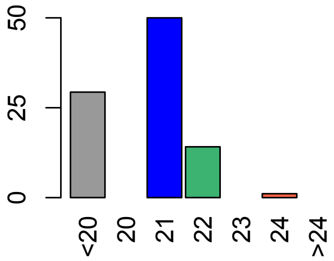


Degradome hits not found for sRNA

**Allenscore:** 4.5

**Target Interaction:**

```
5'- GCAGGGCAGACAUCUGAGAGGA   AT3G06380.1
     | | | | | | | : | | | |   : | | | | | | |
3'- AGUCCCGUUUGUAAGCUCUCCU   CI_cgr-dp_8594
```

**Target Site:** 1320

**Superfamily:** SupFam_14

**miRNA in ccm:** N/A

**Confirmed Targeting**

| *2nd-siRNA* | *NanoPARE* |
|---|---|
| *ccm* | *ccm* |
| *cpe-2015* | *cpe-2015* |
| *cpe-2017* | |
| *cgr-dp* | *cgr-dp* |
| *cgr-pm* | |
| *cgr-mass* | |
| *cin* | *cin* |

### TLP9

AT3G06380 - CI_cgr-dp_8594

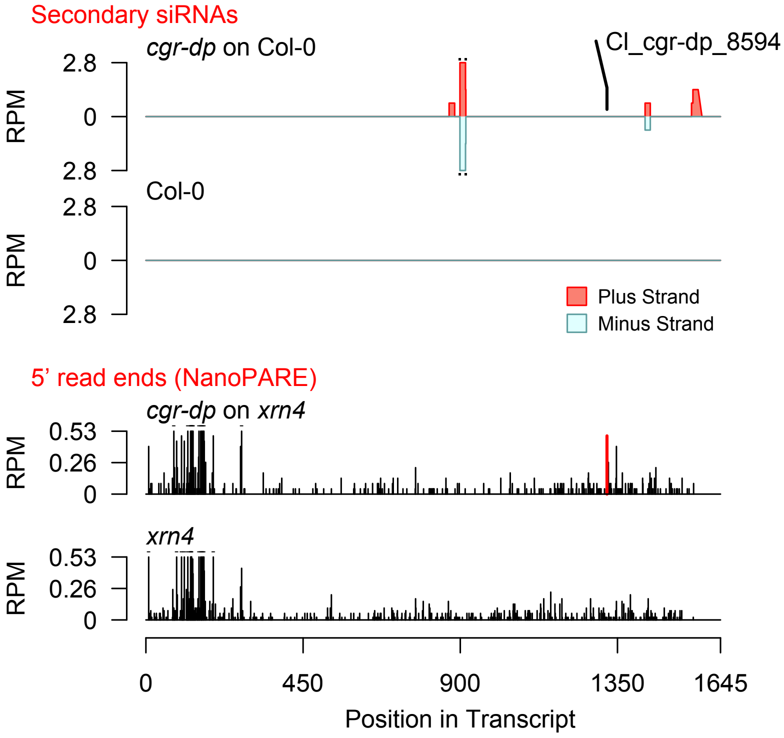


Diff. Exp. secondary siRNA locus not found

**Allenscore:** 5

**Target Interaction:**

```
5'- UGGCUCUUAACUUCUCAGUGU    AT5G65430.1
     | : | |    | | : | | | | | | | | | | | |
3'- AUCGUGAGUUGAAGAGUCAGU    CI_cpe-2015_129
```

**Target Site:** 885

**Superfamily:** SupFam_24

**miRNA in ccm:** Y

**Confirmed Targeting**

| *2nd-siRNA* | *NanoPARE* |
|---|---|
| *ccm* | *ccm* |
| *cpe-2015* | *cpe-2015* |
| *cpe-2017* | |
| *cgr-dp* | *cgr-dp* |
| *cgr-pm* | |
| *cgr-mass* | |
| *cin* | *cin* |

### GRF8

AT5G65430 - CI_cpe-2015_129

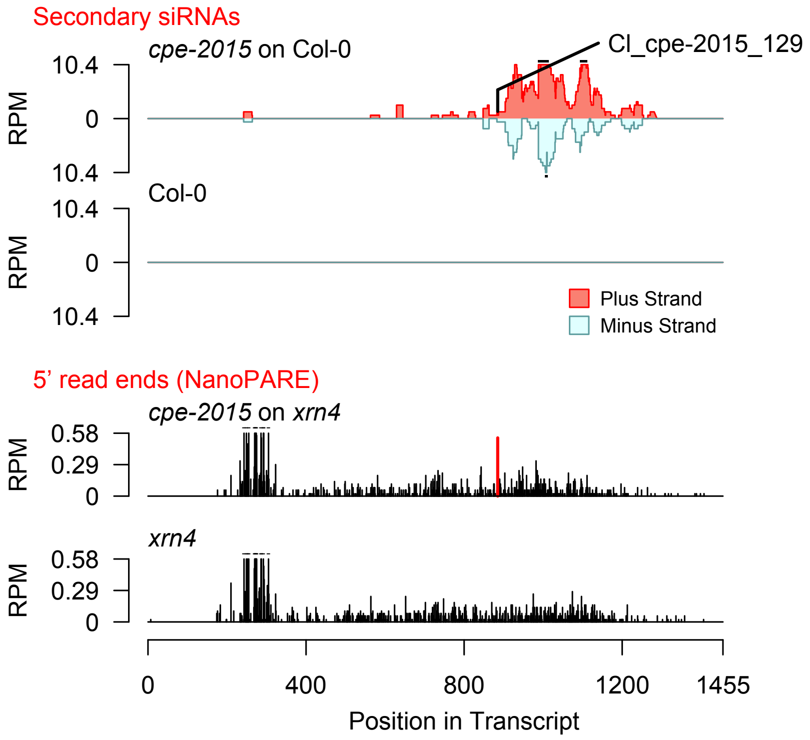


Diff. Exp. secondary siRNA locus not found

**Allenscore:** 4.5

**Target Interaction:**

```
5'- GAGGACUUCAUUACCUUCACA   AT5G24010.1
     : | | | | | | | |   | | | | | | | | | |
3'- UUCCUGAAGAUAUGGAAGUGU   Cl_cgr-dp_14631
```

**Target Site:** 1954

**Superfamily:** SupFam_49

**miRNA in ccm:** Y

**Confirmed Targeting**

| *2nd-siRNA* | *NanoPARE* |
|---|---|
| *ccm* | *ccm* |
| *cpe-2015* | *cpe-2015* |
| *cpe-2017* | |
| *cgr-dp* | *cgr-dp* |
| *cgr-pm* | |
| *cgr-mass* | |
| *cin* | *cin* |

### Protein_kinase_superfamily_protein

AT5G24010 - Cl_cgr-dp_14631

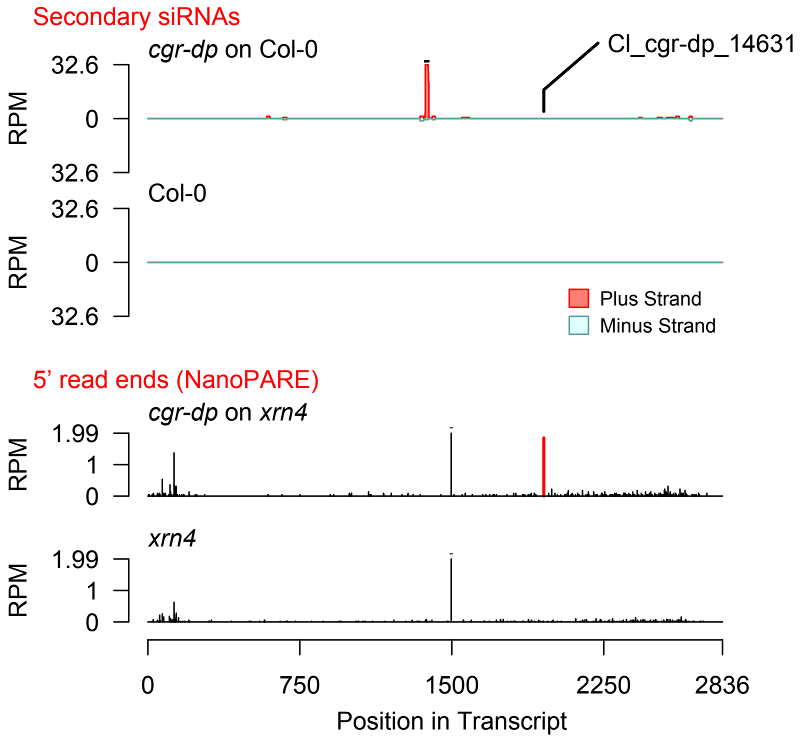


Diff. Exp. secondary siRNA locus not found

**Allenscore:** 4.5

**Target Interaction:**

```
5'- CUGGUGUGCAAGUCAUGGUACG   AT3G62980.1
     | |   | |   | : | | | : | | | | | | | | | :
3'- GAGCAGAUGUUUAGUACCAUGU   Cl_ccm_2
```

**Target Site:** 555

**Superfamily:** SupFam_27

**miRNA in ccm:** Y

**Confirmed Targeting**

| *2nd-siRNA* | *NanoPARE* |
|---|---|
| *ccm* | *ccm* |
| *cpe-2015* | *cpe-2015* |
| *cpe-2017* | |
| *cgr-dp* | *cgr-dp* |
| *cgr-pm* | |
| *cgr-mass* | |
| *cin* | *cin* |

### TIR1

AT3G62980 - Cl_ccm_2

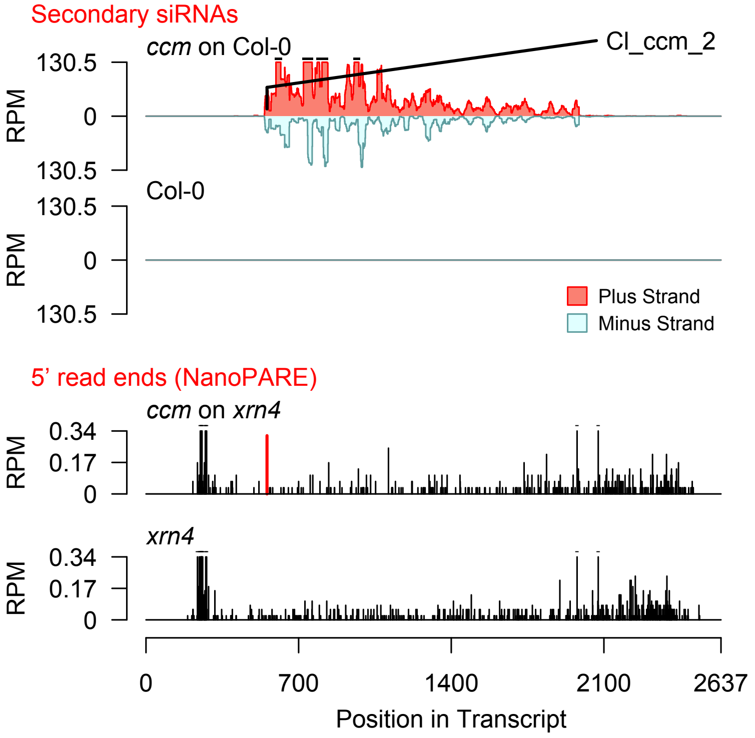


2nd siRNAs: Phase diagram

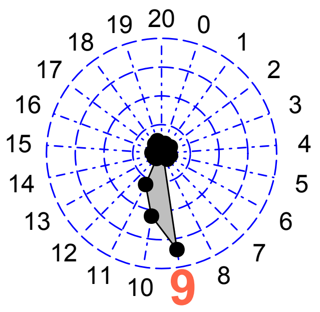


2nd siRNAs: Size distribution

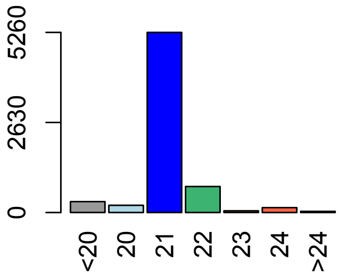

**Allenscore:** 2.5

**Target Interaction:**

```
5'- CUGGUGUGCAAGUCAUGGUACG   AT3G62980.1
      | : | | | | | | | | | | | | | | | | | | :
3'- AAUCACACGUUCAGUACCAUGU   CI_ccm_438
```

**Target Site:** 555

**Superfamily:** SupFam_27

**miRNA in ccm:** Y

**Confirmed Targeting**

| 2nd-siRNA | NanoPARE |
|---|---|
| *ccm* | *ccm* |
| *cpe-2015* | *cpe-2015* |
| *cpe-2017* | |
| *cgr-dp* | *cgr-dp* |
| *cgr-pm* | |
| *cgr-mass* | |
| *cin* | *cin* |

### TIR1

AT3G62980 - CI_ccm_438

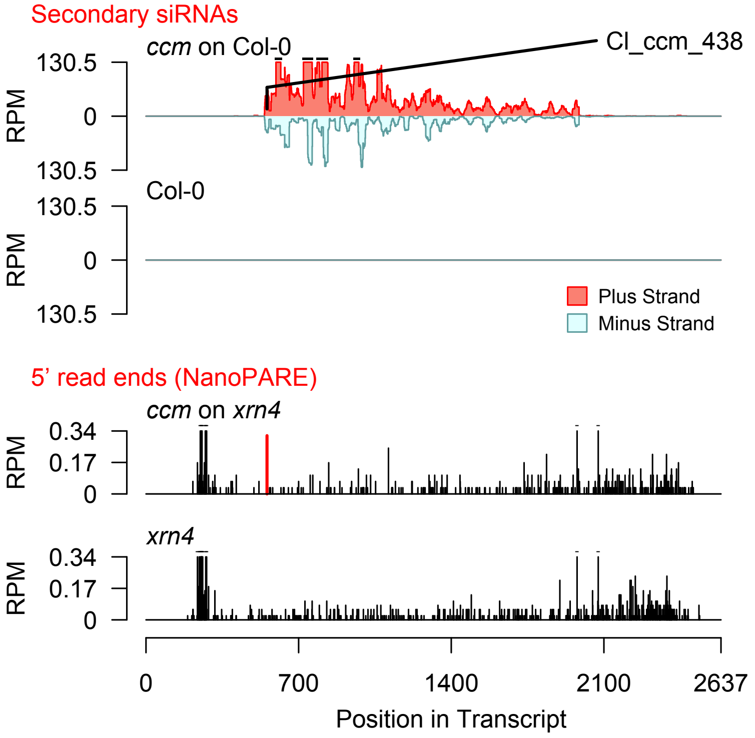


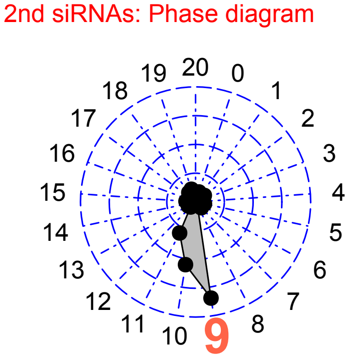


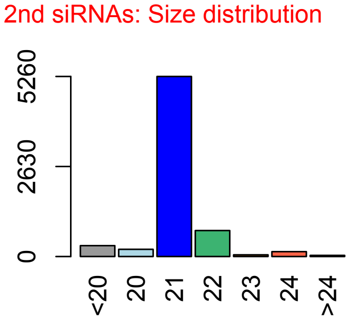

**Allenscore:** 3.5

**Target Interaction:**

```
5'- CUGGUGUGCAAGUCAUGGUACG    AT3G62980.1
     | | | | : | | | | | | : | | | | | | | | | :
3'- AACCAUACGUUUAGUACCAUGU    CI_ccm_614
```

**Target Site:** 555

**Superfamily:** SupFam_27

**miRNA in ccm:** Y

**Confirmed Targeting**

| *2nd-siRNA* | *NanoPARE* |
|---|---|
| *ccm* | *ccm* |
| *cpe-2015* | *cpe-2015* |
| *cpe-2017* | |
| *cgr-dp* | *cgr-dp* |
| *cgr-pm* | |
| *cgr-mass* | |
| *cin* | *cin* |

### TIR1

AT3G62980 - CI_ccm_614

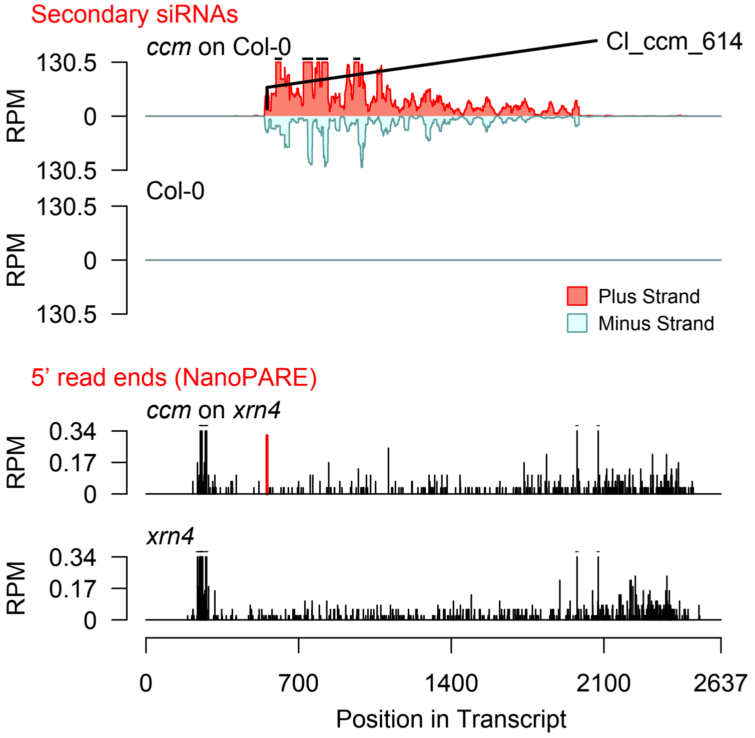


2nd siRNAs: Phase diagram

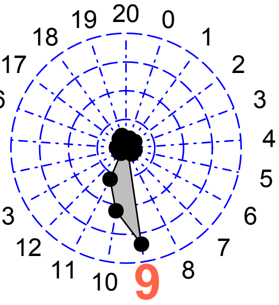


2nd siRNAs: Size distribution

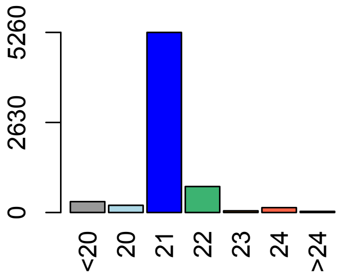

**Allenscore:** 4

**Target Interaction:**

```
5'- CUGGUGUGCAAGUCAUGGUACG   AT3G62980.1
    ||||||||||||:||||||||  :
3'- GACCACACGUUUAGUACCAUAU   CI_ccm_13108
```

**Target Site:** 555

**Superfamily:** SupFam_27

**miRNA in ccm:** Y

**Confirmed Targeting**

| *2nd-siRNA* | *NanoPARE* |
|---|---|
| *ccm* | *ccm* |
| *cpe-2015* | *cpe-2015* |
| *cpe-2017* | |
| *cgr-dp* | *cgr-dp* |
| *cgr-pm* | |
| *cgr-mass* | |
| *cin* | *cin* |

### TIR1

AT3G62980 - CI_ccm_13108

Secondary siRNAs

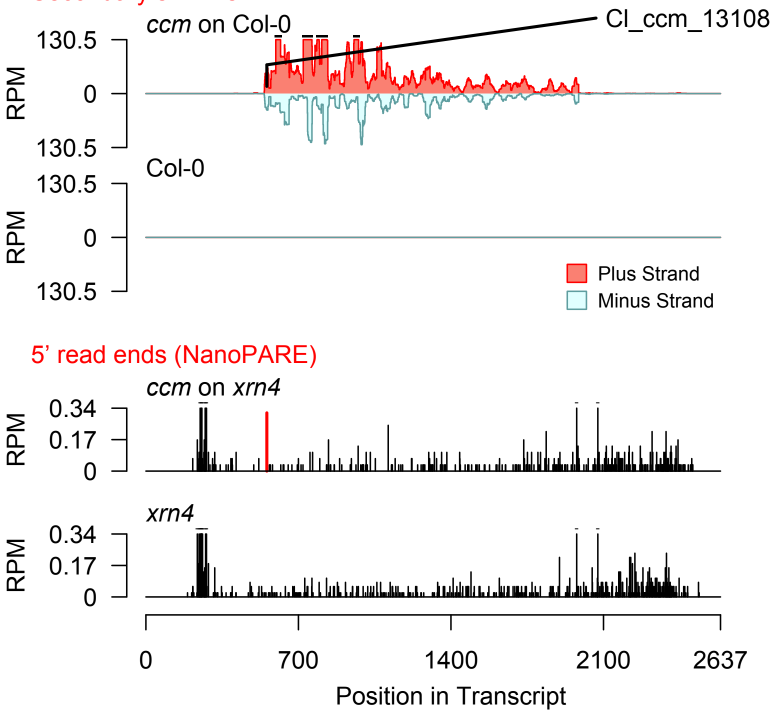


2nd siRNAs: Phase diagram

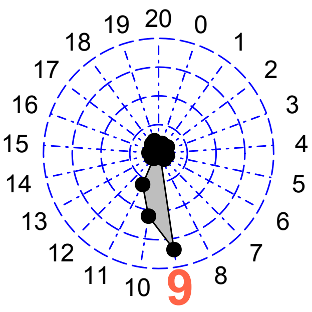


2nd siRNAs: Size distribution

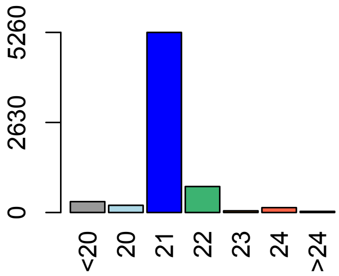

**Allenscore:** 1

**Target Interaction:**

```
5'- GGUGUGCAAGUCAUGGUACG   AT3G62980.1
    | | | | | | | | | | | | | | | | | | | :
3'- CCACACGUUCAGUACCAUGU   Cl_ccm_51538
```

**Target Site:** 555

**Superfamily:** SupFam_27

**miRNA in ccm:** Y

**Confirmed Targeting**

| *2nd-siRNA* | *NanoPARE* |
|---|---|
| *ccm* | *ccm* |
| *cpe-2015* | *cpe-2015* |
| *cpe-2017* | |
| *cgr-dp* | *cgr-dp* |
| *cgr-pm* | |
| *cgr-mass* | |
| *cin* | *cin* |

### TIR1

AT3G62980 - Cl_ccm_51538

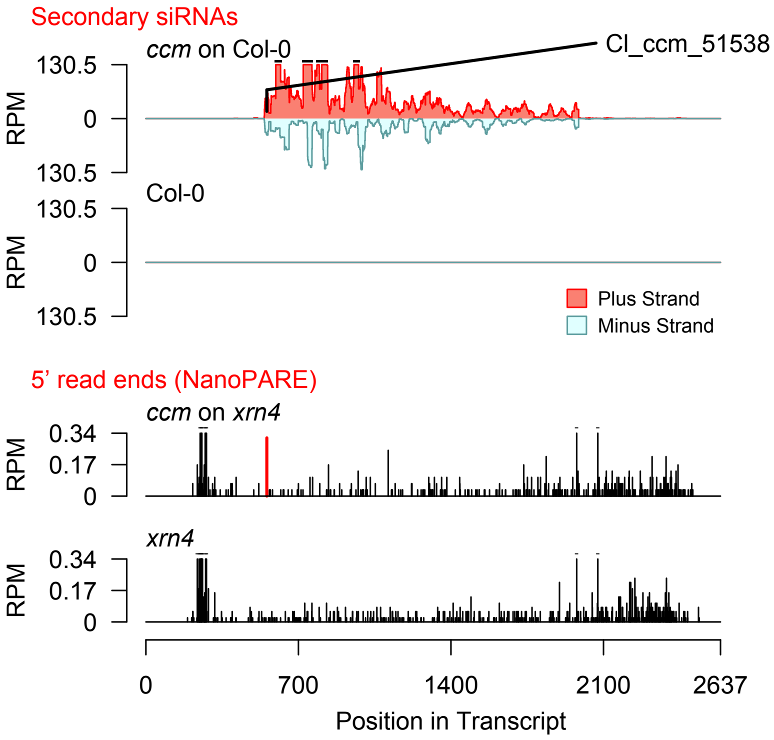


2nd siRNAs: Phase diagram

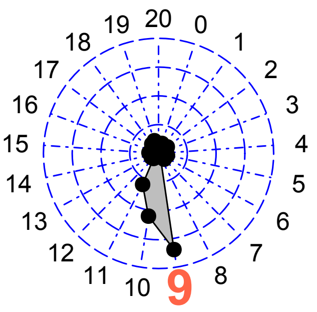


2nd siRNAs: Size distribution

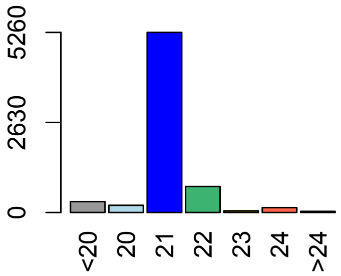

**Allenscore:** 4

**Target Interaction:**

```
5'- CUGGUGUGCAAGUCAUGGUACG   AT3G62980.1
     ||  ||  |||||:|||||||||:
3'- GAGCAGACGUUUAGUACCAUGU   Cl_cpe-2015_7
```

**Target Site:** 555

**Superfamily:** SupFam_27

**miRNA in ccm:** Y

**Confirmed Targeting**

| *2nd-siRNA* | *NanoPARE* |
|---|---|
| *ccm* | *ccm* |
| *cpe-2015* | *cpe-2015* |
| *cpe-2017* | |
| *cgr-dp* | *cgr-dp* |
| *cgr-pm* | |
| *cgr-mass* | |
| *cin* | *cin* |

### TIR1

AT3G62980 - Cl_cpe-2015_7

Secondary siRNAs

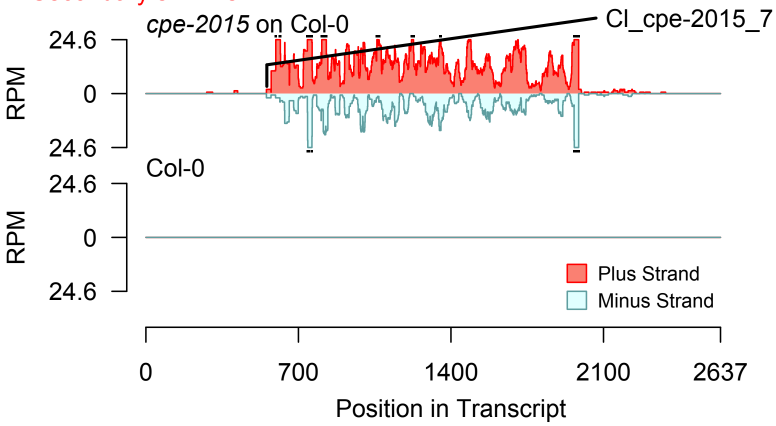


2nd siRNAs: Phase diagram

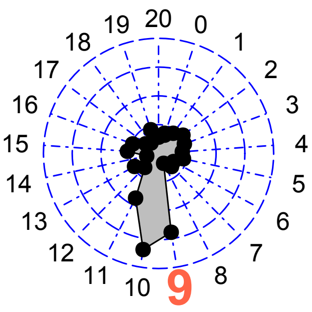


2nd siRNAs: Size distribution

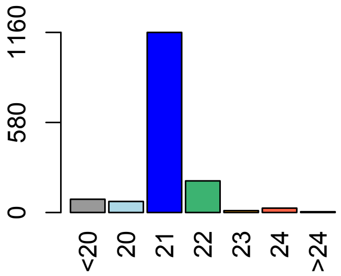


Degradome hits not found for sRNA

**Allenscore:** 4

**Target Interaction:**

```
5'- CUGGUGUGCAAGUCAUGGUACG    AT3G62980.1
     | |   | |   | | | | | : | | | | | | | | | | :
3'- GAGCAGACGUUUAGUACCAUGU    Cl_cpe-2017_11
```

**Target Site:** 555

**Superfamily:** SupFam_27

**miRNA in ccm:** Y

**Confirmed Targeting**

| *2nd-siRNA* | *NanoPARE* |
|---|---|
| *ccm* | *ccm* |
| *cpe-2015* | *cpe-2015* |
| *cpe-2017* | |
| *cgr-dp* | *cgr-dp* |
| *cgr-pm* | |
| *cgr-mass* | |
| *cin* | *cin* |

### TIR1

AT3G62980 - Cl_cpe-2017_11

Secondary siRNAs

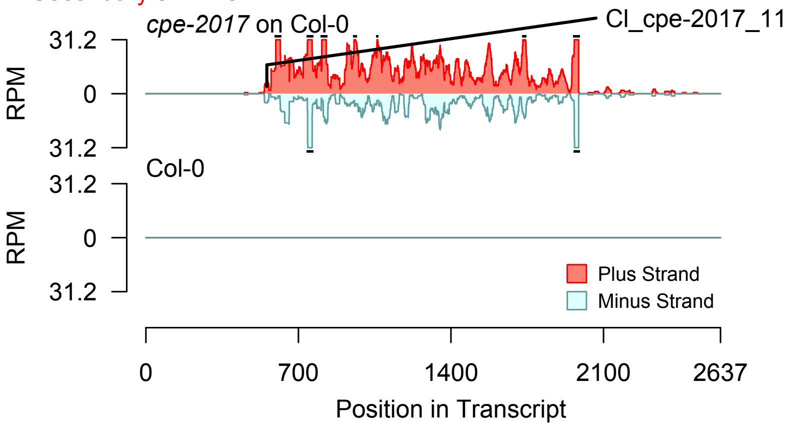


Degradome hits not found for sRNA

2nd siRNAs: Phase diagram

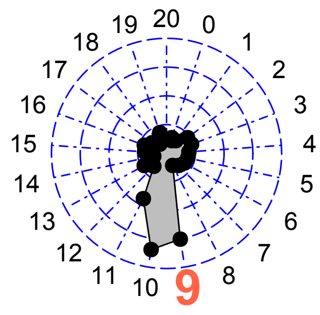


2nd siRNAs: Size distribution

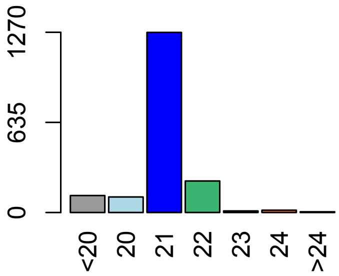

**Allenscore:** 5.5

**Target Interaction:**

```
5'- UGGUGAAAAGUGGAAAUCCUUGU   AT1G20696.1
      :   : | |   | | | | | | | | | | | | | | |
3'- UUAGCUCUUCACCUUUAGGAACU   CI_ccm_4973
```

**Target Site:** 572

**Superfamily:** SupFam_20

**miRNA in ccm:** Y

**Confirmed Targeting**

| *2nd-siRNA* | *NanoPARE* |
|---|---|
| *ccm* | *ccm* |
| *cpe-2015* | *cpe-2015* |
| *cpe-2017* | |
| *cgr-dp* | *cgr-dp* |
| *cgr-pm* | |
| *cgr-mass* | |
| *cin* | *cin* |

### HMGB3

AT1G20696 - CI_ccm_4973

Secondary siRNAs

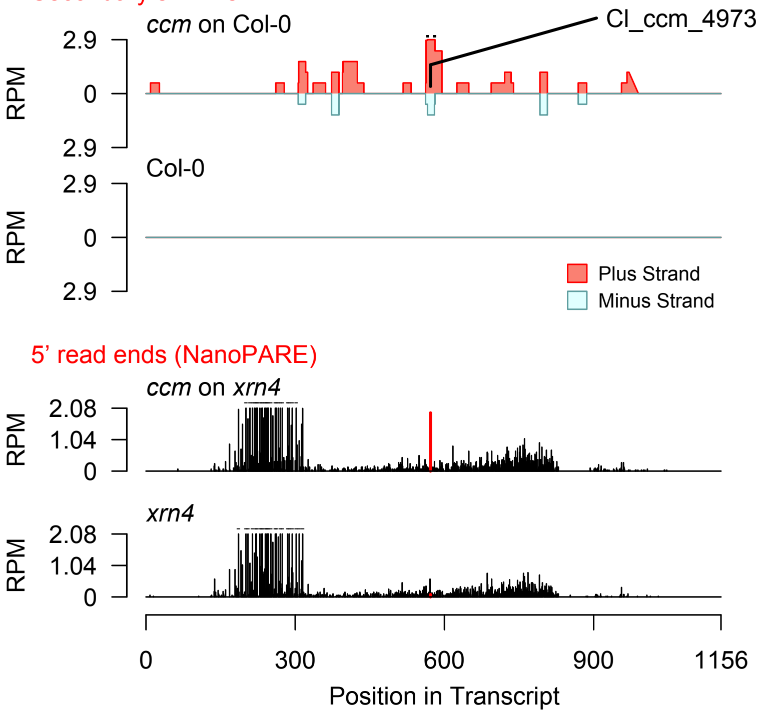


Diff. Exp. secondary siRNA locus not found

**Allenscore:** 2.5

**Target Interaction:**

```
5'- UGAAAAGUGGAAAUCCUUGU   AT1G20696.1
     : | |   | | | | | | | | | | | | | | |
3'- GCUCUUCACCUUUAGGAACU   CI_ccm_1979
```

**Target Site:** 572

**Superfamily:** SupFam_20

**miRNA in ccm:** Y

**Confirmed Targeting**

| *2nd-siRNA* | *NanoPARE* |
|---|---|
| *ccm* | *ccm* |
| *cpe-2015* | *cpe-2015* |
| *cpe-2017* | |
| *cgr-dp* | *cgr-dp* |
| *cgr-pm* | |
| *cgr-mass* | |
| *cin* | *cin* |

### HMGB3

AT1G20696 - CI_ccm_1979

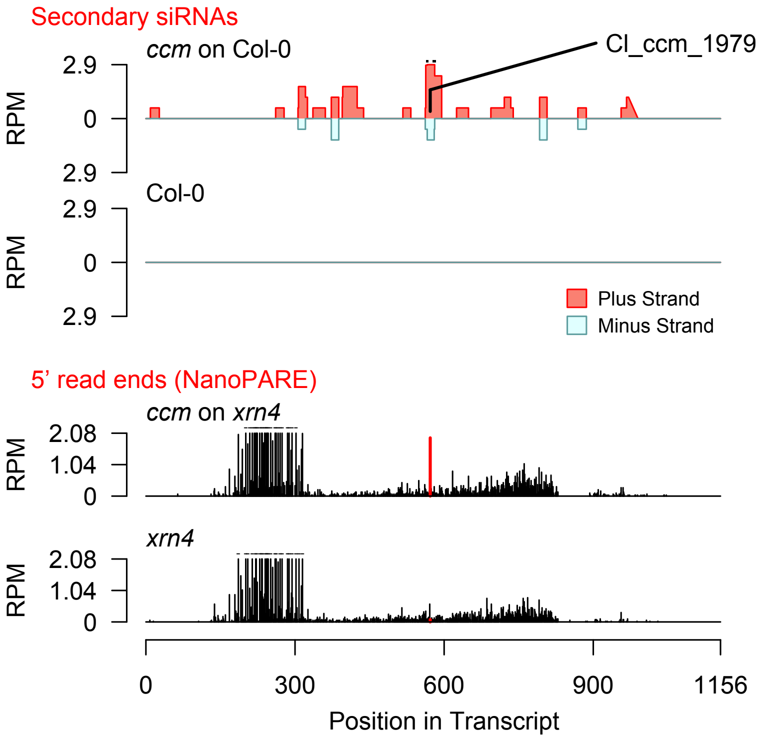


Diff. Exp. secondary siRNA locus not found

**Allenscore:** 3

**Target Interaction:**

```
5'- UCUGGUCUGCAAGUCAUGGU   AT4G03190.1
      ||  ||||||||:|||||||
3'- UGAGCAGACGUUUAGUACCA   CI_cpe-2015_526
```

**Target Site:** 436

**Superfamily:** SupFam_27

**miRNA in ccm:** Y

**Confirmed Targeting**

| *2nd-siRNA* | *NanoPARE* |
|---|---|
| *ccm* | *ccm* |
| *cpe-2015* | *cpe-2015* |
| *cpe-2017* | |
| *cgr-dp* | *cgr-dp* |
| *cgr-pm* | |
| *cgr-mass* | |
| *cin* | *cin* |

### AFB1

AT4G03190 - CI_cpe-2015_526

Secondary siRNAs

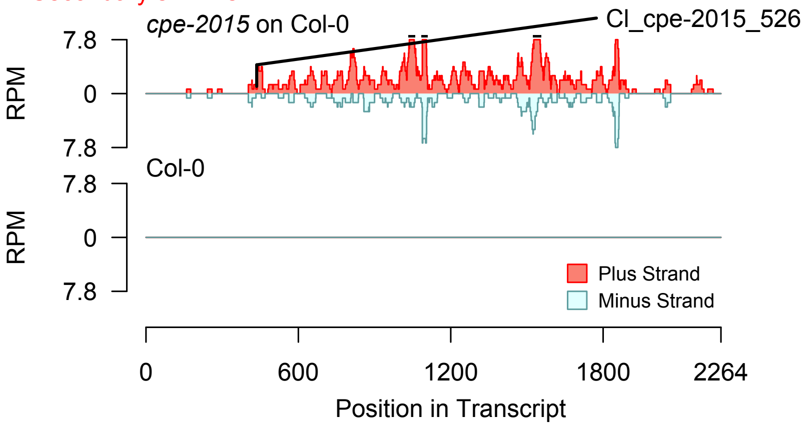


2nd siRNAs: Phase diagram

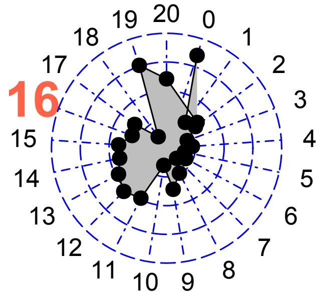


2nd siRNAs: Size distribution

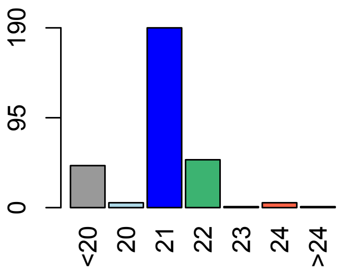


Degradome hits not found for sRNA

**Allenscore:** 3

**Target Interaction:**

```
5'- UCUGGUCUGCAAGUCAUGGU   AT4G03190.1
      ||  ||||||||:|||||||
3'- UGAGCAGACGUUUAGUACCA   CI_cpe-2017_486
```

**Target Site:** 436

**Superfamily:** SupFam_27

**miRNA in ccm:** Y

**Confirmed Targeting**

| *2nd-siRNA* | *NanoPARE* |
|---|---|
| *ccm* | *ccm* |
| *cpe-2015* | *cpe-2015* |
| *cpe-2017* | |
| *cgr-dp* | *cgr-dp* |
| *cgr-pm* | |
| *cgr-mass* | |
| *cin* | *cin* |

### AFB1

AT4G03190 - CI_cpe-2017_486

Secondary siRNAs

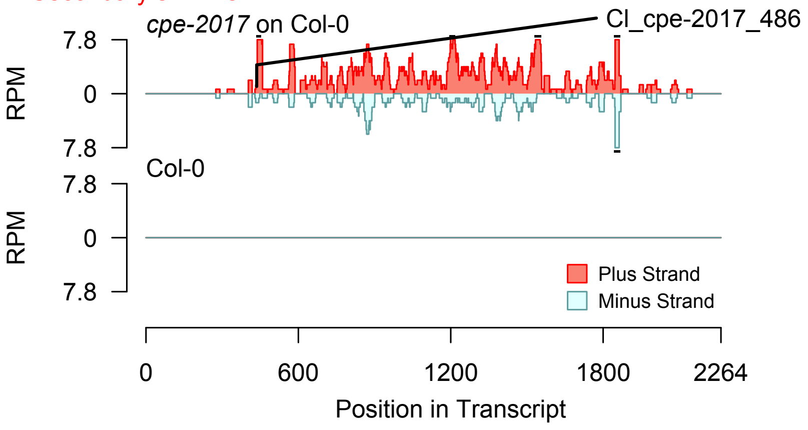


2nd siRNAs: Phase diagram

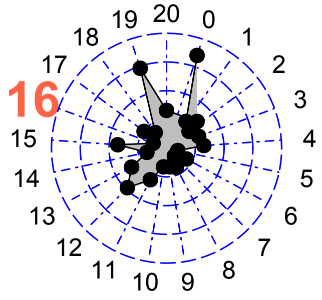


2nd siRNAs: Size distribution

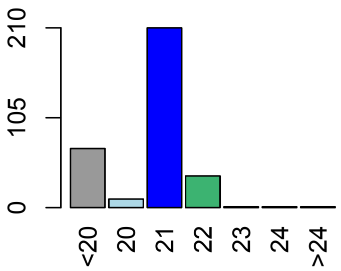


Degradome hits not found for sRNA

**Allenscore:** 5

**Target Interaction:**

```
5'- UUCUCUGGUCUGCAAGUCAUGG   AT4G03190.1
        |||  |||||||||:||||||
3'- UUAAGAGCAGACGUUUAGUACC   Cl_cpe-2017_29015
```

**Target Site:** 435

**Superfamily:** SupFam_27

**miRNA in ccm:** Y

**Confirmed Targeting**

| *2nd-siRNA* | *NanoPARE* |
|---|---|
| *ccm* | *ccm* |
| *cpe-2015* | *cpe-2015* |
| *cpe-2017* | |
| *cgr-dp* | *cgr-dp* |
| *cgr-pm* | |
| *cgr-mass* | |
| *cin* | *cin* |

### AFB1

AT4G03190 - Cl_cpe-2017_29015

Secondary siRNAs

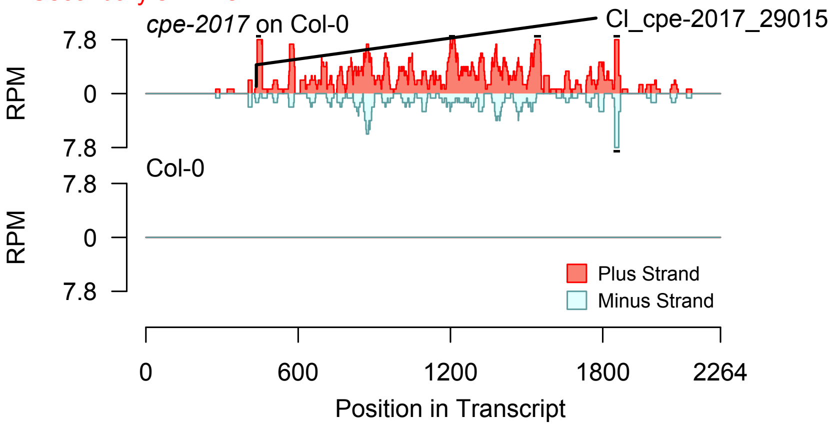


Degradome hits not found for sRNA

2nd siRNAs: Phase diagram

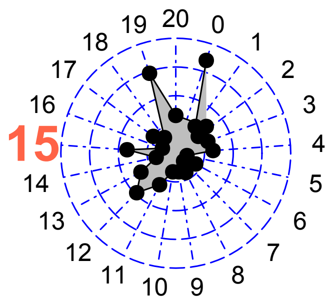


2nd siRNAs: Size distribution

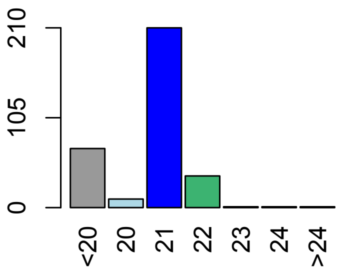

**Allenscore:** 2

**Target Interaction:**

```
5'- CUUGUAUGCAAAUCAUGGUACA   AT3G26810.1
     | | : | |   | : | | | | | | | | | | | | | |
3'- GAGCAGAUGUUUAGUACCAUGU   CI_ccm_2
```

**Target Site:** 781

**Superfamily:** SupFam_27

**miRNA in ccm:** Y

**Confirmed Targeting**

| *2nd-siRNA* | *NanoPARE* |
|---|---|
| *ccm* | *ccm* |
| *cpe-2015* | *cpe-2015* |
| *cpe-2017* | |
| *cgr-dp* | *cgr-dp* |
| *cgr-pm* | |
| *cgr-mass* | |
| *cin* | *cin* |

### AFB2

AT3G26810 - CI_ccm_2

Secondary siRNAs

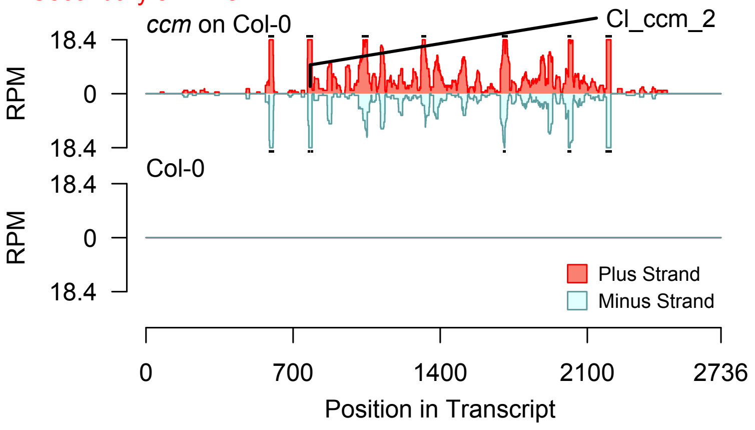


2nd siRNAs: Phase diagram

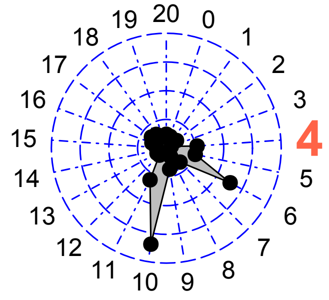


2nd siRNAs: Size distribution

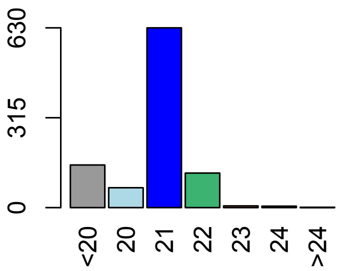


Degradome hits not found for sRNA

**Allenscore:** 3

**Target Interaction:**

```
5'- CUUGUAUGCAAAUCAUGGUACA   AT3G26810.1
      |  | | | | | | | | | | | | | | | | | | |
3'- AACCAUACGUUUAGUACCAUGU   Cl_ccm_614
```

**Target Site:** 781

**Superfamily:** SupFam_27

**miRNA in ccm:** Y

**Confirmed Targeting**

| *2nd-siRNA* | *NanoPARE* |
|---|---|
| *ccm* | *ccm* |
| *cpe-2015* | *cpe-2015* |
| *cpe-2017* | |
| *cgr-dp* | *cgr-dp* |
| *cgr-pm* | |
| *cgr-mass* | |
| *cin* | *cin* |

### AFB2

AT3G26810 - Cl_ccm_614

Secondary siRNAs

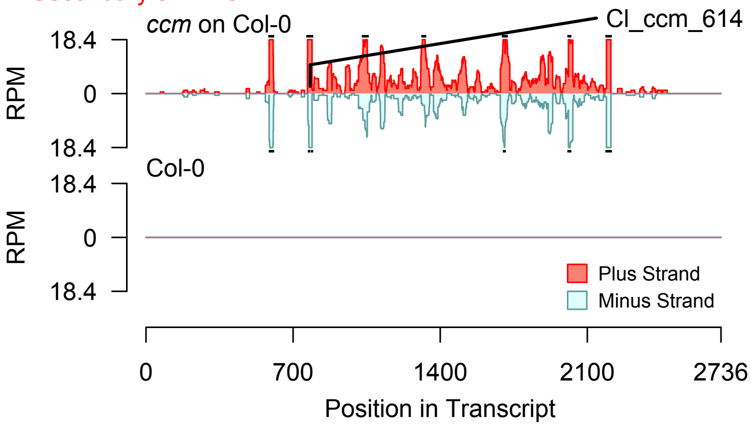


2nd siRNAs: Phase diagram

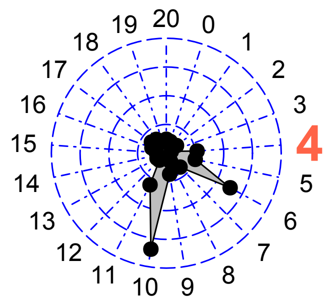


2nd siRNAs: Size distribution

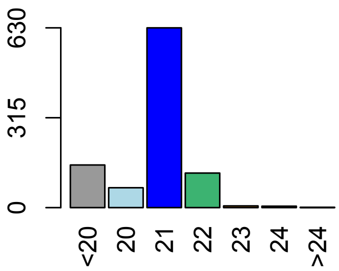


Degradome hits not found for sRNA

**Allenscore:** 1.5

**Target Interaction:**

```
5'- CUUGUAUGCAAAUCAUGGUACA    AT3G26810.1
     | | : | |   | | | | | | | | | | | | | | | |
3'- GAGCAGACGUUUAGUACCAUGU    Cl_cpe-2015_7
```

**Target Site:** 781

**Superfamily:** SupFam_27

**miRNA in ccm:** Y

**Confirmed Targeting**

| *2nd-siRNA* | *NanoPARE* |
|---|---|
| *ccm* | *ccm* |
| *cpe-2015* | *cpe-2015* |
| *cpe-2017* | |
| *cgr-dp* | *cgr-dp* |
| *cgr-pm* | |
| *cgr-mass* | |
| *cin* | *cin* |

### AFB2

AT3G26810 - Cl_cpe-2015_7

Secondary siRNAs

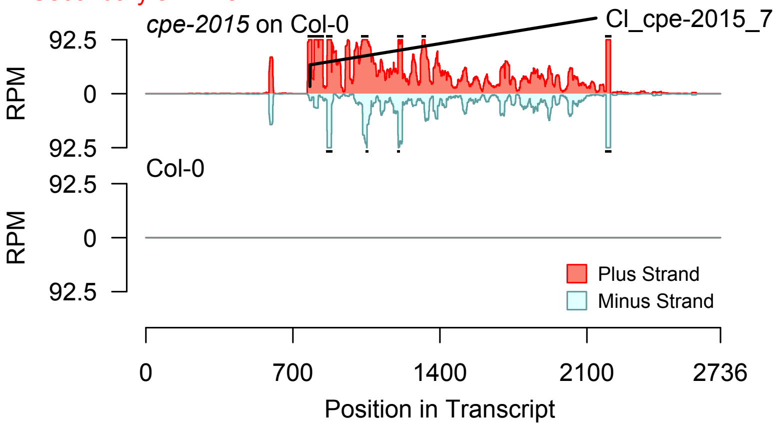


2nd siRNAs: Phase diagram

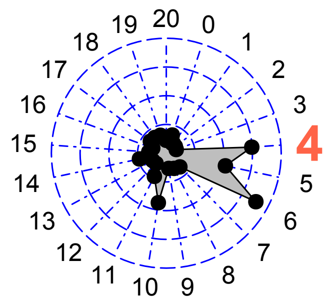


2nd siRNAs: Size distribution

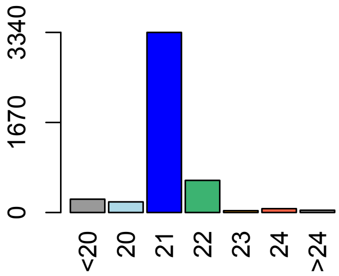


Degradome hits not found for sRNA

**Allenscore:** 1.5

**Target Interaction:**

```
5'- CUUGUAUGCAAAUCAUGGUACA    AT3G26810.1
     | | : | |   | | | | | | | | | | | | | | |
3'- GAGCAGACGUUUAGUACCAUGU    Cl_cpe-2017_11
```

**Target Site:** 781

**Superfamily:** SupFam_27

**miRNA in ccm:** Y

**Confirmed Targeting**

| *2nd-siRNA* | *NanoPARE* |
|---|---|
| *ccm* | *ccm* |
| *cpe-2015* | *cpe-2015* |
| *cpe-2017* | |
| *cgr-dp* | *cgr-dp* |
| *cgr-pm* | |
| *cgr-mass* | |
| *cin* | *cin* |

### AFB2

AT3G26810 - Cl_cpe-2017_11

Secondary siRNAs

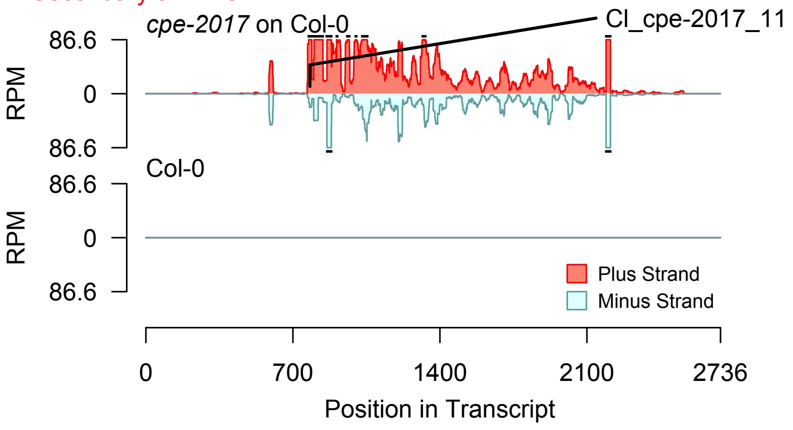


Degradome hits not found for sRNA

2nd siRNAs: Phase diagram

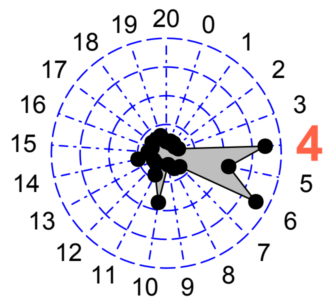


2nd siRNAs: Size distribution

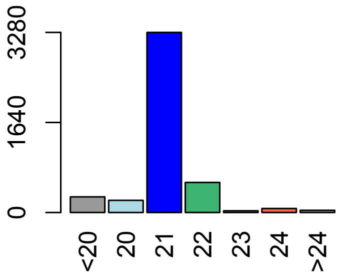

**Allenscore:** 1.5

**Target Interaction:**

```
5'- GGGAUGAGAAAGUGAAGGUUCU    AT3G27300.1
     | : | | | | | | | | | | | | | | | | | | |
3'- CUCUACUCUUUCACUUCCAAGU    CI_cpe-2015_537
```

**Target Site:** 1099

**Superfamily:** SupFam_312

**miRNA in ccm:** Y

**Confirmed Targeting**

| *2nd-siRNA* | *NanoPARE* |
|---|---|
| *ccm* | *ccm* |
| *cpe-2015* | *cpe-2015* |
| *cpe-2017* | |
| *cgr-dp* | *cgr-dp* |
| *cgr-pm* | |
| *cgr-mass* | |
| *cin* | *cin* |

# G6PD5

AT3G27300 - CI_cpe-2015_537

Secondary siRNAs

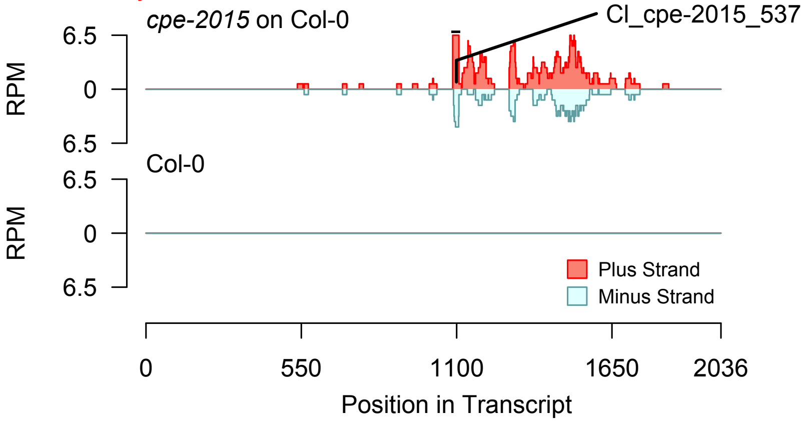


2nd siRNAs: Phase diagram

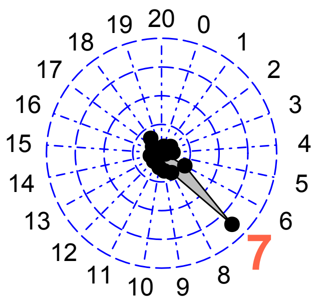


2nd siRNAs: Size distribution

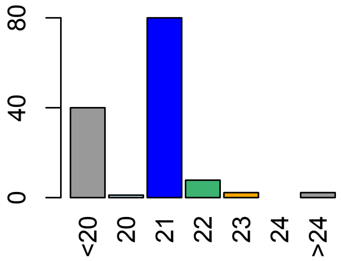


Degradome hits not found for sRNA

**Allenscore:** 1.5

**Target Interaction:**

```
5'- GGGAUGAGAAAGUGAAGGUUCU   AT3G27300.1
     | : | | | | | | | | | | | | | | | | | | |
3'- CUCUACUCUUUCACUUCCAAGU   Cl_cpe-2017_615
```

**Target Site:** 1099

**Superfamily:** SupFam_312

**miRNA in ccm:** Y

**Confirmed Targeting**

| *2nd-siRNA* | *NanoPARE* |
|---|---|
| *ccm* | *ccm* |
| *cpe-2015* | *cpe-2015* |
| *cpe-2017* | |
| *cgr-dp* | *cgr-dp* |
| *cgr-pm* | |
| *cgr-mass* | |
| *cin* | *cin* |

# G6PD5

AT3G27300 - Cl_cpe-2017_615

Secondary siRNAs

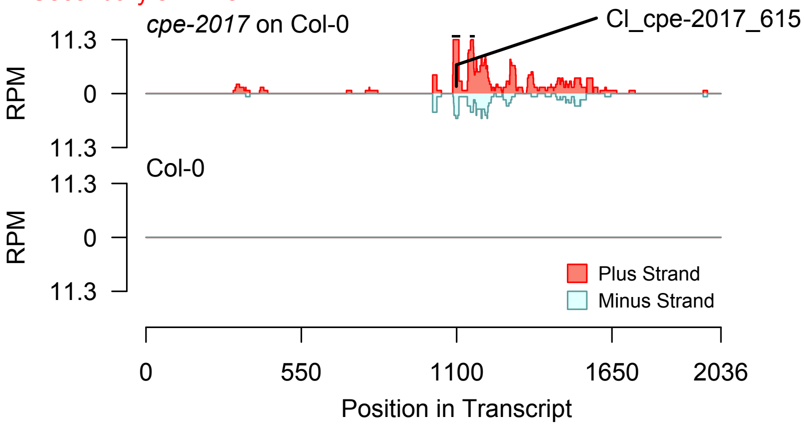


2nd siRNAs: Phase diagram

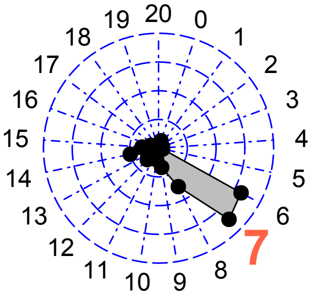


2nd siRNAs: Size distribution

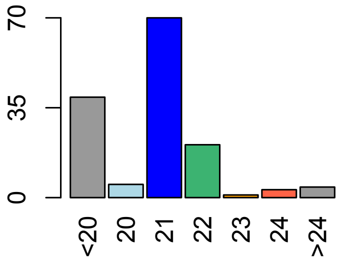


Degradome hits not found for sRNA

**Allenscore:** 1.5

**Target Interaction:**

```
5'- GGGAUGAGAAAGUGAAGGUUCU   AT3G27300.1
     | : | | | | | | | | | | | | | | | | | | | |
3'- CUCUACUCUUUCACUUCCAAGU   CI_ccm_5718
```

**Target Site:** 1099

**Superfamily:** SupFam_312

**miRNA in ccm:** Y

**Confirmed Targeting**

| *2nd-siRNA* | *NanoPARE* |
|---|---|
| *ccm* | *ccm* |
| *cpe-2015* | *cpe-2015* |
| *cpe-2017* | |
| *cgr-dp* | *cgr-dp* |
| *cgr-pm* | |
| *cgr-mass* | |
| *cin* | *cin* |

# G6PD5

AT3G27300 - CI_ccm_5718

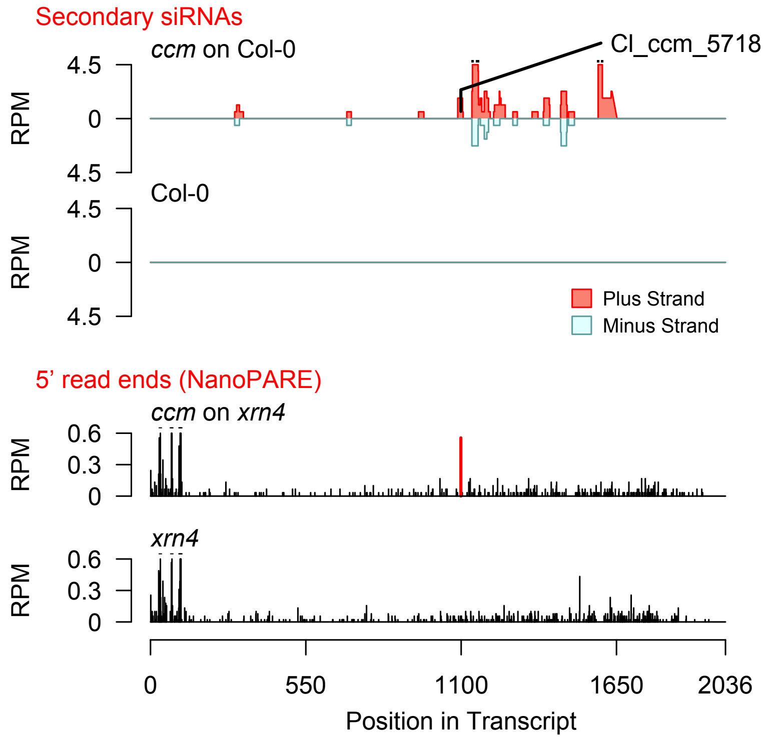


Diff. Exp. secondary siRNA locus not found

**Allenscore:** 4

**Target Interaction:**

```
5'- CUGGUUACUUAGACCCAGAGUA   AT4G00330.1
      |||||||||||| |||||||
3'- AACCAAUGAAUCUAGGUCUCAC   Cl_cpe-2015_439
```

**Target Site:** 1015

**Superfamily:** SupFam_257

**miRNA in ccm:** N/A

**Confirmed Targeting**

| *2nd-siRNA* | *NanoPARE* |
|---|---|
| *ccm* | *ccm* |
| *cpe-2015* | *cpe-2015* |
| *cpe-2017* | |
| *cgr-dp* | *cgr-dp* |
| *cgr-pm* | |
| *cgr-mass* | |
| *cin* | *cin* |

### CRCK2

AT4G00330 - Cl_cpe-2015_439

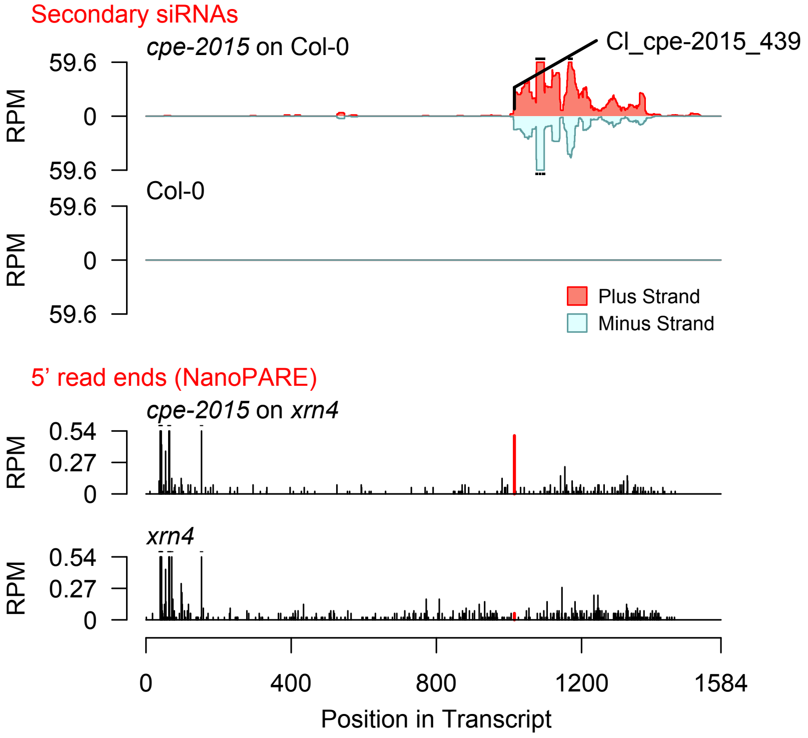


2nd siRNAs: Phase diagram

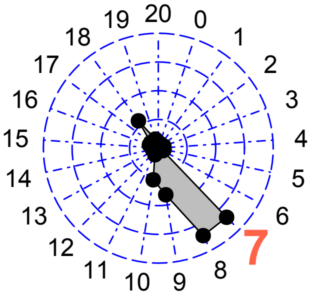


2nd siRNAs: Size distribution

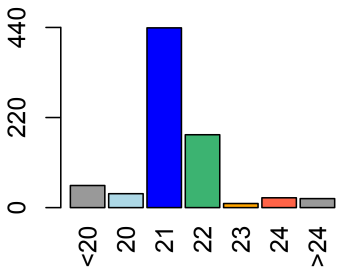

**Allenscore:** 3.5

**Target Interaction:**

```
5'- GGUUACUUAGACCCAGAGUA   AT4G00330.1
     : | | | | | | | | | | |   | | | | | | |
3'- UCAAUGAAUCUAGGUCUCAC   CI_cpe-2015_45087
```

**Target Site:** 1015

**Superfamily:** SupFam_257

**miRNA in ccm:** N/A

**Confirmed Targeting**

| *2nd-siRNA* | *NanoPARE* |
|---|---|
| *ccm* | *ccm* |
| *cpe-2015* | *cpe-2015* |
| *cpe-2017* | |
| *cgr-dp* | *cgr-dp* |
| *cgr-pm* | |
| *cgr-mass* | |
| *cin* | *cin* |

### CRCK2

AT4G00330 - CI_cpe-2015_45087

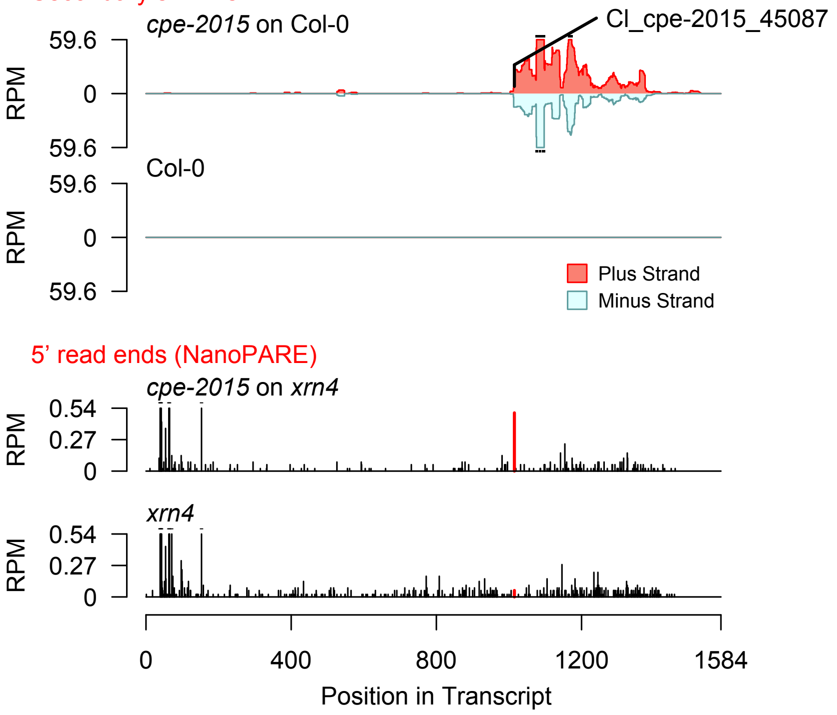


2nd siRNAs: Phase diagram

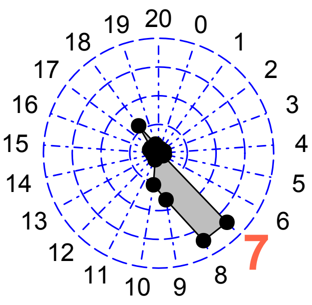


2nd siRNAs: Size distribution

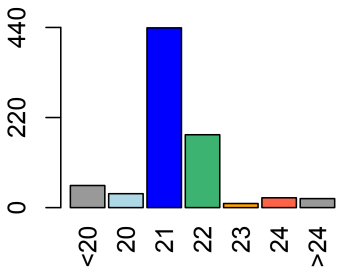

**Allenscore:** 4

**Target Interaction:**

```
5'- CUGGUUACUUAGACCCAGAGUA   AT4G00330.1
      |||||||||||| |||||||
3'- AACCAAUGAAUCUAGGUCUCAC   CI_cpe-2017_503
```

**Target Site:** 1015

**Superfamily:** SupFam_257

**miRNA in ccm:** N/A

**Confirmed Targeting**

| *2nd-siRNA* | *NanoPARE* |
|---|---|
| *ccm* | *ccm* |
| *cpe-2015* | *cpe-2015* |
| *cpe-2017* | |
| *cgr-dp* | *cgr-dp* |
| *cgr-pm* | |
| *cgr-mass* | |
| *cin* | *cin* |

### CRCK2

AT4G00330 - CI_cpe-2017_503

Secondary siRNAs

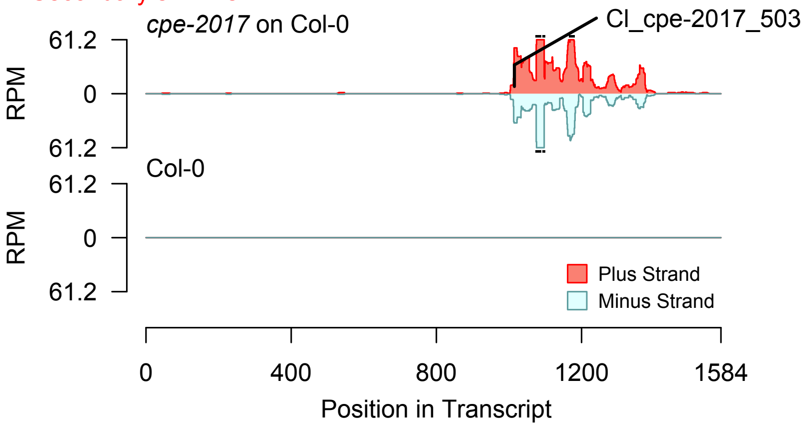


2nd siRNAs: Phase diagram

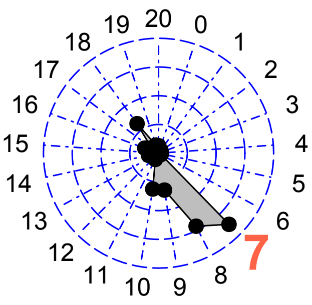


2nd siRNAs: Size distribution

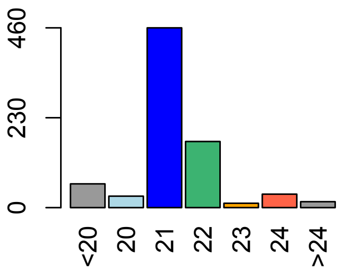


Degradome hits not found for sRNA

**Allenscore:** 5

**Target Interaction:**

```
5'- GGUUACUUAGACCCAGAGUA   AT4G00330.1
     | | | | | | | | | | |   |   | | | | |
3'- CCAAUGAAUCUAGAUCUCAC   Cl_cpe-2017_25056
```

**Target Site:** 1015

**Superfamily:** SupFam_257

**miRNA in ccm:** N/A

**Confirmed Targeting**

| *2nd-siRNA* | *NanoPARE* |
|---|---|
| *ccm* | *ccm* |
| *cpe-2015* | *cpe-2015* |
| *cpe-2017* | |
| *cgr-dp* | *cgr-dp* |
| *cgr-pm* | |
| *cgr-mass* | |
| *cin* | *cin* |

### CRCK2

AT4G00330 - Cl_cpe-2017_25056

Secondary siRNAs

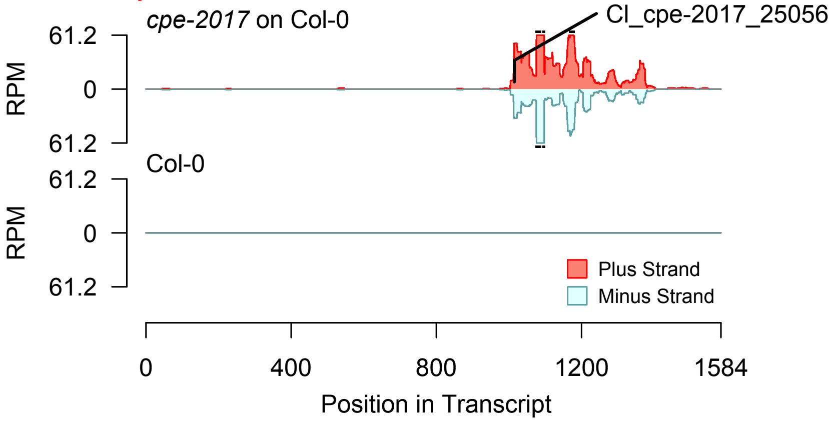


2nd siRNAs: Phase diagram

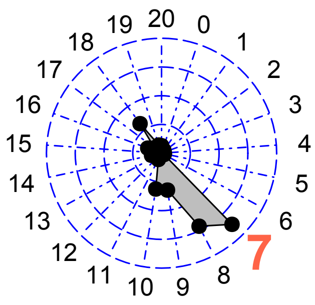


2nd siRNAs: Size distribution

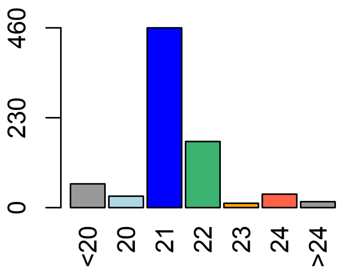


Degradome hits not found for sRNA

**Allenscore:** 5

**Target Interaction:**

```
5'- CAUCU-UCUUCUUCCCUCGAUUCA    AT3G51990.1
     |||||  ||||||||||| ||  ||||
3'- GUAGACAGAAGAAGGG-GC-AAGU    CI_cpe-2015_44
```

**Target Site:** 221

**Superfamily:** SupFam_269

**miRNA in ccm:** Y

**Confirmed Targeting**

| *2nd-siRNA* | *NanoPARE* |
|---|---|
| *ccm* | *ccm* |
| *cpe-2015* | *cpe-2015* |
| *cpe-2017* | |
| *cgr-dp* | *cgr-dp* |
| *cgr-pm* | |
| *cgr-mass* | |
| *cin* | *cin* |

### Protein_kinase_superfamily_protein

AT3G51990 - CI_cpe-2015_44

Secondary siRNAs

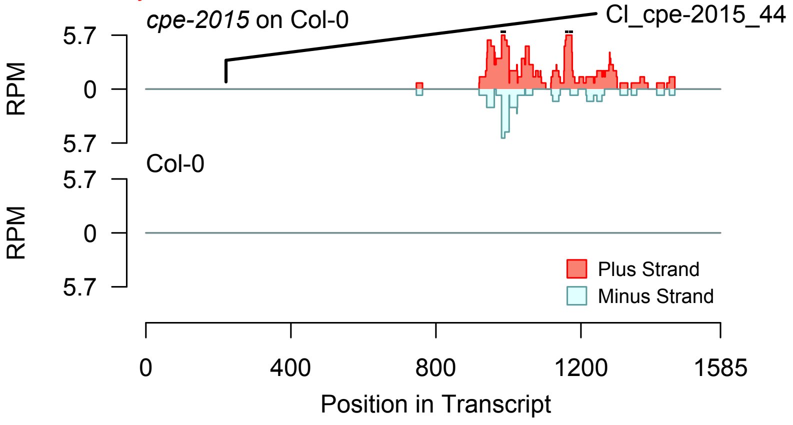


Degradome hits not found for sRNA

2nd siRNAs: Phase diagram

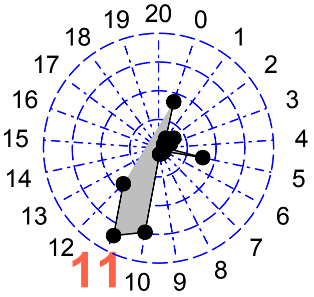


2nd siRNAs: Size distribution

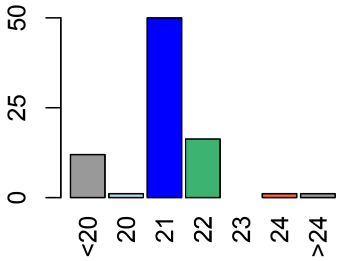

**Allenscore:** 5.5

**Target Interaction:**

```
5'- GGUUACUUAGAUCCAGAUUA    AT3G51990.1
     : | | | | | | | | | | | | | | | |    |
3'- UCAAUGAAUCUAGGUCUCAC    CI_cpe-2015_45087
```

**Target Site:** 921

**Superfamily:** SupFam_257

**miRNA in ccm:** N/A

**Confirmed Targeting**

| *2nd-siRNA* | *NanoPARE* |
|---|---|
| *ccm* | *ccm* |
| *cpe-2015* | *cpe-2015* |
| *cpe-2017* | |
| *cgr-dp* | *cgr-dp* |
| *cgr-pm* | |
| *cgr-mass* | |
| *cin* | *cin* |

### Protein_kinase_superfamily_protein

AT3G51990 - CI_cpe-2015_45087

Secondary siRNAs

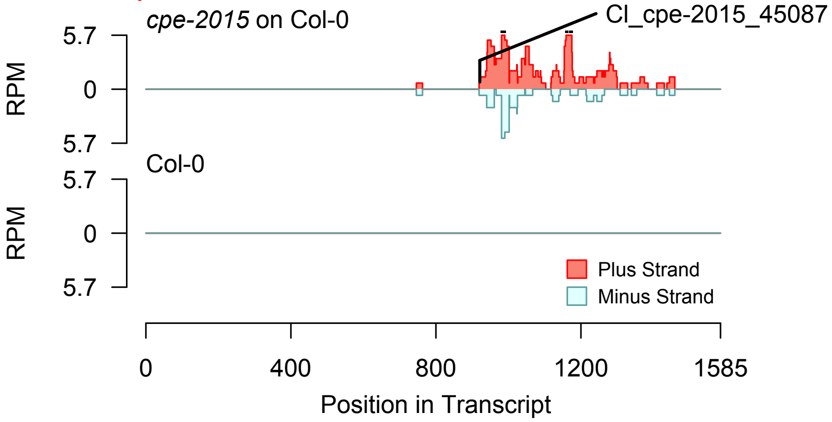


2nd siRNAs: Phase diagram

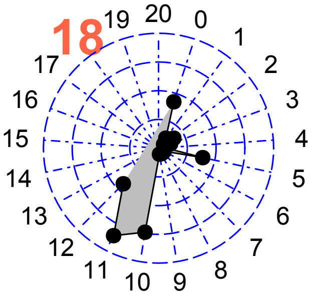


2nd siRNAs: Size distribution

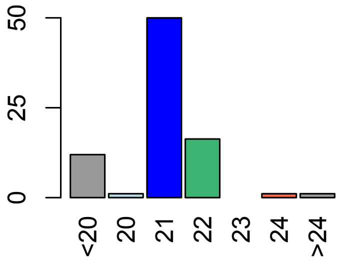


Degradome hits not found for sRNA

**Allenscore:** 5

**Target Interaction:**

```
5'- CAUCU - UCUUCUUCCCUCGAUUCA    AT3G51990.1
     | | | | |   | | | | | | | | | | |   | |   | | | |
3'- GUAGACAGAAGAAGGG - GC - AAGU    Cl_cpe-2017_47
```

**Target Site:** 221

**Superfamily:** SupFam_269

**miRNA in ccm:** Y

**Confirmed Targeting**

| *2nd-siRNA* | *NanoPARE* |
|---|---|
| *ccm* | *ccm* |
| *cpe-2015* | *cpe-2015* |
| *cpe-2017* | |
| *cgr-dp* | *cgr-dp* |
| *cgr-pm* | |
| *cgr-mass* | |
| *cin* | *cin* |

### Protein_kinase_superfamily_protein

AT3G51990 - Cl_cpe-2017_47

Secondary siRNAs

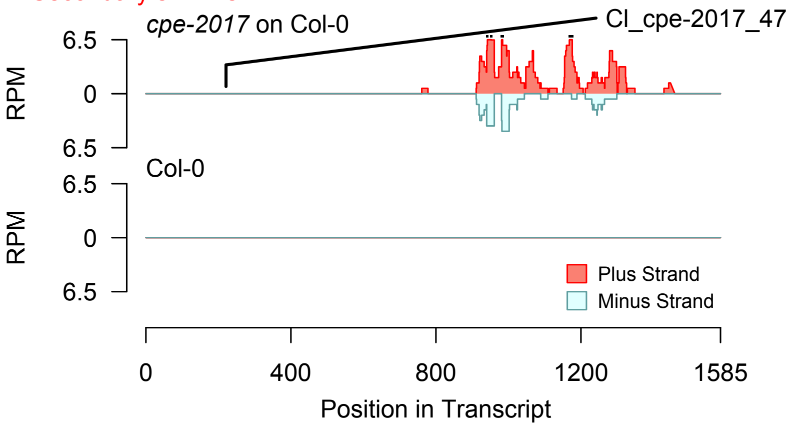


2nd siRNAs: Phase diagram

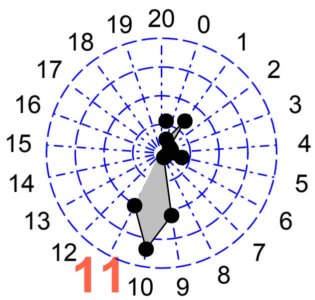


2nd siRNAs: Size distribution

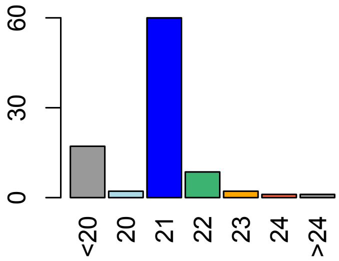


Degradome hits not found for sRNA

**Allenscore:** 3

**Target Interaction:**

```
5'- ACCGGAAGAAGAUGAGAAGCUUA   AT1G63910.1
     |||  ||||||||||||:|||||:|
3'- UGGGCUUCUUCUACUUUUCGAGU   Cl_cgr-dp_279
```

**Target Site:** 216

**Superfamily:** SupFam_37

**miRNA in ccm:** N/A

**Confirmed Targeting**

| *2nd-siRNA* | *NanoPARE* |
|---|---|
| *ccm* | *ccm* |
| *cpe-2015* | *cpe-2015* |
| *cpe-2017* | |
| *cgr-dp* | *cgr-dp* |
| *cgr-pm* | |
| *cgr-mass* | |
| *cin* | *cin* |

### AtMYB103

AT1G63910 - Cl_cgr-dp_279

Secondary siRNAs

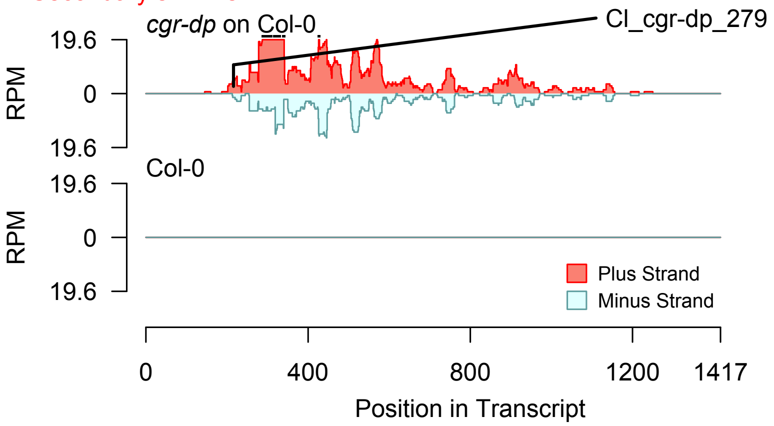


Degradome hits not found for sRNA

2nd siRNAs: Phase diagram

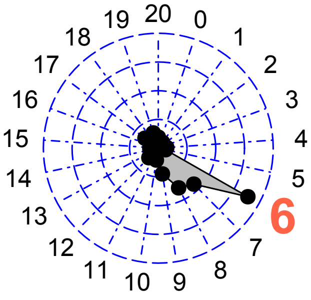


2nd siRNAs: Size distribution

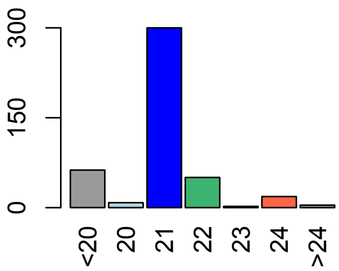

**Allenscore:** 3

**Target Interaction:**

```
5'- GGAAGAGAUGAGAAGCUUA   AT1G63910.1
     | | | | | | | | | | | : | | | | | : |
3'- GCUUCUUCUACUUUUCGAGU  Cl_cgr-dp_37384
```

**Target Site:** 216

**Superfamily:** SupFam_37

**miRNA in ccm:** N/A

**Confirmed Targeting**

| *2nd-siRNA* | *NanoPARE* |
|---|---|
| *ccm* | *ccm* |
| *cpe-2015* | *cpe-2015* |
| *cpe-2017* | |
| *cgr-dp* | *cgr-dp* |
| *cgr-pm* | |
| *cgr-mass* | |
| *cin* | *cin* |

### AtMYB103

AT1G63910 - Cl_cgr-dp_37384

Secondary siRNAs

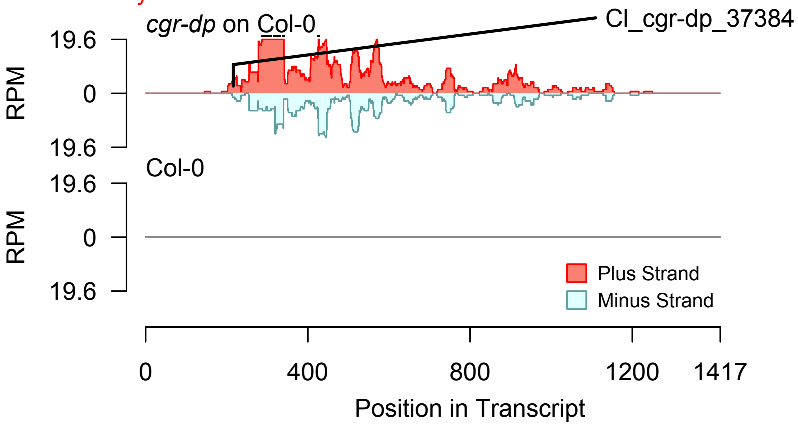


2nd siRNAs: Phase diagram

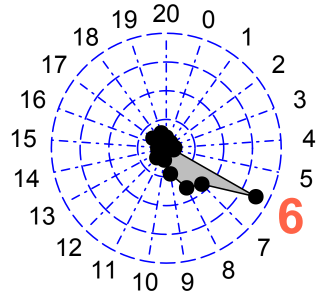


2nd siRNAs: Size distribution

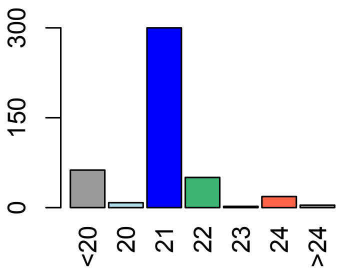


Degradome hits not found for sRNA

**Allenscore:** 2

**Target Interaction:**

```
5'- ACCGGAAGAAGAUGAGAAGCUUA    AT1G63910.1
    ||||||||||||||||:||||||:|
3'- UGGCCUUCUUCUACUUUUCGAGU    Cl_cgr-pm_3376
```

**Target Site:** 216

**Superfamily:** SupFam_37

**miRNA in ccm:** N/A

**Confirmed Targeting**

| 2nd-siRNA | NanoPARE |
|---|---|
| *ccm* | *ccm* |
| *cpe-2015* | *cpe-2015* |
| *cpe-2017* | |
| *cgr-dp* | *cgr-dp* |
| *cgr-pm* | |
| *cgr-mass* | |
| *cin* | *cin* |

### AtMYB103

AT1G63910 - Cl_cgr-pm_3376

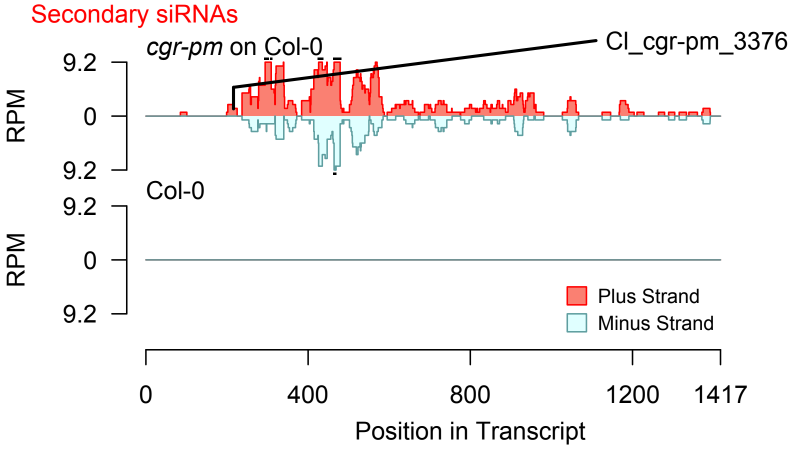


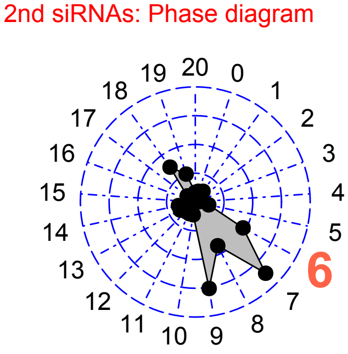


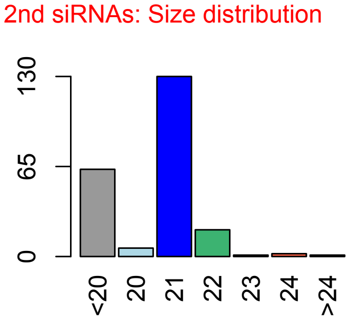


Degradome hits not found for sRNA

**Allenscore:** 4

**Target Interaction:**

```
5'- CCGGAAGAAGAUGAGAAGCUUA   AT1G63910.1
     | |   | | | | | | | | : | | | | | : | | : |
3'- GGACUUCUUCUGCUCUUUGAGU   Cl_cgr-pm_5111
```

**Target Site:** 216

**Superfamily:** SupFam_37

**miRNA in ccm:** N/A

**Confirmed Targeting**

| *2nd-siRNA* | *NanoPARE* |
|---|---|
| *ccm* | *ccm* |
| *cpe-2015* | *cpe-2015* |
| *cpe-2017* | |
| *cgr-dp* | *cgr-dp* |
| *cgr-pm* | |
| *cgr-mass* | |
| *cin* | *cin* |

### AtMYB103

AT1G63910 - Cl_cgr-pm_5111

Secondary siRNAs

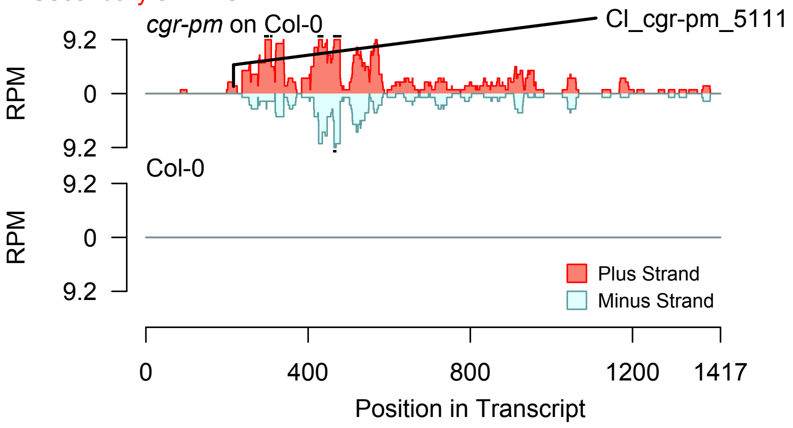


2nd siRNAs: Phase diagram

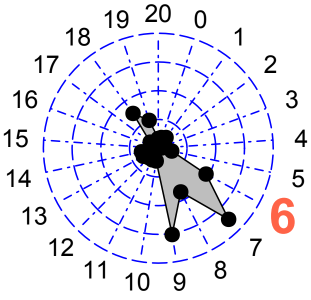


2nd siRNAs: Size distribution

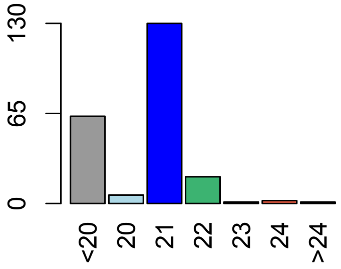


Degradome hits not found for sRNA

**Allenscore:** 4

**Target Interaction:**

```
5'- CCGGAAGAAGAUGAGAAGCUUA   AT1G63910.1
     | |   | | | | | | | | : | | | | | : | | : |
3'- GGACUUCUUCUGCUCUUUGAGU   Cl_cgr-mass_3016
```

**Target Site:** 216

**Superfamily:** SupFam_37

**miRNA in ccm:** N/A

**Confirmed Targeting**

| *2nd-siRNA* | *NanoPARE* |
|---|---|
| *ccm* | *ccm* |
| *cpe-2015* | *cpe-2015* |
| *cpe-2017* | |
| *cgr-dp* | *cgr-dp* |
| *cgr-pm* | |
| *cgr-mass* | |
| *cin* | *cin* |

### AtMYB103

AT1G63910 - Cl_cgr-mass_3016

Secondary siRNAs

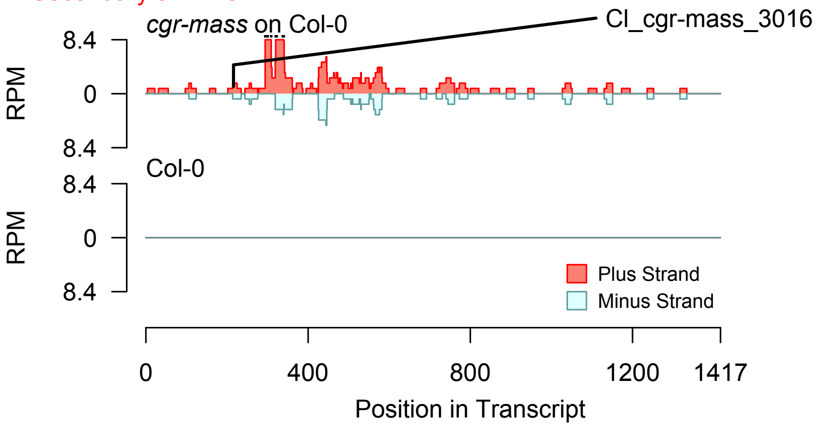


2nd siRNAs: Phase diagram

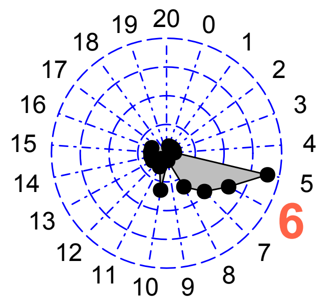


2nd siRNAs: Size distribution

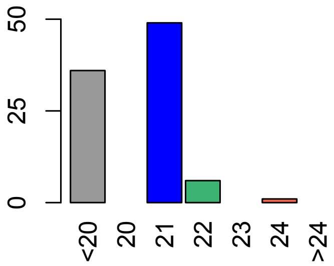


Degradome hits not found for sRNA

**Allenscore:** 2

**Target Interaction:**

```
5'- ACCGGAAGAAGAUGAGAAGCUUA    AT1G63910.1
     | | | | | | | | | | | | | | | | : | | | | | : |
3'- UGGCCUUCUUCUACUUUUCGAGU    Cl_cgr-mass_4415
```

**Target Site:** 216

**Superfamily:** SupFam_37

**miRNA in ccm:** N/A

**Confirmed Targeting**

| *2nd-siRNA* | *NanoPARE* |
|---|---|
| *ccm* | *ccm* |
| *cpe-2015* | *cpe-2015* |
| *cpe-2017* | |
| *cgr-dp* | *cgr-dp* |
| *cgr-pm* | |
| *cgr-mass* | |
| *cin* | *cin* |

### AtMYB103

AT1G63910 - Cl_cgr-mass_4415

Secondary siRNAs

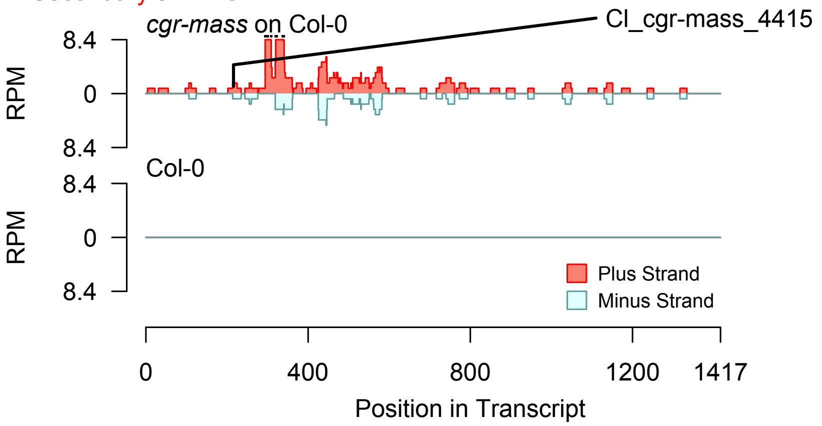


Degradome hits not found for sRNA

2nd siRNAs: Phase diagram

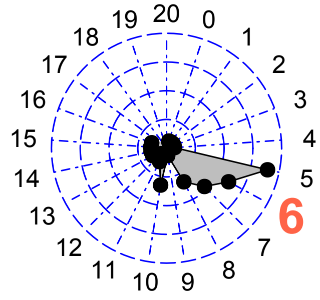


2nd siRNAs: Size distribution

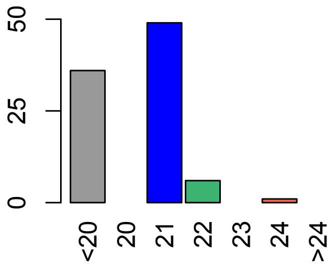

**Allenscore:** 6

**Target Interaction:**

```
5'- UCGUCAGCUCAAUACUUAUGGU   AT1G46264.1
       |  |||||||||||  |||||
3'- CUCUGUCGAGUUAUGUAUACCU   Cl_ccm_19
```

**Target Site:** 457

**Superfamily:** SupFam_116

**miRNA in ccm:** Y

**Confirmed Targeting**

| *2nd-siRNA* | *NanoPARE* |
|---|---|
| *ccm* | *ccm* |
| *cpe-2015* | *cpe-2015* |
| *cpe-2017* | |
| *cgr-dp* | *cgr-dp* |
| *cgr-pm* | |
| *cgr-mass* | |
| *cin* | *cin* |

### HSFB4

AT1G46264 - Cl_ccm_19

Secondary siRNAs

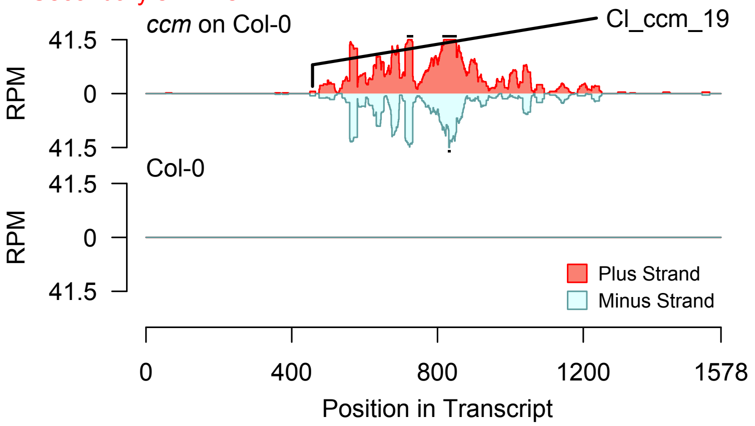


2nd siRNAs: Phase diagram

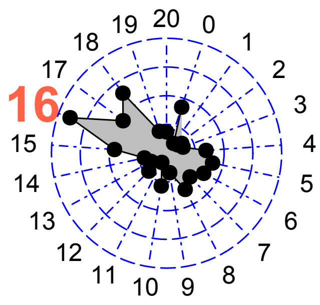


2nd siRNAs: Size distribution

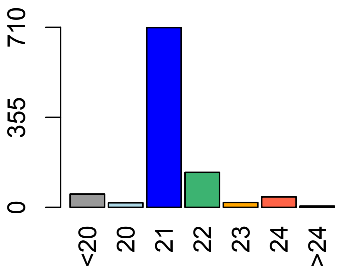


Degradome hits not found for sRNA

**Allenscore:** 6

**Target Interaction:**

```
5'- UCGUCAGCUCAAUACUUAUGGU   AT1G46264.1
       |  | | | | | | | | | | |   | | | | | |
3'- CUCUGUCGAGUUAUGUAUACCU   CI_cpe-2017_72
```

**Target Site:** 457

**Superfamily:** SupFam_116

**miRNA in ccm:** Y

**Confirmed Targeting**

| *2nd-siRNA* | *NanoPARE* |
|---|---|
| *ccm* | *ccm* |
| *cpe-2015* | *cpe-2015* |
| *cpe-2017* | |
| *cgr-dp* | *cgr-dp* |
| *cgr-pm* | |
| *cgr-mass* | |
| *cin* | *cin* |

### HSFB4

AT1G46264 - CI_cpe-2017_72

Secondary siRNAs

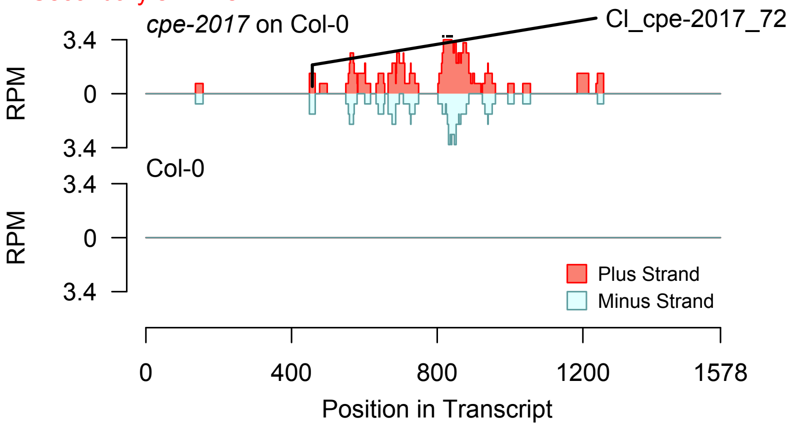


2nd siRNAs: Phase diagram

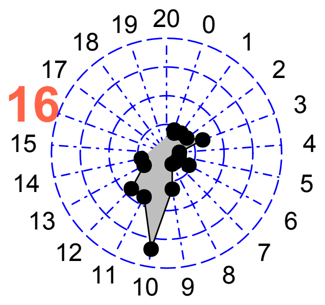


2nd siRNAs: Size distribution

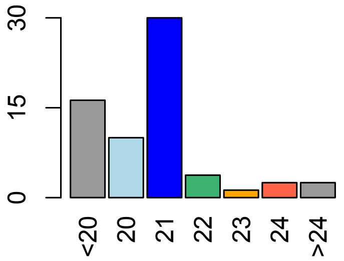


Degradome hits not found for sRNA

**Allenscore:** 5

**Target Interaction:**

```
5'- UCGUCAGCUCAAUACUUAUGGU    AT1G46264.1
     : | :  ||||||||||| |||||
3'- GGUUGUCGAGUUAUGUAUACCU    Cl_cpe-2017_3993
```

**Target Site:** 457

**Superfamily:** SupFam_116

**miRNA in ccm:** Y

**Confirmed Targeting**

| *2nd-siRNA* | *NanoPARE* |
|---|---|
| *ccm* | *ccm* |
| *cpe-2015* | *cpe-2015* |
| *cpe-2017* | |
| *cgr-dp* | *cgr-dp* |
| *cgr-pm* | |
| *cgr-mass* | |
| *cin* | *cin* |

### HSFB4

AT1G46264 - Cl_cpe-2017_3993

Secondary siRNAs

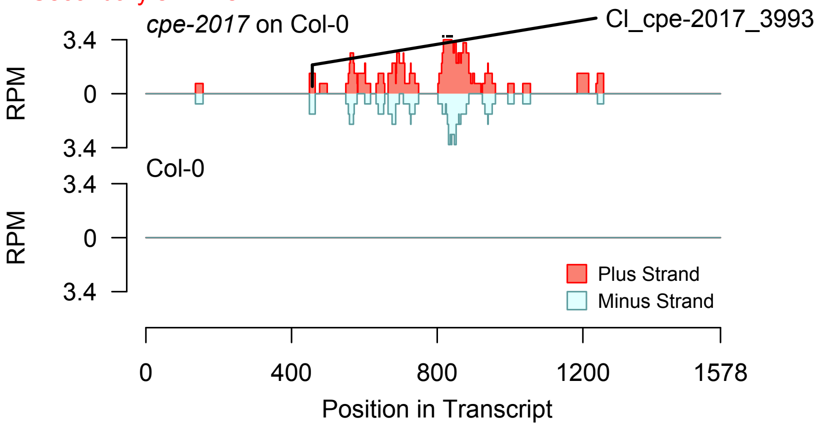


2nd siRNAs: Phase diagram

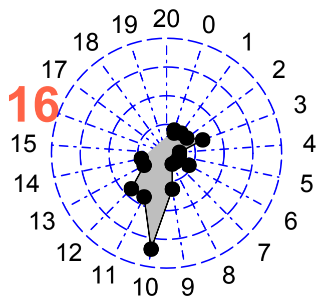


2nd siRNAs: Size distribution

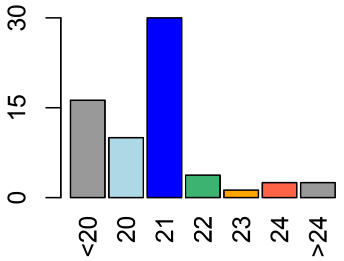


Degradome hits not found for sRNA

**Allenscore:** 4.5

**Target Interaction:**

```
5'- UACGGUUACAUUGCACCAGAAUA    AT5G61480.1
    : |   | | | | | |   : | | | | | | | | | | | |
3'- GUCCCAAUGCGACGUGGUCUUAU    CI_ccm_818
```

**Target Site:** 3083

**Superfamily:** SupFam_124

**miRNA in ccm:** N

**Confirmed Targeting**

| *2nd-siRNA* | *NanoPARE* |
|---|---|
| *ccm* | *ccm* |
| *cpe-2015* | *cpe-2015* |
| *cpe-2017* | |
| *cgr-dp* | *cgr-dp* |
| *cgr-pm* | |
| *cgr-mass* | |
| *cin* | *cin* |

### PXY

AT5G61480 - CI_ccm_818

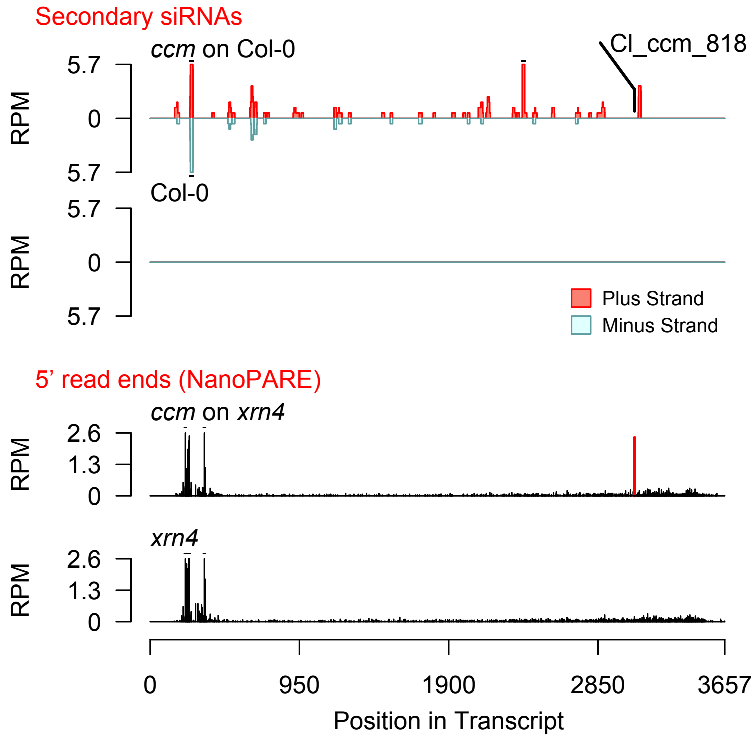


Diff. Exp. secondary siRNA locus not found

**Allenscore:** 3

**Target Interaction:**

```
5'- AUGGAUGGUAAGUUUUAUGUGA    AT5G60570.1
     ||||||||  |||:||||:||||
3'- UACCUACCUUUCGAAAUGCACU    Cl_ccm_670
```

**Target Site:** 1307

**Superfamily:** SupFam_281

**miRNA in ccm:** Y

**Confirmed Targeting**

| *2nd-siRNA* | *NanoPARE* |
|---|---|
| *ccm* | *ccm* |
| *cpe-2015* | *cpe-2015* |
| *cpe-2017* | |
| *cgr-dp* | *cgr-dp* |
| *cgr-pm* | |
| *cgr-mass* | |
| *cin* | *cin* |

### Galactose_oxidase_superfamily_protein

AT5G60570 - Cl_ccm_670

Secondary siRNAs

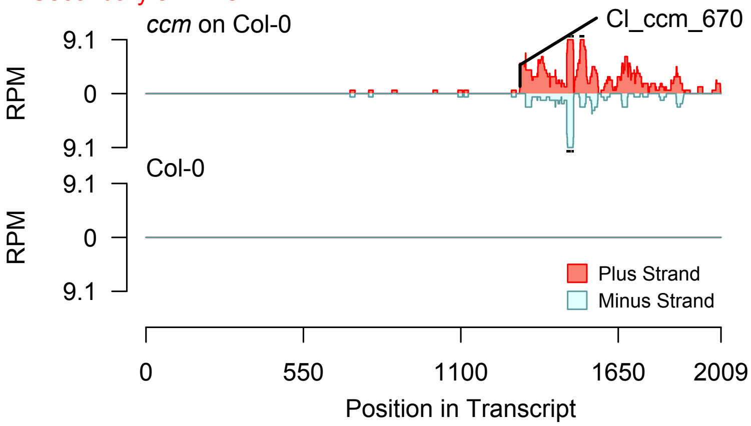


Degradome hits not found for sRNA

2nd siRNAs: Phase diagram

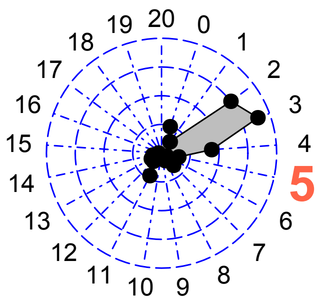


2nd siRNAs: Size distribution

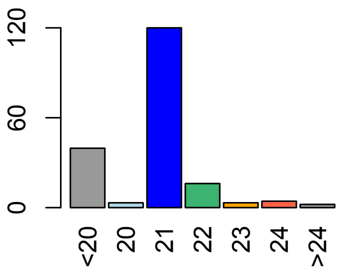

**Allenscore:** 3

**Target Interaction:**

```
5'- AUGGAUGGUAAGUUUUAUGUGA   AT5G60570.1
     | | | | | | | |   | | | : | | | | : | | | |
3'- UACCUACCUUUCGAAAUGCACU   Cl_cpe-2015_316
```

**Target Site:** 1307

**Superfamily:** SupFam_281

**miRNA in ccm:** Y

**Confirmed Targeting**

| *2nd-siRNA* | *NanoPARE* |
|---|---|
| *ccm* | *ccm* |
| *cpe-2015* | *cpe-2015* |
| *cpe-2017* | |
| *cgr-dp* | *cgr-dp* |
| *cgr-pm* | |
| *cgr-mass* | |
| *cin* | *cin* |

### Galactose_oxidase_superfamily_protein

AT5G60570 - Cl_cpe-2015_316

Secondary siRNAs

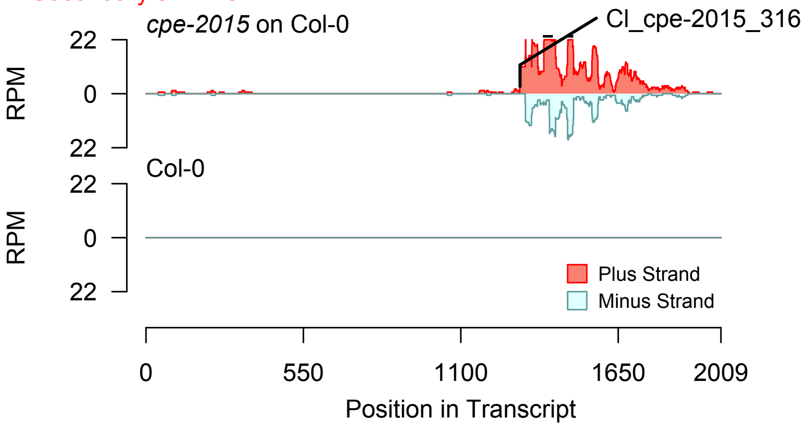


2nd siRNAs: Phase diagram

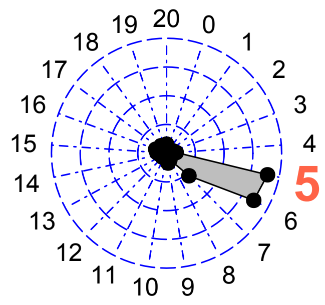


2nd siRNAs: Size distribution

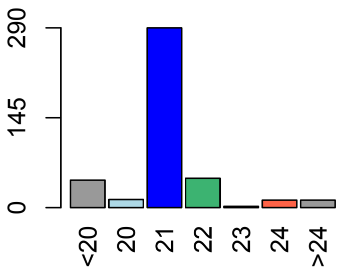


Degradome hits not found for sRNA

**Allenscore:** 3

**Target Interaction:**

```
5'- AUGGAUGGUAAGUUUUAUGUGA    AT5G60570.1
     | | | | | | | |   | | | : | | | | : | | | |
3'- UACCUACCUUUCGAAAUGCACU    Cl_cpe-2017_305
```

**Target Site:** 1307

**Superfamily:** SupFam_281

**miRNA in ccm:** Y

**Confirmed Targeting**

| 2nd-siRNA | NanoPARE |
|---|---|
| *ccm* | *ccm* |
| *cpe-2015* | *cpe-2015* |
| *cpe-2017* | |
| *cgr-dp* | *cgr-dp* |
| *cgr-pm* | |
| *cgr-mass* | |
| *cin* | *cin* |

### Galactose_oxidase_superfamily_protein

AT5G60570 - Cl_cpe-2017_305

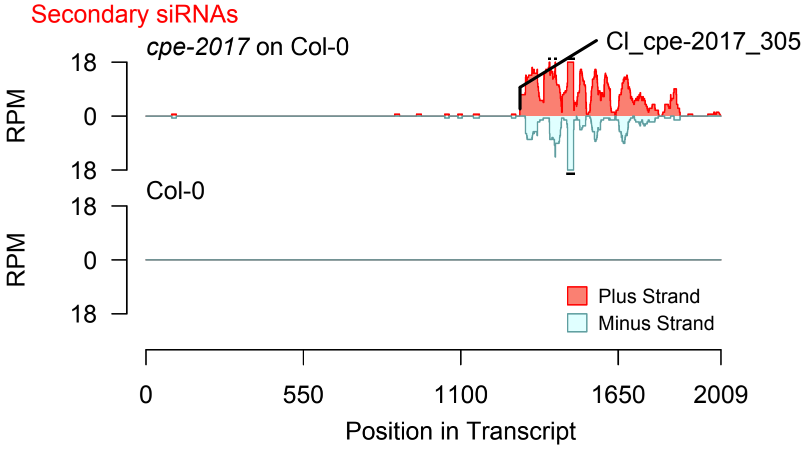


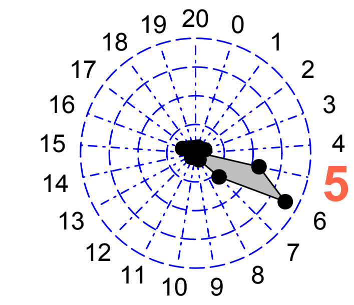


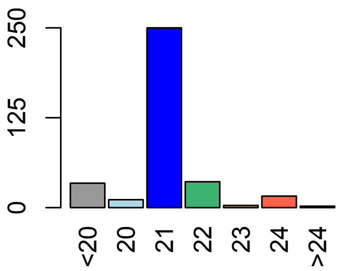


Degradome hits not found for sRNA

**Allenscore:** 2.5

**Target Interaction:**

```
5'- GAGGACUUCUAUAUCUUCACC   AT1G65800.1
     : | | | | | | | | | | | | : | | | | | |
3'- UUCCUGAAGAUAUGGAAGUGU   Cl_cgr-dp_14631
```

**Target Site:** 2196

**Superfamily:** SupFam_49

**miRNA in ccm:** Y

**Confirmed Targeting**

| *2nd-siRNA* | *NanoPARE* |
|---|---|
| *ccm* | *ccm* |
| *cpe-2015* | *cpe-2015* |
| *cpe-2017* | |
| *cgr-dp* | *cgr-dp* |
| *cgr-pm* | |
| *cgr-mass* | |
| *cin* | *cin* |

# RK2

AT1G65800 - Cl_cgr-dp_14631

Secondary siRNAs

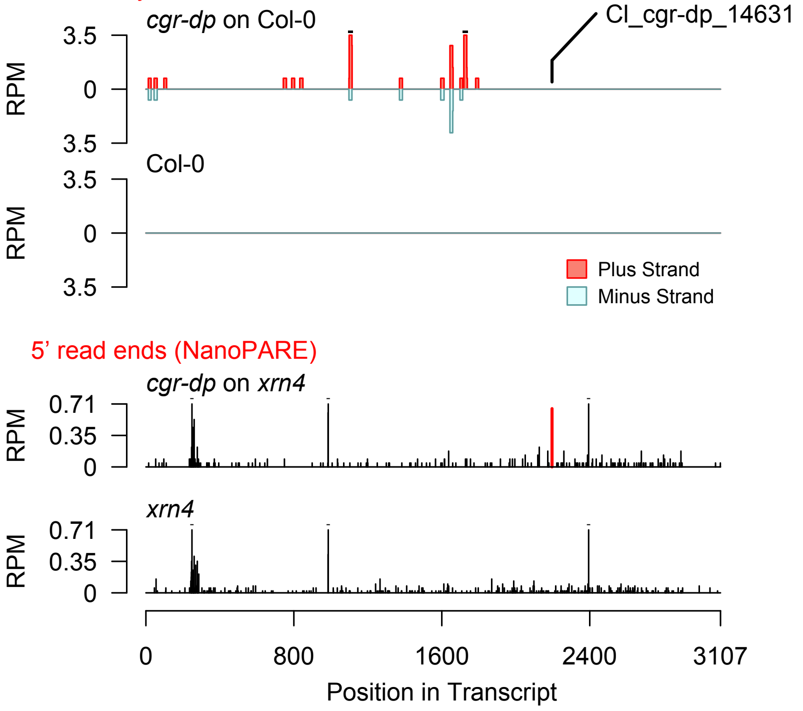


Diff. Exp. secondary siRNA locus not found

**Allenscore:** 2

**Target Interaction:**

```
5'- GGUUAUUUAGAUCCAGAGUA   AT3G53840.1
    : | | | | : | | | | | | | | | | | | |
3'- UCAAUGAAUCUAGGUCUCAC   CI_cpe-2015_45087
```

**Target Site:** 1583

**Superfamily:** SupFam_257

**miRNA in ccm:** N/A

**Confirmed Targeting**

| *2nd-siRNA* | *NanoPARE* |
|---|---|
| *ccm* | *ccm* |
| *cpe-2015* | *cpe-2015* |
| *cpe-2017* | |
| *cgr-dp* | *cgr-dp* |
| *cgr-pm* | |
| *cgr-mass* | |
| *cin* | *cin* |

### Protein_kinase_superfamily_protein

AT3G53840 - CI_cpe-2015_45087

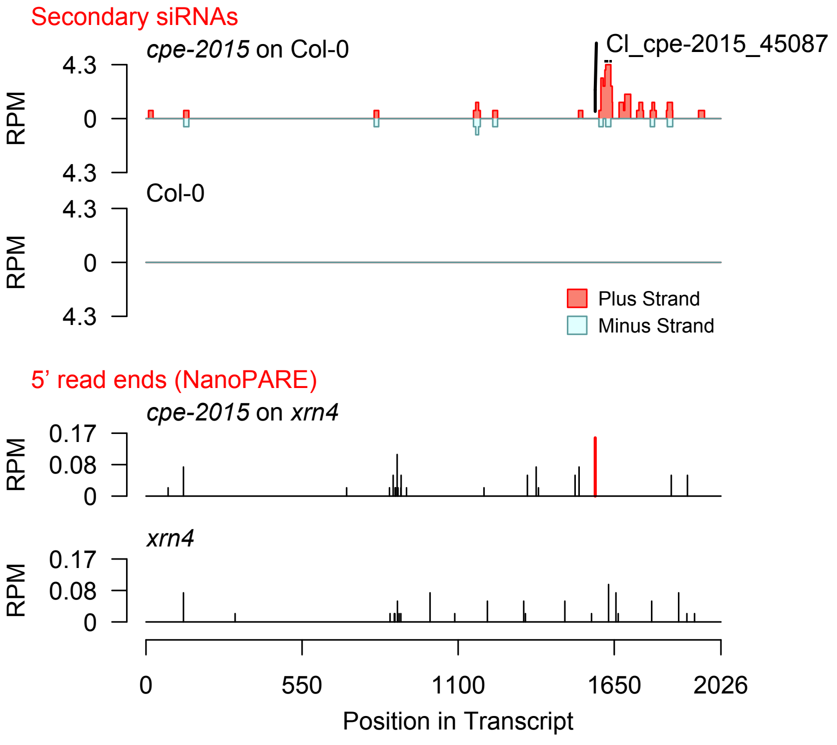


Diff. Exp. secondary siRNA locus not found

**Allenscore:** 1

**Target Interaction:**

```
5'- GCUGAAGCACAUGAACGACUUA    AT1G67310.1
    ||  ||||||||||||||||||||
3'- CG-CUUCGUGUACUUGCUGAAU    CI_ccm_23
```

**Target Site:** 334

**Superfamily:** SupFam_39

**miRNA in ccm:** Y

**Confirmed Targeting**

| *2nd-siRNA* | *NanoPARE* |
|---|---|
| *ccm* | *ccm* |
| *cpe-2015* | *cpe-2015* |
| *cpe-2017* | |
| *cgr-dp* | *cgr-dp* |
| *cgr-pm* | |
| *cgr-mass* | |
| *cin* | *cin* |

### Calmodulin-binding_transcription_activat

AT1G67310 - Cl_ccm_23

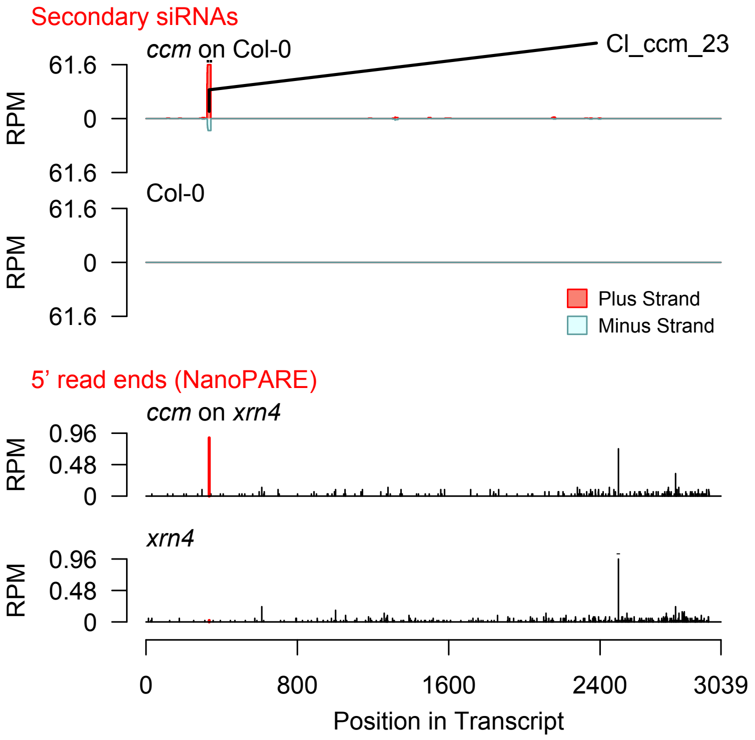


Diff. Exp. secondary siRNA locus not found

**Allenscore:** 4

**Target Interaction:**

```
5'- UGCUGAAGCACAUGAACGACUUA   AT1G67310.1
      ||  ||||||||||||||  |||||
3'- UCG-CUUCGUGUACUUGGUGAAU   CI_cgr-dp_47
```

**Target Site:** 334

**Superfamily:** SupFam_39

**miRNA in ccm:** Y

**Confirmed Targeting**

| *2nd-siRNA* | *NanoPARE* |
|---|---|
| *ccm* | *ccm* |
| *cpe-2015* | *cpe-2015* |
| *cpe-2017* | |
| *cgr-dp* | *cgr-dp* |
| *cgr-pm* | |
| *cgr-mass* | |
| *cin* | *cin* |

### Calmodulin-binding_transcription_activat

AT1G67310 - CI_cgr-dp_47

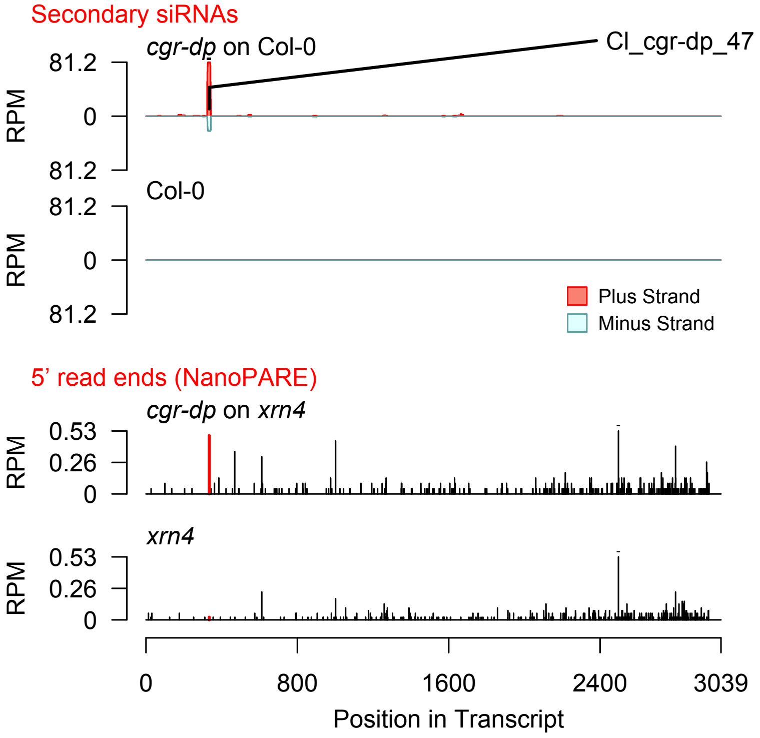


Diff. Exp. secondary siRNA locus not found

**Allenscore:** 3.5

**Target Interaction:**

```
5'- AUGGAUGUUAUCUGCCAAGCA   AT5G11200.1
     | | | | | | | | : | | | : : | | | | | | |
3'- UACCUACAGUAGGUGGUUCGA   Cl_cin_42282
```

**Target Site:** 359

**Superfamily:** SupFam_1585

**miRNA in ccm:** N/A

**Confirmed Targeting**

| *2nd-siRNA* | *NanoPARE* |
|---|---|
| *ccm* | *ccm* |
| *cpe-2015* | *cpe-2015* |
| *cpe-2017* | |
| *cgr-dp* | *cgr-dp* |
| *cgr-pm* | |
| *cgr-mass* | |
| *cin* | *cin* |

### UAP56b

AT5G11200 - Cl_cin_42282

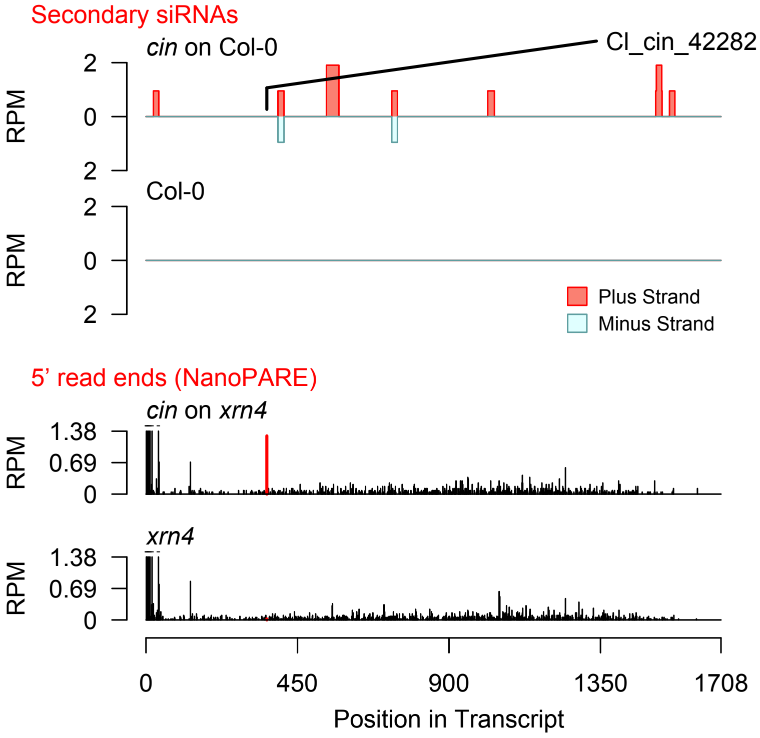


Diff. Exp. secondary siRNA locus not found

**Allenscore:** 1

**Target Interaction:**

```
5'- CUAGUCUACGAGUUUAUGCAAA   AT2G39660.1
     | | | | | | | | | | | : | | | | | | | | | |
3'- GAUCAGAUGCUUAAAUACGUUU   CI_ccm_7
```

**Target Site:** 870

**Superfamily:** SupFam_1

**miRNA in ccm:** Y

**Confirmed Targeting**

| *2nd-siRNA* | *NanoPARE* |
|---|---|
| *ccm* | *ccm* |
| *cpe-2015* | *cpe-2015* |
| *cpe-2017* | |
| *cgr-dp* | *cgr-dp* |
| *cgr-pm* | |
| *cgr-mass* | |
| *cin* | *cin* |

### BIK1

AT2G39660 - CI_ccm_7

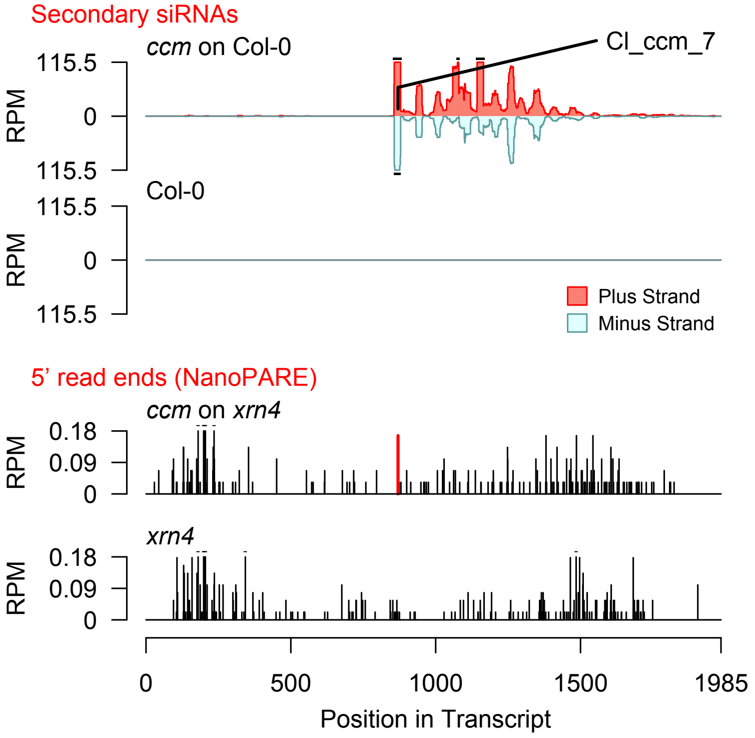


2nd siRNAs: Phase diagram

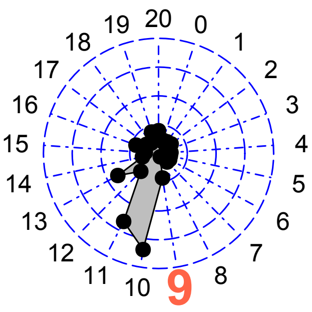


2nd siRNAs: Size distribution

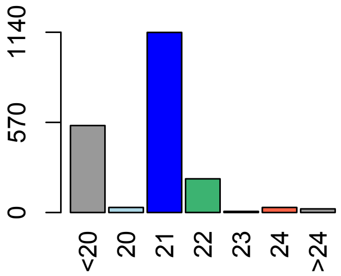

**Allenscore:** 2

**Target Interaction:**

```
5'- AGUCUACGAGUUUAUGCAAA   AT2G39660.1
      ||||||||:||||||||||
3'- CCAGAUGCUUAAAUACGUUU   CI_ccm_7146
```

**Target Site:** 870

**Superfamily:** SupFam_1

**miRNA in ccm:** Y

**Confirmed Targeting**

| *2nd-siRNA* | *NanoPARE* |
|---|---|
| *ccm* | *ccm* |
| *cpe-2015* | *cpe-2015* |
| *cpe-2017* | |
| *cgr-dp* | *cgr-dp* |
| *cgr-pm* | |
| *cgr-mass* | |
| *cin* | *cin* |

### BIK1

AT2G39660 - CI_ccm_7146

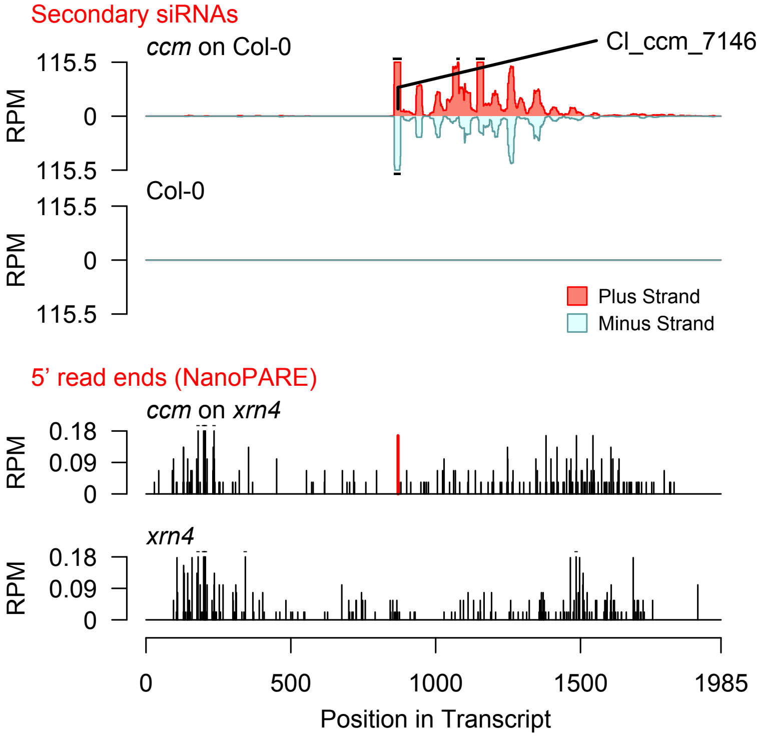


2nd siRNAs: Phase diagram

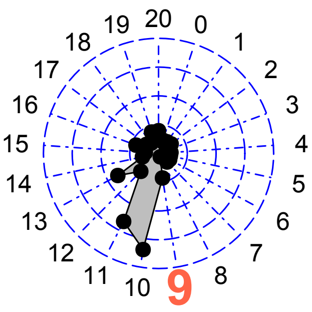


2nd siRNAs: Size distribution

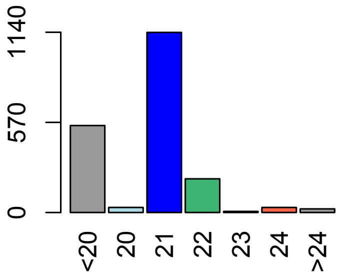

**Allenscore:** 2

**Target Interaction:**

```
5'- UAUGGGUACGCCGCGCCUGAGUA   AT2G39660.1
      ||||  |||||||||||||||||
3'- UUACCGAUGCGGCGCGGACUCAU   CI_ccm_117
```

**Target Site:** 1156

**Superfamily:** SupFam_190

**miRNA in ccm:** Y

**Confirmed Targeting**

| *2nd-siRNA* | *NanoPARE* |
|---|---|
| *ccm* | *ccm* |
| *cpe-2015* | *cpe-2015* |
| *cpe-2017* | |
| *cgr-dp* | *cgr-dp* |
| *cgr-pm* | |
| *cgr-mass* | |
| *cin* | *cin* |

### BIK1

AT2G39660 - CI_ccm_117

Secondary siRNAs

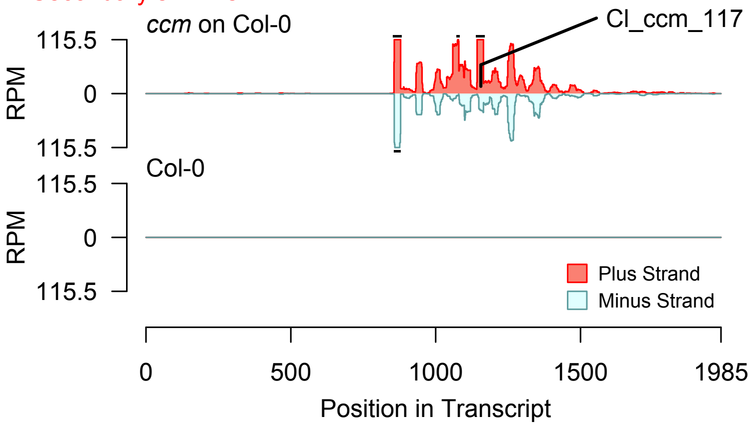


2nd siRNAs: Phase diagram

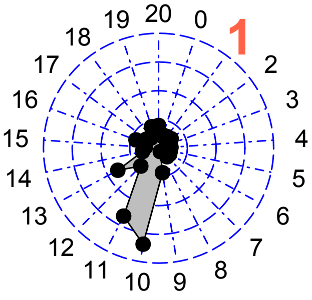


2nd siRNAs: Size distribution

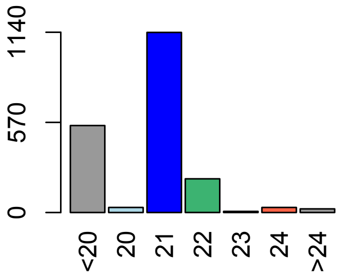


Degradome hits not found for sRNA

**Allenscore:** 1

**Target Interaction:**

```
5'- GGGUACGCCGCGCCUGAGUA   AT2G39660.1
     | |   | | | | | | | | | | | | | | | | |
3'- CCGAUGCGGCGCGGACUCAU   CI_ccm_23740
```

**Target Site:** 1156

**Superfamily:** SupFam_190

**miRNA in ccm:** Y

**Confirmed Targeting**

| 2nd-siRNA | NanoPARE |
|---|---|
| *ccm* | *ccm* |
| *cpe-2015* | *cpe-2015* |
| *cpe-2017* | |
| *cgr-dp* | *cgr-dp* |
| *cgr-pm* | |
| *cgr-mass* | |
| *cin* | *cin* |

### BIK1

AT2G39660 - CI_ccm_23740

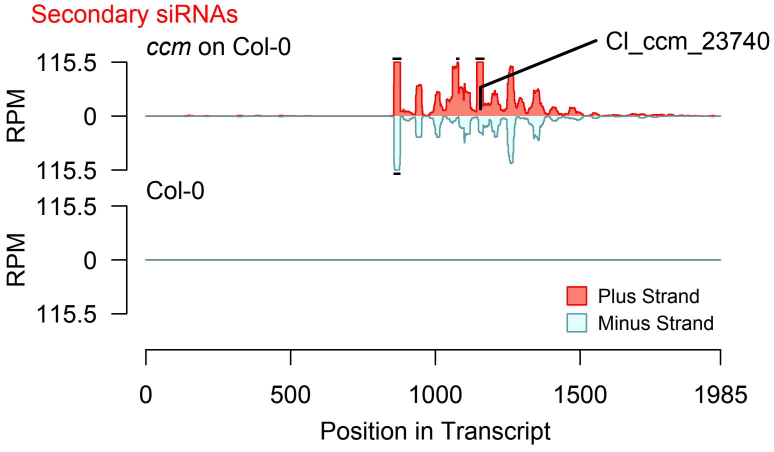


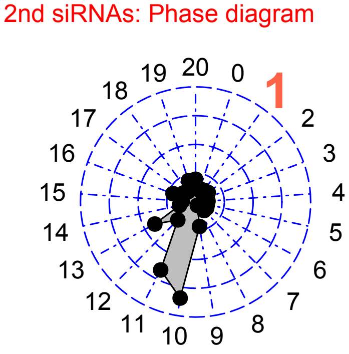


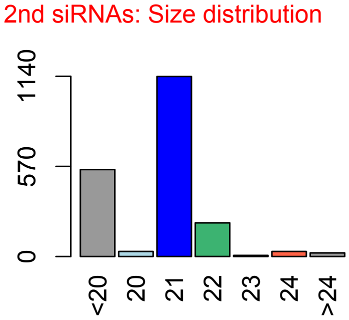


Degradome hits not found for sRNA

**Allenscore:** 1

**Target Interaction:**

```
5'- CUAGUCUACGAGUUUAUGCAAA    AT2G39660.1
    |||||||||||:||||||||||
3'- GAUCAGAUGCUUAAAUACGUUU    CI_cpe-2015_5
```

**Target Site:** 870

**Superfamily:** SupFam_1

**miRNA in ccm:** Y

**Confirmed Targeting**

| *2nd-siRNA* | *NanoPARE* |
|---|---|
| *ccm* | *ccm* |
| *cpe-2015* | *cpe-2015* |
| *cpe-2017* | |
| *cgr-dp* | *cgr-dp* |
| *cgr-pm* | |
| *cgr-mass* | |
| *cin* | *cin* |

### BIK1

AT2G39660 - CI_cpe-2015_5

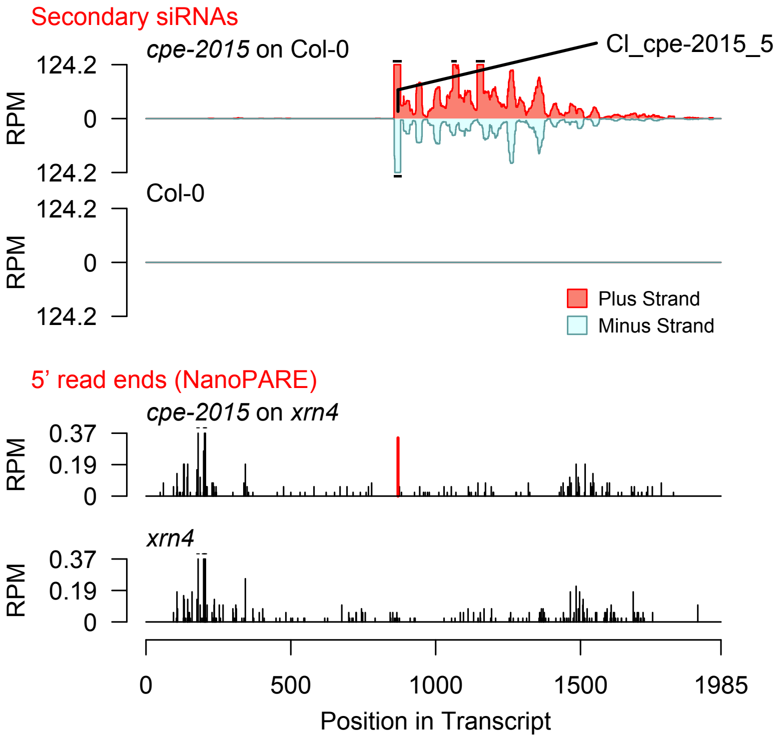


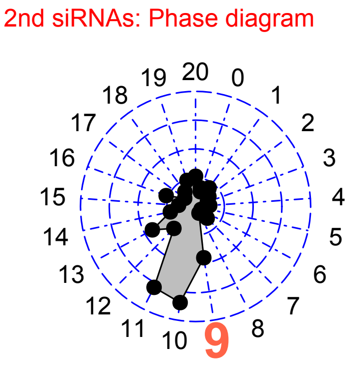


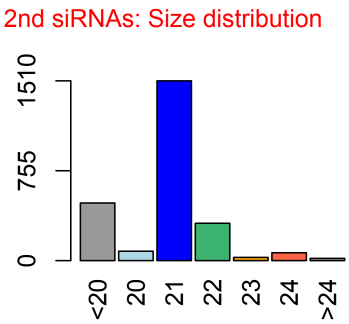

**Allenscore:** 2

**Target Interaction:**

```
5'- AGUCUACGAGUUUAUGCAAA   AT2G39660.1
     ||||||||:||||||||||
3'- CCAGAUGCUUAAAUACGUUU   Cl_cpe-2015_4852
```

**Target Site:** 870

**Superfamily:** SupFam_1

**miRNA in ccm:** Y

**Confirmed Targeting**

| *2nd-siRNA* | *NanoPARE* |
|---|---|
| *ccm* | *ccm* |
| *cpe-2015* | *cpe-2015* |
| *cpe-2017* | |
| *cgr-dp* | *cgr-dp* |
| *cgr-pm* | |
| *cgr-mass* | |
| *cin* | *cin* |

### BIK1

AT2G39660 - Cl_cpe-2015_4852

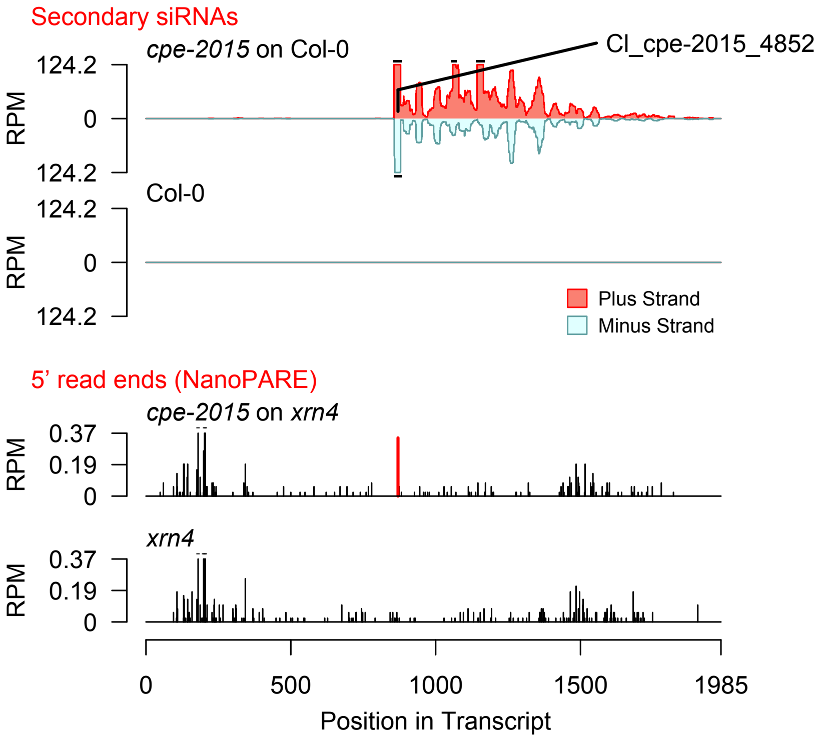


2nd siRNAs: Phase diagram

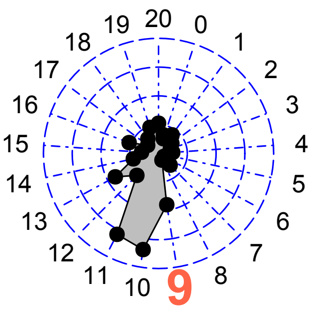


2nd siRNAs: Size distribution

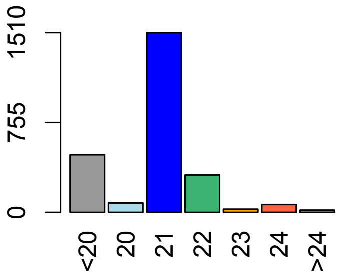

**Allenscore:** 2

**Target Interaction:**

```
5'- UAUGGGUACGCCGCGCCUGAGUA    AT2G39660.1
       ||||  |||||||||||||||||
3'- UUACCGAUGCGGCGCGGACUCAU    CI_cpe-2015_953
```

**Target Site:** 1156

**Superfamily:** SupFam_190

**miRNA in ccm:** Y

**Confirmed Targeting**

| *2nd-siRNA* | *NanoPARE* |
|---|---|
| *ccm* | *ccm* |
| *cpe-2015* | *cpe-2015* |
| *cpe-2017* | |
| *cgr-dp* | *cgr-dp* |
| *cgr-pm* | |
| *cgr-mass* | |
| *cin* | *cin* |

### BIK1

AT2G39660 - CI_cpe-2015_953

Secondary siRNAs

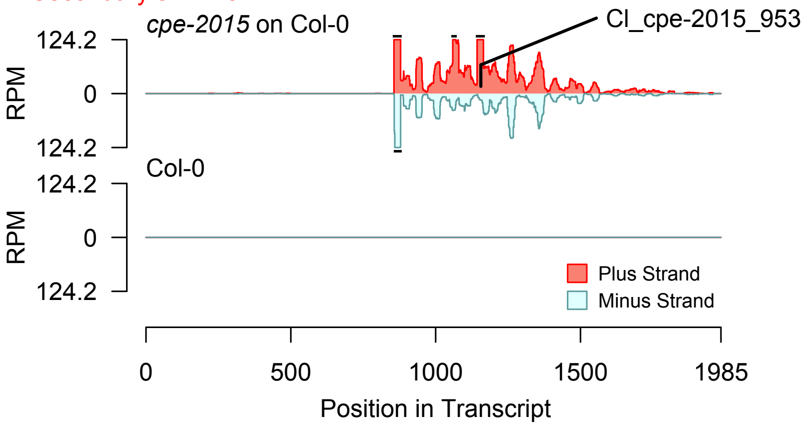


2nd siRNAs: Phase diagram

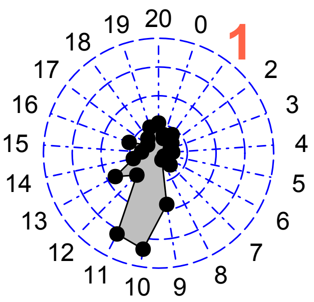


2nd siRNAs: Size distribution

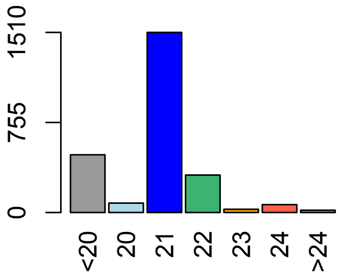


Degradome hits not found for sRNA

**Allenscore:** 1

**Target Interaction:**

```
5'- CUAGUCUACGAGUUUAUGCAAA   AT2G39660.1
    |||||||||||:|||||||||
3'- GAUCAGAUGCUUAAAUACGUUU   Cl_cpe-2017_9
```

**Target Site:** 870

**Superfamily:** SupFam_1

**miRNA in ccm:** Y

**Confirmed Targeting**

| *2nd-siRNA* | *NanoPARE* |
|---|---|
| *ccm* | *ccm* |
| *cpe-2015* | *cpe-2015* |
| *cpe-2017* | |
| *cgr-dp* | *cgr-dp* |
| *cgr-pm* | |
| *cgr-mass* | |
| *cin* | *cin* |

### BIK1

AT2G39660 - Cl_cpe-2017_9

Secondary siRNAs

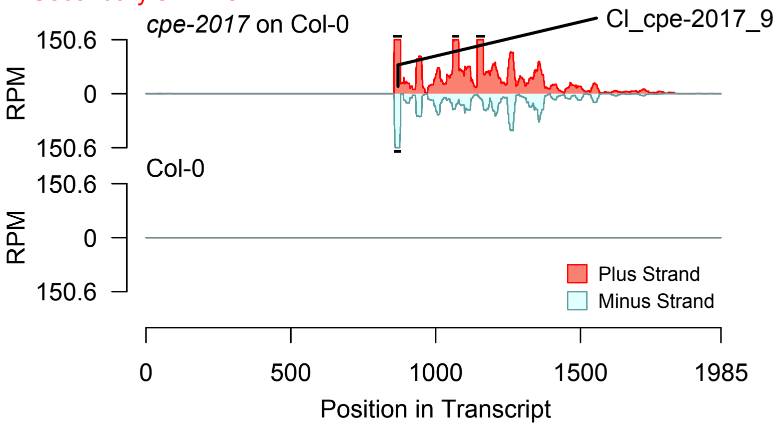


Degradome hits not found for sRNA

2nd siRNAs: Phase diagram

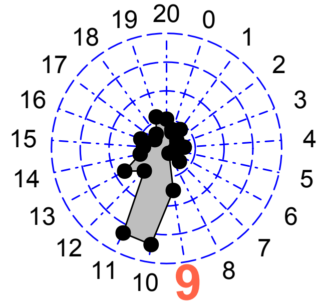


2nd siRNAs: Size distribution

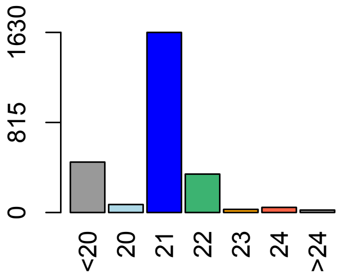

**Allenscore:** 2

**Target Interaction:**

```
5'- AGUCUACGAGUUUAUGCAAA   AT2G39660.1
      |||||||||:||||||||||
3'- CCAGAUGCUUAAAUACGUUU   CI_cpe-2017_5671
```

**Target Site:** 870

**Superfamily:** SupFam_1

**miRNA in ccm:** Y

**Confirmed Targeting**

| *2nd-siRNA* | *NanoPARE* |
|---|---|
| *ccm* | *ccm* |
| *cpe-2015* | *cpe-2015* |
| *cpe-2017* | |
| *cgr-dp* | *cgr-dp* |
| *cgr-pm* | |
| *cgr-mass* | |
| *cin* | *cin* |

### BIK1

AT2G39660 - CI_cpe-2017_5671

Secondary siRNAs

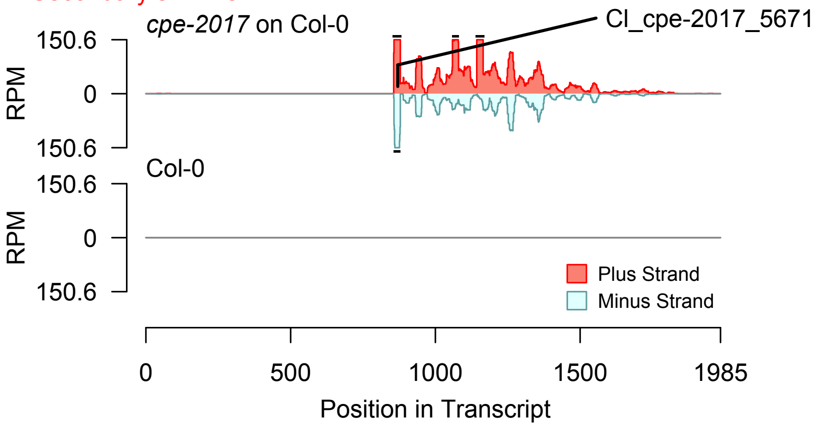


Degradome hits not found for sRNA

2nd siRNAs: Phase diagram

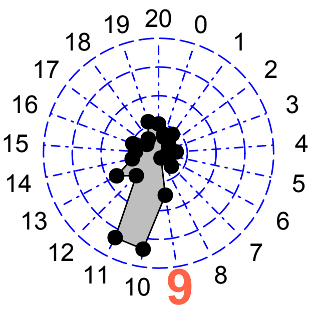


2nd siRNAs: Size distribution

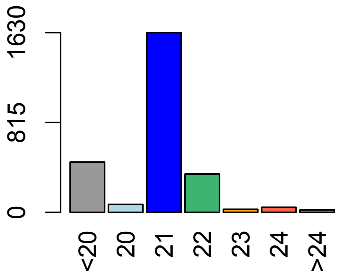

**Allenscore:** 2

**Target Interaction:**

```
5'- UAUGGGUACGCCGCGCCUGAGUA   AT2G39660.1
      ||||  ||||||||||||||||
3'- UUACCGAUGCGGCGCGGACUCAU   CI_cpe-2017_695
```

**Target Site:** 1156

**Superfamily:** SupFam_190

**miRNA in ccm:** Y

**Confirmed Targeting**

| *2nd-siRNA* | *NanoPARE* |
|---|---|
| *ccm* | *ccm* |
| *cpe-2015* | *cpe-2015* |
| *cpe-2017* | |
| *cgr-dp* | *cgr-dp* |
| *cgr-pm* | |
| *cgr-mass* | |
| *cin* | *cin* |

### BIK1

AT2G39660 - CI_cpe-2017_695

Secondary siRNAs

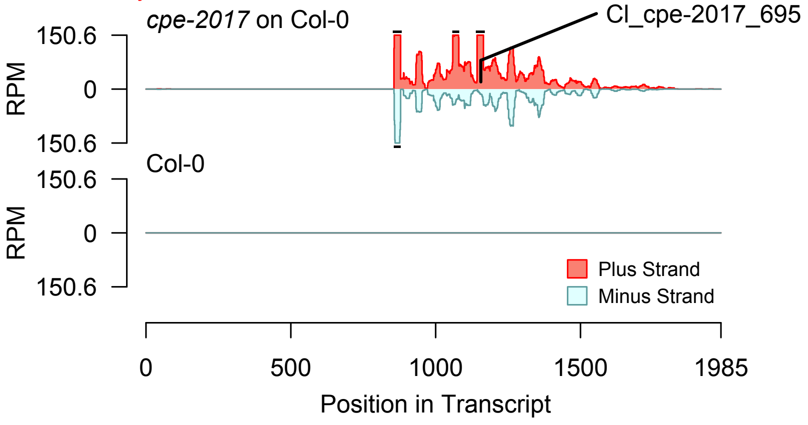


2nd siRNAs: Phase diagram

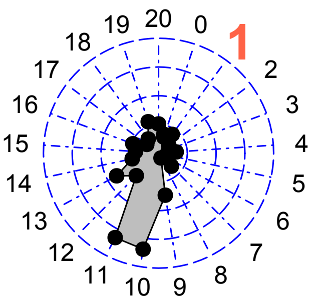


2nd siRNAs: Size distribution

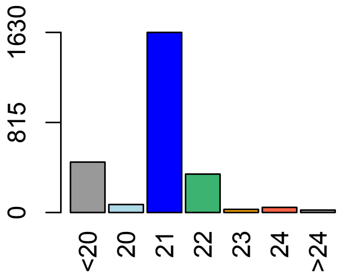


Degradome hits not found for sRNA

**Allenscore:** 4.5

**Target Interaction:**

```
5'- UUGUUUAUGAAAGGGAAUACA   AT1G10900.1
       | : | | | | | | | | | | : | | | | | |
3'- UUCGAAUACUUUCUCUUAUGA   Cl_cgr-dp_539
```

**Target Site:** 1185

**Superfamily:** SupFam_370

**miRNA in ccm:** N/A

**Confirmed Targeting**

| *2nd-siRNA* | *NanoPARE* |
|---|---|
| *ccm* | *ccm* |
| *cpe-2015* | *cpe-2015* |
| *cpe-2017* | |
| *cgr-dp* | *cgr-dp* |
| *cgr-pm* | |
| *cgr-mass* | |
| *cin* | *cin* |

### Phosphatidylinositol-4-phosphate_5-kinas

AT1G10900 - Cl_cgr-dp_539

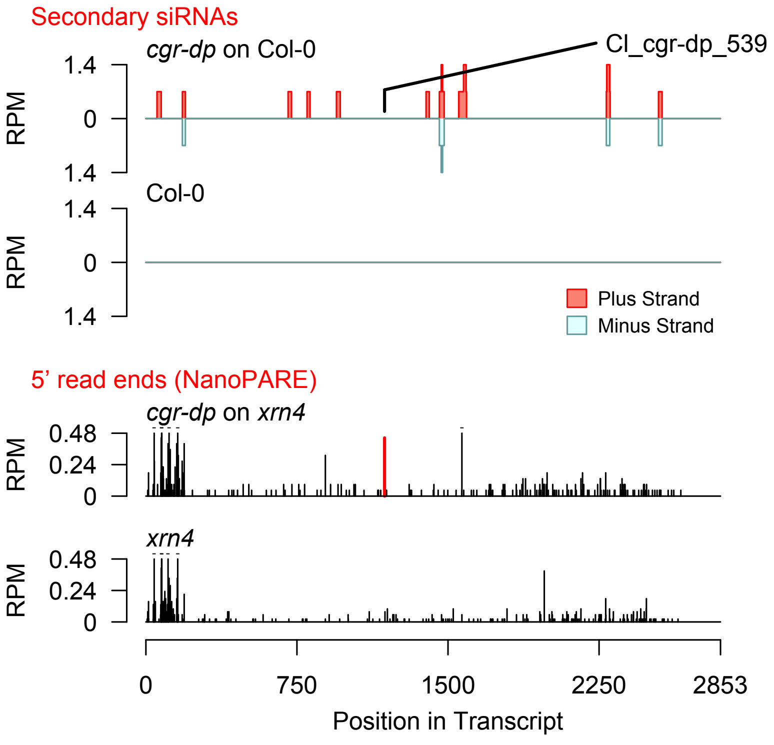


Diff. Exp. secondary siRNA locus not found

**Allenscore:** 4.5

**Target Interaction:**

```
5'- AGUGAGGAACUUAGAGGACUCG   AT3G09830.1
     | :   : | | : | | | | | | | | | | | | | | :
3'- UUCUUCUUUGAAUCUCCUGAGU   Cl_cgr-dp_42
```

**Target Site:** 689

**Superfamily:** SupFam_81

**miRNA in ccm:** N/A

**Confirmed Targeting**

| *2nd-siRNA* | *NanoPARE* |
|---|---|
| *ccm* | *ccm* |
| *cpe-2015* | *cpe-2015* |
| *cpe-2017* | |
| *cgr-dp* | *cgr-dp* |
| *cgr-pm* | |
| *cgr-mass* | |
| *cin* | *cin* |

### PCRK1

AT3G09830 - Cl_cgr-dp_42

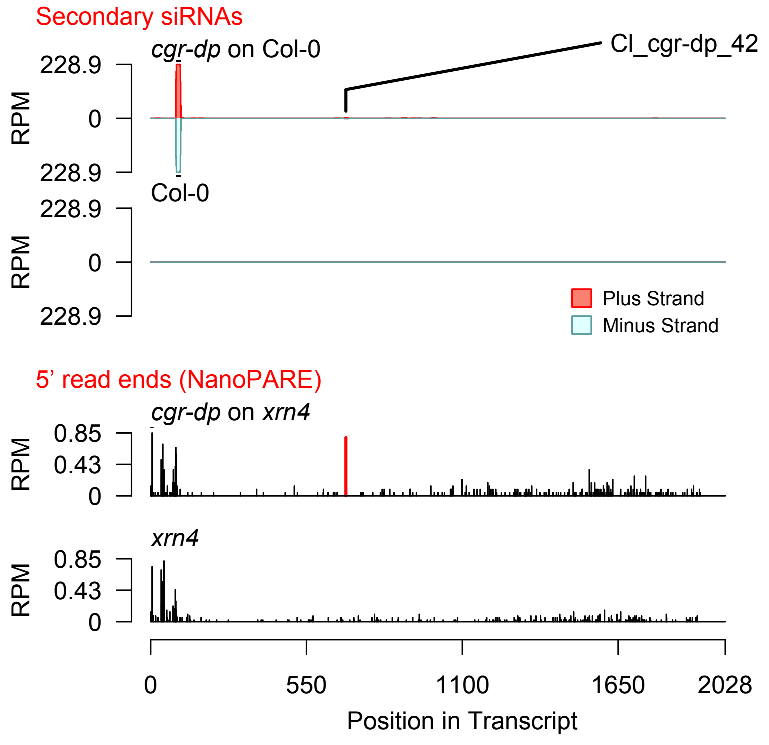


Diff. Exp. secondary siRNA locus not found

**Allenscore:** 4

**Target Interaction:**

```
5'- UGAGCUACUUACACCAAAAUA   AT4G38470.1
     | : | |   | | | :   | | | | | | | | | | |
3'- AUUCUAUGGUUGUGGUUUUAU   Cl_cin_14840
```

**Target Site:** 1743

**Superfamily:** SupFam_189

**miRNA in ccm:** N/A

**Confirmed Targeting**

| *2nd-siRNA* | *NanoPARE* |
|---|---|
| *ccm* | *ccm* |
| *cpe-2015* | *cpe-2015* |
| *cpe-2017* | |
| *cgr-dp* | *cgr-dp* |
| *cgr-pm* | |
| *cgr-mass* | |
| *cin* | *cin* |

### STY46

AT4G38470 - Cl_cin_14840

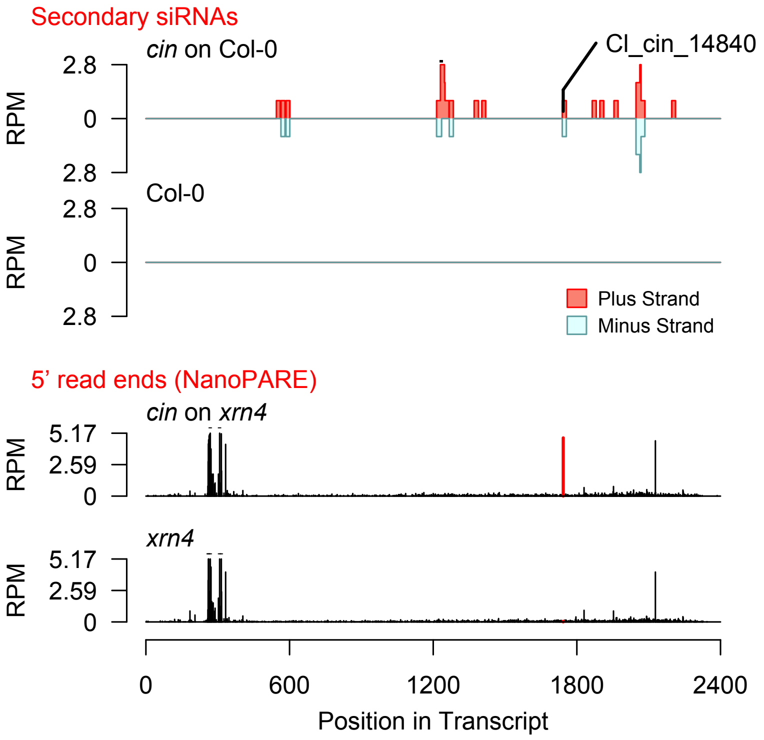


Diff. Exp. secondary siRNA locus not found

**Allenscore:** 3.5

**Target Interaction:**

```
5'- UGAGCUACUUACACCAAAAUA   AT4G38470.1
     | | | | | | | | : |   | | | | | | | | | |
3'- CCUCGAUGGACGUGGUUUUAU   Cl_cin_176
```

**Target Site:** 1743

**Superfamily:** SupFam_189

**miRNA in ccm:** N/A

**Confirmed Targeting**

| *2nd-siRNA* | *NanoPARE* |
|---|---|
| *ccm* | *ccm* |
| *cpe-2015* | *cpe-2015* |
| *cpe-2017* | |
| *cgr-dp* | *cgr-dp* |
| *cgr-pm* | |
| *cgr-mass* | |
| *cin* | *cin* |

### STY46

AT4G38470 - Cl_cin_176

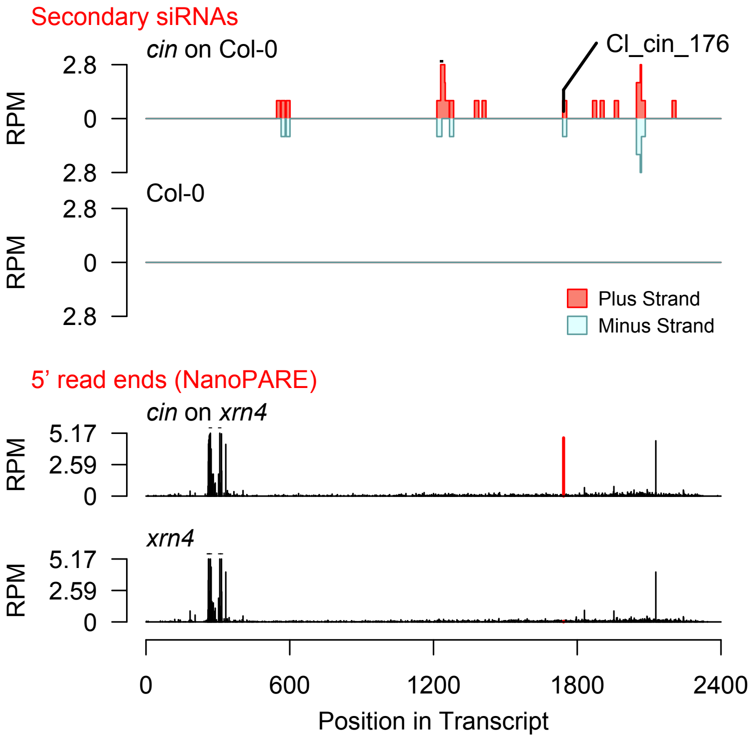


Diff. Exp. secondary siRNA locus not found

**Allenscore:** 4.5

**Target Interaction:**

```
5'- UGAGCUACUUACACCAAAAUA   AT4G38470.1
      | | | | | | | : | | | |   | | | | | : |
3'- UCUCGAUGGAUGUAGUUUUGU   CI_cin_21889
```

**Target Site:** 1743

**Superfamily:** SupFam_189

**miRNA in ccm:** N/A

**Confirmed Targeting**

| *2nd-siRNA* | *NanoPARE* |
| --- | --- |
| *ccm* | *ccm* |
| *cpe-2015* | *cpe-2015* |
| *cpe-2017* | |
| *cgr-dp* | *cgr-dp* |
| *cgr-pm* | |
| *cgr-mass* | |
| *cin* | *cin* |

### STY46

AT4G38470 - CI_cin_21889

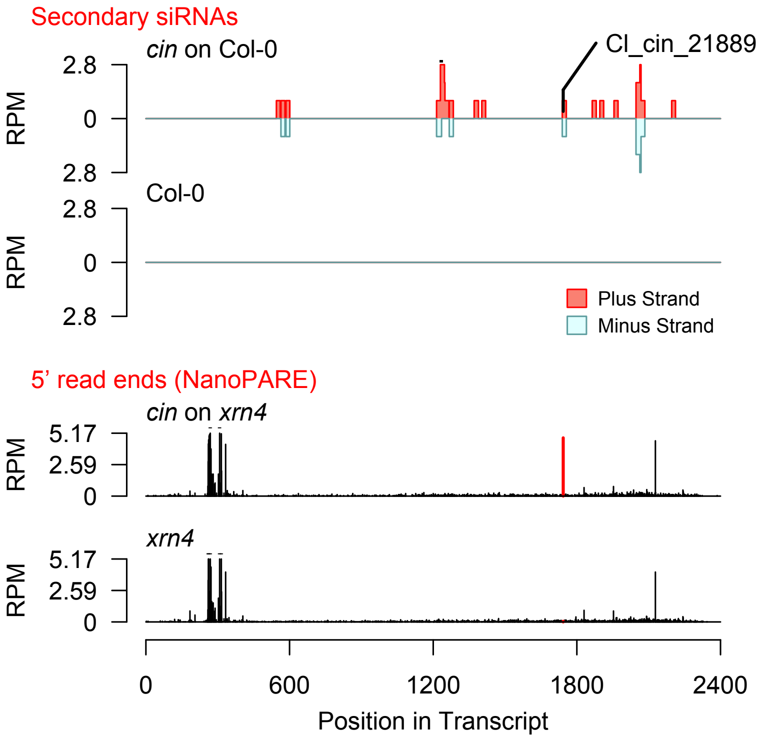


Diff. Exp. secondary siRNA locus not found

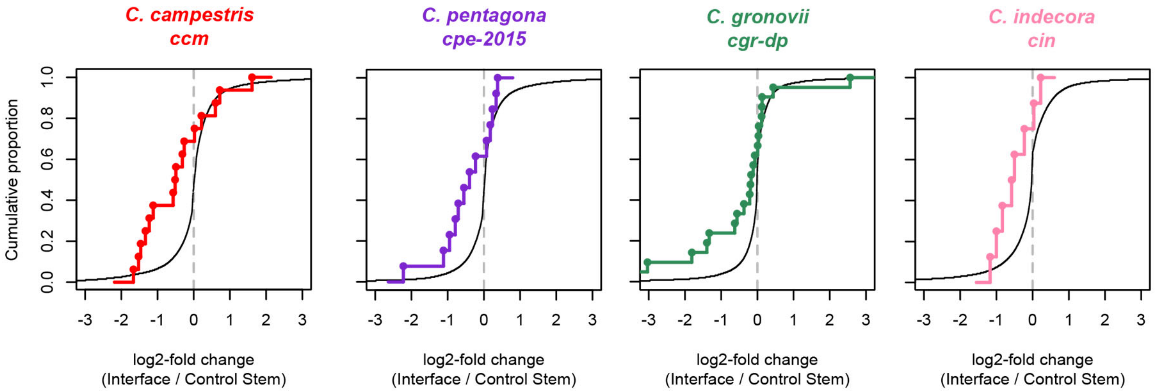


**Fig. S6. Analysis of gene expression in host-parasite interfaces**
Cumulative density plots of interface / control stem ratios for host genes expressed in *Cuscuta*-host interfaces. All genes shown with black line. Colored line and dots indicate genes which have confirmed targeting by a sRNA in the indicated *Cuscuta* isolate.

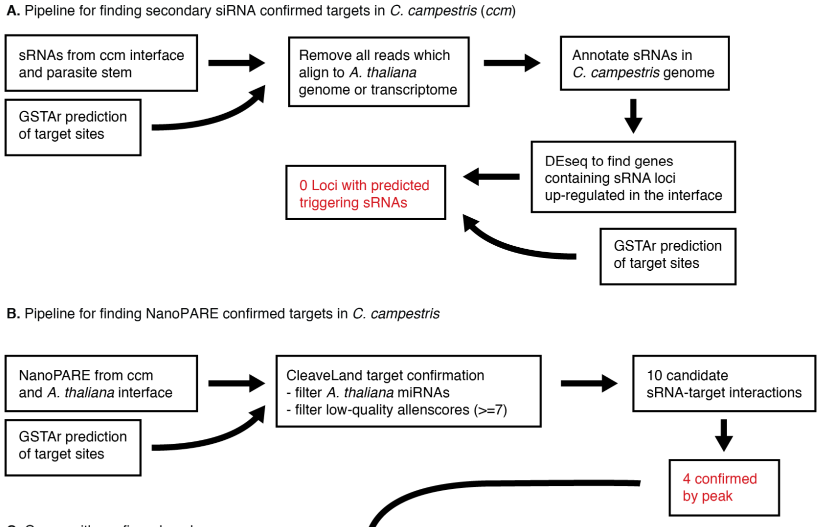


**C.** Genes with confirmed peaks

| Gene ID | Blast2Go annotation | sRNA | AllenScore |
|---|---|---|---|
| *Cc004391* | NDR1/HIN1-like protein 6 | Cl_ccm_15561 | 6 |
| *Cc025031* | V-type proton ATPase subunit E | Cl_ccm_23769 | 6 |
| *Cc035710* | hypothetical protein CTI12_AA531400 | Cl_ccm_9724 | 6 |
| *Cc041968* | threonine synthase, chloroplastic | Cl_ccm_328 | 3.5 |

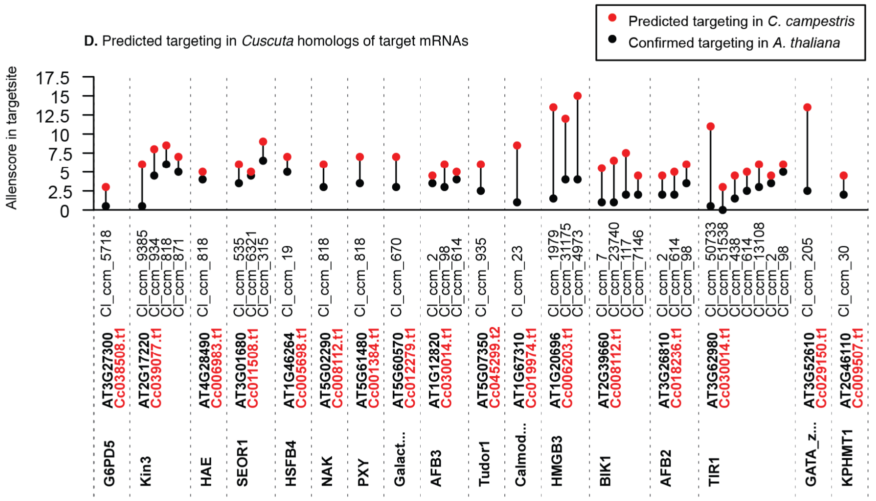


**Fig. S7. Experimental flowchart for confirming self-targeting of *C. campestris* mRNAs by HI-sRNAs**
(**A**) Pipeline for confirmation by the presence of secondary siRNAs. (**B**) Pipeline for confirmation by the 5' transcript sequencing (NanoPARE). (**C**) List of all mRNAs with strong evidence for self-targeting. (**D**) Target prediction scores for confirmed *A. thaliana* mRNA targets (black) and best-blast-hit homologs in *C. campestris* (red). All sRNAs with predicted targeting are shown.

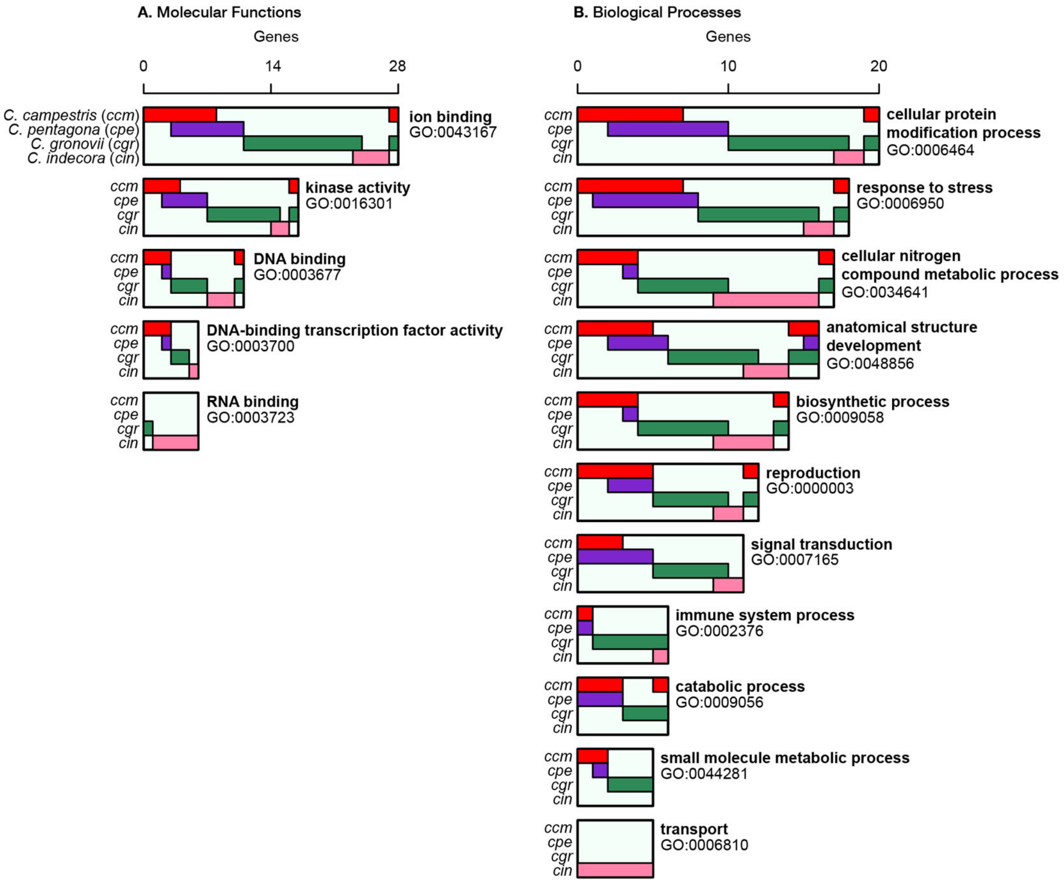


**Fig. S8. Most common GO terms for confirmed target genes**
(**A**) GO terms for molecular function with a nodescore >= 5.0, demonstrating the species for which the interaction is confirmed with colored bars. Locations where bars overlap indicate genes where both species have confirmed targeting. (**B**) Same as with A, but for biological processes.

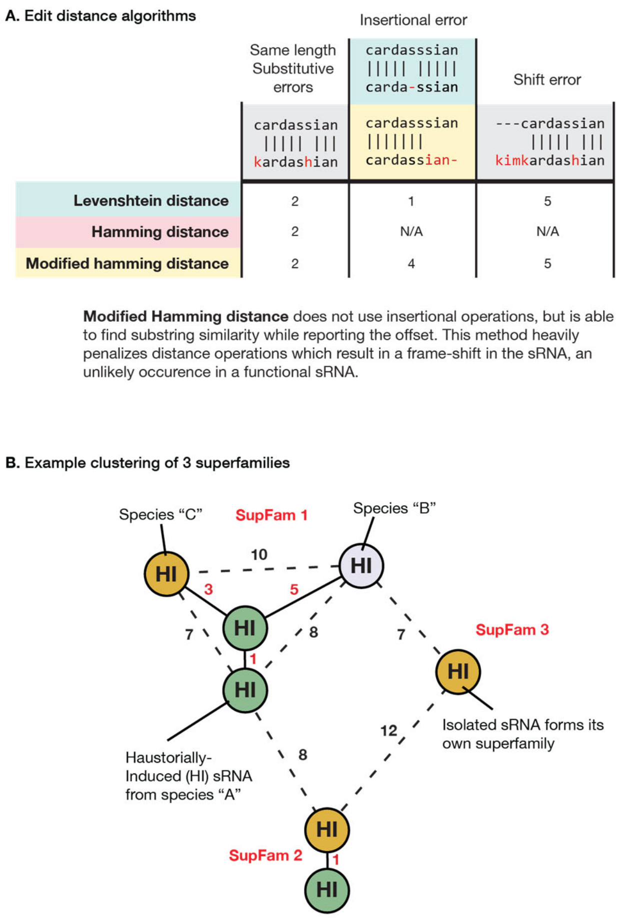


**Fig. S9. Clustering method for forming HI-sRNA superfamilies**
(**A**) Example demonstrating implementation of the "modified hamming distance" (mHD) when comparing strings. Levenshtein edit distance is tolerant of insertions and deletions, yet the mHD does not allow these operations, making a high penalty to strings which contain insertional errors while shift errors are penalized the same. (**B**) Example of clustering seven HI-sRNAs into three superfamilies using mHD. Species are indicated by color; clustering is independent of species. Edges close enough to form a cluster (solid line, red distance number) and inadequate edges (dashed line, black distance number) connect HI-sRNA nodes. Cutoff for clustering is an mHD distance of five or less and it is not required that all nodes in a cluster must meet this threshold (must have one adequate edge to join a cluster).

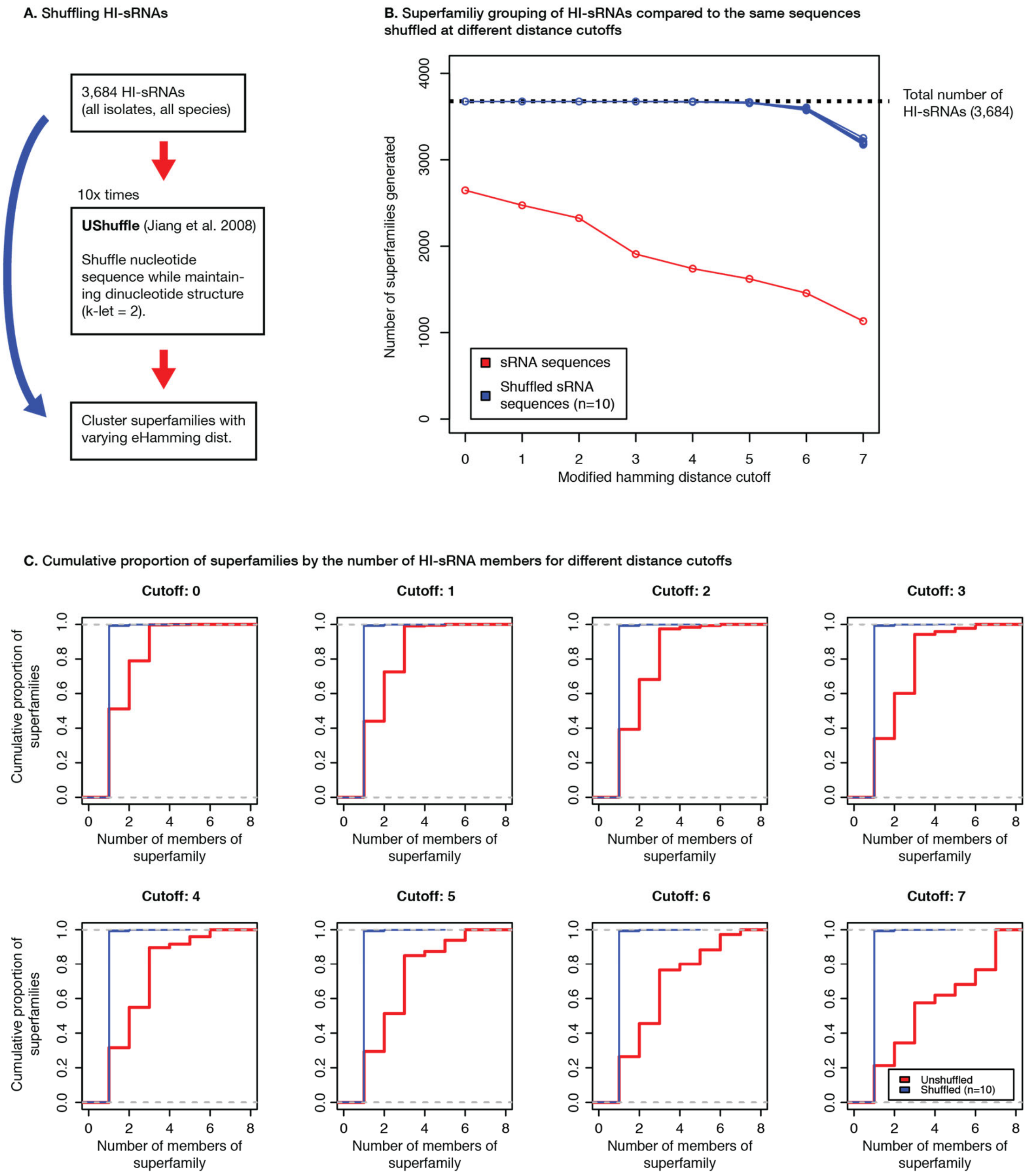


**Fig. S10. Testing distance cutoff parameters for superfamily formation**
(**A**) Experimental pipeline for testing cutoff. sRNA libraries are shuffled using UShuffle maintaining dinucleotide composition. (**B**) Number of superfamilies formed for real HI-sRNAs and shuffled libraries by maximum distance allowed for cluster formation. Smaller count of superfamilies means that more HI-sRNAs are successfully clustering with each other. (**C**) The same analysis as in B, except demonstrating the cumulative density of superfamilies by the number of sRNAs grouped in them. Larger cutoffs yield larger superfamilies, with shuffled libraries remaining unable to form clusters larger than one or two.

**Fig. S11. Target interactions with significant correlation of variation in superfamily and target site**
Multiple sequence alignments of HI-sRNA superfamilies which have significant correlations between sRNA positional variation and target site variation. Alignment of eudicot homologs around target site also shown, with nucleotide and amino acid Shannon entropy shown as bits. Vertical red lines indicate the frame. Dots indicate the number of possible synonymous nucleotides at a position for the confirmed target's sequence. Nucleotide positions are in reference to the position in the multiple sequence alignment.

**AT5G66850 - MAPKKK5**
**36/36 homologs found in phytozome eudicots containing targetsite**

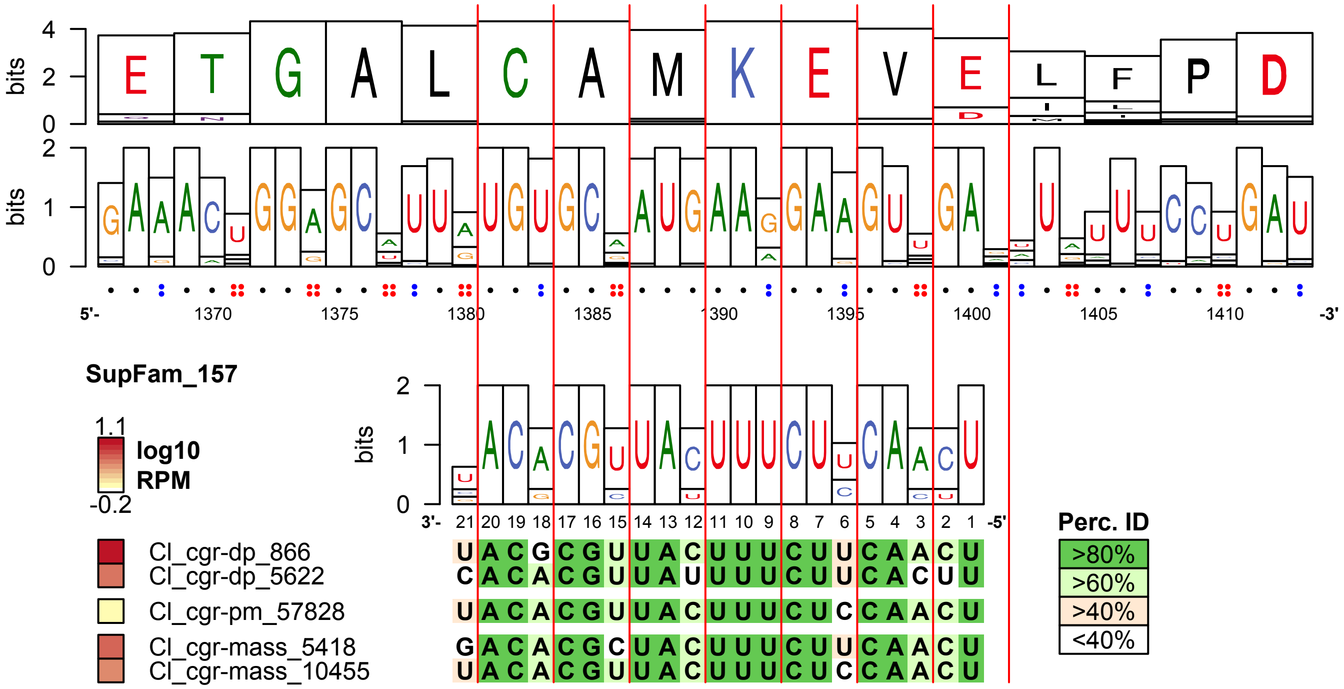

**AT3G45640 - MPK3**

**35/36 homologs found in phytozome eudicots containing targetsite**

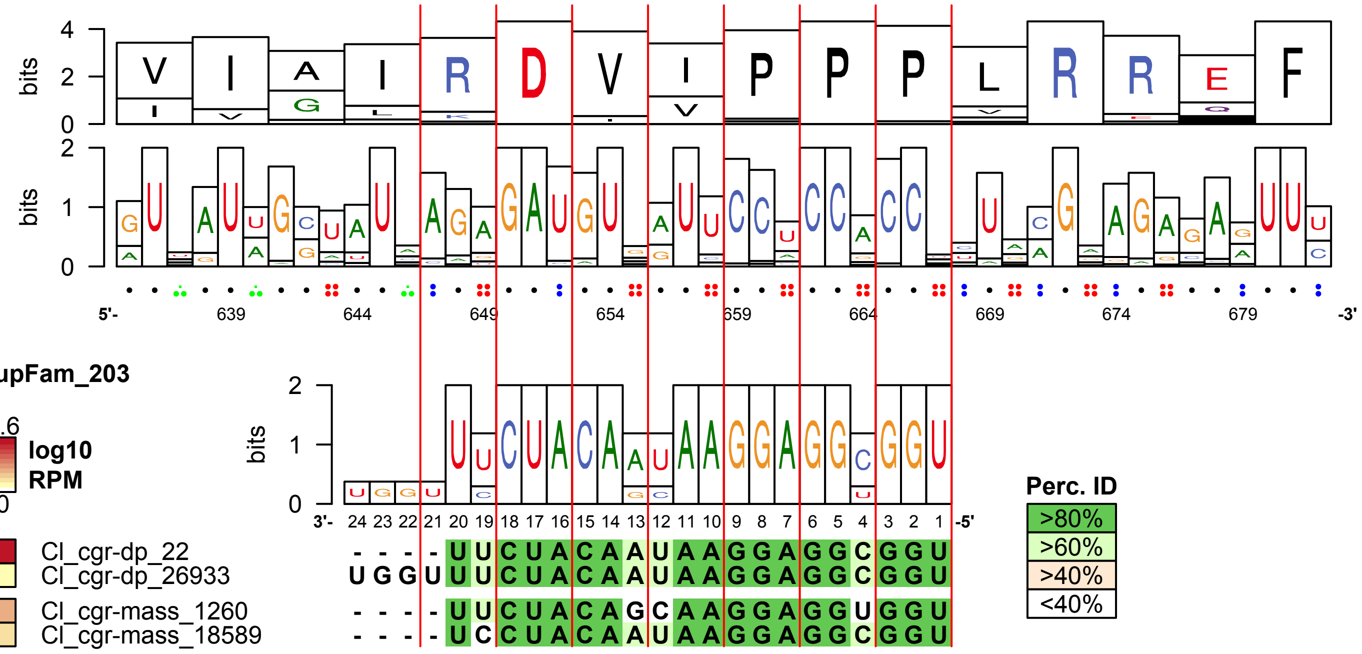

**AT1G53440 - Leucine-rich_repeat_transmembrane_protein_kinase**
**36/36 homologs found in phytozome eudicots containing targetsite**

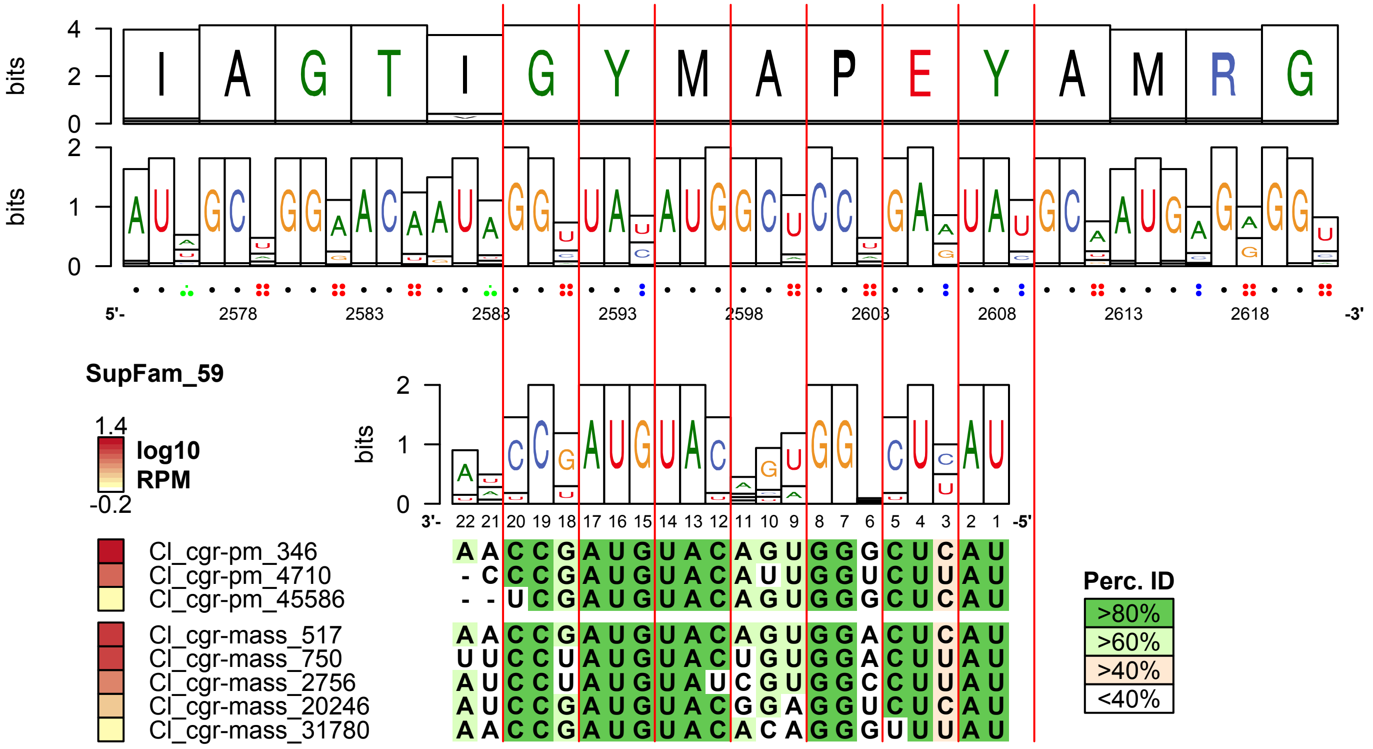

**AT4G28490 - HAE**
**34/36 homologs found in phytozome eudicots containing targetsite**

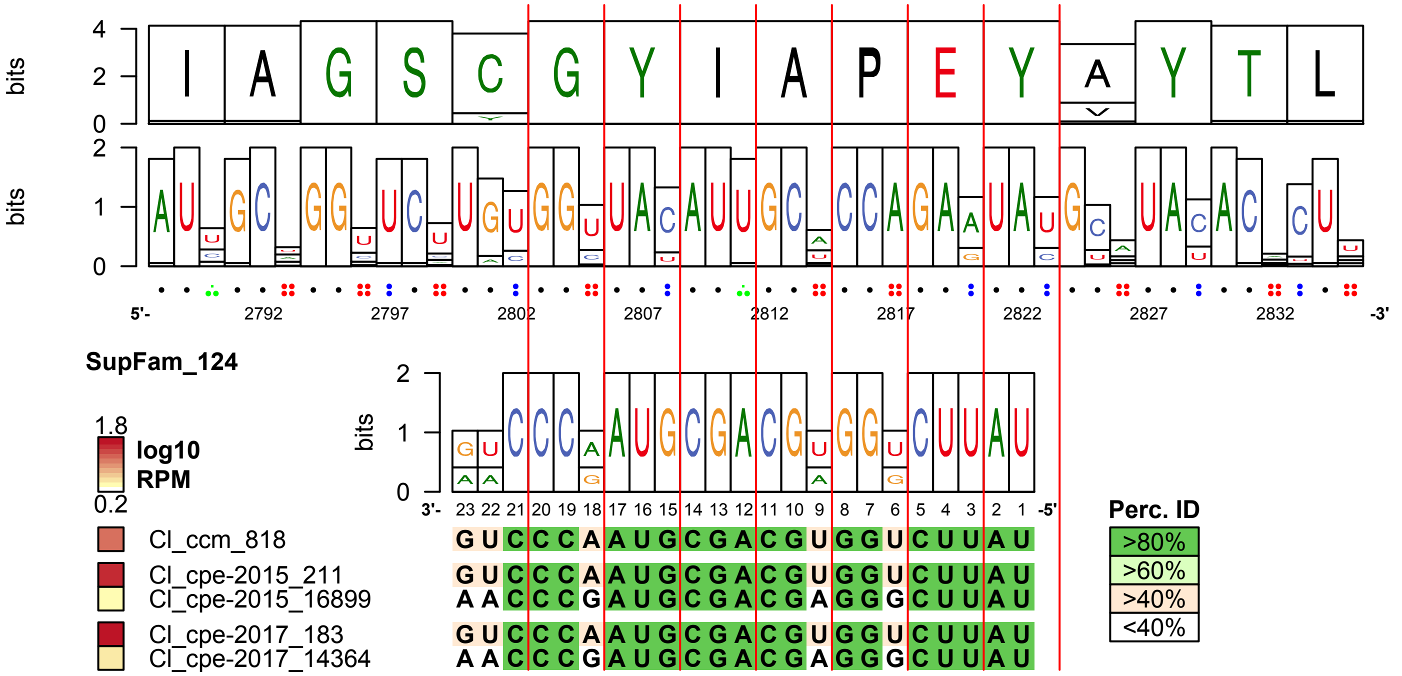

**AT1G12820 - AFB3**
**36/36 homologs found in phytozome eudicots containing targetsite**

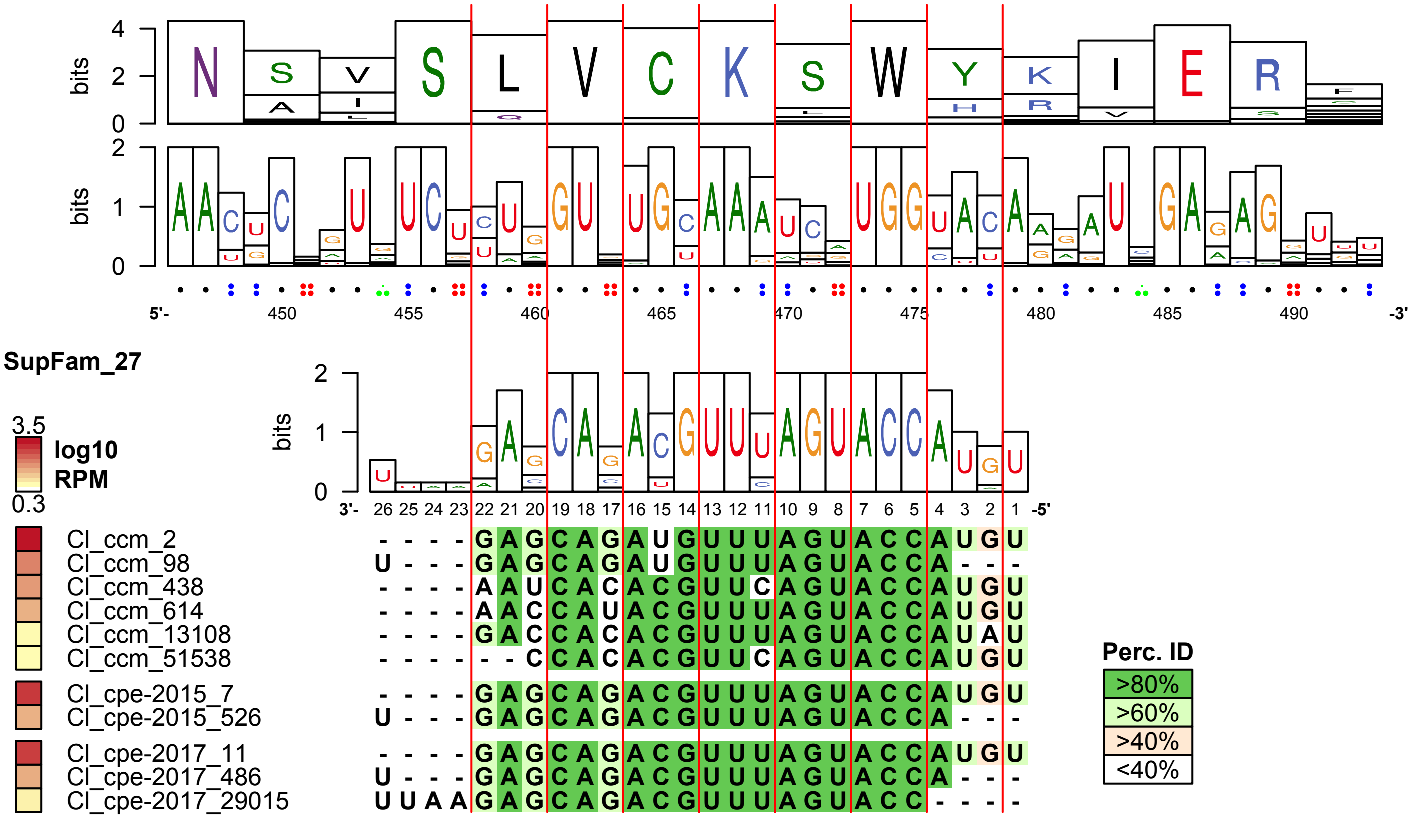

**AT3G01680 - SEOR1**

**34/36 homologs found in phytozome eudicots containing targetsite**

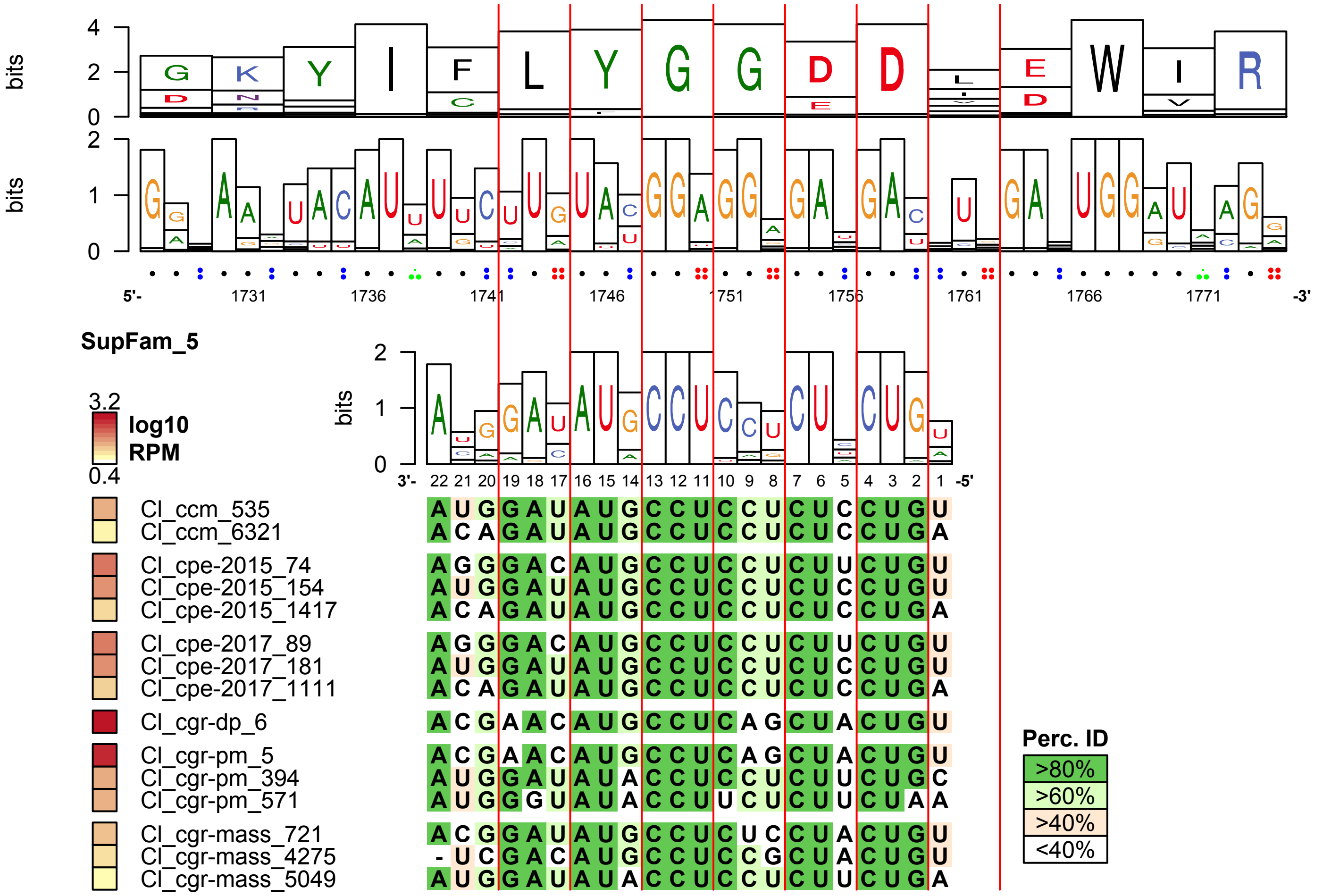

**AT4G39400 - BRI1**
**36/36 homologs found in phytozome eudicots containing targetsite**

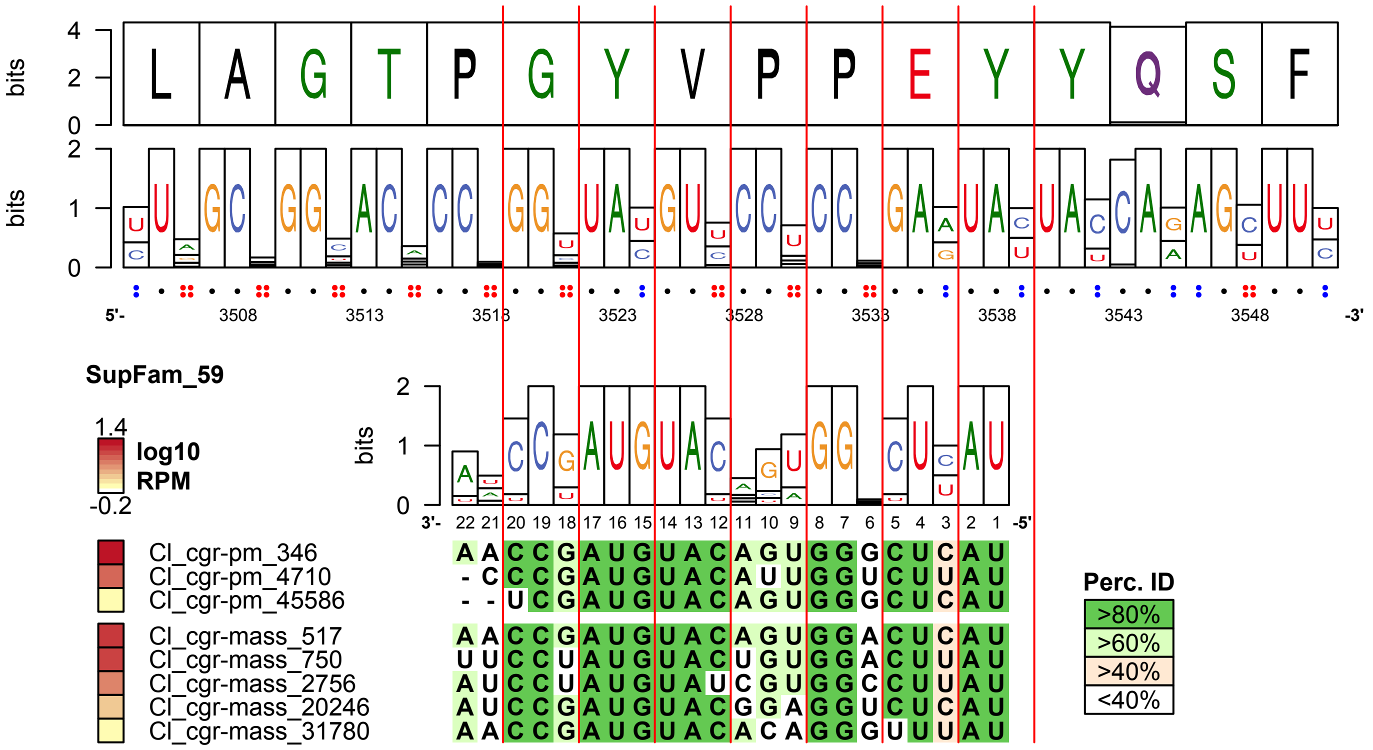

**AT5G02290 - NAK**

**36/36 homologs found in phytozome eudicots containing targetsite**

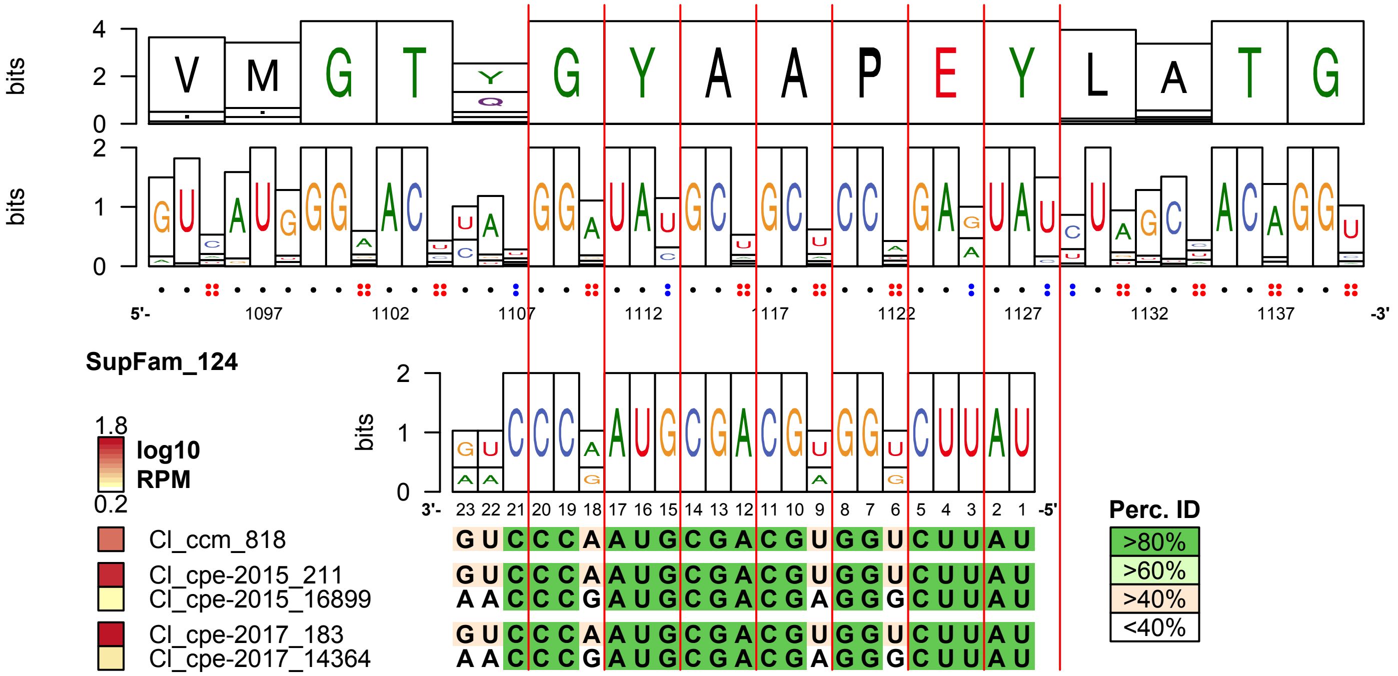

**AT2G02800 - APK2B**
**33/36 homologs found in phytozome eudicots containing targetsite**

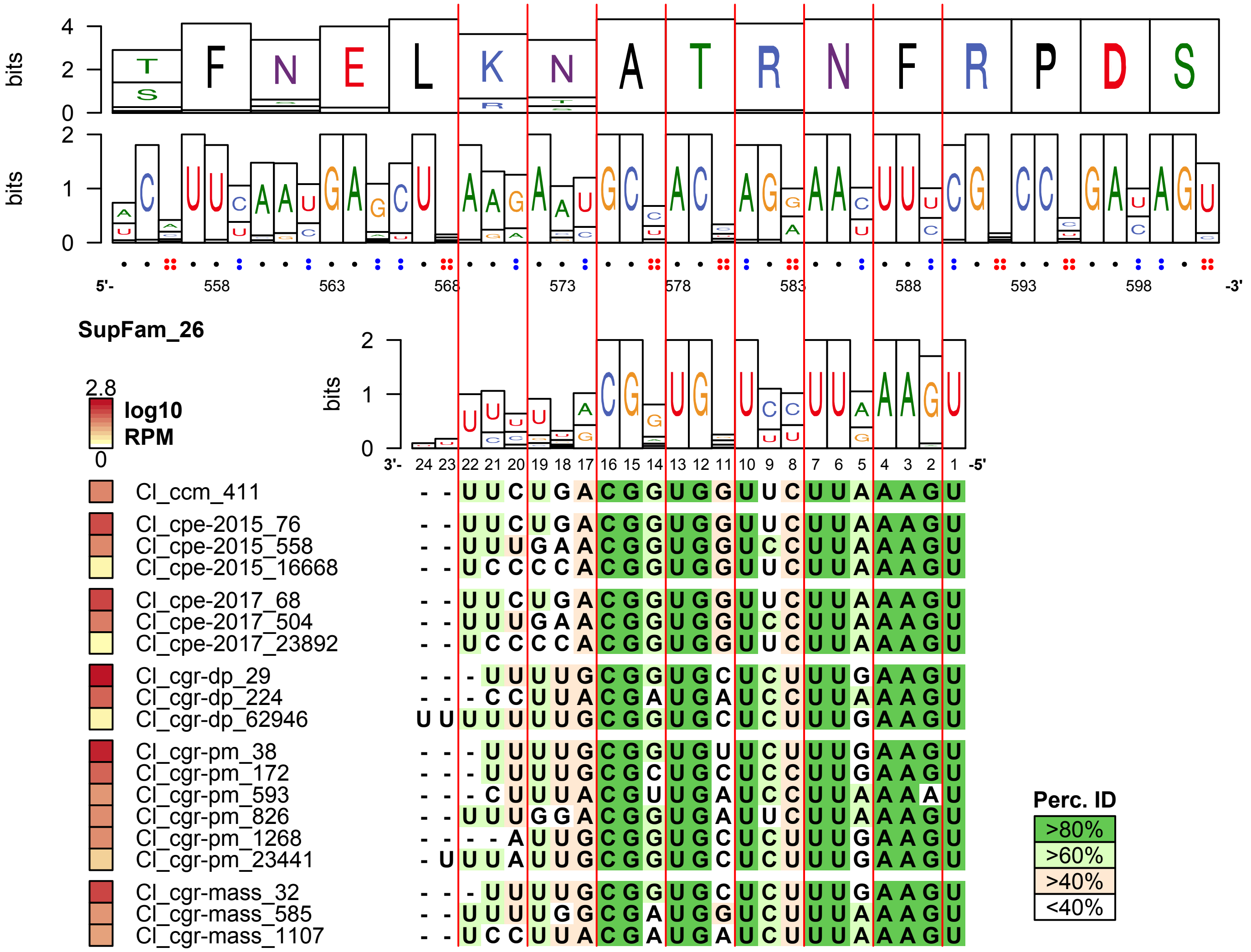

**AT5G65430 - GRF8**
**36/36 homologs found in phytozome eudicots containing targetsite**

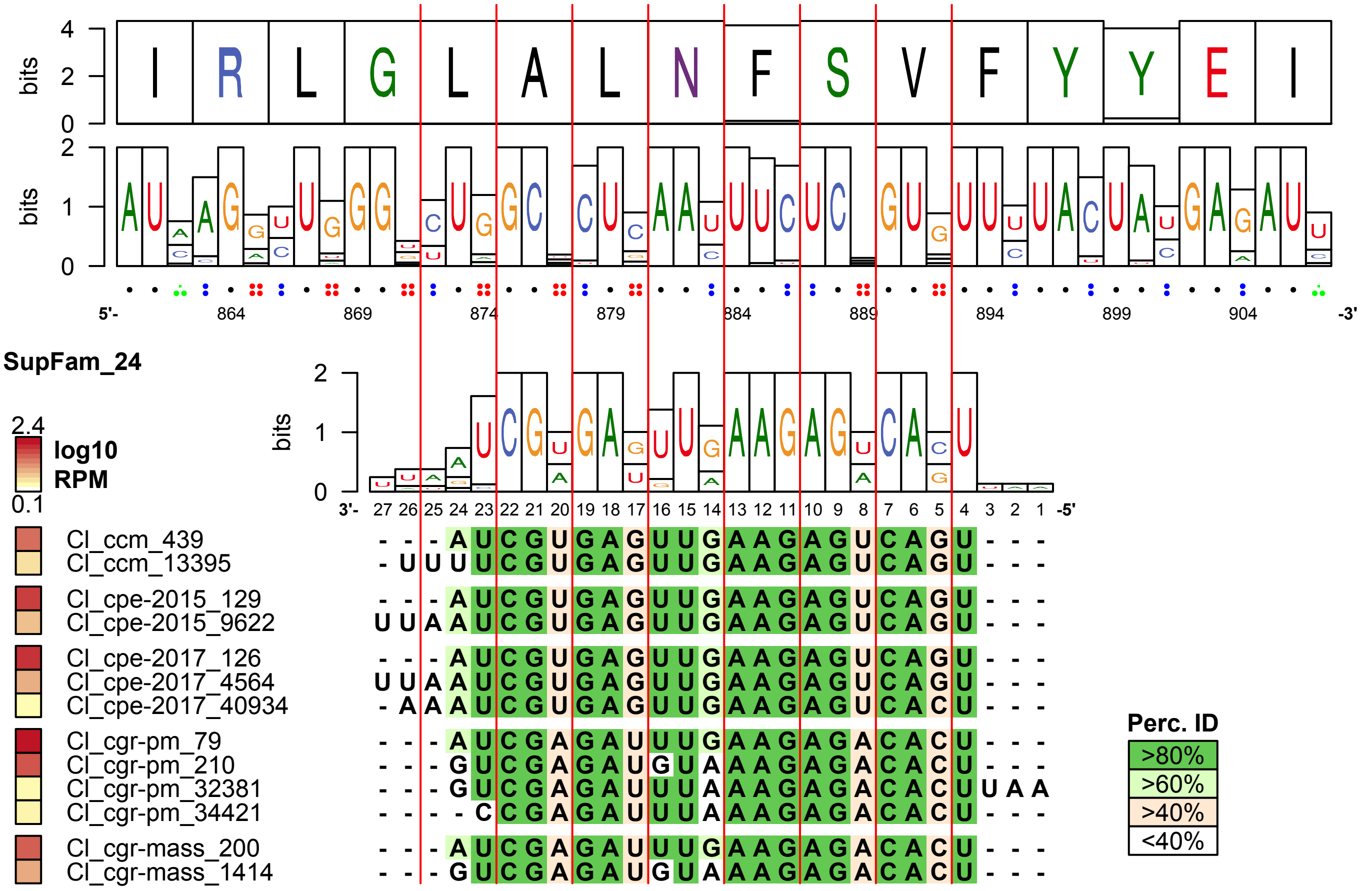

**AT5G24010 - Protein_kinase_superfamily_protein**
**36/36 homologs found in phytozome eudicots containing targetsite**

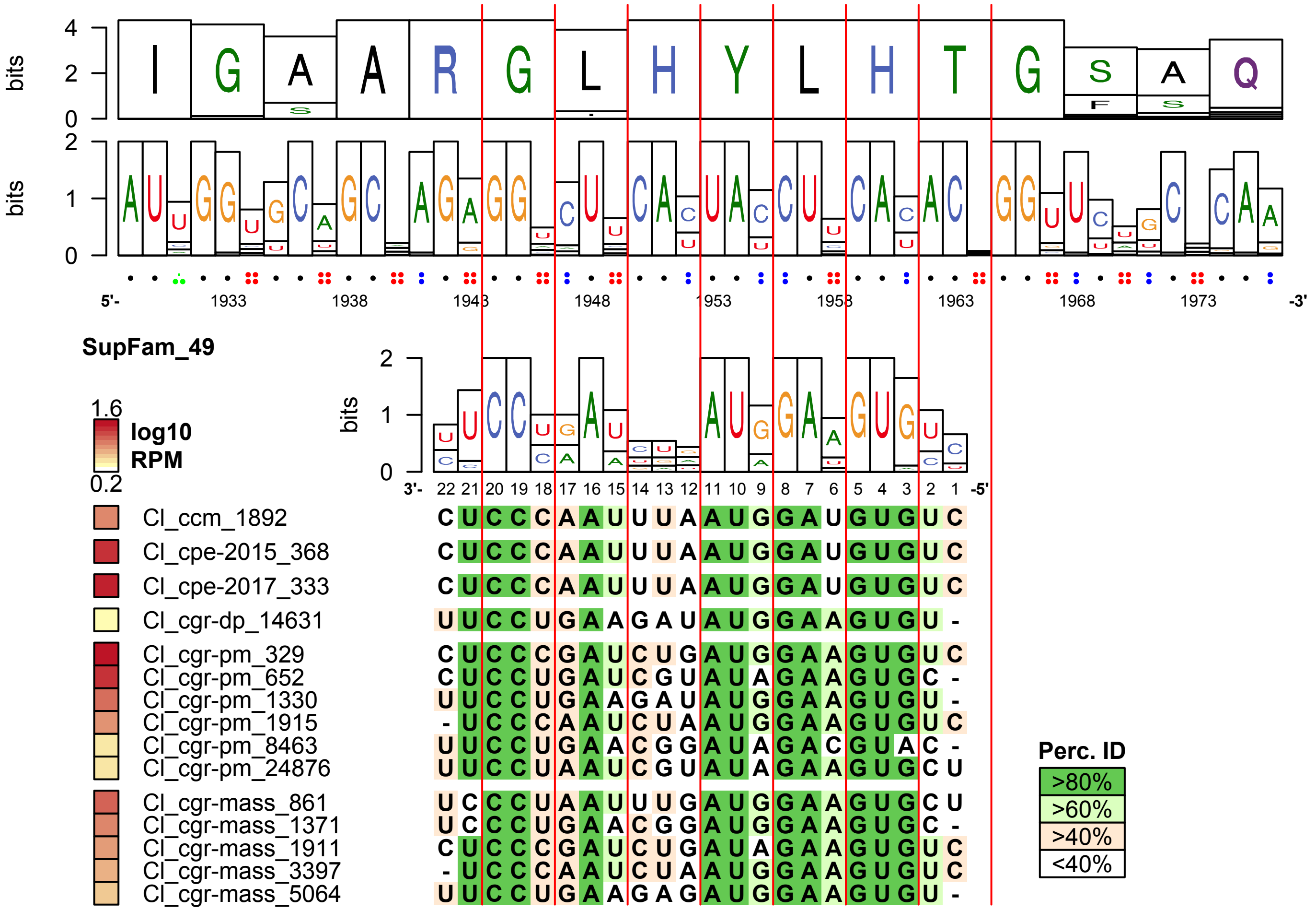

**AT3G62980 - TIR1**
**35/36 homologs found in phytozome eudicots containing targetsite**

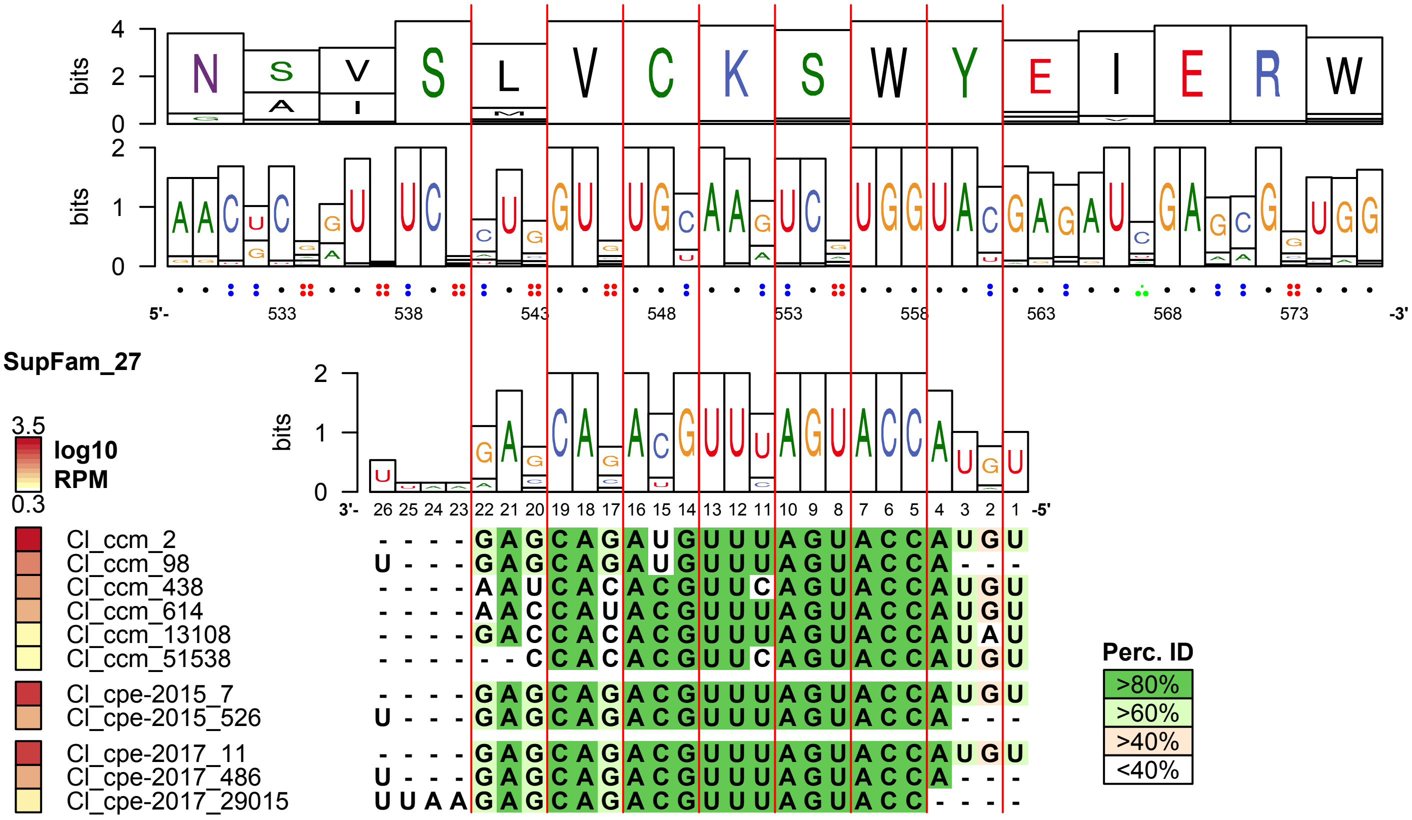

**AT1G20696 - HMGB3**
**35/36 homologs found in phytozome eudicots containing targetsite**

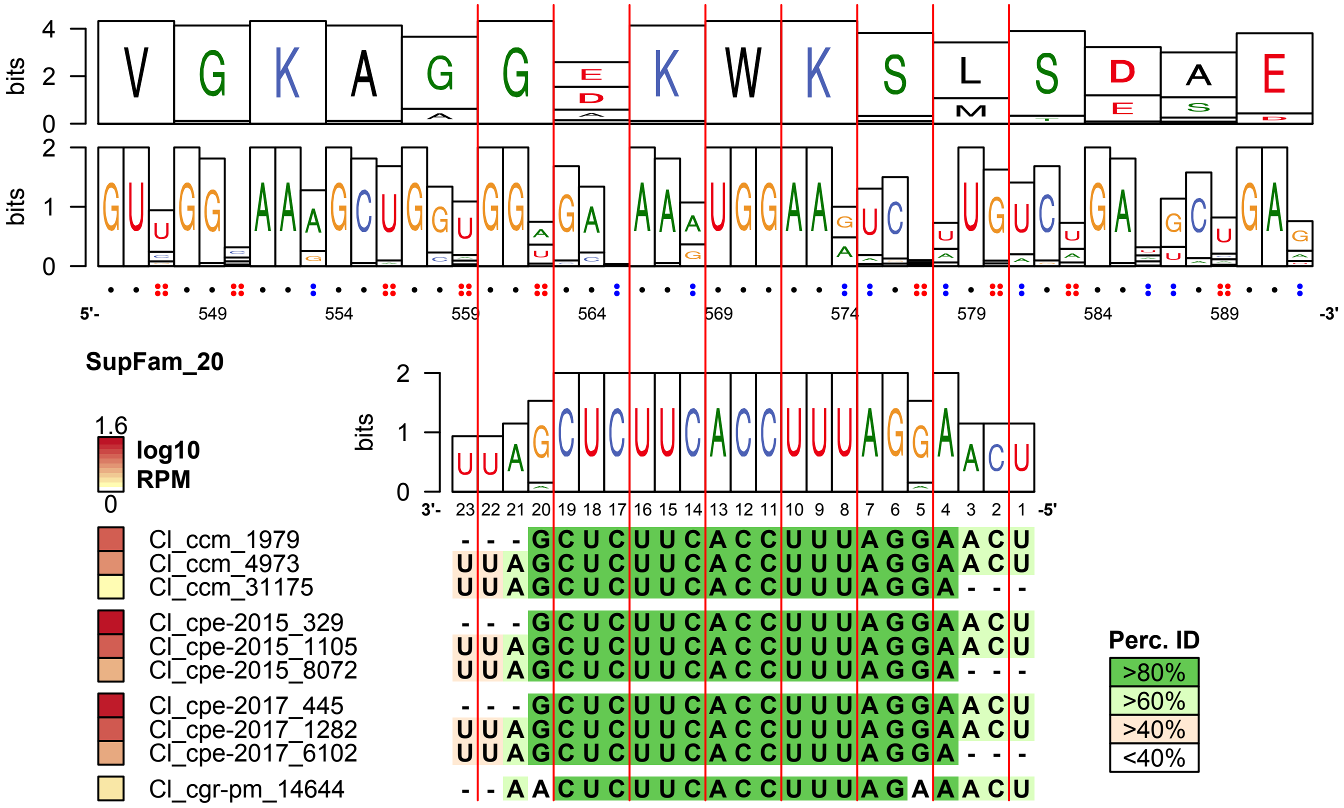

**AT4G03190 - AFB1**
**36/36 homologs found in phytozome eudicots containing targetsite**

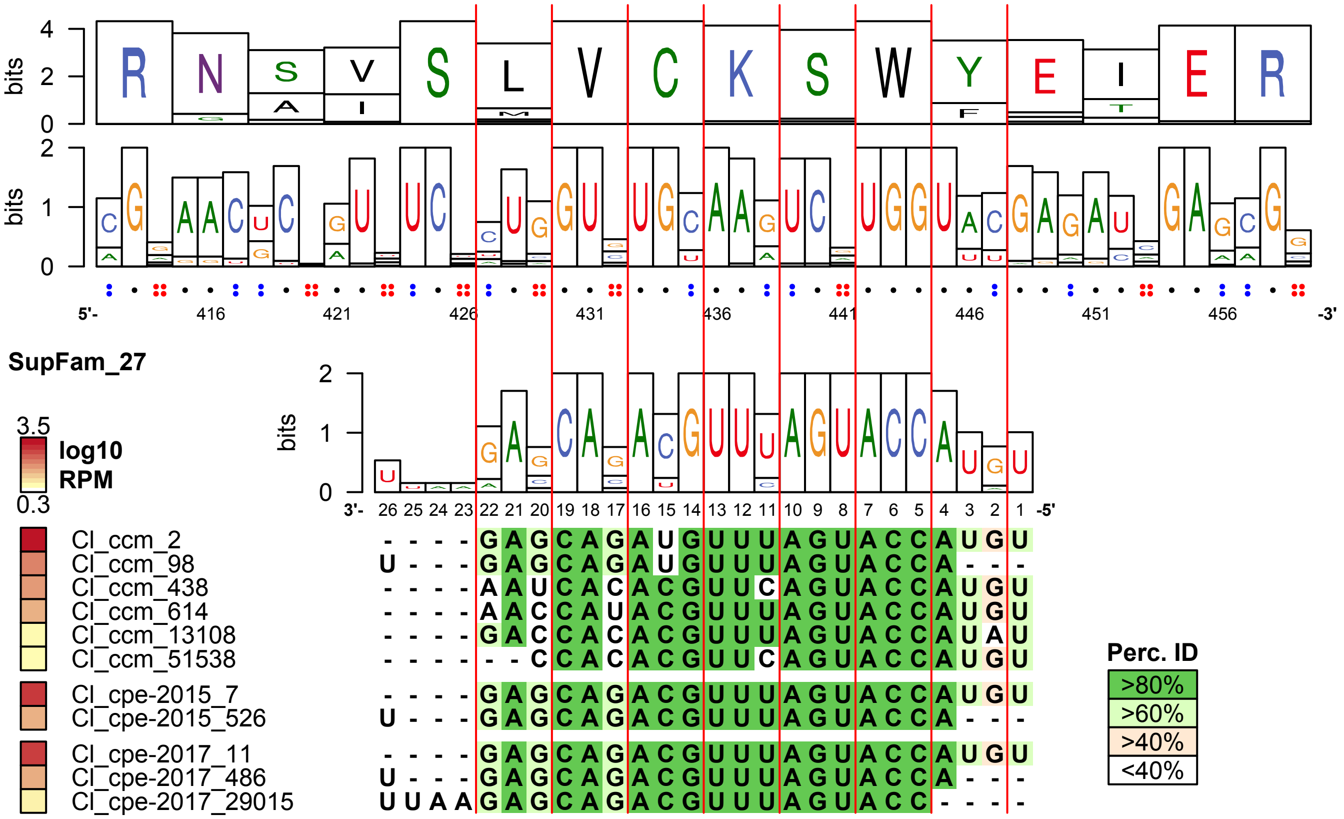

**AT3G26810 - AFB2**

**36/36 homologs found in phytozome eudicots containing targetsite**

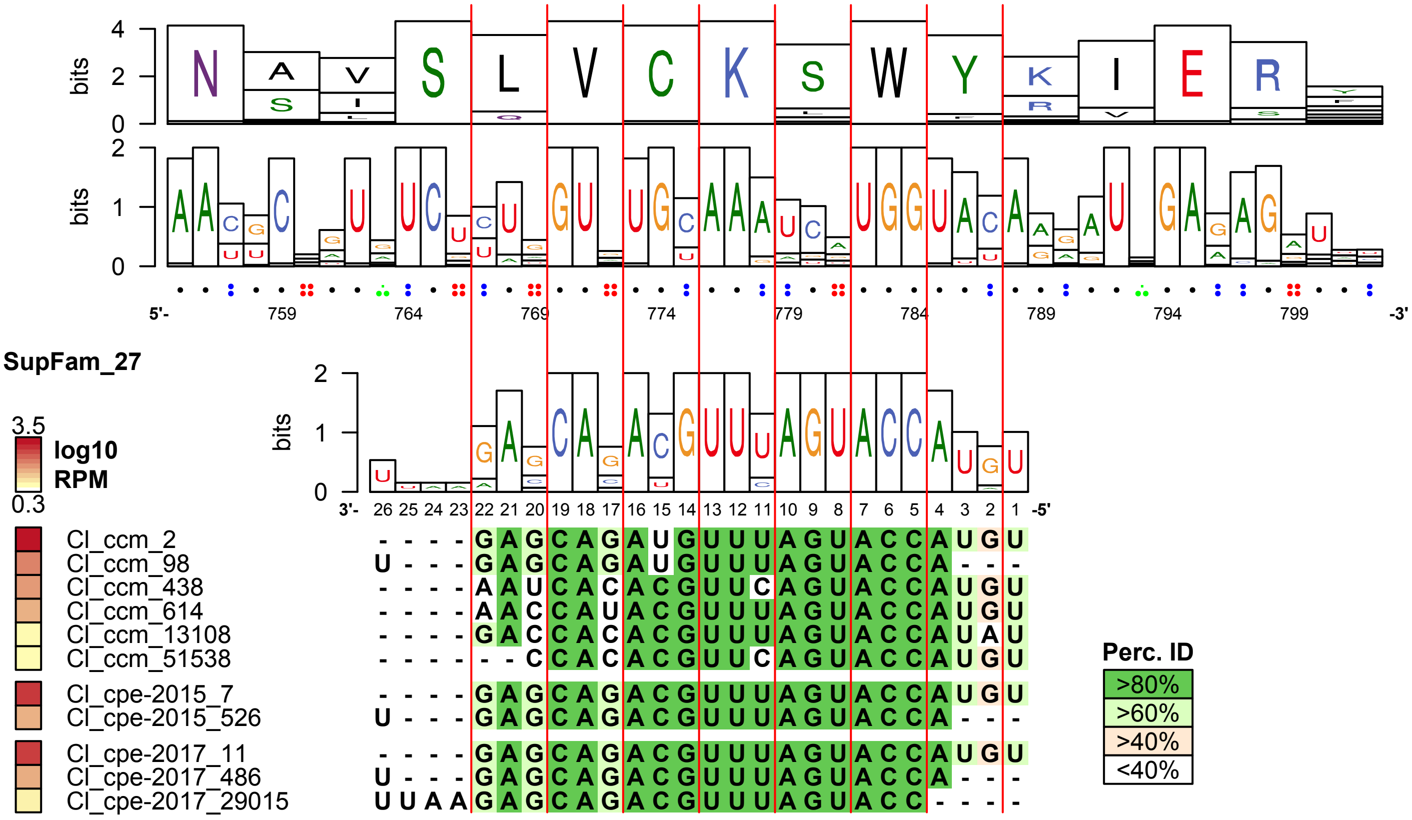

**AT1G63910 - AtMYB103**
**36/36 homologs found in phytozome eudicots containing targetsite**

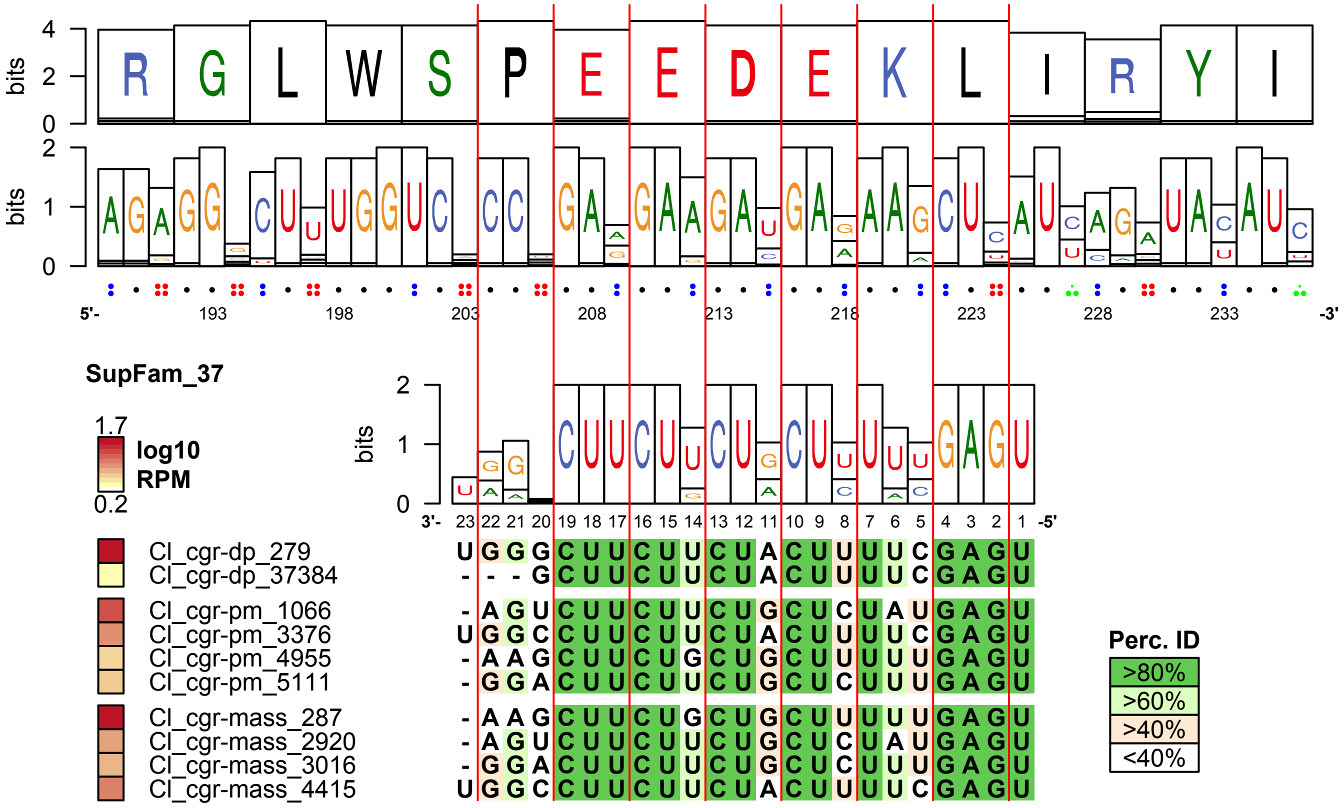

**AT1G65800 - RK2**
**35/36 homologs found in phytozome eudicots containing targetsite**

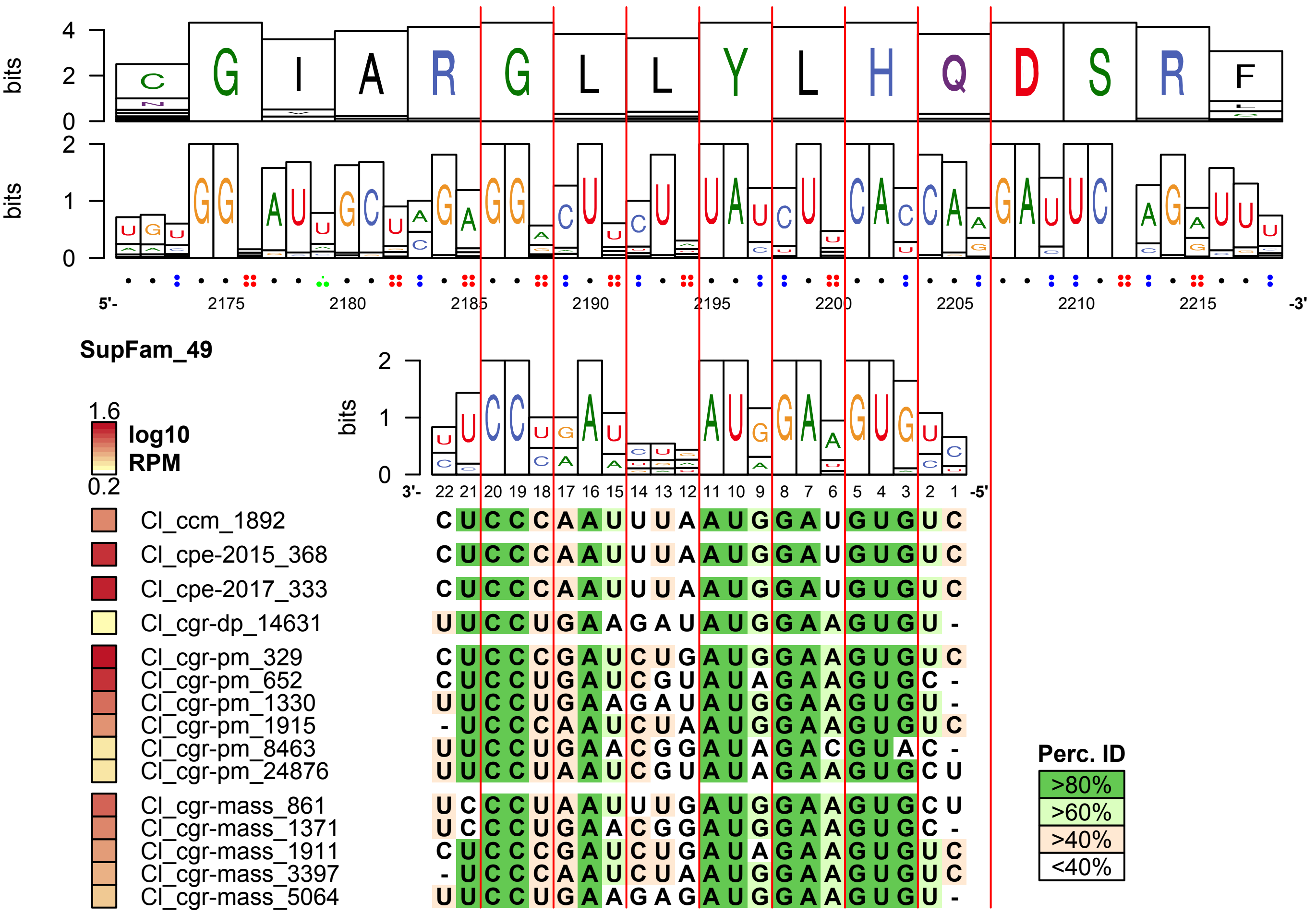

**AT2G39660 - BIK1**

**36/36 homologs found in phytozome eudicots containing targetsite**

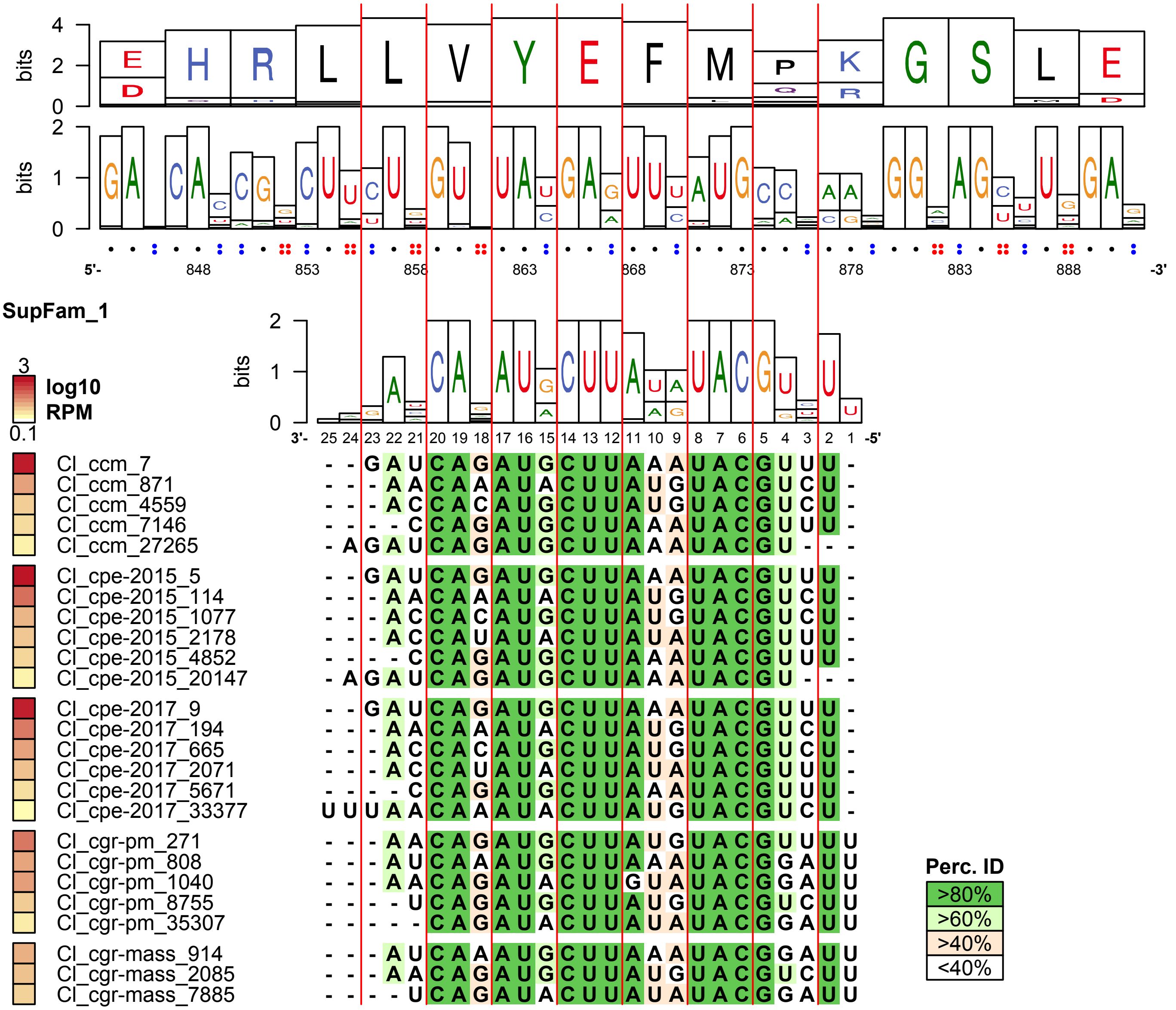

**Fig. S12. Target interactions of *A. thaliana* homologs with conserved target motifs**
Multiple sequence alignments sRNA of superfamilies and conserved target motifs found in *Arabidopsis* transcriptome, with nucleotide and amino acid Shannon entropy shown as bits. Vertical red lines indicate the frame. Dots indicate the number of possible synonymous nucleotides at a position for the confirmed target's sequence. Nucleotide positions are in reference to the position in the multiple sequence alignment. Color of gene names indicates if there is evidence for targeting in NanoPARE data (black - 0 replicates; orange - 1 or 2 replicates; red - 3 replicates, confirmed interaction)

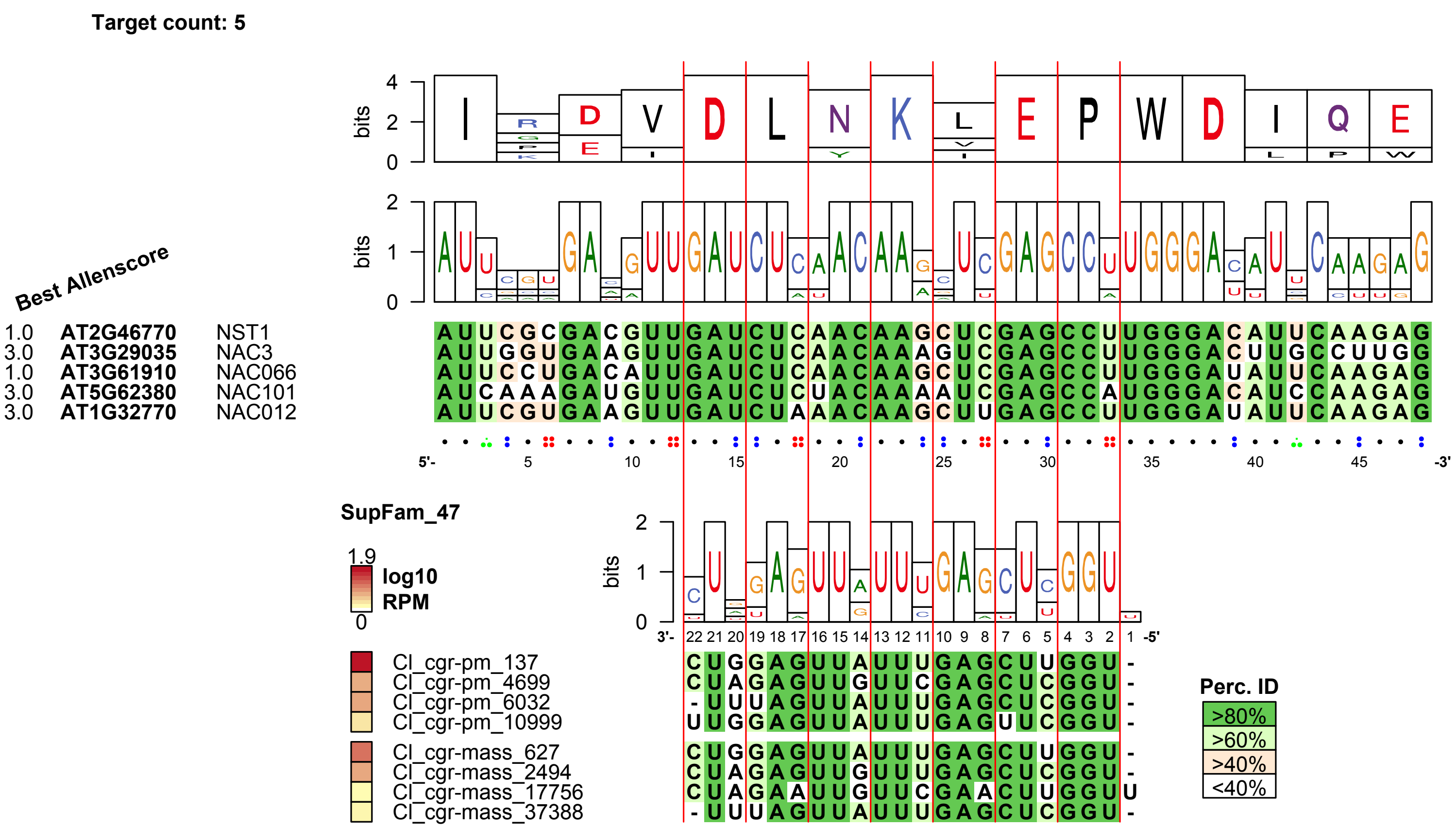

Target count: 5
Best Allenscore
1.0 AT2G46770 NST1 AUUCGCGACGUUGAUCUCAACAAGCUCGAGCCUUGGGACAUUCAAGAG
3.0 AT3G29035 NAC3 AUUGGUGAAGUUGAUCUCAACAAAGUCGAGCCUUGGGACUUGCCUUGG
1.0 AT3G61910 NAC066 AUUCCUGACAUUGAUCUCAACAAGCUCGAGCCUUGGGAUAUUCAAGAG
3.0 AT5G62380 NAC101 AUCAAAGAUGUUGAUCUCUACAAAAUCGAGCCAUGGGACAUCCAAGAG
3.0 AT1G32770 NAC012 AUUCGUGAAGUUGAUCUAAACAAGCUUGAGCCUUGGGAUAUUCAAGAG
5'- 5 10 15 20 25 30 35 40 45 -3'
SupFam_47
1.9 log10 RPM 0
3'- 22 21 20 19 18 17 16 15 14 13 12 11 10 9 8 7 6 5 4 3 2 1 -5'
Cl_cgr-pm_137 CUGGAGUUAUUUGAGCUUGGU -
Cl_cgr-pm_4699 CUAGAGUUGUUCGAGCUCGGU -
Cl_cgr-pm_6032 - UUUAGUUAUUUGAGCUCGGU -
Cl_cgr-pm_10999 UUGGAGUUAUUUGAGUUCGGU -
Cl_cgr-mass_627 CUGGAGUUAUUUGAGCUUGGU -
Cl_cgr-mass_2494 CUAGAGUUGUUUGAGCUCGGU -
Cl_cgr-mass_17756 CUAGAAUUGUUCGAACUUGGUU
Cl_cgr-mass_37388 - UUUAGUUAUUUGAGCUCGGU -
Perc. ID
>80%
>60%
>40%
<40%

**Target count: 5**

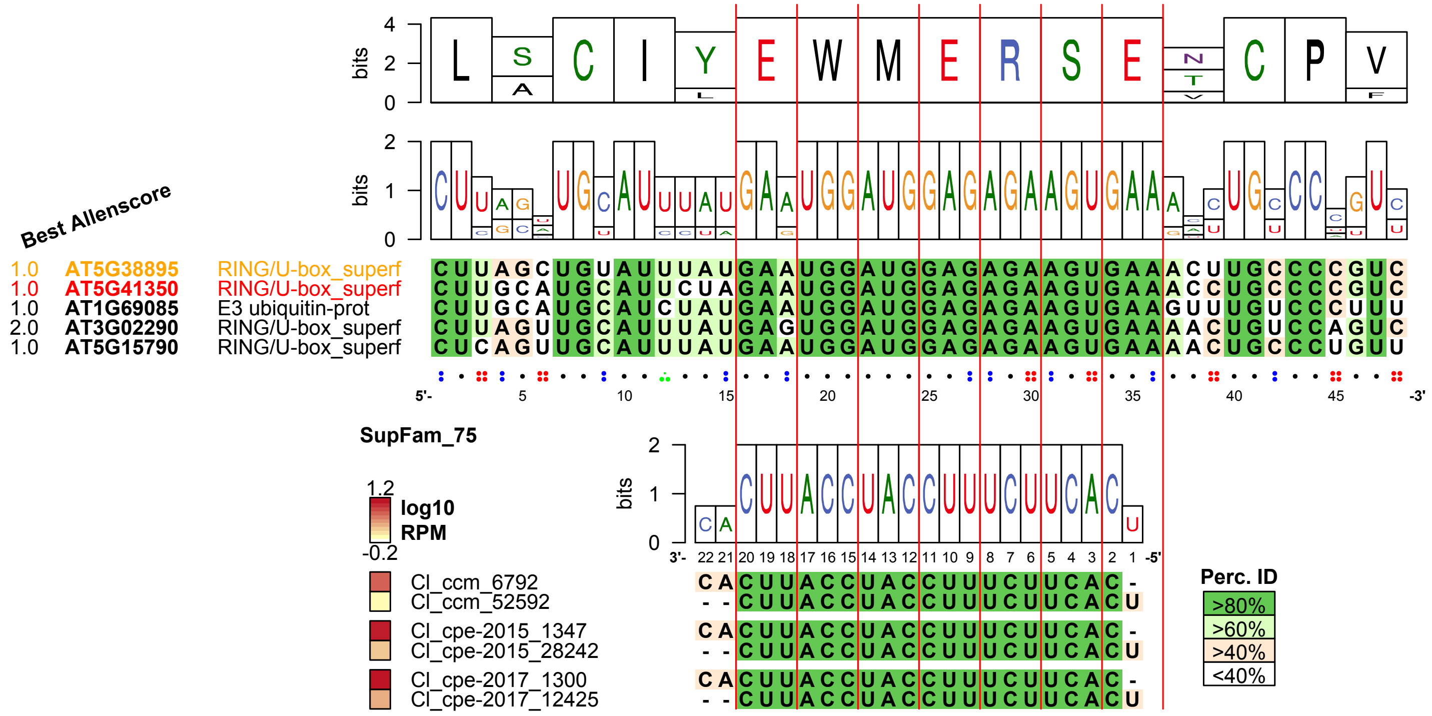

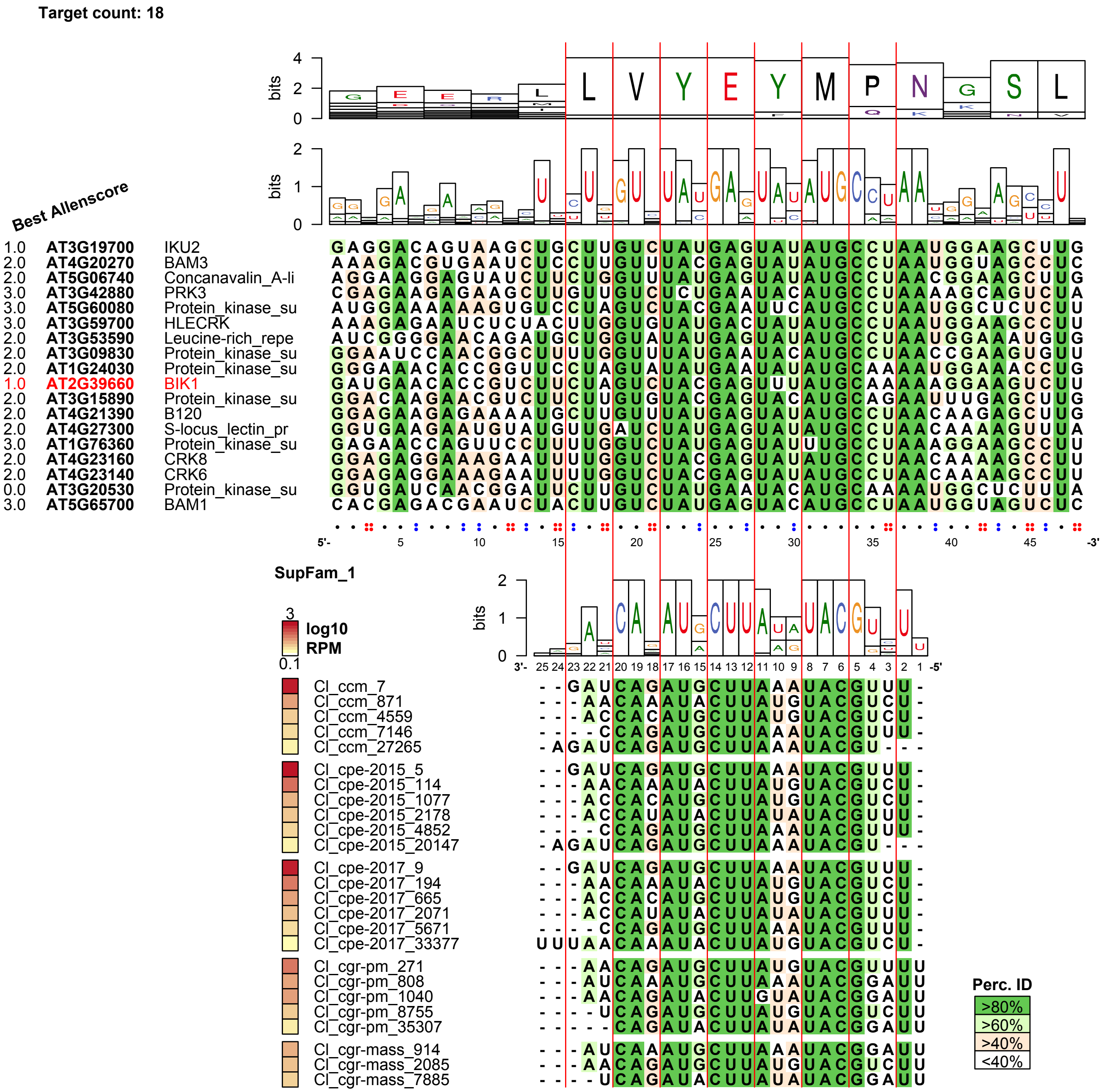

Target count: 18
Best Allenscore
1.0 AT3G19700 IKU2
2.0 AT4G20270 BAM3
2.0 AT5G06740 Concanavalin_A-li
3.0 AT3G42880 PRK3
3.0 AT5G60080 Protein_kinase_su
3.0 AT3G59700 HLECRK
2.0 AT3G53590 Leucine-rich_repe
2.0 AT3G09830 Protein_kinase_su
2.0 AT1G24030 Protein_kinase_su
1.0 AT2G39660 BIK1
2.0 AT3G15890 Protein_kinase_su
2.0 AT4G21390 B120
2.0 AT4G27300 S-locus_lectin_pr
3.0 AT1G76360 Protein_kinase_su
2.0 AT4G23160 CRK8
2.0 AT4G23140 CRK6
0.0 AT3G20530 Protein_kinase_su
3.0 AT5G65700 BAM1
SupFam_1
log10 RPM
Cl_ccm_7
Cl_ccm_871
Cl_ccm_4559
Cl_ccm_7146
Cl_ccm_27265
Cl_cpe-2015_5
Cl_cpe-2015_114
Cl_cpe-2015_1077
Cl_cpe-2015_2178
Cl_cpe-2015_4852
Cl_cpe-2015_20147
Cl_cpe-2017_9
Cl_cpe-2017_194
Cl_cpe-2017_665
Cl_cpe-2017_2071
Cl_cpe-2017_5671
Cl_cpe-2017_33377
Cl_cgr-pm_271
Cl_cgr-pm_808
Cl_cgr-pm_1040
Cl_cgr-pm_8755
Cl_cgr-pm_35307
Cl_cgr-mass_914
Cl_cgr-mass_2085
Cl_cgr-mass_7885
Perc. ID
>80%
>60%
>40%
<40%

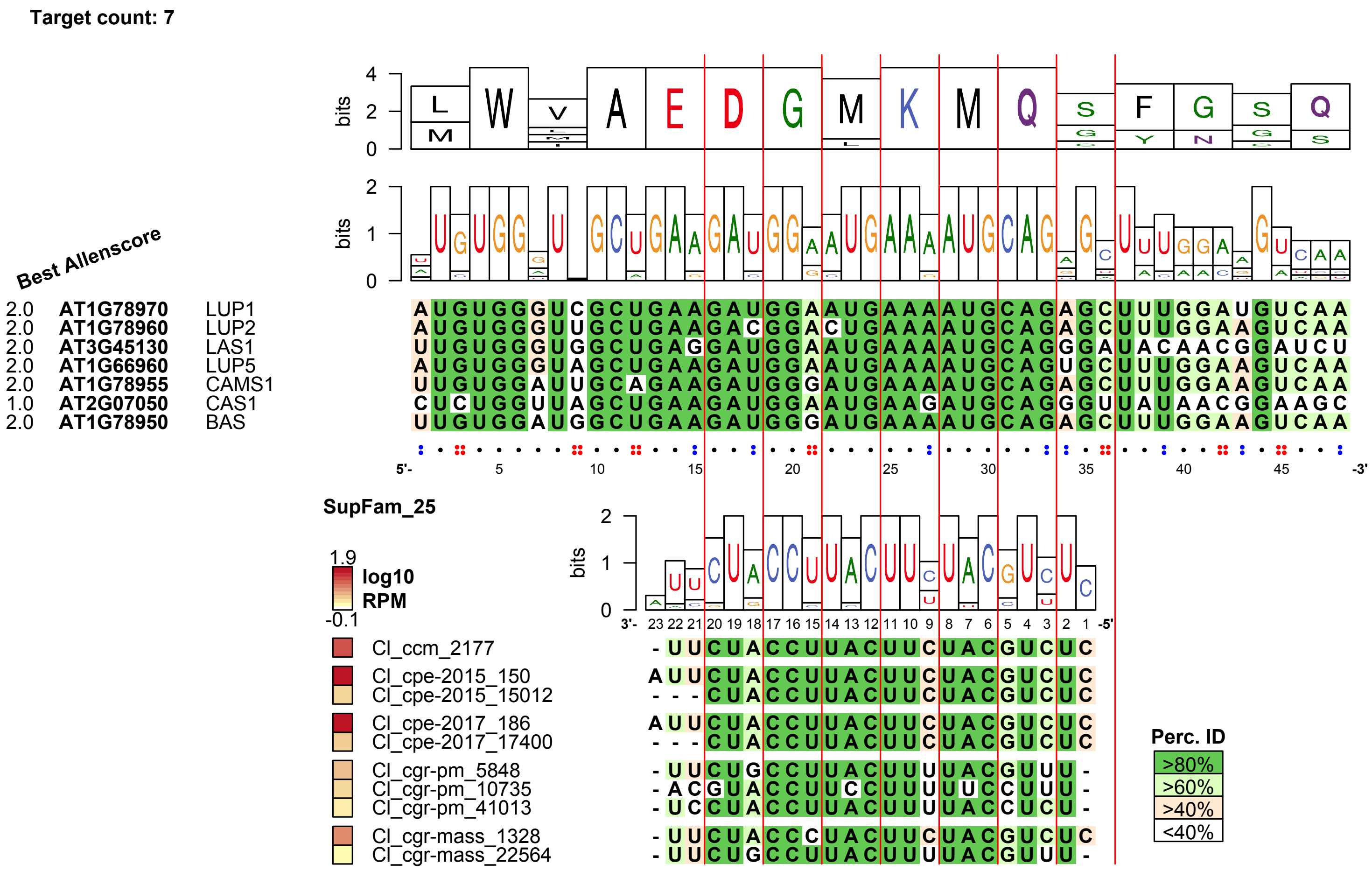

Target count: 7
Best Allenscore
L M W V A E D G M K M Q S G F Y G N S G Q S
bits
4 2 0
2 1 0
2.0 AT1G78970 LUP1 AUGUGGGUCGCUGAAGAUGGAAUGAAAAUGCAGAGCUUUGGAUGUCAA
2.0 AT1G78960 LUP2 AUGUGGGUUGCUGAAGACGGACUGAAAAUGCAGAGCUUUGGAAGUCAA
2.0 AT3G45130 LAS1 UUGUGGGUGGCUGAGGAUGGAAUGAAAAUGCAGGGAUACAACGGAUCU
2.0 AT1G66960 LUP5 AUGUGGGUAGCUGAAGAUGGAAUGAAAAUGCAGUGCUUUGGAAGUCAA
2.0 AT1G78955 CAMS1 UUGUGGAUUGCAGAAGAUGGGAUGAAAAUGCAGAGCUUUGGAAGUCAA
1.0 AT2G07050 CAS1 CUCUGGUUAGCUGAAGAUGGAAUGAAGAUGCAGGGUUAUAACGGAAGC
2.0 AT1G78950 BAS UUGUGGAUGGCUGAAGAUGGGAUGAAAAUGCAGAGCUUUGGAAGUCAA
5'- 5 10 15 20 25 30 35 40 45 -3'
SupFam_25
1.9 log10 RPM -0.1
3'- 23 22 21 20 19 18 17 16 15 14 13 12 11 10 9 8 7 6 5 4 3 2 1 -5'
Cl_ccm_2177 -UUCUACCUUACUUCUACGUCUC
Cl_cpe-2015_150 AUUCUACCUUACUUCUACGUCUC
Cl_cpe-2015_15012 ---CUACCUUACUUCUACGUCUC
Cl_cpe-2017_186 AUUCUACCUUACUUCUACGUCUC
Cl_cpe-2017_17400 ---CUACCUUACUUCUACGUCUC
Cl_cgr-pm_5848 -UUCUGCCUUACUUUUACGUUU-
Cl_cgr-pm_10735 -ACGUACCUUCCUUUUUCCUUU-
Cl_cgr-pm_41013 -UCCUACCUUACUUUUACCUCU-
Cl_cgr-mass_1328 -UUCUACCCUACUUCUACGUCUC
Cl_cgr-mass_22564 -UUCUGCCUUACUUUUACGUUU-
Perc. ID
>80%
>60%
>40%
<40%

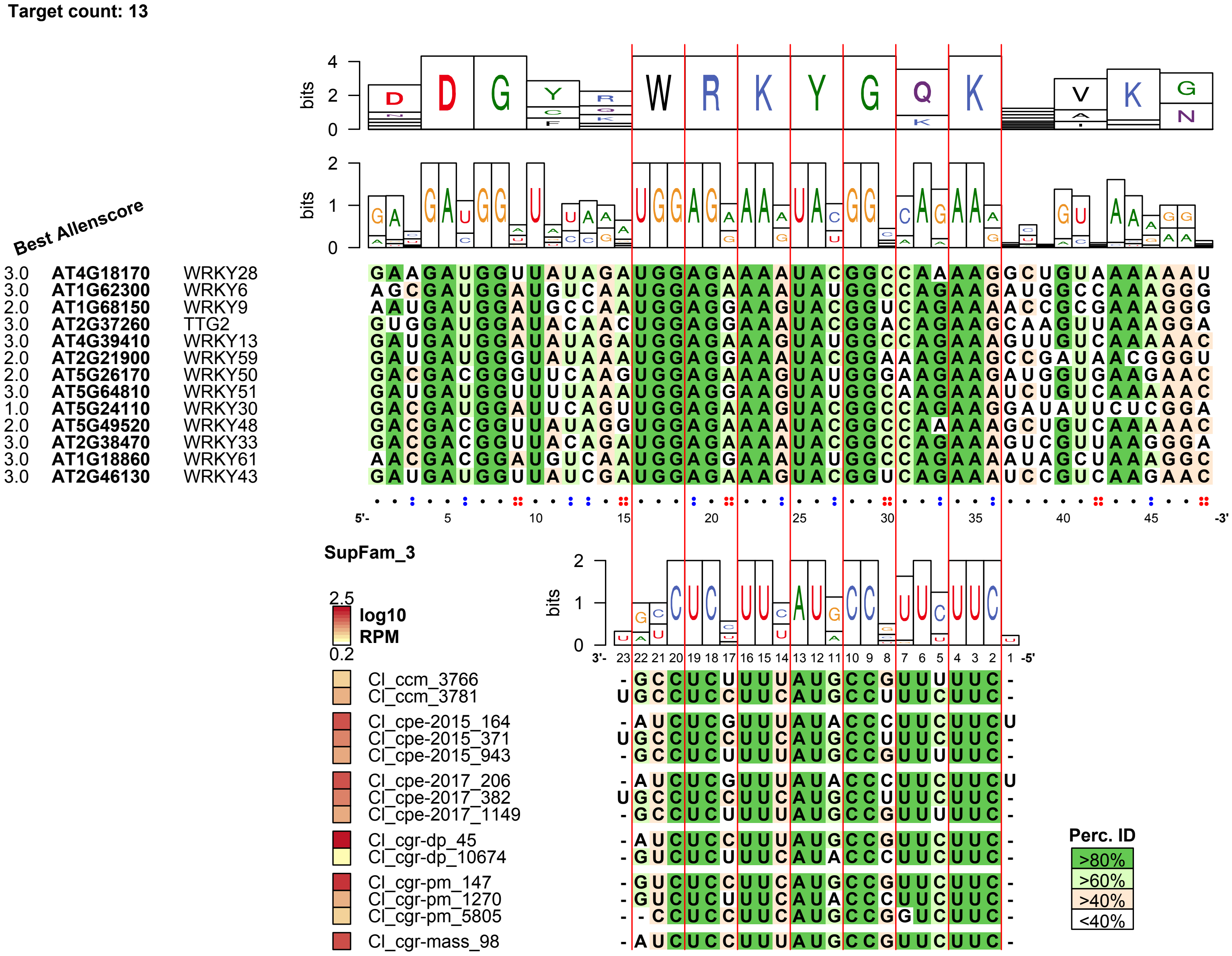

Target count: 13
Best Allenscore
3.0 AT4G18170 WRKY28
3.0 AT1G62300 WRKY6
2.0 AT1G68150 WRKY9
3.0 AT2G37260 TTG2
3.0 AT4G39410 WRKY13
2.0 AT2G21900 WRKY59
2.0 AT5G26170 WRKY50
3.0 AT5G64810 WRKY51
1.0 AT5G24110 WRKY30
2.0 AT5G49520 WRKY48
3.0 AT2G38470 WRKY33
3.0 AT1G18860 WRKY61
3.0 AT2G46130 WRKY43
bits
SupFam_3
log10 RPM
Cl_ccm_3766
Cl_ccm_3781
Cl_cpe-2015_164
Cl_cpe-2015_371
Cl_cpe-2015_943
Cl_cpe-2017_206
Cl_cpe-2017_382
Cl_cpe-2017_1149
Cl_cgr-dp_45
Cl_cgr-dp_10674
Cl_cgr-pm_147
Cl_cgr-pm_1270
Cl_cgr-pm_5805
Cl_cgr-mass_98
Perc. ID
>80%
>60%
>40%
<40%

**Target count: 13**

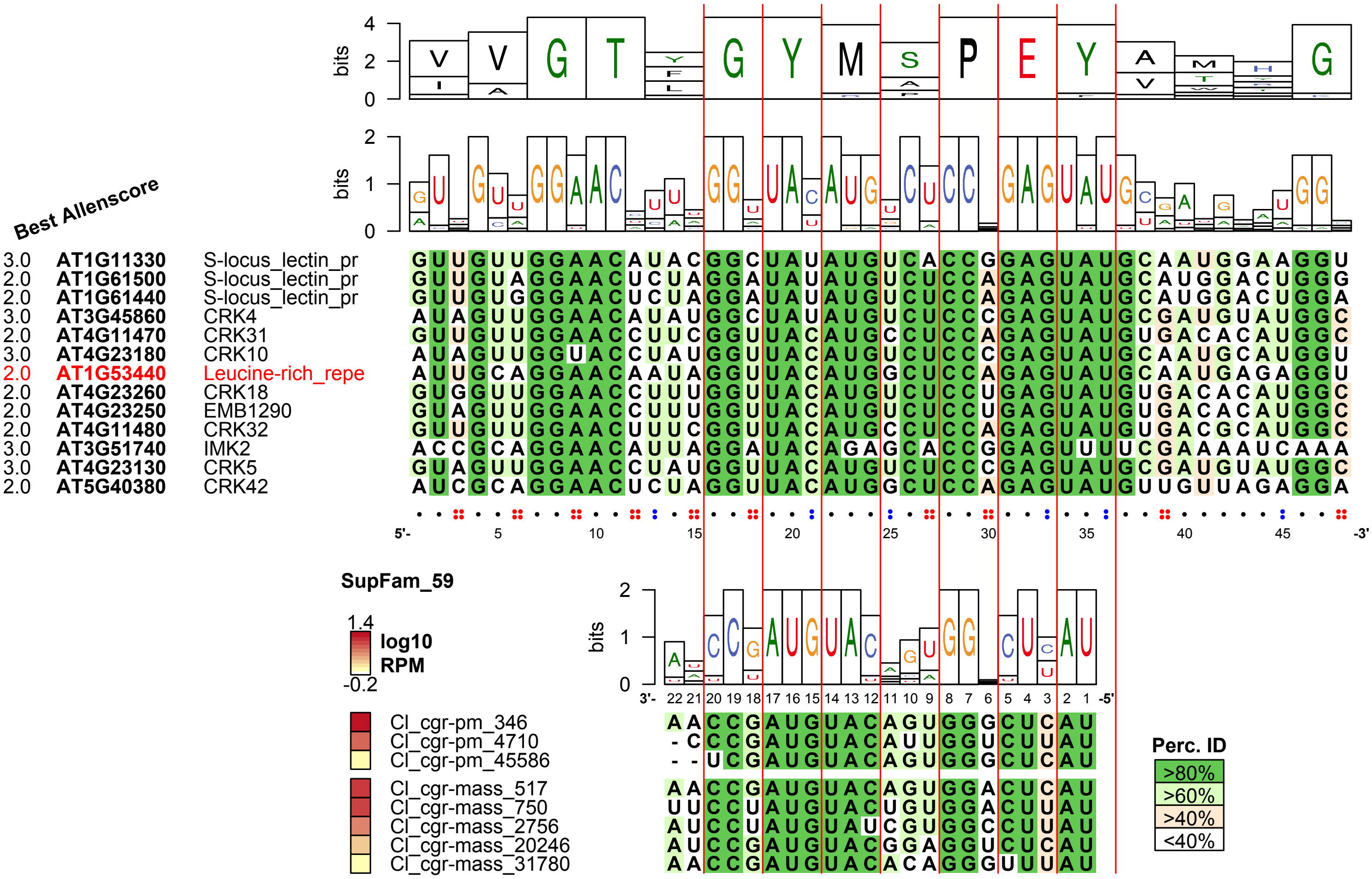

**Target count: 21**

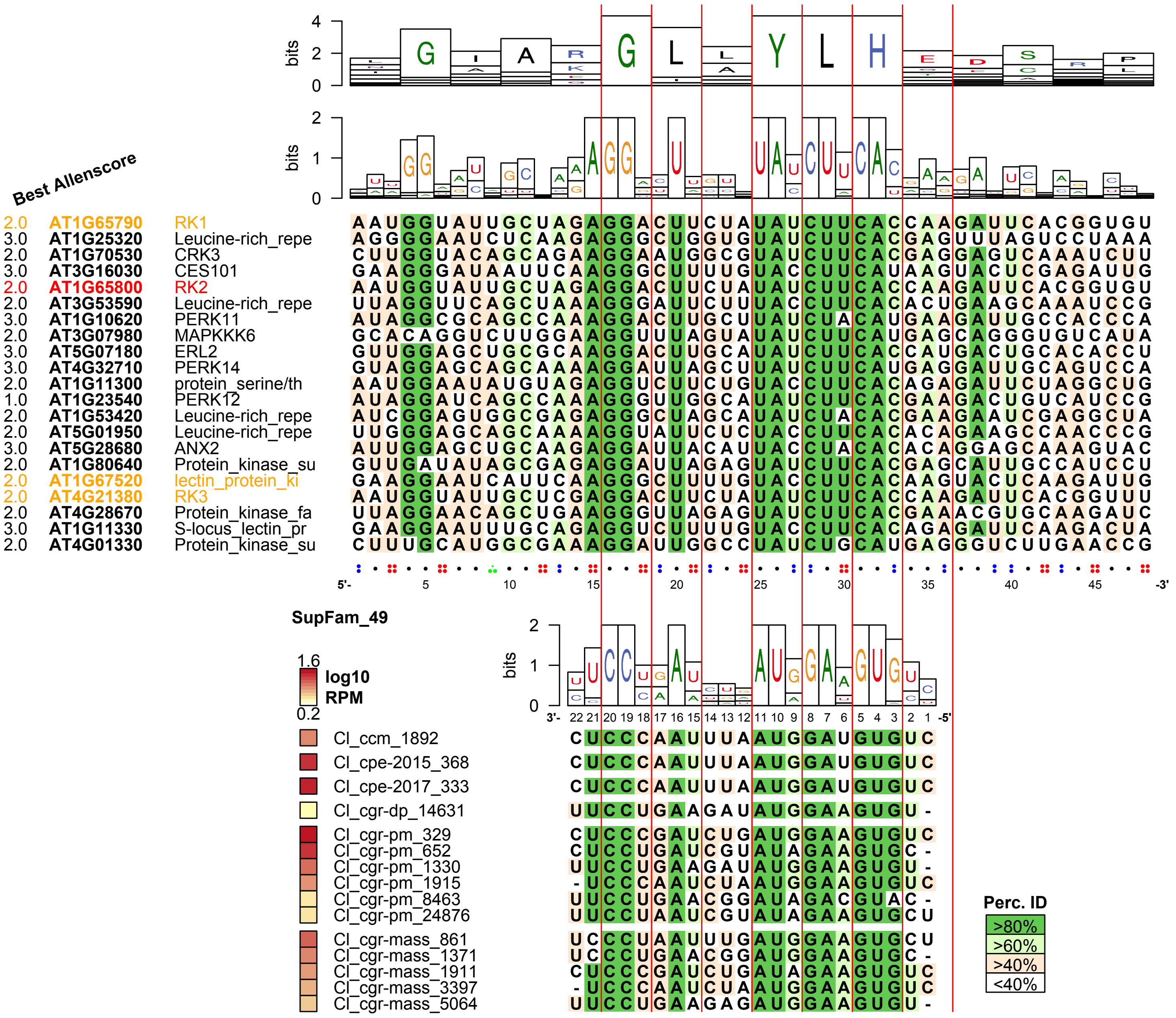

**Target count: 10**

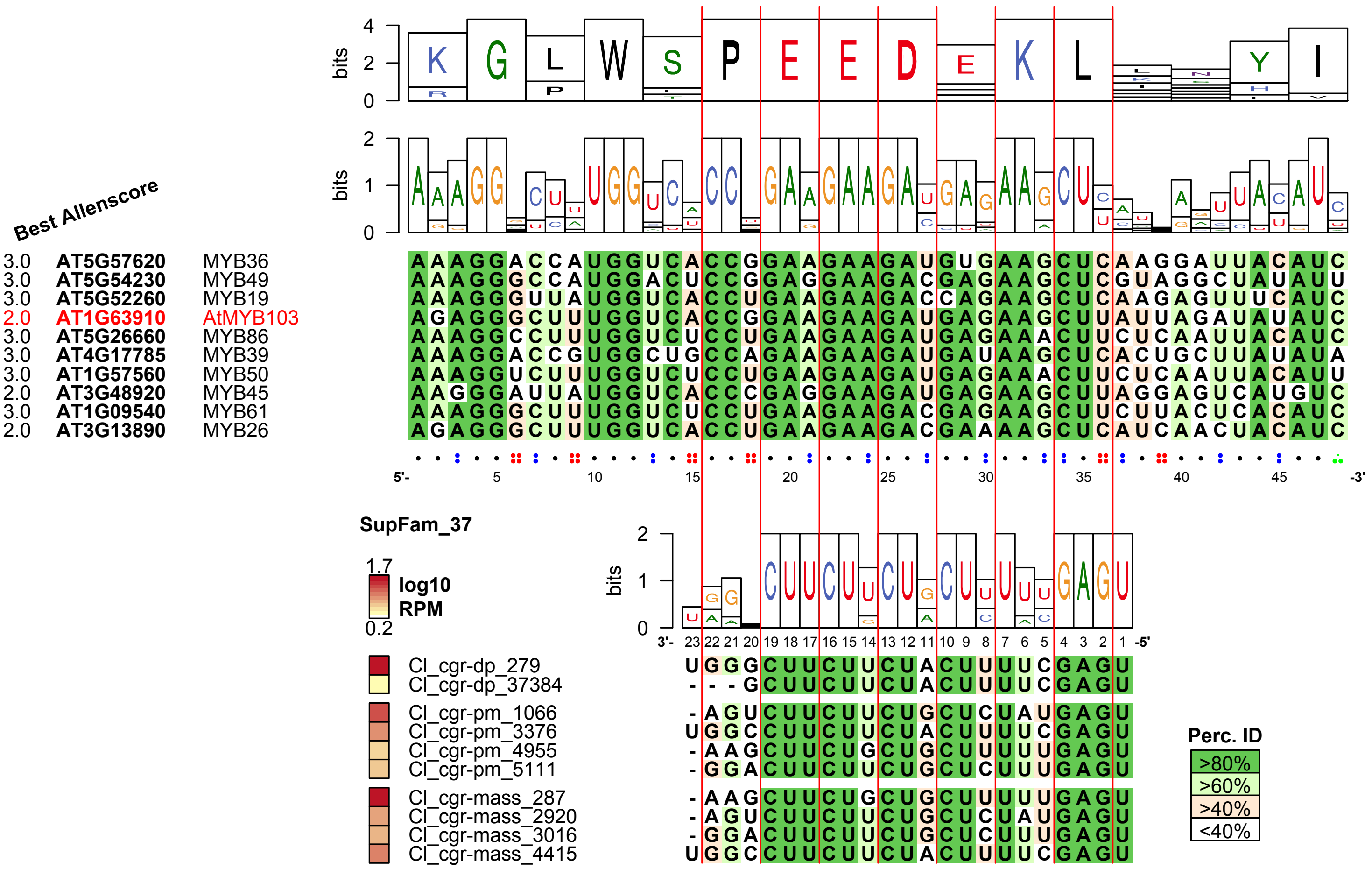

Target count: 5

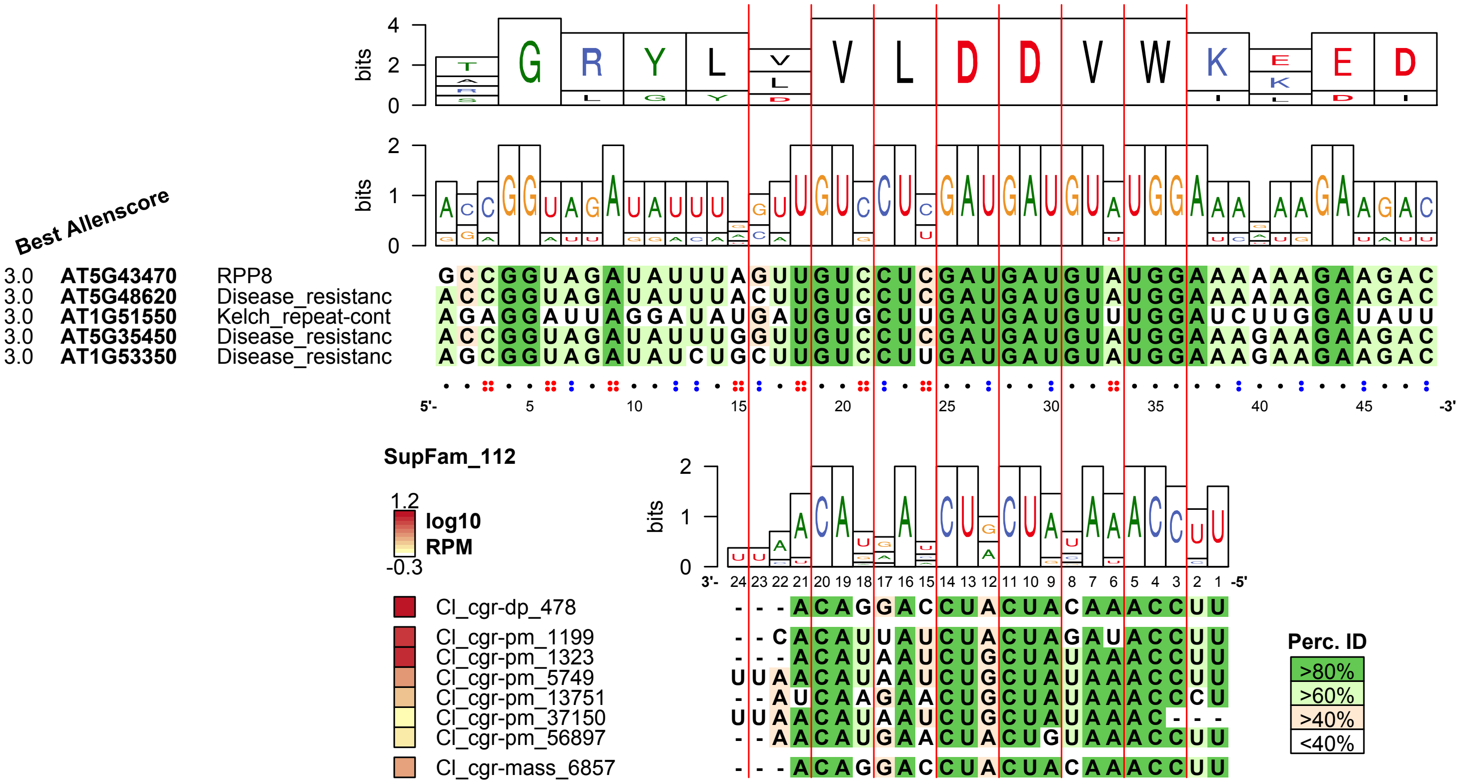

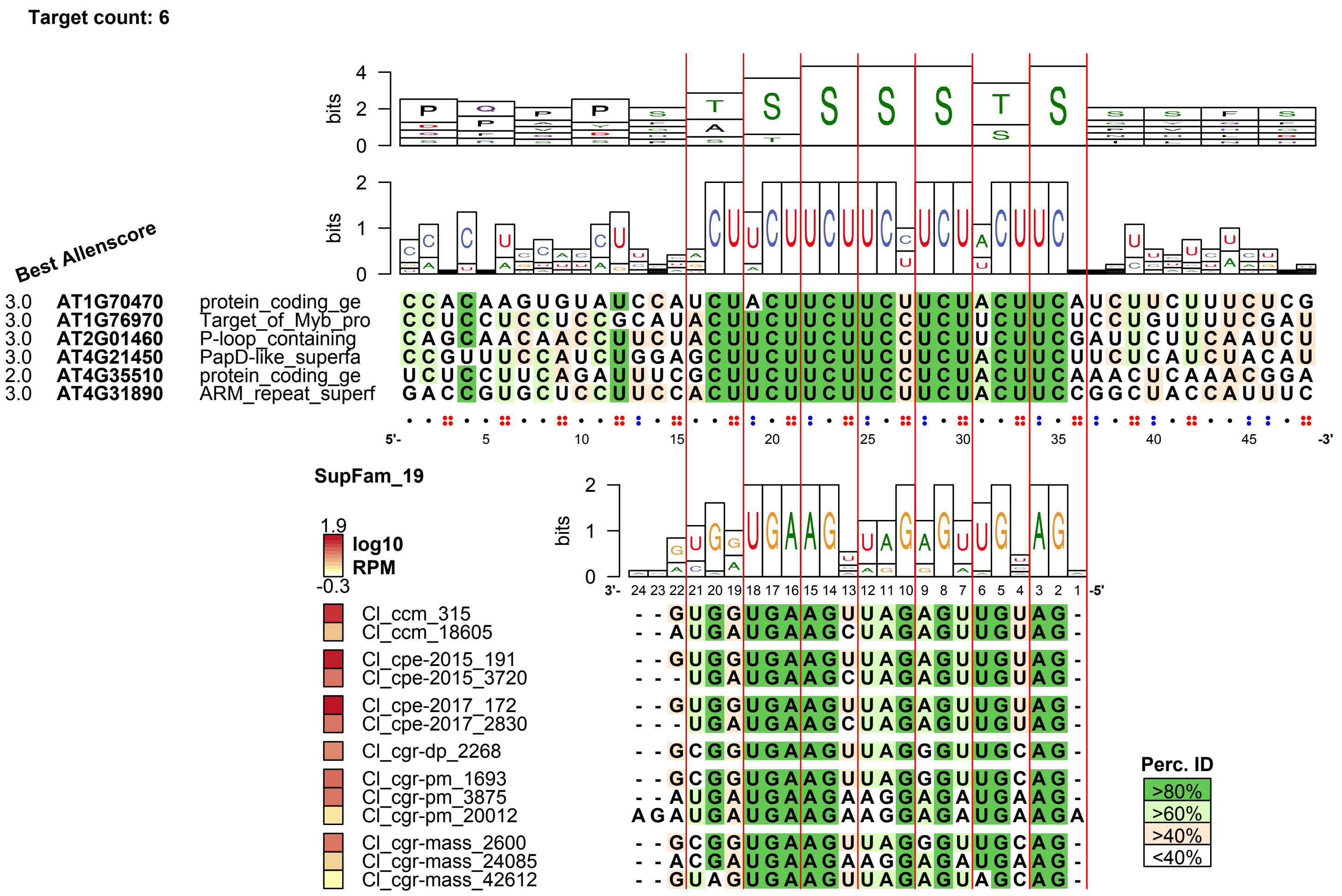

Target count: 6
Best Allenscore
bits
3.0 AT1G70470 protein_coding_ge
3.0 AT1G76970 Target_of_Myb_pro
3.0 AT2G01460 P-loop_containing
3.0 AT4G21450 PapD-like_superfa
2.0 AT4G35510 protein_coding_ge
3.0 AT4G31890 ARM_repeat_superf
5'-
-3'
SupFam_19
1.9
log10
RPM
-0.3
Cl_ccm_315
Cl_ccm_18605
Cl_cpe-2015_191
Cl_cpe-2015_3720
Cl_cpe-2017_172
Cl_cpe-2017_2830
Cl_cgr-dp_2268
Cl_cgr-pm_1693
Cl_cgr-pm_3875
Cl_cgr-pm_20012
Cl_cgr-mass_2600
Cl_cgr-mass_24085
Cl_cgr-mass_42612
3'-
-5'
Perc. ID
>80%
>60%
>40%
<40%

**Table S1. List of all libraries and tissues prepared in this study**

All libraries are available under the SRA BioProject: PRJNA543296

| SRA accession | file_name | library_id | tissue_id | experiment | tissue | host | parasite_species | parasite isolate | parasite_origin | biological replicate |
|---|---|---|---|---|---|---|---|---|---|---|
| SRR9216091 | 1_sRNA_ccm_IN_rep1.fastq.gz | lib1_sRNA_ccm_IN_rep1 | tissue1_ccm_IN_rep1 | sRNA-seq | (IN) interface | *A. thaliana* (Col-0) | *C. campestris* | *ccm* | germinated seedling | rep_1 |
| SRR9216090 | 2_sRNA_ccm_PS_rep1.fastq.gz | lib2_sRNA_ccm_PS_rep1 | tissue2_ccm_PS_rep1 | sRNA-seq | (PS) parasite stem | *A. thaliana* (Col-0) | *C. campestris* | *ccm* | germinated seedling | rep_1 |
| SRR9216089 | 3_sRNA_ccm_IN_rep2.fastq.gz | lib3_sRNA_ccm_IN_rep2 | tissue3_ccm_IN_rep2 | sRNA-seq | (IN) interface | *A. thaliana* (Col-0) | *C. campestris* | *ccm* | germinated seedling | rep_2 |
| SRR9216088 | 4_sRNA_ccm_PS_rep2.fastq.gz | lib4_sRNA_ccm_PS_rep2 | tissue4_ccm_PS_rep2 | sRNA-seq | (PS) parasite stem | *A. thaliana* (Col-0) | *C. campestris* | *ccm* | germinated seedling | rep_2 |
| SRR9216087 | 5_sRNA_ccm_IN_rep3.fastq.gz | lib5_sRNA_ccm_IN_rep3 | tissue5_ccm_IN_rep3 | sRNA-seq | (IN) interface | *A. thaliana* (Col-0) | *C. campestris* | *ccm* | germinated seedling | rep_3 |
| SRR9216086 | 6_sRNA_ccm_PS_rep3.fastq.gz | lib6_sRNA_ccm_PS_rep3 | tissue6_ccm_PS_rep3 | sRNA-seq | (PS) parasite stem | *A. thaliana* (Col-0) | *C. campestris* | *ccm* | germinated seedling | rep_3 |
| SRR9216085 | 7_sRNA_cgr-dp_IN_rep1.fastq.gz | lib7_sRNA_cgr-dp_IN_rep1 | tissue7_cgr-dp_IN_rep1 | sRNA-seq | (IN) interface | *A. thaliana* (Col-0) | *C. gronovii* | *cgr-dp* | germinated seedling | rep_1 |
| SRR9216084 | 8_sRNA_cgr-dp_PS_rep1.fastq.gz | lib8_sRNA_cgr-dp_PS_rep1 | tissue8_cgr-dp_PS_rep1 | sRNA-seq | (PS) parasite stem | *A. thaliana* (Col-0) | *C. gronovii* | *cgr-dp* | germinated seedling | rep_1 |
| SRR9216092 | 9_sRNA_cgr-dp_IN_rep2.fastq.gz | lib9_sRNA_cgr-dp_IN_rep2 | tissue9_cgr-dp_IN_rep2 | sRNA-seq | (IN) interface | *A. thaliana* (Col-0) | *C. gronovii* | *cgr-dp* | germinated seedling | rep_2 |
| SRR9216112 | 10_sRNA_cgr-dp_PS_rep2.fastq.gz | lib10_sRNA_cgr-dp_PS_rep2 | tissue10_cgr-dp_PS_rep2 | sRNA-seq | (PS) parasite stem | *A. thaliana* (Col-0) | *C. gronovii* | *cgr-dp* | germinated seedling | rep_2 |
| SRR9216081 | 11_sRNA_cgr-dp_IN_rep3.fastq.gz | lib11_sRNA_cgr-dp_IN_rep3 | tissue11_cgr-dp_IN_rep3 | sRNA-seq | (IN) interface | *A. thaliana* (Col-0) | *C. gronovii* | *cgr-dp* | germinated seedling | rep_3 |
| SRR9216080 | 12_sRNA_cgr-dp_PS_rep3.fastq.gz | lib12_sRNA_cgr-dp_PS_rep3 | tissue12_cgr-dp_PS_rep3 | sRNA-seq | (PS) parasite stem | *A. thaliana* (Col-0) | *C. gronovii* | *cgr-dp* | germinated seedling | rep_3 |
| SRR9216083 | 13_sRNA_cgr-pm_IN_rep1.fastq.gz | lib13_sRNA_cgr-pm_IN_rep1 | tissue13_cgr-pm_IN_rep1 | sRNA-seq | (IN) interface | *A. thaliana* (Col-0) | *C. gronovii* | *cgr-pm* | germinated seedling | rep_1 |
| SRR9216082 | 14_sRNA_cgr-pm_PS_rep1.fastq.gz | lib14_sRNA_cgr-pm_PS_rep1 | tissue14_cgr-pm_PS_rep1 | sRNA-seq | (PS) parasite stem | *A. thaliana* (Col-0) | *C. gronovii* | *cgr-pm* | germinated seedling | rep_1 |
| SRR9216077 | 15_sRNA_cgr-pm_IN_rep2.fastq.gz | lib15_sRNA_cgr-pm_IN_rep2 | tissue15_cgr-pm_IN_rep2 | sRNA-seq | (IN) interface | *A. thaliana* (Col-0) | *C. gronovii* | *cgr-pm* | germinated seedling | rep_2 |
| SRR9216076 | 16_sRNA_cgr-pm_PS_rep2.fastq.gz | lib16_sRNA_cgr-pm_PS_rep2 | tissue16_cgr-pm_PS_rep2 | sRNA-seq | (PS) parasite stem | *A. thaliana* (Col-0) | *C. gronovii* | *cgr-pm* | germinated seedling | rep_2 |
| SRR9216079 | 17_sRNA_cgr-pm_IN_rep3.fastq.gz | lib17_sRNA_cgr-pm_IN_rep3 | tissue17_cgr-pm_IN_rep3 | sRNA-seq | (IN) interface | *A. thaliana* (Col-0) | *C. gronovii* | *cgr-pm* | germinated seedling | rep_3 |
| SRR9216078 | 18_sRNA_cgr-pm_PS_rep3.fastq.gz | lib18_sRNA_cgr-pm_PS_rep3 | tissue18_cgr-pm_PS_rep3 | sRNA-seq | (PS) parasite stem | *A. thaliana* (Col-0) | *C. gronovii* | *cgr-pm* | germinated seedling | rep_3 |
| SRR9216075 | 19_sRNA_cgr-mass_IN_rep1.fastq.gz | lib19_sRNA_cgr-mass_IN_rep1 | tissue19_cgr-mass_IN_rep1 | sRNA-seq | (IN) interface | *A. thaliana* (Col-0) | *C. gronovii* | *cgr-mass* | germinated seedling | rep_1 |
| SRR9216074 | 20_sRNA_cgr-mass_PS_rep1.fastq.gz | lib20_sRNA_cgr-mass_PS_rep1 | tissue20_cgr-mass_PS_rep1 | sRNA-seq | (PS) parasite stem | *A. thaliana* (Col-0) | *C. gronovii* | *cgr-mass* | germinated seedling | rep_1 |
| SRR9216057 | 21_sRNA_cgr-mass_IN_rep2.fastq.gz | lib21_sRNA_cgr-mass_IN_rep2 | tissue21_cgr-mass_IN_rep2 | sRNA-seq | (IN) interface | *A. thaliana* (Col-0) | *C. gronovii* | *cgr-mass* | germinated seedling | rep_2 |
| SRR9216058 | 22_sRNA_cgr-mass_PS_rep2.fastq.gz | lib22_sRNA_cgr-mass_PS_rep2 | tissue22_cgr-mass_PS_rep2 | sRNA-seq | (PS) parasite stem | *A. thaliana* (Col-0) | *C. gronovii* | *cgr-mass* | germinated seedling | rep_2 |
| SRR9216059 | 23_sRNA_cgr-mass_IN_rep3.fastq.gz | lib23_sRNA_cgr-mass_IN_rep3 | tissue23_cgr-mass_IN_rep3 | sRNA-seq | (IN) interface | *A. thaliana* (Col-0) | *C. gronovii* | *cgr-mass* | germinated seedling | rep_3 |
| SRR9216060 | 24_sRNA_cgr-mass_PS_rep3.fastq.gz | lib24_sRNA_cgr-mass_PS_rep3 | tissue24_cgr-mass_PS_rep3 | sRNA-seq | (PS) parasite stem | *A. thaliana* (Col-0) | *C. gronovii* | *cgr-mass* | germinated seedling | rep_3 |
| SRR9216061 | 25_sRNA_cpe-2015_IN_rep1.fastq.gz | lib25_sRNA_cpe-2015_IN_rep1 | tissue25_cpe-2015_IN_rep1 | sRNA-seq | (IN) interface | *A. thaliana* (Col-0) | *C. pentagona* | *cpe-2015* | germinated seedling | rep_1 |
| SRR9216062 | 26_sRNA_cpe-2015_PS_rep1.fastq.gz | lib26_sRNA_cpe-2015_PS_rep1 | tissue26_cpe-2015_PS_rep1 | sRNA-seq | (PS) parasite stem | *A. thaliana* (Col-0) | *C. pentagona* | *cpe-2015* | germinated seedling | rep_1 |
| SRR9216063 | 27_sRNA_cpe-2015_IN_rep2.fastq.gz | lib27_sRNA_cpe-2015_IN_rep2 | tissue27_cpe-2015_IN_rep2 | sRNA-seq | (IN) interface | *A. thaliana* (Col-0) | *C. pentagona* | *cpe-2015* | germinated seedling | rep_2 |
| SRR9216064 | 28_sRNA_cpe-2015_PS_rep2.fastq.gz | lib28_sRNA_cpe-2015_PS_rep2 | tissue28_cpe-2015_PS_rep2 | sRNA-seq | (PS) parasite stem | *A. thaliana* (Col-0) | *C. pentagona* | *cpe-2015* | germinated seedling | rep_2 |
| SRR9216065 | 29_sRNA_cpe-2015_IN_rep3.fastq.gz | lib29_sRNA_cpe-2015_IN_rep3 | tissue29_cpe-2015_IN_rep3 | sRNA-seq | (IN) interface | *A. thaliana* (Col-0) | *C. pentagona* | *cpe-2015* | germinated seedling | rep_3 |
| SRR9216101 | 30_sRNA_cpe-2015_PS_rep3.fastq.gz | lib30_sRNA_cpe-2015_PS_rep3 | tissue30_cpe-2015_PS_rep3 | sRNA-seq | (PS) parasite stem | *A. thaliana* (Col-0) | *C. pentagona* | *cpe-2015* | germinated seedling | rep_3 |
| SRR9216046 | 31_sRNA_cpe-2017_IN_rep1.fastq.gz | lib31_sRNA_cpe-2017_IN_rep1 | tissue31_cpe-2017_IN_rep1 | sRNA-seq | (IN) interface | *A. thaliana* (Col-0) | *C. pentagona* | *cpe-2017* | germinated seedling | rep_1 |
| SRR9216045 | 32_sRNA_cpe-2017_PS_rep1.fastq.gz | lib32_sRNA_cpe-2017_PS_rep1 | tissue32_cpe-2017_PS_rep1 | sRNA-seq | (PS) parasite stem | *A. thaliana* (Col-0) | *C. pentagona* | *cpe-2017* | germinated seedling | rep_1 |
| SRR9216044 | 33_sRNA_cpe-2017_IN_rep2.fastq.gz | lib33_sRNA_cpe-2017_IN_rep2 | tissue33_cpe-2017_IN_rep2 | sRNA-seq | (IN) interface | *A. thaliana* (Col-0) | *C. pentagona* | *cpe-2017* | germinated seedling | rep_2 |
| SRR9216043 | 34_sRNA_cpe-2017_PS_rep2.fastq.gz | lib34_sRNA_cpe-2017_PS_rep2 | tissue34_cpe-2017_PS_rep2 | sRNA-seq | (PS) parasite stem | *A. thaliana* (Col-0) | *C. pentagona* | *cpe-2017* | germinated seedling | rep_2 |
| SRR9216042 | 35_sRNA_cpe-2017_IN_rep3.fastq.gz | lib35_sRNA_cpe-2017_IN_rep3 | tissue35_cpe-2017_IN_rep3 | sRNA-seq | (IN) interface | *A. thaliana* (Col-0) | *C. pentagona* | *cpe-2017* | germinated seedling | rep_3 |
| SRR9216041 | 36_sRNA_cpe-2017_PS_rep3.fastq.gz | lib36_sRNA_cpe-2017_PS_rep3 | tissue36_cpe-2017_PS_rep3 | sRNA-seq | (PS) parasite stem | *A. thaliana* (Col-0) | *C. pentagona* | *cpe-2017* | germinated seedling | rep_3 |
| SRR9216040 | 37_sRNA_cin_IN_rep1.fastq.gz | lib37_sRNA_cin_IN_rep1 | tissue37_cin_IN_rep1 | sRNA-seq | (IN) interface | *A. thaliana* (Col-0) | *C. indecora* | *cin* | ~5 cm tendril tip from adult plant | rep_1 |
| SRR9216039 | 38_sRNA_cin_PS_rep1.fastq.gz | lib38_sRNA_cin_PS_rep1 | tissue38_cin_PS_rep1 | sRNA-seq | (PS) parasite stem | *A. thaliana* (Col-0) | *C. indecora* | *cin* | ~5 cm tendril tip from adult plant | rep_1 |
| SRR9216048 | 39_sRNA_cin_IN_rep2.fastq.gz | lib39_sRNA_cin_IN_rep2 | tissue39_cin_IN_rep2 | sRNA-seq | (IN) interface | *A. thaliana* (Col-0) | *C. indecora* | *cin* | ~5 cm tendril tip from adult plant | rep_2 |
| SRR9216047 | 40_sRNA_cin_PS_rep2.fastq.gz | lib40_sRNA_cin_PS_rep2 | tissue40_cin_PS_rep2 | sRNA-seq | (PS) parasite stem | *A. thaliana* (Col-0) | *C. indecora* | *cin* | ~5 cm tendril tip from adult plant | rep_2 |
| SRR9216095 | 41_sRNA_cin_IN_rep3.fastq.gz | lib41_sRNA_cin_IN_rep3 | tissue41_cin_IN_rep3 | sRNA-seq | (IN) interface | *A. thaliana* (Col-0) | *C. indecora* | *cin* | ~5 cm tendril tip from adult plant | rep_3 |
| SRR9216051 | 42_sRNA_cin_PS_rep3.fastq.gz | lib42_sRNA_cin_PS_rep3 | tissue42_cin_PS_rep3 | sRNA-seq | (PS) parasite stem | *A. thaliana* (Col-0) | *C. indecora* | *cin* | ~5 cm tendril tip from adult plant | rep_3 |
| SRR9216049 | 43_sRNA_ath_CS_rep1.fastq.gz | lib43_sRNA_ath_CS_rep1 | tissue43_ath_CS_rep1 | sRNA-seq | (CS) control, unparasitized stem | *A. thaliana* (Col-0) | | | | rep_1 |
| SRR9216050 | 44_sRNA_ath_CS_rep2.fastq.gz | lib44_sRNA_ath_CS_rep2 | tissue44_ath_CS_rep2 | sRNA-seq | (CS) control, unparasitized stem | *A. thaliana* (Col-0) | | | | rep_2 |
| SRR9216053 | 45_sRNA_ath_CS_rep3.fastq.gz | lib45_sRNA_ath_CS_rep3 | tissue45_ath_CS_rep3 | sRNA-seq | (CS) control, unparasitized stem | *A. thaliana* (Col-0) | | | | rep_3 |
| SRR9216054 | 46_PARE_ccm_HIN_rep1.fastq.gz | lib46_PARE_ccm_HIN_rep1 | tissue46_ccm_HIN_rep1 | NanoPARE | (HIN) host stem from interface | *A. thaliana* (*xrn4*) | *C. campestris* | *ccm* | germinated seedling | rep_1 |
| SRR9216052 | 47_PARE_ccm_HIN_rep2.fastq.gz | lib47_PARE_ccm_HIN_rep2 | tissue47_ccm_HIN_rep2 | NanoPARE | (HIN) host stem from interface | *A. thaliana* (*xrn4*) | *C. campestris* | *ccm* | germinated seedling | rep_2 |
| SRR9216098 | 48_PARE_ccm_HIN_rep3.fastq.gz | lib48_PARE_ccm_HIN_rep3 | tissue48_ccm_HIN_rep3 | NanoPARE | (HIN) host stem from interface | *A. thaliana* (*xrn4*) | *C. campestris* | *ccm* | germinated seedling | rep_3 |
| SRR9216055 | 49_PARE_cgr-dp_HIN_rep1.fastq.gz | lib49_PARE_cgr-dp_HIN_rep1 | tissue49_cgr-dp_HIN_rep1 | NanoPARE | (HIN) host stem from interface | *A. thaliana* (*xrn4*) | *C. gronovii* | *cgr-dp* | germinated seedling | rep_1 |
| SRR9216056 | 50_PARE_cgr-dp_HIN_rep2.fastq.gz | lib50_PARE_cgr-dp_HIN_rep2 | tissue50_cgr-dp_HIN_rep2 | NanoPARE | (HIN) host stem from interface | *A. thaliana* (*xrn4*) | *C. gronovii* | *cgr-dp* | germinated seedling | rep_2 |
| SRR9216030 | 51_PARE_cgr-dp_HIN_rep3.fastq.gz | lib51_PARE_cgr-dp_HIN_rep3 | tissue51_cgr-dp_HIN_rep3 | NanoPARE | (HIN) host stem from interface | *A. thaliana* (*xrn4*) | *C. gronovii* | *cgr-dp* | germinated seedling | rep_3 |
| SRR9216029 | 52_PARE_cpe-2015_HIN_rep1.fastq.gz | lib52_PARE_cpe-2015_HIN_rep1 | tissue52_cpe-2015_HIN_rep1 | NanoPARE | (HIN) host stem from interface | *A. thaliana* (*xrn4*) | *C. pentagona* | *cpe-2015* | germinated seedling | rep_1 |
| SRR9216032 | 53_PARE_cpe-2015_HIN_rep2.fastq.gz | lib53_PARE_cpe-2015_HIN_rep2 | tissue53_cpe-2015_HIN_rep2 | NanoPARE | (HIN) host stem from interface | *A. thaliana* (*xrn4*) | *C. pentagona* | *cpe-2015* | germinated seedling | rep_2 |
| SRR9216031 | 54_PARE_cpe-2015_HIN_rep3.fastq.gz | lib54_PARE_cpe-2015_HIN_rep3 | tissue54_cpe-2015_HIN_rep3 | NanoPARE | (HIN) host stem from interface | *A. thaliana* (*xrn4*) | *C. pentagona* | *cpe-2015* | germinated seedling | rep_3 |
| SRR9216034 | 55_PARE_cin_HIN_rep1.fastq.gz | lib55_PARE_cin_HIN_rep1 | tissue55_cin_HIN_rep1 | NanoPARE | (HIN) host stem from interface | *A. thaliana* (*xrn4*) | *C. indecora* | *cin* | ~5 cm tendril tip from adult plant | rep_1 |
| SRR9216033 | 56_PARE_cin_HIN_rep2.fastq.gz | lib56_PARE_cin_HIN_rep2 | tissue56_cin_HIN_rep2 | NanoPARE | (HIN) host stem from interface | *A. thaliana* (*xrn4*) | *C. indecora* | *cin* | ~5 cm tendril tip from adult plant | rep_2 |
| SRR9216036 | 57_PARE_cin_HIN_rep3.fastq.gz | lib57_PARE_cin_HIN_rep3 | tissue57_cin_HIN_rep3 | NanoPARE | (HIN) host stem from interface | *A. thaliana* (*xrn4*) | *C. indecora* | *cin* | ~5 cm tendril tip from adult plant | rep_3 |
| SRR9216035 | 58_PARE_xrn4_CS_rep1.fastq.gz | lib58_PARE_xrn4_CS_rep1 | tissue58_xrn4_CS_rep1 | NanoPARE | (CS) control, unparasitized stem | *A. thaliana* (*xrn4*) | | | | rep_1 |
| SRR9216038 | 59_PARE_xrn4_CS_rep2.fastq.gz | lib59_PARE_xrn4_CS_rep2 | tissue59_xrn4_CS_rep2 | NanoPARE | (CS) control, unparasitized stem | *A. thaliana* (*xrn4*) | | | | rep_2 |
| SRR9216037 | 60_PARE_xrn4_CS_rep3.fastq.gz | lib60_PARE_xrn4_CS_rep3 | tissue60_xrn4_CS_rep3 | NanoPARE | (CS) control, unparasitized stem | *A. thaliana* (*xrn4*) | | | | rep_3 |

| | | | | | | | | | | |
|---|---|---|---|---|---|---|---|---|---|---|
| SRR9216111 | 61_PARE_ccm_IN_rep1.fastq.gz | lib61_PARE_ccm_IN_rep1 | tissue1_ccm_IN_rep1 | NanoPARE | (IN) interface | *A. thaliana* (Col-0) | *C. campestris* | *ccm* | germinated seedling | rep_1 |
| SRR9216113 | 62_PARE_ccm_IN_rep2.fastq.gz | lib62_PARE_ccm_IN_rep2 | tissue3_ccm_IN_rep2 | NanoPARE | (IN) interface | *A. thaliana* (Col-0) | *C. campestris* | *ccm* | germinated seedling | rep_2 |
| SRR9216114 | 63_PARE_ccm_IN_rep3.fastq.gz | lib63_PARE_ccm_IN_rep3 | tissue5_ccm_IN_rep3 | NanoPARE | (IN) interface | *A. thaliana* (Col-0) | *C. campestris* | *ccm* | germinated seedling | rep_3 |
| SRR9216116 | 64_PARE_cgr-dp_IN_rep1.fastq.gz | lib64_PARE_cgr-dp_IN_rep1 | tissue7_cgr-dp_IN_rep1 | NanoPARE | (IN) interface | *A. thaliana* (Col-0) | *C. gronovii* | *cgr-dp* | germinated seedling | rep_1 |
| SRR9216105 | 65_PARE_cgr-dp_IN_rep2.fastq.gz | lib65_PARE_cgr-dp_IN_rep2 | tissue9_cgr-dp_IN_rep2 | NanoPARE | (IN) interface | *A. thaliana* (Col-0) | *C. gronovii* | *cgr-dp* | germinated seedling | rep_2 |
| SRR9216107 | 66_PARE_cgr-dp_IN_rep3.fastq.gz | lib66_PARE_cgr-dp_IN_rep3 | tissue11_cgr-dp_IN_rep3 | NanoPARE | (IN) interface | *A. thaliana* (Col-0) | *C. gronovii* | *cgr-dp* | germinated seedling | rep_3 |
| SRR9216108 | 67_PARE_cpe-2015_IN_rep1.fastq.gz | lib67_PARE_cpe-2015_IN_rep1 | tissue25_cpe-2015_IN_rep1 | NanoPARE | (IN) interface | *A. thaliana* (Col-0) | *C. pentagona* | *cpe-2015* | germinated seedling | rep_1 |
| SRR9216110 | 68_PARE_cpe-2015_IN_rep2.fastq.gz | lib68_PARE_cpe-2015_IN_rep2 | tissue27_cpe-2015_IN_rep2 | NanoPARE | (IN) interface | *A. thaliana* (Col-0) | *C. pentagona* | *cpe-2015* | germinated seedling | rep_2 |
| SRR9216117 | 69_PARE_cpe-2015_IN_rep3.fastq.gz | lib69_PARE_cpe-2015_IN_rep3 | tissue29_cpe-2015_IN_rep3 | NanoPARE | (IN) interface | *A. thaliana* (Col-0) | *C. pentagona* | *cpe-2015* | germinated seedling | rep_3 |
| SRR9216118 | 70_PARE_cin_IN_rep1.fastq.gz | lib70_PARE_cin_IN_rep1 | tissue37_cin_IN_rep1 | NanoPARE | (IN) interface | *A. thaliana* (Col-0) | *C. indecora* | *cin* | ~5 cm tendril tip from adult plant | rep_1 |
| SRR9216100 | 71_PARE_cin_IN_rep2.fastq.gz | lib71_PARE_cin_IN_rep2 | tissue39_cin_IN_rep2 | NanoPARE | (IN) interface | *A. thaliana* (Col-0) | *C. indecora* | *cin* | ~5 cm tendril tip from adult plant | rep_2 |
| SRR9216099 | 72_PARE_cin_IN_rep3.fastq.gz | lib72_PARE_cin_IN_rep3 | tissue41_cin_IN_rep3 | NanoPARE | (IN) interface | *A. thaliana* (Col-0) | *C. indecora* | *cin* | ~5 cm tendril tip from adult plant | rep_3 |
| SRR9216097 | 73_PARE_ath_CS_rep1.fastq.gz | lib73_PARE_ath_CS_rep1 | tissue43_ath_CS_rep1 | NanoPARE | (CS) control, unparasitized stem | *A. thaliana* (Col-0) | | | | rep_1 |
| SRR9216096 | 74_PARE_ath_CS_rep2.fastq.gz | lib74_PARE_ath_CS_rep2 | tissue44_ath_CS_rep2 | NanoPARE | (CS) control, unparasitized stem | *A. thaliana* (Col-0) | | | | rep_2 |
| SRR9216104 | 75_PARE_ath_CS_rep3.fastq.gz | lib75_PARE_ath_CS_rep3 | tissue45_ath_CS_rep3 | NanoPARE | (CS) control, unparasitized stem | *A. thaliana* (Col-0) | | | | rep_3 |
| SRR9216103 | 76_mRNA_ccm_IN_rep1.fastq.gz | lib76_mRNA_ccm_IN_rep1 | tissue1_ccm_IN_rep1 | mRNA-seq | (IN) interface | *A. thaliana* (Col-0) | *C. campestris* | *ccm* | germinated seedling | rep_1 |
| SRR9216115 | 77_mRNA_ccm_IN_rep2.fastq.gz | lib77_mRNA_ccm_IN_rep2 | tissue3_ccm_IN_rep2 | mRNA-seq | (IN) interface | *A. thaliana* (Col-0) | *C. campestris* | *ccm* | germinated seedling | rep_2 |
| SRR9216102 | 78_mRNA_ccm_IN_rep3.fastq.gz | lib78_mRNA_ccm_IN_rep3 | tissue5_ccm_IN_rep3 | mRNA-seq | (IN) interface | *A. thaliana* (Col-0) | *C. campestris* | *ccm* | germinated seedling | rep_3 |
| SRR9216094 | 79_mRNA_cgr-dp_IN_rep1.fastq.gz | lib79_mRNA_cgr-dp_IN_rep1 | tissue7_cgr-dp_IN_rep1 | mRNA-seq | (IN) interface | *A. thaliana* (Col-0) | *C. gronovii* | *cgr-dp* | germinated seedling | rep_1 |
| SRR9216093 | 80_mRNA_cgr-dp_IN_rep2.fastq.gz | lib80_mRNA_cgr-dp_IN_rep2 | tissue9_cgr-dp_IN_rep2 | mRNA-seq | (IN) interface | *A. thaliana* (Col-0) | *C. gronovii* | *cgr-dp* | germinated seedling | rep_2 |
| SRR9216071 | 81_mRNA_cgr-dp_IN_rep3.fastq.gz | lib81_mRNA_cgr-dp_IN_rep3 | tissue11_cgr-dp_IN_rep3 | mRNA-seq | (IN) interface | *A. thaliana* (Col-0) | *C. gronovii* | *cgr-dp* | germinated seedling | rep_3 |
| SRR9216109 | 82_mRNA_cpe-2015_IN_rep1.fastq.gz | lib82_mRNA_cpe-2015_IN_rep1 | tissue25_cpe-2015_IN_rep1 | mRNA-seq | (IN) interface | *A. thaliana* (Col-0) | *C. pentagona* | *cpe-2015* | germinated seedling | rep_1 |
| SRR9216069 | 83_mRNA_cpe-2015_IN_rep2.fastq.gz | lib83_mRNA_cpe-2015_IN_rep2 | tissue27_cpe-2015_IN_rep2 | mRNA-seq | (IN) interface | *A. thaliana* (Col-0) | *C. pentagona* | *cpe-2015* | germinated seedling | rep_2 |
| SRR9216070 | 84_mRNA_cpe-2015_IN_rep3.fastq.gz | lib84_mRNA_cpe-2015_IN_rep3 | tissue29_cpe-2015_IN_rep3 | mRNA-seq | (IN) interface | *A. thaliana* (Col-0) | *C. pentagona* | *cpe-2015* | germinated seedling | rep_3 |
| SRR9216106 | 85_mRNA_cin_IN_rep1.fastq.gz | lib85_mRNA_cin_IN_rep1 | tissue37_cin_IN_rep1 | mRNA-seq | (IN) interface | *A. thaliana* (Col-0) | *C. indecora* | *cin* | ~5 cm tendril tip from adult plant | rep_1 |
| SRR9216068 | 86_mRNA_cin_IN_rep2.fastq.gz | lib86_mRNA_cin_IN_rep2 | tissue39_cin_IN_rep2 | mRNA-seq | (IN) interface | *A. thaliana* (Col-0) | *C. indecora* | *cin* | ~5 cm tendril tip from adult plant | rep_2 |
| SRR9216066 | 87_mRNA_cin_IN_rep3.fastq.gz | lib87_mRNA_cin_IN_rep3 | tissue41_cin_IN_rep3 | mRNA-seq | (IN) interface | *A. thaliana* (Col-0) | *C. indecora* | *cin* | ~5 cm tendril tip from adult plant | rep_3 |
| SRR9216067 | 88_mRNA_ath_CS_rep1.fastq.gz | lib88_mRNA_ath_CS_rep1 | tissue43_ath_CS_rep1 | mRNA-seq | (CS) control, unparasitized stem | *A. thaliana* (Col-0) | | | | rep_1 |
| SRR9216072 | 89_mRNA_ath_CS_rep2.fastq.gz | lib89_mRNA_ath_CS_rep2 | tissue44_ath_CS_rep2 | mRNA-seq | (CS) control, unparasitized stem | *A. thaliana* (Col-0) | | | | rep_2 |
| SRR9216073 | 90_mRNA_ath_CS_rep3.fastq.gz | lib90_mRNA_ath_CS_rep3 | tissue45_ath_CS_rep3 | mRNA-seq | (CS) control, unparasitized stem | *A. thaliana* (Col-0) | | | | rep_3 |

**Table S2. List of primers used in this study**

| Name | Sequence (5'->3') | Description | Experiment | comment |
|---|---|---|---|---|
| MJA1 | CGAAATCGGTAGACGCTACG | TrnL-F forward | TrnL-F genotyping | From (*32,33*) |
| MJA2 | ATTTGAACTGGTGACACGAG | TrnL-F reverse | TrnL-F genotyping | From (*32,33*) |
| NJ410 | /5Phos/AGATCGGAAGAGCACACGTCT/3SpC3/ | 3' SR Adaptor | sRNA-seq | /5Phos/ indicates 5' phosphorylation; /3SpC3/ indicates C3 Spacer for blocking |
| NJ391 | AGACGTGTGCTCTTCCGATCT | NEB SR RT primer | sRNA-seq | |
| NJ411 | rGrUrUrCrArGrArGrUrUrCrUrArCrArGrUrCrCrGrArCrGrArUrC | 5' SR RNA adaptor | sRNA-seq | rN indicates RNA base |
| NEB1-1 | CAAGCAGAAGACGGCATACGAGATCGTGATGTGACTGGAGTTCAGACGTGTGCTCTTCCGATC*T | NEB barcode Set 1-1 | sRNA-seq | * indicates phosphorothioate bond |
| NEB1-2 | CAAGCAGAAGACGGCATACGAGATACATCGGTGACTGGAGTTCAGACGTGTGCTCTTCCGATC*T | NEB barcode Set 1-2 | sRNA-seq | * indicates phosphorothioate bond |
| NEB1-3 | CAAGCAGAAGACGGCATACGAGATGCCTAAGTGACTGGAGTTCAGACGTGTGCTCTTCCGATC*T | NEB barcode Set 1-3 | sRNA-seq | * indicates phosphorothioate bond |
| NEB1-4 | CAAGCAGAAGACGGCATACGAGATTGGTCAGTGACTGGAGTTCAGACGTGTGCTCTTCCGATC*T | NEB barcode Set 1-4 | sRNA-seq | * indicates phosphorothioate bond |
| NEB1-5 | CAAGCAGAAGACGGCATACGAGATCACTGTGTGACTGGAGTTCAGACGTGTGCTCTTCCGATC*T | NEB barcode Set 1-5 | sRNA-seq | * indicates phosphorothioate bond |
| NEB1-6 | CAAGCAGAAGACGGCATACGAGATATTGGCGTGACTGGAGTTCAGACGTGTGCTCTTCCGATC*T | NEB barcode Set 1-6 | sRNA-seq | * indicates phosphorothioate bond |
| NEB1-7 | CAAGCAGAAGACGGCATACGAGATGATCTGGTGACTGGAGTTCAGACGTGTGCTCTTCCGATC*T | NEB barcode Set 1-7 | sRNA-seq | * indicates phosphorothioate bond |
| NEB1-8 | CAAGCAGAAGACGGCATACGAGATTCAAGTGTGACTGGAGTTCAGACGTGTGCTCTTCCGATC*T | NEB barcode Set 1-8 | sRNA-seq | * indicates phosphorothioate bond |
| NEB1-9 | CAAGCAGAAGACGGCATACGAGATCTGATCGTGACTGGAGTTCAGACGTGTGCTCTTCCGATC*T | NEB barcode Set 1-9 | sRNA-seq | * indicates phosphorothioate bond |
| NEB1-10 | CAAGCAGAAGACGGCATACGAGATAAGCTAGTGACTGGAGTTCAGACGTGTGCTCTTCCGATC*T | NEB barcode Set 1-10 | sRNA-seq | * indicates phosphorothioate bond |
| NEB1-11 | CAAGCAGAAGACGGCATACGAGATGTAGCCGTGACTGGAGTTCAGACGTGTGCTCTTCCGATC*T | NEB barcode Set 1-11 | sRNA-seq | * indicates phosphorothioate bond |
| NEB1-12 | CAAGCAGAAGACGGCATACGAGATTACAAGGTGACTGGAGTTCAGACGTGTGCTCTTCCGATC*T | NEB barcode Set 1-12 | sRNA-seq | * indicates phosphorothioate bond |
| NEB2-1 | CAAGCAGAAGACGGCATACGAGATTGTTGACTGTGACTGGAGTTCAGACGTGTGCTCTTCCGATC*T | NEB barcode Set 2-1 | sRNA-seq | * indicates phosphorothioate bond |
| NEB2-2 | CAAGCAGAAGACGGCATACGAGATACGGAACTGTGACTGGAGTTCAGACGTGTGCTCTTCCGATC*T | NEB barcode Set 2-2 | sRNA-seq | * indicates phosphorothioate bond |
| NEB2-3 | CAAGCAGAAGACGGCATACGAGATTCTGACATGTGACTGGAGTTCAGACGTGTGCTCTTCCGATC*T | NEB barcode Set 2-3 | sRNA-seq | * indicates phosphorothioate bond |
| NEB2-4 | CAAGCAGAAGACGGCATACGAGATGCGGACGGGTGACTGGAGTTCAGACGTGTGCTCTTCCGATC*T | NEB barcode Set 2-4 | sRNA-seq | * indicates phosphorothioate bond |
| NEB2-5 | CAAGCAGAAGACGGCATACGAGATGTGCGGACGTGACTGGAGTTCAGACGTGTGCTCTTCCGATC*T | NEB barcode Set 2-5 | sRNA-seq | * indicates phosphorothioate bond |
| NEB2-6 | CAAGCAGAAGACGGCATACGAGATCGTTTCACGTGACTGGAGTTCAGACGTGTGCTCTTCCGATC*T | NEB barcode Set 2-6 | sRNA-seq | * indicates phosphorothioate bond |
| NEB2-7 | CAAGCAGAAGACGGCATACGAGATAAGGCCACGTGACTGGAGTTCAGACGTGTGCTCTTCCGATC*T | NEB barcode Set 2-7 | sRNA-seq | * indicates phosphorothioate bond |
| NEB2-8 | CAAGCAGAAGACGGCATACGAGATTCCGAAACGTGACTGGAGTTCAGACGTGTGCTCTTCCGATC*T | NEB barcode Set 2-8 | sRNA-seq | * indicates phosphorothioate bond |
| NEB2-9 | CAAGCAGAAGACGGCATACGAGATTACGTACGGTGACTGGAGTTCAGACGTGTGCTCTTCCGATC*T | NEB barcode Set 2-9 | sRNA-seq | * indicates phosphorothioate bond |
| NEB2-10 | CAAGCAGAAGACGGCATACGAGATATCCACTCGTGACTGGAGTTCAGACGTGTGCTCTTCCGATC*T | NEB barcode Set 2-10 | sRNA-seq | * indicates phosphorothioate bond |
| NEB2-11 | CAAGCAGAAGACGGCATACGAGATATATCAGTGTGACTGGAGTTCAGACGTGTGCTCTTCCGATC*T | NEB barcode Set 2-11 | sRNA-seq | * indicates phosphorothioate bond |
| NEB2-12 | CAAGCAGAAGACGGCATACGAGATAAAGGAATGTGACTGGAGTTCAGACGTGTGCTCTTCCGATC*T | NEB barcode Set 2-12 | sRNA-seq | * indicates phosphorothioate bond |
| NEB3-1 | CAAGCAGAAGACGGCATACGAGATCTCTACGTGACTGGAGTTCAGACGTGTGCTCTTCCGATC*T | NEB barcode Set 3-1 | sRNA-seq | * indicates phosphorothioate bond |
| NEB3-2 | CAAGCAGAAGACGGCATACGAGATGCTACCGTGACTGGAGTTCAGACGTGTGCTCTTCCGATC*T | NEB barcode Set 3-2 | sRNA-seq | * indicates phosphorothioate bond |
| NEB3-3 | CAAGCAGAAGACGGCATACGAGATGCTCATGTGACTGGAGTTCAGACGTGTGCTCTTCCGATC*T | NEB barcode Set 3-3 | sRNA-seq | * indicates phosphorothioate bond |
| NEB3-4 | CAAGCAGAAGACGGCATACGAGATCTTTTGGTGACTGGAGTTCAGACGTGTGCTCTTCCGATC*T | NEB barcode Set 3-4 | sRNA-seq | * indicates phosphorothioate bond |
| NEB3-5 | CAAGCAGAAGACGGCATACGAGATTAGTTGGTGACTGGAGTTCAGACGTGTGCTCTTCCGATC*T | NEB barcode Set 3-5 | sRNA-seq | * indicates phosphorothioate bond |
| NEB3-6 | CAAGCAGAAGACGGCATACGAGATATCGTGGTGACTGGAGTTCAGACGTGTGCTCTTCCGATC*T | NEB barcode Set 3-6 | sRNA-seq | * indicates phosphorothioate bond |
| NEB3-7 | CAAGCAGAAGACGGCATACGAGATTGAGTGGTGACTGGAGTTCAGACGTGTGCTCTTCCGATC*T | NEB barcode Set 3-7 | sRNA-seq | * indicates phosphorothioate bond |
| NEB3-8 | CAAGCAGAAGACGGCATACGAGATCGCCTGGTGACTGGAGTTCAGACGTGTGCTCTTCCGATC*T | NEB barcode Set 3-8 | sRNA-seq | * indicates phosphorothioate bond |
| NEB3-9 | CAAGCAGAAGACGGCATACGAGATGCCATGGTGACTGGAGTTCAGACGTGTGCTCTTCCGATC*T | NEB barcode Set 3-9 | sRNA-seq | * indicates phosphorothioate bond |
| NEB3-10 | CAAGCAGAAGACGGCATACGAGATAAAATGGTGACTGGAGTTCAGACGTGTGCTCTTCCGATC*T | NEB barcode Set 3-10 | sRNA-seq | * indicates phosphorothioate bond |
| NEB3-11 | CAAGCAGAAGACGGCATACGAGATTGTTGGGTGACTGGAGTTCAGACGTGTGCTCTTCCGATC*T | NEB barcode Set 3-11 | sRNA-seq | * indicates phosphorothioate bond |
| NEB3-12 | CAAGCAGAAGACGGCATACGAGATCGATTAGTGACTGGAGTTCAGACGTGTGCTCTTCCGATC*T | NEB barcode Set 3-12 | sRNA-seq | * indicates phosphorothioate bond |
| NEB4-1 | CAAGCAGAAGACGGCATACGAGATCCGGTGGTGACTGGAGTTCAGACGTGTGCTCTTCCGATC*T | NEB barcode Set 4-1 | sRNA-seq | * indicates phosphorothioate bond |
| NEB4-2 | CAAGCAGAAGACGGCATACGAGATATTCCGGTGACTGGAGTTCAGACGTGTGCTCTTCCGATC*T | NEB barcode Set 4-2 | sRNA-seq | * indicates phosphorothioate bond |
| NEB4-3 | CAAGCAGAAGACGGCATACGAGATAGCTAGGTGACTGGAGTTCAGACGTGTGCTCTTCCGATC*T | NEB barcode Set 4-3 | sRNA-seq | * indicates phosphorothioate bond |
| NEB4-4 | CAAGCAGAAGACGGCATACGAGATGTATAGGTGACTGGAGTTCAGACGTGTGCTCTTCCGATC*T | NEB barcode Set 4-4 | sRNA-seq | * indicates phosphorothioate bond |
| NEB4-5 | CAAGCAGAAGACGGCATACGAGATTGGATCACGTGACTGGAGTTCAGACGTGTGCTCTTCCGATC*T | NEB barcode Set 4-5 | sRNA-seq | * indicates phosphorothioate bond |
| NEB4-6 | CAAGCAGAAGACGGCATACGAGATGTCGTCGTGACTGGAGTTCAGACGTGTGCTCTTCCGATC*T | NEB barcode Set 4-6 | sRNA-seq | * indicates phosphorothioate bond |
| NEB4-7 | CAAGCAGAAGACGGCATACGAGATGCTGTAGTGACTGGAGTTCAGACGTGTGCTCTTCCGATC*T | NEB barcode Set 4-7 | sRNA-seq | * indicates phosphorothioate bond |
| NEB4-8 | CAAGCAGAAGACGGCATACGAGATATTATAGTGACTGGAGTTCAGACGTGTGCTCTTCCGATC*T | NEB barcode Set 4-8 | sRNA-seq | * indicates phosphorothioate bond |
| NEB4-9 | CAAGCAGAAGACGGCATACGAGATGAATGAGTGACTGGAGTTCAGACGTGTGCTCTTCCGATC*T | NEB barcode Set 4-9 | sRNA-seq | * indicates phosphorothioate bond |
| NJ412 | AATGATACGGCGACCACCGAGATCTACACGTTCAGAGTTCTACAGTCCG*A | NEB SR primer | sRNA-seq | * indicates phosphorothioate bond |
| NJ392 | /5Biosg/AAGCAGTGGTATCAACGCAGAGTACrGrG+G | Template switching oligo | NanoPARE and mRNA-seq | rN indicates RNA base; +N indicates LNA base; /5Biosg/ indicates 5' biotinylation. Used as described in (*12*) |
| NJ393 | AAGCAGTGGTATCAACGCAGAGTACTTTTTTTTTTTTTTTTTTTTTTTTTTTTTTVN | Anchored oligo-dT RT primer | NanoPARE and mRNA-seq | Used as described in (*12*) |
| NJ394 | AAGCAGTGGTATCAACGCAGAGT | ISPCR primers for preamplification | NanoPARE and mRNA-seq | Used as described in (*12*) |
| NJ396 | AATGATACGGCGACCACCGAGATCTACACTATCCTCTCTAGCAAGCAGTGGTATCAACGCAGAGTACGGG | 5' TSO enrichment - H503 | NanoPARE | Forward |
| NJ397 | AATGATACGGCGACCACCGAGATCTACACGTAAGGAGCTAGCAAGCAGTGGTATCAACGCAGAGTACGGG | 5' TSO enrichment - H505 | NanoPARE | Forward |
| NJ398 | AATGATACGGCGACCACCGAGATCTACACACTGCATACTAGCAAGCAGTGGTATCAACGCAGAGTACGGG | 5' TSO enrichment - H506 | NanoPARE | Forward |
| NJ399 | AATGATACGGCGACCACCGAGATCTACACTATCCTCTTCGTCGGCAGCGTC | Nextera Tn5.1 - H503 | mRNA-seq | Forward |
| NJ400 | AATGATACGGCGACCACCGAGATCTACACGTAAGGAGTCGTCGGCAGCGTC | Nextera Tn5.1 - H505 | mRNA-seq | Forward |
| NJ401 | AATGATACGGCGACCACCGAGATCTACACACTGCATATCGTCGGCAGCGTC | Nextera Tn5.1 - H506 | mRNA-seq | Forward |
| NJ402 | CAAGCAGAAGACGGCATACGAGATTCGCCTTAGTCTCGTGGGCTCGG | Nextera Tn5.2 - H701 | NanoPARE and mRNA-seq | Reverse |
| NJ403 | CAAGCAGAAGACGGCATACGAGATCTAGTACGGTCTCGTGGGCTCGG | Nextera Tn5.2 - H702 | NanoPARE and mRNA-seq | Reverse |
| NJ404 | CAAGCAGAAGACGGCATACGAGATTTCTGCCTGTCTCGTGGGCTCGG | Nextera Tn5.2 - H703 | NanoPARE and mRNA-seq | Reverse |
| NJ405 | CAAGCAGAAGACGGCATACGAGATAGGAGTCCGTCTCGTGGGCTCGG | Nextera Tn5.2 - H705 | NanoPARE and mRNA-seq | Reverse |
| NJ406 | CAAGCAGAAGACGGCATACGAGATCATGCCTAGTCTCGTGGGCTCGG | Nextera Tn5.2 - H706 | NanoPARE and mRNA-seq | Reverse |
| NJ395 | CTAGCAAGCAGTGGTATCAACGCAGAGTACGGG | Sequencing primer read 1 | NanoPARE and mRNA-seq | Alternative sequencing primers used in sequencing |
| NJ416 | CCCGTACTCTGCGTTGATACCACTGCTTGCTAG | Sequencing primer i5 index | NanoPARE and mRNA-seq | Alternative sequencing primers used in sequencing |

**Table S3. Eudicot genomic resources used in this study**
All available in phytozome version v12.1.6

| Genome version | cDNA files | CDS files |
|---|---|---|
| *Amaranthus hypochondriacus v1.0* | Acoerulea_322_v3.1.transcript.fa | Acoerulea_322_v3.1.cds.fa |
| *Aquilegia coerulea v3.1* | Ahalleri_264_v1.1.transcript.fa | Ahalleri_264_v1.1.cds.fa |
| *Arabidopsis halleri v1.1* | Ahypochondriacus_459_v2.1.transcript.fa | Ahypochondriacus_459_v2.1.cds.fa |
| *Arabidopsis lyrata v2.1* | Alyrata_384_v2.1.transcript.fa | Alyrata_384_v2.1.cds.fa |
| *Arabidopsis thaliana TAIR10* | Araport11_cDNA_no-repeats.no-whitespace.fa | Athaliana_447_Araport11.cds.fa |
| *Boechera stricta v1.2* | Bstricta_278_v1.2.transcript.fa | Bstricta_278_v1.2.cds.fa |
| *Brassica oleracea capitata v1.0* | Boleraceacapitata_446_v1.0.transcript.fa | Boleraceacapitata_446_v1.0.cds.fa |
| *Brassica rapa FPsc v1.3* | BrapaFPsc_277_v1.3.transcript.fa | BrapaFPsc_277_v1.3.cds.fa |
| *Capsella grandiflora v1.1* | Cgrandiflora_266_v1.1.transcript.fa | Cgrandiflora_266_v1.1.cds.fa |
| *Capsella rubella v1.0* | Crubella_474_v1.1.transcript.fa | Crubella_474_v1.1.cds.fa |
| *Carica papaya ASGPBv0.4* | Cpapaya_113_ASGPBv0.4.transcript.fa | Cpapaya_113_ASGPBv0.4.cds.fa |
| *Citrus clementina v1.0* | Cclementina_182_v1.0.transcript.fa | Cclementina_182_v1.0.cds.fa |
| *Citrus sinensis v1.1* | Csinensis_154_v1.1.transcript.fa | Csinensis_154_v1.1.cds.fa |
| *Cucumis sativus v1.0* | Csativus_122_v1.0.transcript.fa | Csativus_122_cds.fa |
| *Daucus carota v2.0* | Dcarota_388_v2.0.transcript.fa | Dcarota_388_v2.0.cds.fa |
| *Eucalyptus grandis v2.0* | Egrandis_297_v2.0.transcript.fa | Egrandis_297_v2.0.cds.fa |
| *Eutrema salsugineum v1.0* | Esalsugineum_173_v1.0.transcript.fa | Esalsugineum_173_v1.0.cds.fa |
| *Fragaria vesca v1.1* | Fvesca_501_v2.0.a2.transcript.fa | Fvesca_501_v2.0.a2.cds.fa |
| *Glycine max Wm82.a2.v1* | Gmax_275_Wm82.a2.v1.transcript.fa | Gmax_275_Wm82.a2.v1.cds.fa |
| *Gossypium raimondii v2.1* | Graimondii_221_v2.1.transcript.fa | Graimondii_221_v2.1.cds.fa |
| *Kalanchoe fedtschenkoi v1.1* | Klaxiflora_309_v1.1.transcript.fa | Klaxiflora_309_v1.1.cds.fa |
| *Kalanchoe laxiflora v1.1* | Kfedtschenkoi_382_v1.1.transcript.fa | Kfedtschenkoi_382_v1.1.cds.fa |
| *Linum usitatissimum v1.0* | Lusitatissimum_200_v1.0.transcript.fa | Lusitatissimum_200_v1.0.cds.fa |
| *Malus domestica v1.0* | Mdomestica_491_v1.1.transcript.fa | Mdomestica_491_v1.1.cds.fa |
| *Manihot esculenta v6.1* | Mesculenta_305_v6.1.transcript.fa | Mesculenta_305_v6.1.cds.fa |
| *Medicago truncatula Mt4.0v1* | Mtruncatula_285_Mt4.0v1.transcript.fa | Mtruncatula_285_Mt4.0v1.cds.fa |
| *Mimulus guttatus v2.0* | Mguttatus_256_v2.0.transcript.fa | Mguttatus_256_v2.0.cds.fa |
| *Phaseolus vulgaris v2.1* | Ppersica_298_v2.1.transcript.fa | Ppersica_298_v2.1.cds.fa |
| *Populus trichocarpa v3.0* | Ptrichocarpa_444_v3.1.transcript.fa | Ptrichocarpa_444_v3.1.cds.fa |
| *Prunus persica v2.1* | Pvulgaris_442_v2.1.transcript.fa | Pvulgaris_442_v2.1.cds.fa |
| *Ricinus communis v0.1* | Rcommunis_119_v0.1.transcript.fa | Rcommunis_119_v0.1.cds.fa |
| *Salix purpurea v1.0* | Spurpurea_289_v1.0.transcript.fa | Spurpurea_289_v1.0.cds.fa |
| *Solanum tuberosum v4.03* | Stuberosum_448_v4.03.transcript.fa | Stuberosum_448_v4.03.cds.fa |
| *Theobroma cacao v1.1* | Tcacao_233_v1.1.transcript.fa | Tcacao_233_v1.1.cds.fa |
| *Trifolium pratense v2* | Tpratense_385_v2.transcript.fa | Tpratense_385_v2.cds.fa |
| *Vitis vinifera Genoscope.12X* | Vvinifera_457_v2.1.transcript.fa | Vvinifera_457_v2.1.cds.fa |

**Data S1 (separate file - FASTA format). Alignment of TrnL-F sequences from *Cuscuta***

**Data S2 (separate file - tab-delimited text format). Comprehensive list of HI-sRNAs discovered in this study**
